# Complete blood count with differential: An effective diagnostic for IBS subtype in the context of BMI?

**DOI:** 10.1101/608208

**Authors:** JM Robinson, CM Boulineaux, KR Butler, PV Joseph, MT Murray, SN Pocock, LB Sherwin, SC Turkington, KR Weaver, WA Henderson

**Author notes:** Correspondence should be addressed to Wendy A. Henderson, PhD, Investigator & Chief, Digestive Disorder Unit, Division of Intramural Research, NINR, National Institutes of Health, Department of Health and Human Services, 10 Center Drive, 2-1341 Bethesda, MD 20892.

## Abstract

The Complete Blood Count with Differential hematological assay is a mainstay diagnostic for point-of-care clinical diagnoses for a spectrum of diseases including infection, inflammation, anemia, and leukemia, and CBC-D profiles are under investigation as early prognostic biomarkers for leukemias and other diseases. Chronic abdominal pain (CAP) and irritable bowel syndrome (IBS) are prevalent gastrointestinal disorders in the United States, with obesity among the most common comorbidities. Often, IBS-like gastrointestinal (GI) symptoms persist after resolution of known inflammation and/or enteropathogenic infection, and current literature contains significant discussion of the extent to which IBS is within the biological spectrum of inflammatory disease. Obesity is also associated with generalized signatures of inflammation and may confound accurate diagnoses. We performed ANOVA, multiple means comparisons, statistical analyses of CBC data from our "Brain-Gut Natural History" (BGNH) clinical cohort, with additional ELISA assays for lipopolysaccharide binding protein (LBP), IL-10, cortisol, and ACTH, signatures of immune-inflammatory response and Hypothalamic-Pituitary-Adrenal (HPA) axis activity, respectively. BGNH cohort includes healthy and overweight individuals diagnosed with IBS diarrhea-(IBS-D) and constipation-predominant (IBS-C) subtypes. We identified several potentially significant markers for IBS-D and IBS-C, notably IL-10, mean platelet volume (MPV), with LBP and monocyte percent also showing some statistical significance. Weight also showed significant results for overweight vs. normal weight, regardless of IBS subtype, particularly for Cortisol. CBC-D predictive profiles for IBS subtype and weight were identified using discriminant functions analysis and show that predictivity of marker profiles have poor performance relative to their normal weight subsets. Further refinement of this analysis will be performed utilizing increased sample size, additional molecular profiles, and enhanced statistical analysis.

## Introduction

Chronic abdominal pain (CAP) and irritable bowel syndrome (IBS), with diarrhea-(IBS-D) and constipation-predominant (IBS-C) subtypes, are among the most common gastrointestinal disorders in Western nations, however identification of mechanistic associations have proven difficult given the heterogeneity of possible underlying mechanisms [1–4]. Gastrointestinal pain and diarrhea are also symptoms of enteropathogenic infection and inflammatory bowel disease (IBD). Significant discussion appears in recent literature that IBS (or a proportion of IBS cases) may represent a mild, subclinical extreme of the IBD ‘spectrum’, which many authors have supported and provided evidence for [5–9]. Complete blood count with differential (CBC-D) hematological test is the routine multiparameter test providing clinicians with identification of hematological perturbation and infectious immune responses [10]. CBC-D has recently been the focus of discovery of prognostic biomarkers in diseases including T-cell lymphoma [11], peripheral arterial disease [12], and many others.

MVP, ESR, and CRP parameters of CBC-D have each been reported as statistically-supported indicators of IBD. [13, 14] CBC monocyte count is also indicator of inflammation [15]. CD14, LPS, Cortisol, and IL-10 are reported indicators of immune response or inflammatory suppression, respectively. sCD14 is an indicator of monocyte activation [16], and both sCD14 and LPS are play roles in innate immunity via interactions with the TLR4 receptor [17]. IL-10 is an important anti-inflammatory cytokine associated with IBD [18, 19], with significant application as anti-inflammatory therapeutic compound. [20] Regulation of macrophage activity by IL-10 appears particularly important in intestinal inflammation. IL-10 is produced by macrophages in response to LPS exposure [21]. IL-10 activates its own expression in macrophages via Stat3 transcription factor [22]. Mutations in IL-10 or IL-10 receptors (IL-10R) result in uncontrolled inflammation, and absence of IL-10R results in spontaneous inflammation. [23, 24] IL-10 exposure results in differentiation of inflammation-regulatory macrophage subtypes. [25, 26] IL-10 also affects T-cell activity in colitis. [27, 28] The combination of parameters available in CBC-D and ELISA results therefore provide not only a potential diagnostic profile, but insight into pathophysiology as well.

We sought to identify potential signatures of immune/inflammatory responses associated with weight and IBS-subtype using CBC-D and other ELISA data from our clinical cohort “Brain-Gut Interactions in Overweight and Normal Weight Patients with Chronic Abdominal Pain” (09-NR-0064) natural history protocol. We performed exploratory statistical analyses using ANOVA and multiple means comparisons of clinical CBC-D data with cortisol and ACTH, as well as ELISA tests for IL-10, serum CD14 and lipopolysaccharide binding protein (LBP). Finally, we applied the discriminant functions analysis (Fischer’s discriminant) to provide additional quantification for how differential CBC-D/ELISA signal could be predictive for weight and IBS subtype.

## Methods

### Clinical Protocol (NCT00824941)

Participants in this study gave informed consent at the National Institutes of Health’s Hatfield Clinical Research Center under our IRB-approved “Brain-Gut Interactions in Overweight and Normal Weight Patients with Chronic Abdominal Pain” (09-NR-0064) natural history clinical protocol. Participants must be between the ages of 13-45 years, and, if female, have regular menstrual cycles. Participants were evaluated for CAP using the Gastrointestinal Pain Pointer (GIPP) tool [29], and were diagnosed for IBS and IBS-subtype under the Rome III Criteria, [2] with subtypes classified as diarrhea-predominant IBS (IBS-D) and constipation-predominant IBS (IBS-C). The participants diagnosed with mixed-type IBS (IBS-M) or un-subtyped IBS (IBS-U) were excluded from this analysis due to small sample size. Participants were classified by body mass index (BMI) into “normal” (BMI < 25) and “overweight” (BMI ≥ 25) groups. Measurements of height and weight (duplicate) were averaged to calculate patients’ BMI. Table 1A shows summary statistics for the cohort’s phenotypic distribution.

**TABLE 1A.**
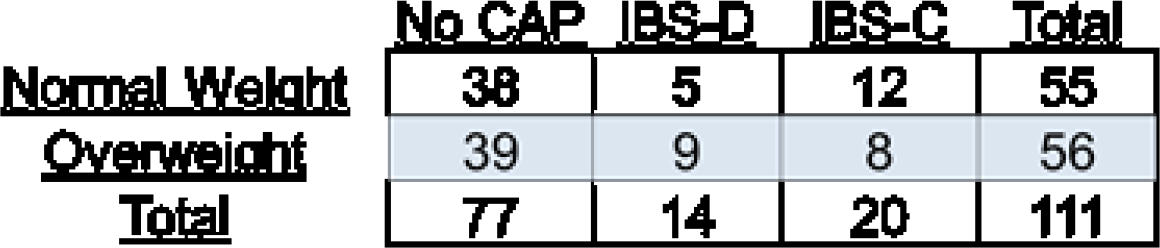
Participant phenotype matrix (No CAP = Healthy Control (HC))

|  | No CAP | IBS-D | IBS-C | Total |
| --- | --- | --- | --- | --- |
| <u>Normal Weight</u> | 38 | 5 | 12 | 55 |
| <u>Overweight</u> | 39 | 9 | 8 | 56 |
| <u>Total</u> | 77 | 14 | 20 | 111 |

**TABLE 1B.** Results of Oneway Means Comparisons Tests. (W0=Normal Weight, W1=Overweight) (* p≤.05; ** p≤.01; *** p≤.001; **** p≤.0001)

| Cohort subsets: | Total cohort | W0 | IBS-D | W1 | HC | Total cohort | IBS-C | W0 | W1 | Total cohort |
| --- | --- | --- | --- | --- | --- | --- | --- | --- | --- | --- |
| Means comparison: | HC v IBSD | HC v IBSD | W0 v W1 | HC v IBSD | W0 v W1 | W0 v W1 | W0 v W1 | HC v IBSC | HC v IBSC | HC v IBSC |
| CBC datapoint: |  |  |  |  |  |  |  |  |  |  |
| Monocytes Percent | * |  |  |  |  |  |  |  |  |  |
| IgM | * |  |  |  |  |  |  |  |  |  |
| LBP | ** | * |  |  |  |  |  |  |  |  |
| MPV | ** | *** | ** |  |  |  |  |  |  |  |
| IL10 |  | ** | ** |  |  |  |  |  |  |  |
| Eosinophil_PCT |  | * | * |  |  |  |  |  |  |  |
| Cortisol |  | * | *** |  | * | **** | ** |  |  |  |
| CRP |  |  |  |  | ** | ** | ** |  |  |  |
| ESR |  |  |  |  | * | ** | ** | ** |  |  |
| MCH |  |  |  |  |  | * | ** | * | * |  |
| Lymphocytes |  |  |  |  | * | * |  |  |  |  |
| Platelet Count |  |  |  |  | * | * |  |  |  |  |
| IgE |  |  |  |  |  |  | * |  |  |  |
| sCD14 |  |  |  |  |  |  | * | * |  |  |
| IgA |  |  |  |  |  |  |  | * |  |  |
| Lymphocytes Percent |  |  |  |  |  |  |  |  |  | * |
| Neutrophils Percent |  |  |  |  |  |  |  |  |  | * |

### Complete Blood Count with Differential (CBC-D)

In the protocol’s clinical visits, fasting blood was collected in the morning from participants. Standard blood panels are submitted to NIH-CRC Department of Laboratory Medicine (DLM) for CBC-D, serum cortisol and ACTH. The full matrix used in this study is found in Sup. Data 1, summarized in Table 1A.

CBC-D data was compiled and curated from the NIH CRC Biomedical Translational Research Information System (BTRIS) [30] and Clinical Research Information System (CRIS) databases (https://cris.cc.nih.gov/index.html). CBC data provides absolute counts by volume for overall WBCs, and absolute and relative counts for monocytes, lymphocytes, neutrophils, and eosinophils. WBC differential counts were compiled including absolute counts and relative abundance of each cell category. Absolute counts (x10^6^ cells/mL) for overall WBCs, neutrophils, monocytes, lymphocytes, basophils, and eosinophils and relative abundance (percentage of overall WBC count) for neutrophils, monocytes, lymphocytes, basophils, and eosinophils were evaluated. Serum immunoglobulin tests were also analyzed for IgA, IgG, IgM, and IgE. Other CBC-D data collected for each participant included MPV, MCH, HCT, ESR, LDH, serum cortisol, ACTH, CRP, platelet counts, and red blood cell counts.

### ELISA quantification of serum protein biomarkers

Research blood was collected for concurrent processing of serum and plasma using standard laboratory protocols. ELISAs were performed according to the manufacturer’s instructions for IL-10 (Quantikine^®^HS ELISA Human IL-10 Immunoassay, R&D Systems, Inc., Cat# HS100B), sCD14 (Quantikine^®^ ELISA Human sCD14 Immunoassay, R&D Systems, Inc., Cat# DC140), I-FABP (Human I-FABP ELISA Kit, Hycult biotech, Cat# HK406), and LBP (Human Lipopolysaccharide Binding Protein ELISA Kit, Cell Sciences, Inc., Cat# CKH113).

### Stool culture tests

Stool culture tests for enteropathogenic microbes are a standard diagnostic for enteropathogenic infection in the intestine, which also produce diarrheal and chronic pain symptomology. [31] In our cohort, diagnostic testing was performed using enteric bacterial culture, ova and parasite exam, anti-*Helicobacter pylori* IgG antibody detection, polymerase chain reaction assay for *Clostridium difficile*, and/or multiplex gastrointestinal pathogen panel testing, as indicated. (Results provided in Sup. Data 1)

### Statistical Analyses

#### Student’s T-tests and means comparisons

ANOVA and multiple means comparison tests were performed using JMP13 software [32] for each predictive variable (weight: 0 = normal, 1 = overweight; IBS subtype 0 = healthy, 1 = IBS-D, 2 = IBS-C) (SAS-JMP 13 manual). Complete statistical reports were generated for 1-and 2-tailed tests on ANOVA and multiple means comparisons results:

1. Normal and Overweight subsets each tested for HC vs IBS-D and HC vs IBS-C (Sup. Data 2: Stats Subtype subset by Weight)
2. HC, IBS-D, and IBS-C subsets each tested for Normal vs. Overweight (Sup. Data 3: Stats Weight subset by Subtype)

Box-and-Whisker plots for multiple comparisons were produced with R package ggplot2. P-values are reported for 1-tailed t-tests on multiple means comparisons that showed statistically significant results (Fig. 1, Table 1B).

**Figure 1.**
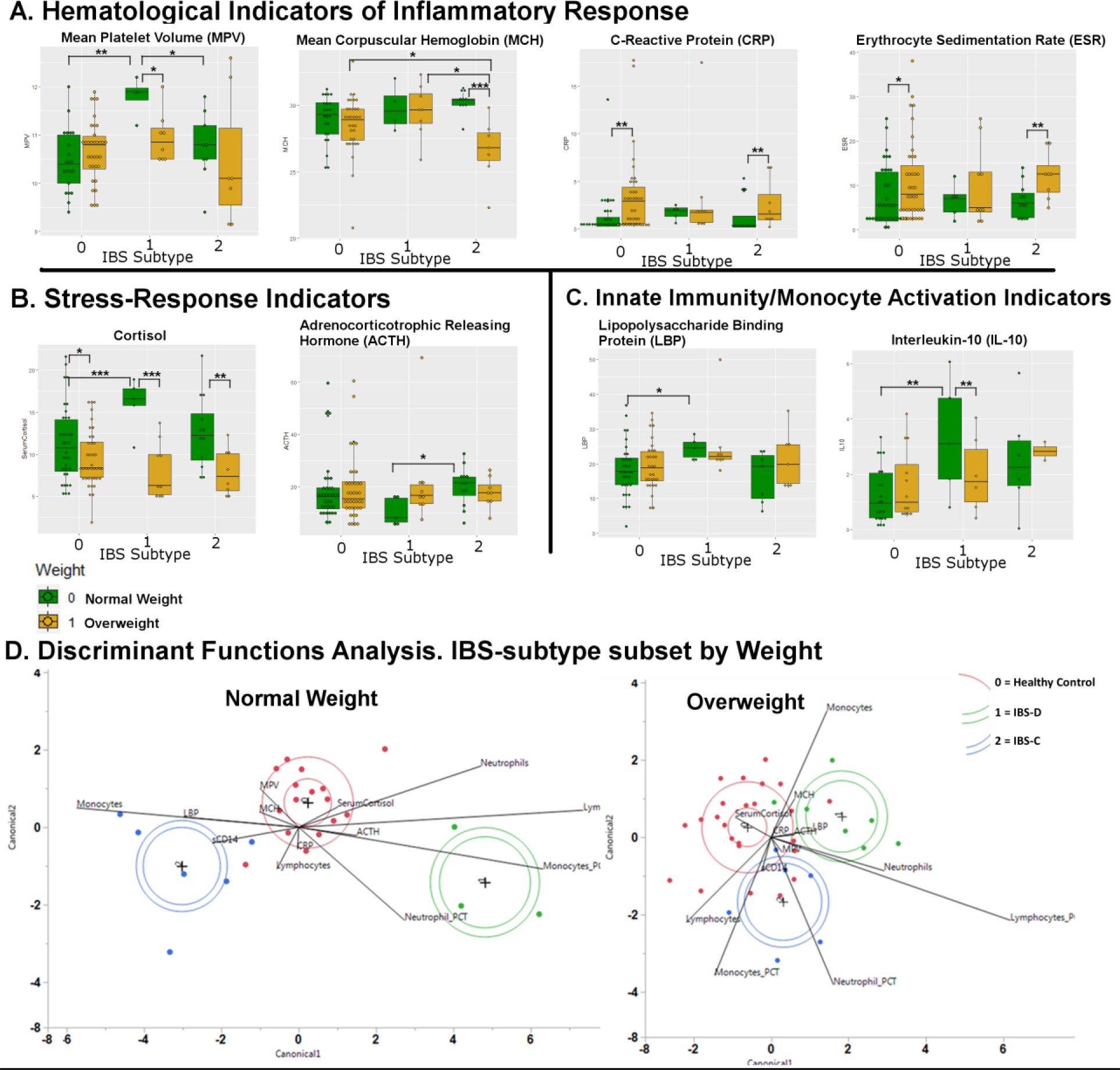
Multiple Means and Discriminant Analysis Results of Complete Blood Count Data Indicate Differential Profiles Associated with IBS-subtype and Weight Phenotypes. **A-C.** Blood indicators show significant weigh-associated differential readouts. Each of MPV, cortisol, LPB, and IL-10 are significantly higher for IBS-D/Normal Weight vs. IBS-D/Overweight. **D.** Discriminant functions analysis using these cohort data points, subset by Normal Weight and Overweight, shows diminished predictivity for IBS-subtype in Overweight vs. Normal Weight.

#### Discriminant Functions Analysis

Discriminant functions analysis is a multivariate, supervised grouping method (a generalization of Fischer’s linear discriminant) used to predict categorical classification variables based on multiple continuous variables. [33] Data were subset into normal and overweight groups, and discriminant analysis was used on each subset using JMP13. [34] Complete results including canonical plots are provided (Sup. Data 4 (normal weight subset), Sup. 5 (overweight subset)).

## Results and Discussion

### CBC and serum ELISA panels demonstrate a signature of immune/inflammatory suppression associated with the IBS-D/Normal Weight cohort subset

(Table 1B, Fig. 1A-C, Fig. 2A,B,C, Sup. Data 1-3). Significant IBS-D associated increases in MPV, LBP, cortisol, and IL-10 are observed, suggesting innate immune activation and inflammatory response (represented by increased LBP and MPV, respectively) with a corresponding signature of immunosuppressive response (cortisol, IL-10) (Fig. 1A,B, Table 1B). Increased MPV is associated with inflammatory disorders, including IBD [13, 14]. IL-10 is a key inflammatory suppression cytokine associated with IBD and is an important developmental switch in monocyte -> macrophage differentiation, acting through the Stat3 transcription factor, its known effector [22, 24–26]. IL-10 itself is produced by Monocytes, regulatory B-cells and other immune cells in response to LPS [21]. In our data, IBS-D is associated with a weakly significant decrease in Monocyte percentage based on the CBC-D test, while absolute Monocyte counts, contrary to current IBD literature have been reported be increased [15].

**Figure 2.**
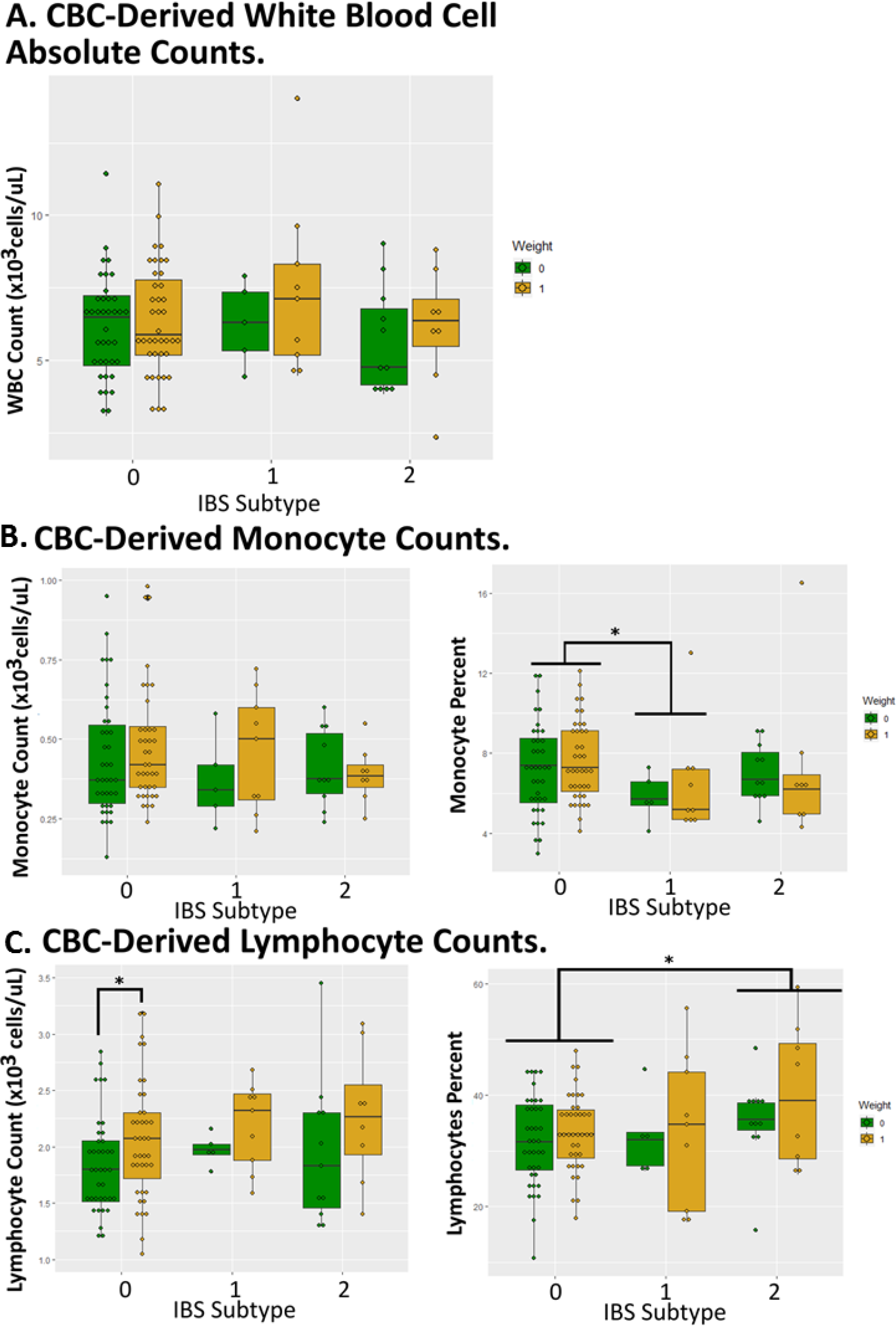
White Blood Cell Counts Show Perturbations Associated with IBS-subtype and Weight Phenotypes. **A.** CBC/D-derived absolute WBC counts between weight and IBS-subtype groups were not significantly different. **B.** CBC-D-derived Monocyte percentage showed significant decrease in IBS-D relative to healthy control group (C). **C.** CBC-D-derived Lymphocyte absolute counts showed significant increase in overweight relative to normal weight individuals, and Lymphocyte percentage was significantly increased in IBS-C vs Healthy Controls.

This pattern, observed in our IBS-D/Normal Weight cohort subset, is consistent with a suppressed immune/inflammatory response. Stool culture testing was performed on a subset of our cohort, however positive stool culture results were not significantly correlated with IBS symptoms; in fact, in many cases, IBS subjects had negative stool cultures, while Healthy Controls showed positive results (Sup. Data 1). Many researchers have supported the possibility that IBS is representative of a ‘low-grade inflammation’ along a spectrum of inflammatory disease [6–9]. Individuals with resolved enteropathogenic infection or IBD in remission often report persistent visceral hypersensitivity with IBS-like symptomology (diarrhea or constipation) [35–37].

### CBC and serum ELISA panels identify overweight-associated effect across IBS-subtypes, and additional inflammatory signatures in IBS-C/overweight subsets

(Table 1B, Fig. 1A-C, Sup. Data) Cortisol levels are suppressed in <u>all</u> overweight subsets, across IBS subtypes including in the Healthy Controls subset, though effect size is particularly strong for IBS-D Normal vs Overweight given the significant elevation of cortisol in the IBS-D/Normal Weight subset. Compared with other cohort subsets, the IBS-D/Overweight subset shows suppressed differential for CRP and ESR, which are significantly elevated in Healthy Control/Overweight and IBS-C/Overweight subsets. IL-10 is not significantly altered in the IBS-D/Overweight subset as it is in IBS-D/Normal Weight subset. Strikingly, among the overweight subset there are *no significant differences in any parameter* between Healthy Controls and IBS-D, with parameters associated with innate immune/inflammatory suppression profile seen in IBS-D/Normal Weight subset being absent in the IBS-D/Overweight subset. Interestingly, the MCH parameter is significantly lowered in IBS-C/Overweight subset only. Low MCH is indicative of anemia (iron deficiency), and is an IBD marker of anemia in its own right [38]. Given these results, IBS-C appears to have less significant associations than IBS-D, with perturbations often associated with weight, rather than with the IBS-C subtype.

Obesity and Metabolic Syndrome are especially common co-morbidities for GI disorders, and often co-occur with IBS symptomology [39], and with elevated pain phenotypes in general [40]. Inflammation is associated with obesity, as well as with associated perturbations of the gut microbiota [41, 42]. Overweight subsets from our Healthy Control/Overweight and IBS-C/Overweight subsets are associated with elevated CRP and ESR, both of which are standard diagnostic markers for inflammation associated with IBD [43, 44]. In our cohort, we find CRP and ESR elevation more associated with Healthy Control/Overweight and IBS-C/Overweight than with IBS symptomology, and these may therefore be indicative of inflammatory signature of weight-associated inflammation rather than of IBS-associated inflammation.

### Monocytes and other immune cells exhibit small but significant perturbations in relative abundance based on CBC data

(Table 1B, Fig. 2). Monocyte percent is significantly decreased in CBC-based statistical comparisons between IBS-D vs. HC cohort subsets (Table 1B, Supplementary Data), although these are still within the clinical “normal range” of ~400–1000 cells/ul, or 2–10% of the relative WBC count.

Observations from CBC-D support a suppression of monocyte abundance in the PBMC population associated with IBS-D and overweight phenotypes. Neutrophils are shown as having a decreased relative abundance in the expression assay but have insignificant CBC results.

Additional small yet significant changes are seen in immune cell count and immunoglobulin counts associated with IBS subtype and weight subsets. Changes in percentage counts from CBC for Macrophage, Neutrophil, Lymphocyte, and Eosinophils appear variously associated with IBS-D and IBS-C (Table 1, Fig. 2). While significant, most of individual perturbations are within the normal diagnostic ranges for their respective counts.

### Discriminant functions analysis of CBC data shows diminished predictivity for IBS subtype in OW compared with NW subsets

(Fig. 1D, Supplementary Data 4,5). We tested predictivity of multivariate parameter profiles, subsetting the cohort by weight (normal and overweight) and applied discriminant functions analysis to a multivariate profile of significantly affected variables. Confirming statistical results described above, predictivity (a function of the multi-parameter *eigenvalue* derivative) was good for IBS-D and IBS-C individuals from the normal weight subset, with good separation of IBS-subtypes encompassed within the first canonical coordinate. However, predictivity was poorer between IBS subtypes in the overweight subset, with small distances and overlapping predictions within canonical coordinates 1 and 2. This confirms predictivity for IBS-D with the variable set is significantly degraded in the overweight cohort participants.

## Conclusion

In IBS-D, previous research demonstrated an association with signatures of immune/inflammatory responses, with significant discussion in the literature as to what extent IBS symptoms fall along the continuum of IBD. [45] Here we report perturbation of many parameters associated with IBS-D which have also been reported associated with IBD, including the anti-inflammatory cytokine IL-10 and modulation of monocyte/macrophage activation [18, 19, 21–23, 25, 26, 46–48].

We observe small but significant perturbations associated with IBS subtype and weight corresponding with many previously reported signatures of inflammation and innate immune activation. Weight plays a significant role as a covariate in each of the separate data-type comparisons and is particularly significant in obscuring the diagnostic value of CBC parameters. Analysis of much larger cohorts, and inclusion of additional genetic, medical history, and psychological data has been a successful approach [49].

## Supplementary Materials

1. SupplementaryData_1_Cohort_Phenotypes.xlsx – All data for cohort variables.
2. SupplementaryData_2_Subtype_BY_Weight.pdf – JMP13 Complete Statistical Results for IBS-subtype subset by Weight Class.
3. Supplementary_Data_3_Weight_BY_Subtype.pdf – JMP13 Complete Statistical Results for Weight Class subset by IBS-subtype.
4. Supplementary_Data_4_Discriminant_W0.htm – JMP13 HTML-formatted results for Discriminant Functions Analysis for Normal Weight subset.
5. Supplementary_Data_5_Discriminant_W1.htm – JMP13 HTML-formatted results for Discriminant Functions Analysis for Overweight subset.

## Supporting information

Supplementary Data 1. Phenotypes

Supplementary Data 2. IBS by Weight

Supplementary Data 3. Weight by IBS

Supplementary Data 4. Discriminant W0

Supplementary Data 5. Discriminant W1

## Glossary of Abbreviations

ACTH: adrenocorticotropic hormone
BGNH: Brain-Gut Natural History protocol
BTRIS: Biomedical Translational Research Information System
CAP: chronic abdominal pain
CBC-D: complete blood count with differential
CRIS: Clinical Research Information System
CRP: c-reactive protein
ESR: erythrocyte sedimentation rate
HC: healthy control
HCT: hematocrit
IBS: irritable bowel syndrome
IBS-C: irritable bowel syndrome, constipation-predominant
IBS-D: irritable bowel syndrome, diarrhea-predominant
IL-10: interleukin-10
LBP: lipopolysaccharide binding protein
LDH: lactate dehydrogenase
MCH: mean corpuscular hemoglobin
MPV: mean platelet volume
NW: normal weight
OW: overweight
PBMC: peripheral blood mononuclear cells
WBC: white blood cell

## Author Contributions

**Robinson** conceived this analysis as an Associate Investigator on the clinical protocol. Robinson mentored and trained Boulineaux in statistical analysis, performed statistical analyses in JMP and R, wrote, revised, and edited the text, and produced the figures. **Boulineaux** compiled and curated subjects’ clinical data from the NIH CRIS and BTRIS databases, performed statistical analyses in JMP, contributed to and edited the text. **Turkington** edited the text. **Joseph** and **Sherwin** performed clinical phenotyping for determination BMI and IBS-subtype diagnosis and edited the text. **Pocock**, **Murray**, and **Butler** compiled and curated stool culture test results. **Henderson** conceived and manages the clinical protocol, provided mentoring and supervision of the project, contributed to and edited the text.

## Conflicts of Interest

All authors declare no conflicts of interest.

## Acknowledgements

Research reported in this publication was supported by the U.S. Department of Health and Human Services, National Institutes of Health, National Institute of Nursing Research, Division of Intramural Research to Wendy A. Henderson, 1Z1ANR000018-01-8; Intramural Research Training Awards to Jeffrey M. Robinson, Christina M. Boulineaux, Simon C. Turkington, Paule V. Joseph, LeeAnne B. Sherwin, Kristen R. Weaver, Kierra R. Butler, Megan T. Murray. Authors thank Dr. Margaret Heitkemper for providing comments on previous drafts. Authors acknowledge NIH Clinical Research Center, Department of Laboratory Medicine (DLM), for contributing CBC data for analysis, and NIH Library for providing computational and software resources for statistical analysis.

## References

1. Drossman, D.A., et al., U.S. householder survey of functional gastrointestinal disorders. Prevalence, sociodemography, and health impact. Dig Dis Sci, 1993. 38(9): p. 1569–80.

2. Ford, A.C., et al., Validation of the Rome III criteria for the diagnosis of irritable bowel syndrome in secondary care. Gastroenterology, 2013. 145(6): p. 1262–70 e1.

3. Dang, J., et al., Systematic review of diagnostic criteria for IBS demonstrates poor validity and utilization of Rome III. Neurogastroenterol Motil, 2012. 24(9): p. 853–e397.

4. Manning, A.P., et al., Towards positive diagnosis of the irritable bowel. Br Med J, 1978. 2(6138): p. 653–4.

5. Sinagra, E., et al., Inflammation in irritable bowel syndrome: Myth or new treatment target? World J Gastroenterol, 2016. 22(7): p. 2242–55.

6. Bercik, P., E.F. Verdu, and S.M. Collins, Is irritable bowel syndrome a low-grade inflammatory bowel disease? Gastroenterol Clin North Am, 2005. 34(2): p. 235–45, vi-vii.

7. Abdul Rani, R., R.A. Raja Ali, and Y.Y. Lee, Irritable bowel syndrome and inflammatory bowel disease overlap syndrome: pieces of the puzzle are falling into place. Intest Res, 2016. 14(4): p. 297–304.

8. Akiho, H., E. Ihara, and K. Nakamura, Low-grade inflammation plays a pivotal role in gastrointestinal dysfunction in irritable bowel syndrome. World J Gastrointest Pathophysiol, 2010. 1(3): p. 97–105.

9. De Giorgio, R. and G. Barbara, Is irritable bowel syndrome an inflammatory disorder? Current Gastroenterology Reports, 2008. 10(4): p. 385–390.

10. N, V., G. Ss, and B. S, Basic examination of blood and bone marrow, in Henry’s Clinical Diagnosis and Management by Laborator Methods, Elsevier, Editor. 2017: St Louis, MO.

11. Feng, X., et al., Complete Blood Count Score Model Integrating Reduced Lymphocyte-Monocyte Ratio, Elevated Neutrophil-Lymphocyte Ratio, and Elevated Platelet-Lymphocyte Ratio Predicts Inferior Clinical Outcomes in Adult T-Lymphoblastic Lymphoma. Oncologist, 2019.

12. Velioglu, Y. and A. Yuksel, Complete blood count parameters in peripheral arterial disease. Aging Male, 2019: p. 1–5.

13. Derkacz, A., P. Olczyk, and K. Komosinska-Vassev, Diagnostic Markers for Nonspecific Inflammatory Bowel Diseases. Dis Markers, 2018. 2018: p. 7451946.

14. Liu, S., et al., Mean platelet volume: a controversial marker of disease activity in Crohn’s disease. Eur J Med Res, 2012. 17(1): p. 27.

15. Mee, A.S., J. Berney, and D.P. Jewell, Monocytes in inflammatory bowel disease: absolute monocyte counts. J Clin Pathol, 1980. 33(10): p. 917–20.

16. Cl, S., et al., Soluble CD14 is a nonspecific marker of monocyte activation. AIDS, 2015. 29(10).

17. Rl, K. and T. PA, Modulatory effects of sCD14 and LBP on LPS-host cell interactions. J Endotoxin Res, 2005. 11(4).

18. Trifunovic, J., et al., Pathologic patterns of interleukin 10 expression--a review. Biochem Med (Zagreb), 2015. 25(1): p. 36–48.

19. Kole, A. and K. Maloy, Control of intestinal inflammation by interleukin-10. 2005.

20. Fioranelli, M. and R. Grazia, Twenty-five years of studies and trials for the therapeutic application of IL-10 immunomodulating properties. From high doses administration to low dose medicine new paradigm. Journal of Integrative Cardiology, 2014. 1(1): p. 2–6.

21. Poitevin, S., et al., Monocyte IL-10 produced in response to lipopolysaccharide modulates thrombin generation by inhibiting tissue factor expression and release of active tissue factor-bound micoparticles. Thrombosis and haemostasis, 2007 97(4): p. 598–607.

22. Staples, K.J., et al., IL-10 induces IL-10 in primary human monocyte-derived macrophages via the transcription factor Stat3. J Immunol, 2007. 178(8): p. 4779–85.

23. Zigmond, E., et al., Macrophage-Restricted Interleukin-10 Receptor Deficiency, but Not IL-10 Deficiency, Causes Severe Spontaneous Colitis. Immunity, 2014. 40(5): p. 720–733.

24. Engelhardt, K.R. and B. Grimbacher, IL-10 in humans: lessons from the gut, IL-10/IL-10 receptor deficiencies, and IL-10 polymorphisms. Curr Top Microbiol Immunol, 2014. 380: p. 1–18.

25. Nguyen, H.H., et al., IL-10 Acts As a Developmental Switch Guiding Monocyte Differentiation to Macrophages during a Murine Peritoneal Infection. Journal of Immunology, 2012. 189(6): p. 3112–3120.

26. Prasse, A., et al., IL-10-producing monocytes differentiate to alternatively activated macrophages and are increased in atopic patients. J Allergy Clin Immunol, 2007. 119(2): p. 464–71.

27. Annacker, O., et al., Interleukin-10 in the regulation of T cell-induced colitis. J Autoimmun, 2003. 20(4): p. 277–9.

28. Berg, D.J., et al., Enterocolitis and colon cancer in interleukin-10-deficient mice are associated with aberrant cytokine production and CD4(+) TH1-like responses. J Clin Invest, 1996. 98(4): p. 1010–20.

29. Henderson, W. A., et al., The Gastrointestinal Pain Pointer: A Valid and Innovative Method to Assess Gastrointestinal Symptoms. Gastroenterol Nurs, 2017. 40(5): p. 357–363.

30. Cimino, J.J., et al., The National Institutes of Health’s Biomedical Translational Research Information System (BTRIS): design, contents, functionality and experience to date. J Biomed Inform, 2014. 52: p. 11–27.

31. Humphries, R.M. and A.J. Linscott, Laboratory diagnosis of bacterial gastroenteritis. Clin Microbiol Rev, 2015. 28(1): p. 3–31.

32. Inc., S.I., JMP 13 Basic Analysis. 2017, Cary, NC: SAS Insititute Inc.

33. Hardle, W. and L. Simar, Applied Multivariate Statistical Analysis. 2 ed. 2007, Berlin: Springer.

34. Inc., S.I., JMP 13 Multivariate Methods, Second Edition. 2 ed. 2917: SAS Institute Inc.

35. Spiller, R. and G. Major, IBS and IBD - separate entities or on a spectrum? Nat Rev Gastroenterol Hepatol, 2016. 13(10): p. 613–21.

36. Grover, M., H. Herfarth, and D.A. Drossman, The functional-organic dichotomy: postinfectious irritable bowel syndrome and inflammatory bowel disease-irritable bowel syndrome. Clin Gastroenterol Hepatol, 2009. 7(1): p. 48–53.

37. Burgell, R.E., A.K. Asthana, and P.R. Gibson, Irritable bowel syndrome in quiescent inflammatory bowel disease: a review. Minerva Gastroenterol Dietol., 2015. 61(4): p. 201–13.

38. Kaitha, S., M. Bashir, and T. Ali, Iron deficiency anemia in inflammatory bowel disease. World J Gastrointest Pathophysiol, 2015. 6(3): p. 62–72.

39. Nam, S.Y., Obesity-Related Digestive Diseases and Their Pathophysiology. Gut and Liver, 2017. 11(3): p. 323–334.

40. Stone, A.A. and J.E. Broderick, Obesity and pain are associated in the United States. Obesity (Silver Spring), 2012. 20(7): p. 1491–5.

41. Cani, P.D. and B.F. Jordan, Gut microbiota-mediated inflammation in obesity: a link with gastrointestinal cancer. Nat Rev Gastroenterol Hepatol, 2018.

42. Cox, A.J., N.P. West, and A.W. Cripps, Obesity, inflammation, and the gut microbiota. Lancet Diabetes Endocrinol, 2015. 3(3): p. 207–15.

43. Fengming, Y. and W. Jianbing, Biomarkers of inflammatory bowel disease. Dis Markers, 2014. 2014: p. 710915.

44. Mitselos, I.V., et al., Association of clinical and inflammatory markers with small bowel capsule endoscopy findings in Crohn’s disease. Eur J Gastroenterol Hepatol, 2018. 30(8): p. 861–867.

45. Liebregts, T., et al., Immune activation in patients with irritable bowel syndrome. Gastroenterology, 2007. 132(3): p. 913–20.

46. Leach, M.W., et al., The role of IL-10 in inflammatory bowel disease: “of mice and men”. Toxicol Pathol, 1999. 27(1): p. 123–33.

47. Manzanillo, P., C. Eidenschenk, and W.J. Ouyang, Deciphering the crosstalk among IL-1 and IL-10 family cytokines in intestinal immunity. Trends in Immunology, 2015. 36(8): p. 471–478.

48. Shah, N., et al., Interleukin-10 and interleukin-10-receptor defects in inflammatory bowel disease. Curr Allergy Asthma Rep, 2012. 12(5): p. 373–9.

49. Sood, R., et al., Systematic review with meta-analysis: the accuracy of diagnosing irritable bowel syndrome with symptoms, biomarkers and/or psychological markers. Aliment Pharmacol Ther, 2015. 42(5): p. 491–503.

