## Supplementary Data 2. IBS by Weight for "Complete blood count with differential: An effective diagnostic for IBS subtype in the context of BMI?"

#### Fit Group

##### Oneway Analysis of MPV By IBS-subtype Weight=0

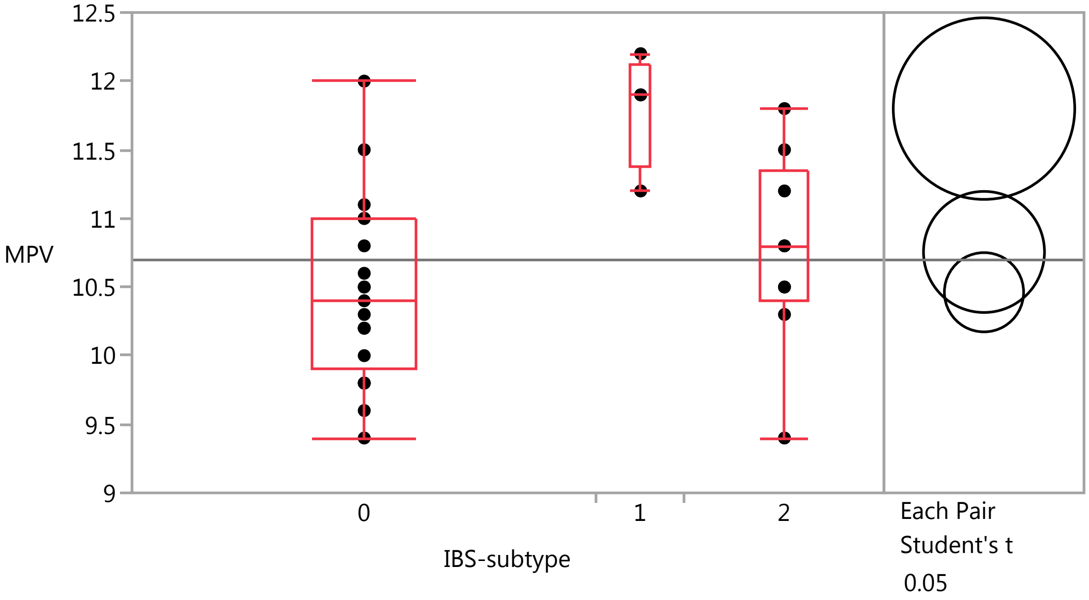


###### Quantiles

| Level | Minimum | 10% | 25% | Median | 75% | 90% | Maximum |
|---|---|---|---|---|---|---|---|
| 0 | 9.4 | 9.64 | 9.9 | 10.4 | 11 | 11.42 | 12 |
| 1 | 11.2 | 11.2 | 11.375 | 11.9 | 12.125 | 12.2 | 12.2 |
| 2 | 9.4 | 9.4 | 10.4 | 10.8 | 11.35 | 11.8 | 11.8 |

###### Means Comparisons

###### Comparisons for each pair using Student's t

###### Confidence Quantile

| t | Alpha |
|---|---|
| 2.03951 | 0.05 |

###### LSD Threshold Matrix

Abs(Dif)-LSD

| | 1 | 2 | 0 |
|---|---|---|---|
| 1 | -0.93576 | 0.24920 | 0.61614 |
| 2 | 0.24920 | -0.62384 | -0.23359 |
| 0 | 0.61614 | -0.23359 | -0.40840 |

Positive values show pairs of means that are significantly different.

###### Connecting Letters Report

| Level | | | Mean |
|---|---|---|---|
| 1 | A | | 11.800000 |
| 2 | | B | 10.755556 |
| 0 | | B | 10.461905 |

Levels not connected by same letter are significantly different.

### Fit Group

#### Oneway Analysis of MPV By IBS-subtype Weight=0

##### Means Comparisons

###### Comparisons for each pair using Student's t

###### Ordered Differences Report

| Level | - Level | Difference | Std Err Dif | Lower CL | Upper CL | p-Value |
|---|---|---|---|---|---|---|
| 1 | 0 | 1.338095 | 0.3539842 | 0.616140 | 2.060051 | 0.0007* |
| 1 | 2 | 1.044444 | 0.3899185 | 0.249200 | 1.839689 | 0.0117* |
| 2 | 0 | 0.293651 | 0.2585135 | -0.233591 | 0.820893 | 0.2647 |

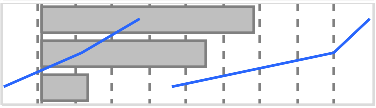

###### Detailed Comparisons Report

###### Comparing 1 with 0

| | | | |
|---|---|---|---|
| Difference | 1.33810 | t Ratio | 3.780099 |
| Std Err Dif | 0.35398 | DF | 31 |
| Upper CL Dif | 2.06005 | Prob > \|t\| | 0.0007* |
| Lower CL Dif | 0.61614 | Prob > t | 0.0003* |
| Confidence | 0.95 | Prob < t | 0.9997 |

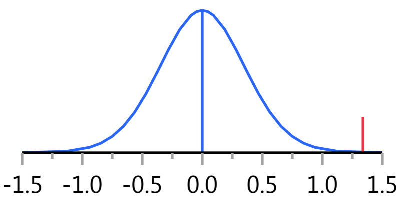


###### Comparing 2 with 0

| | | | |
|---|---|---|---|
| Difference | 0.29365 | t Ratio | 1.135921 |
| Std Err Dif | 0.25851 | DF | 31 |
| Upper CL Dif | 0.82089 | Prob > \|t\| | 0.2647 |
| Lower CL Dif | -0.23359 | Prob > t | 0.1323 |
| Confidence | 0.95 | Prob < t | 0.8677 |

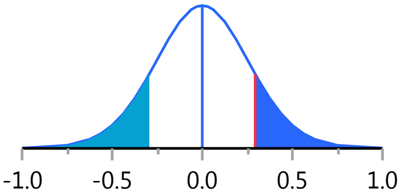


###### Comparing 2 with 1

| | | | |
|---|---|---|---|
| Difference | -1.0444 | t Ratio | -2.67862 |
| Std Err Dif | 0.3899 | DF | 31 |
| Upper CL Dif | -0.2492 | Prob > \|t\| | 0.0117* |
| Lower CL Dif | -1.8397 | Prob > t | 0.9941 |
| Confidence | 0.95 | Prob < t | 0.0059* |

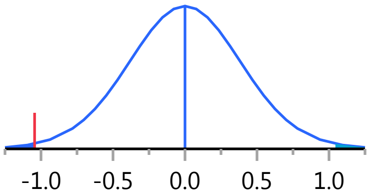


Missing Rows 21

#### Oneway Analysis of MCH By IBS-subtype Weight=0

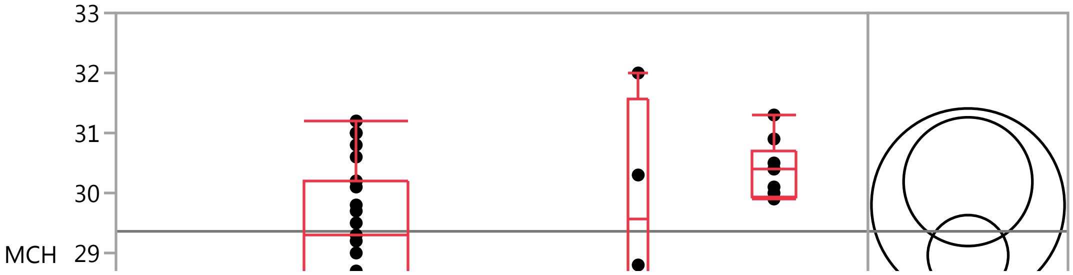

#### Fit Group

##### Oneway Analysis of MCH By IBS-subtype Weight=0

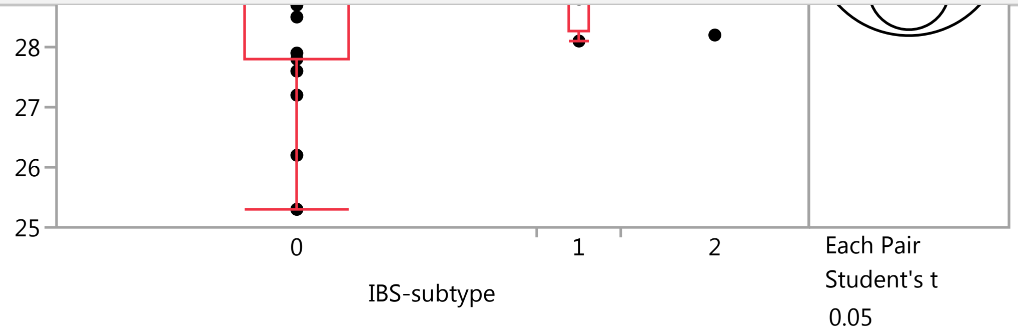


###### Quantiles

| Level | Minimum | 10% | 25% | Median | 75% | 90% | Maximum |
|---|---|---|---|---|---|---|---|
| 0 | 25.3 | 25.66 | 27.8 | 29.3 | 30.2 | 31 | 31.2 |
| 1 | 28.1 | 28.1 | 28.275 | 29.55 | 31.575 | 32 | 32 |
| 2 | 28.2 | 28.2 | 29.95 | 30.4 | 30.7 | 31.3 | 31.3 |

###### Means Comparisons

###### Comparisons for each pair using Student's t

**Confidence Quantile**

| t | Alpha |
|---|---|
| 2.03452 | 0.05 |

**LSD Threshold Matrix**

Abs(Dif)-LSD

| | 2 | 1 | 0 |
|---|---|---|---|
| 2 | -1.5174 | -1.5454 | -0.0376 |
| 1 | -1.5454 | -2.2761 | -0.9046 |
| 0 | -0.0376 | -0.9046 | -0.9492 |

Positive values show pairs of means that are significantly different.

**Connecting Letters Report**

| Level | | Mean |
|---|---|---|
| 2 | A | 30.188889 |
| 1 | A | 29.800000 |
| 0 | A | 28.960870 |

Levels not connected by same letter are significantly different.

**Ordered Differences Report**

| Level | - Level | Difference | Std Err Dif | Lower CL | Upper CL | p-Value |
|---|---|---|---|---|---|---|
| 2 | 0 | 1.228019 | 0.6220604 | -0.03757 | 2.493611 | 0.0568 |
| 1 | 0 | 0.839130 | 0.8570984 | -0.90465 | 2.582910 | 0.3347 |
| 2 | 1 | 0.388889 | 0.9507430 | -1.54541 | 2.323190 | 0.6852 |

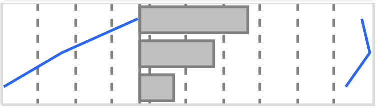

### Fit Group

#### Oneway Analysis of MCH By IBS-subtype Weight=0

##### Means Comparisons

###### Comparisons for each pair using Student's t

###### Detailed Comparisons Report

###### Comparing 1 with 0

| | | | |
|---|---|---|---|
| Difference | 0.8391 | t Ratio | 0.979036 |
| Std Err Dif | 0.8571 | DF | 33 |
| Upper CL Dif | 2.5829 | Prob > \|t\| | 0.3347 |
| Lower CL Dif | -0.9046 | Prob > t | 0.1673 |
| Confidence | 0.95 | Prob < t | 0.8327 |

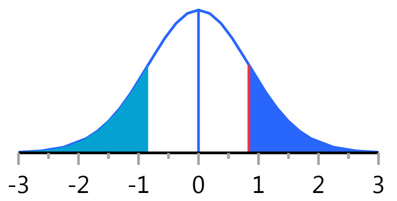


###### Comparing 2 with 0

| | | | |
|---|---|---|---|
| Difference | 1.2280 | t Ratio | 1.974116 |
| Std Err Dif | 0.6221 | DF | 33 |
| Upper CL Dif | 2.4936 | Prob > \|t\| | 0.0568 |
| Lower CL Dif | -0.0376 | Prob > t | 0.0284* |
| Confidence | 0.95 | Prob < t | 0.9716 |

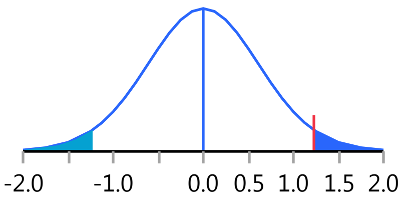


###### Comparing 2 with 1

| | | | |
|---|---|---|---|
| Difference | 0.3889 | t Ratio | 0.409037 |
| Std Err Dif | 0.9507 | DF | 33 |
| Upper CL Dif | 2.3232 | Prob > \|t\| | 0.6852 |
| Lower CL Dif | -1.5454 | Prob > t | 0.3426 |
| Confidence | 0.95 | Prob < t | 0.6574 |

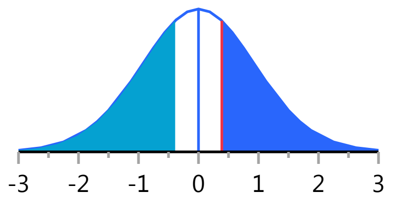


Missing Rows 19

#### Oneway Analysis of SerumCortisol By IBS-subtype Weight=0

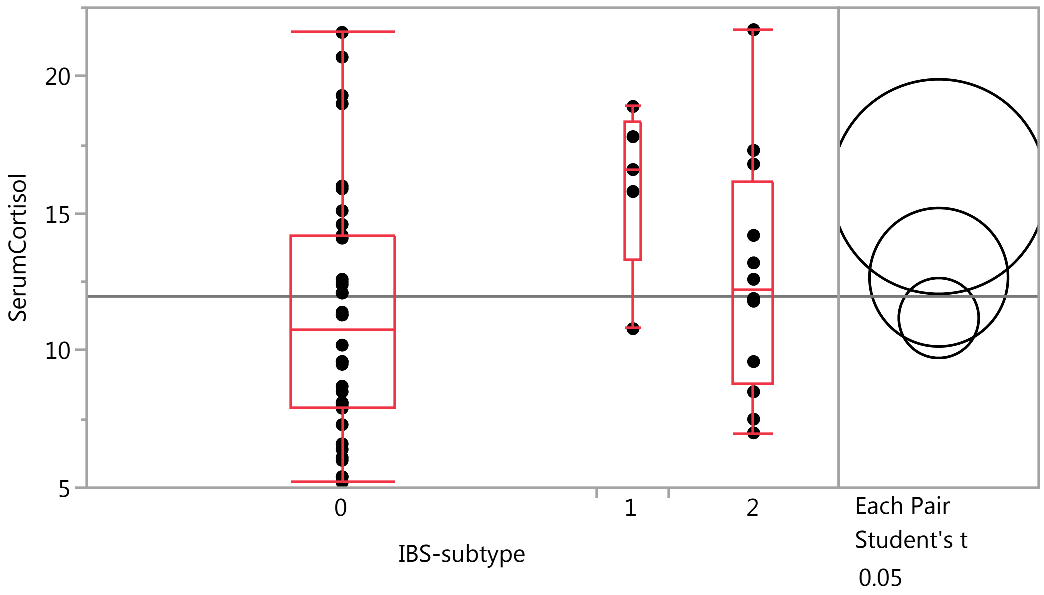

### Fit Group

#### Oneway Analysis of SerumCortisol By IBS-subtype Weight=0

##### Quantiles

| Level | Minimum | 10% | 25% | Median | 75% | 90% | Maximum |
|---|---|---|---|---|---|---|---|
| 0 | 5.2 | 5.82 | 7.925 | 10.75 | 14.175 | 19.09 | 21.6 |
| 1 | 10.8 | 10.8 | 13.3 | 16.6 | 18.35 | 18.9 | 18.9 |
| 2 | 7 | 7.15 | 8.775 | 12.25 | 16.15 | 20.38 | 21.7 |

##### Means Comparisons

###### Comparisons for each pair using Student's t

###### Confidence Quantile

| t | Alpha |
|---|---|
| 2.00856 | 0.05 |

###### LSD Threshold Matrix

Abs(Dif)-LSD

| | 1 | 2 | 0 |
|---|---|---|---|
| 1 | -5.5325 | -1.3513 | 0.6162 |
| 2 | -1.3513 | -3.5712 | -1.4297 |
| 0 | 0.6162 | -1.4297 | -2.0618 |

Positive values show pairs of means that are significantly different.

###### Connecting Letters Report

| Level | | | Mean |
|---|---|---|---|
| 1 | A | | 15.980000 |
| 2 | A | B | 12.675000 |
| 0 | | B | 11.188889 |

Levels not connected by same letter are significantly different.

###### Ordered Differences Report

| Level | - Level | Difference | Std Err Dif | Lower CL | Upper CL | p-Value |
|---|---|---|---|---|---|---|
| 1 | 0 | 4.791111 | 2.078541 | 0.61624 | 8.965984 | 0.0254* |
| 1 | 2 | 3.305000 | 2.318205 | -1.35125 | 7.961252 | 0.1602 |
| 2 | 0 | 1.486111 | 1.451716 | -1.42975 | 4.401969 | 0.3109 |

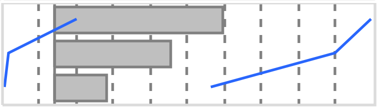

### Fit Group

#### Oneway Analysis of SerumCortisol By IBS-subtype Weight=0

##### Means Comparisons

###### Comparisons for each pair using Student's t

###### Detailed Comparisons Report

###### Comparing 1 with 0

| | | | |
|---|---|---|---|
| Difference | 4.79111 | t Ratio | 2.305035 |
| Std Err Dif | 2.07854 | DF | 50 |
| Upper CL Dif | 8.96598 | Prob > \|t\| | 0.0254* |
| Lower CL Dif | 0.61624 | Prob > t | 0.0127* |
| Confidence | 0.95 | Prob < t | 0.9873 |

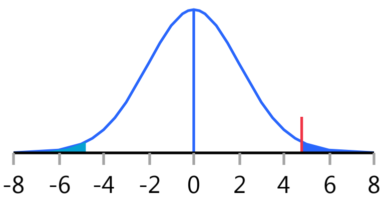


###### Comparing 2 with 0

| | | | |
|---|---|---|---|
| Difference | 1.4861 | t Ratio | 1.023693 |
| Std Err Dif | 1.4517 | DF | 50 |
| Upper CL Dif | 4.4020 | Prob > \|t\| | 0.3109 |
| Lower CL Dif | -1.4297 | Prob > t | 0.1555 |
| Confidence | 0.95 | Prob < t | 0.8445 |

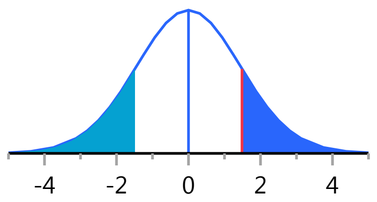


###### Comparing 2 with 1

| | | | |
|---|---|---|---|
| Difference | -3.3050 | t Ratio | -1.42567 |
| Std Err Dif | 2.3182 | DF | 50 |
| Upper CL Dif | 1.3513 | Prob > \|t\| | 0.1602 |
| Lower CL Dif | -7.9613 | Prob > t | 0.9199 |
| Confidence | 0.95 | Prob < t | 0.0801 |

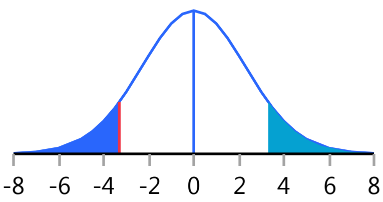


Missing Rows 2

#### Oneway Analysis of LBP By IBS-subtype Weight=0

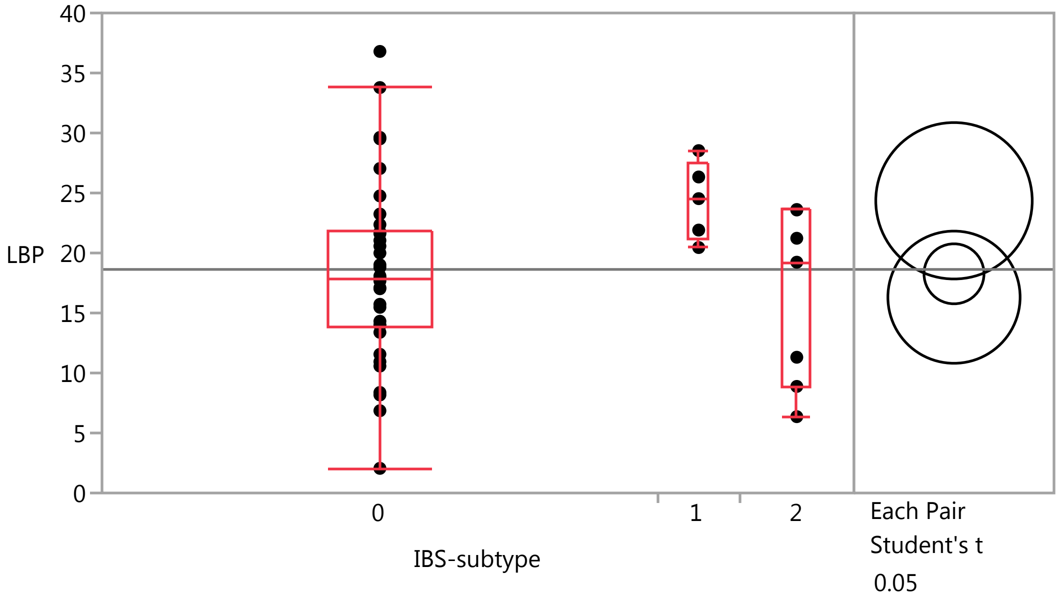

### Fit Group

#### Oneway Analysis of LBP By IBS-subtype Weight=0

##### Quantiles

| Level | Minimum | 10% | 25% | Median | 75% | 90% | Maximum |
|---|---|---|---|---|---|---|---|
| 0 | 2.054 | 8.2745 | 13.813 | 17.911 | 21.90325 | 29.572 | 36.797 |
| 1 | 20.454 | 20.454 | 21.182 | 24.53 | 27.4335 | 28.532 | 28.532 |
| 2 | 6.362 | 6.362 | 8.882 | 19.233 | 23.584 | 23.624 | 23.624 |

##### Means Comparisons

###### Comparisons for each pair using Student's t

###### Confidence Quantile

| t | Alpha |
|---|---|
| 2.01669 | 0.05 |

###### LSD Threshold Matrix

Abs(Dif)-LSD

| | 1 | 0 | 2 |
|---|---|---|---|
| 1 | -9.2183 | -0.8949 | -0.5001 |
| 0 | -0.8949 | -3.5350 | -4.1014 |
| 2 | -0.5001 | -4.1014 | -7.7908 |

Positive values show pairs of means that are significantly different.

###### Connecting Letters Report

| Level | | Mean |
|---|---|---|
| 1 | A | 24.352200 |
| 0 | A | 18.266000 |
| 2 | A | 16.317857 |

Levels not connected by same letter are significantly different.

###### Ordered Differences Report

| Level | - Level | Difference | Std Err Dif | Lower CL | Upper CL | p-Value |
|---|---|---|---|---|---|---|
| 1 | 2 | 8.034343 | 4.231903 | -0.50010 | 16.56879 | 0.0644 |
| 1 | 0 | 6.086200 | 3.461681 | -0.89494 | 13.06734 | 0.0858 |
| 0 | 2 | 1.948143 | 2.999733 | -4.10140 | 7.99768 | 0.5195 |

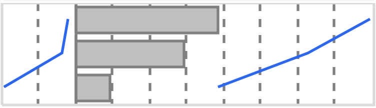

#### Fit Group

##### Oneway Analysis of LBP By IBS-subtype Weight=0

###### Means Comparisons

###### Comparisons for each pair using Student's t

###### Detailed Comparisons Report

**Comparing 1 with 0**

| | | | |
|---|---|---|---|
| Difference | 6.086 | t Ratio | 1.758163 |
| Std Err Dif | 3.462 | DF | 43 |
| Upper CL Dif | 13.067 | Prob > \|t\| | 0.0858 |
| Lower CL Dif | -0.895 | Prob > t | 0.0429* |
| Confidence | 0.95 | Prob < t | 0.9571 |

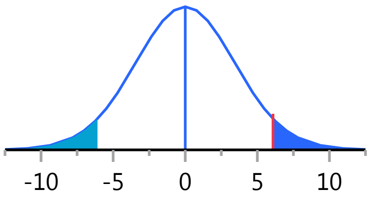


**Comparing 2 with 0**

| | | | |
|---|---|---|---|
| Difference | -1.9481 | t Ratio | -0.64944 |
| Std Err Dif | 2.9997 | DF | 43 |
| Upper CL Dif | 4.1014 | Prob > \|t\| | 0.5195 |
| Lower CL Dif | -7.9977 | Prob > t | 0.7402 |
| Confidence | 0.95 | Prob < t | 0.2598 |

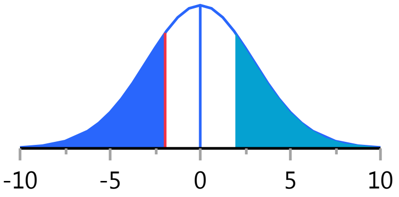


**Comparing 2 with 1**

| | | | |
|---|---|---|---|
| Difference | -8.034 | t Ratio | -1.89852 |
| Std Err Dif | 4.232 | DF | 43 |
| Upper CL Dif | 0.500 | Prob > \|t\| | 0.0644 |
| Lower CL Dif | -16.569 | Prob > t | 0.9678 |
| Confidence | 0.95 | Prob < t | 0.0322* |

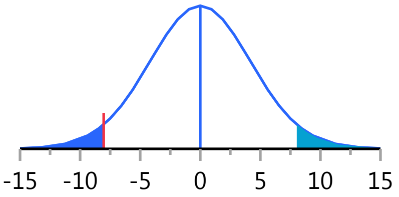


Missing Rows 9

##### Oneway Analysis of ACTH By IBS-subtype Weight=0

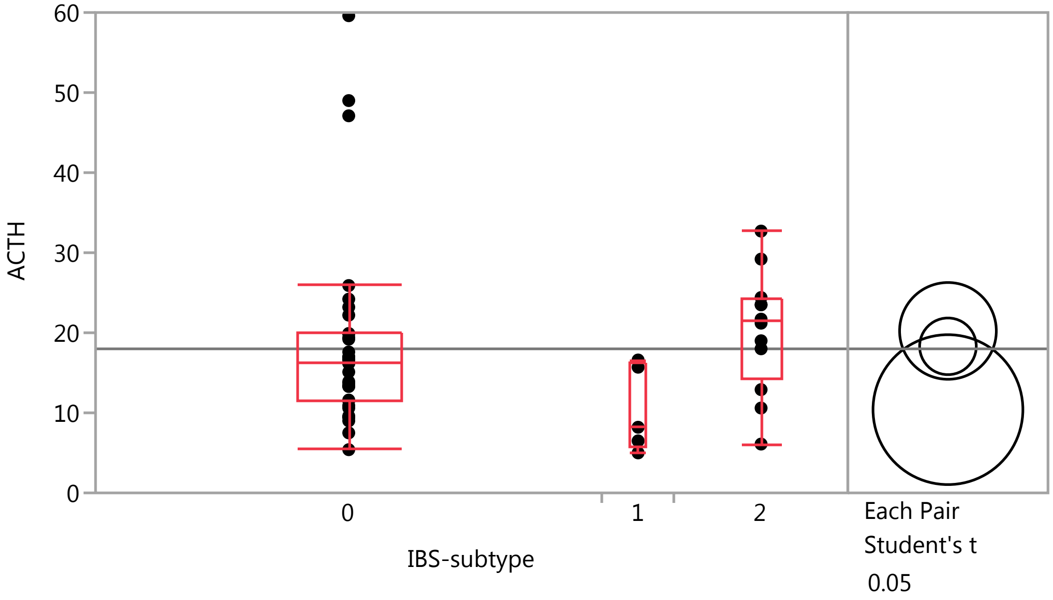

### Fit Group

#### Oneway Analysis of ACTH By IBS-subtype Weight=0

##### Quantiles

| Level | Minimum | 10% | 25% | Median | 75% | 90% | Maximum |
|---|---|---|---|---|---|---|---|
| 0 | 5.4 | 9.24 | 11.5 | 16.2 | 19.9 | 34.38 | 59.6 |
| 1 | 5 | 5 | 5.75 | 8.2 | 16.15 | 16.6 | 16.6 |
| 2 | 6.1 | 7.45 | 14.175 | 21.45 | 24.175 | 31.65 | 32.7 |

##### Means Comparisons

###### Comparisons for each pair using Student's t

###### Confidence Quantile

| t | Alpha |
|---|---|
| 2.00958 | 0.05 |

###### LSD Threshold Matrix

Abs(Dif)-LSD

| | 2 | 0 | 1 |
|---|---|---|---|
| 2 | -8.546 | -5.078 | -1.309 |
| 0 | -5.078 | -5.004 | -2.100 |
| 1 | -1.309 | -2.100 | -13.240 |

Positive values show pairs of means that are significantly different.

###### Connecting Letters Report

| Level | | Mean |
|---|---|---|
| 2 | A | 20.233333 |
| 0 | A | 18.308571 |
| 1 | A | 10.400000 |

Levels not connected by same letter are significantly different.

###### Ordered Differences Report

| Level | - Level | Difference | Std Err Dif | Lower CL | Upper CL | p-Value |
|---|---|---|---|---|---|---|
| 2 | 1 | 9.833333 | 5.544837 | -1.30943 | 20.97610 | 0.0824 |
| 0 | 1 | 7.908571 | 4.980247 | -2.09961 | 17.91675 | 0.1187 |
| 2 | 0 | 1.924762 | 3.484691 | -5.07799 | 8.92751 | 0.5832 |

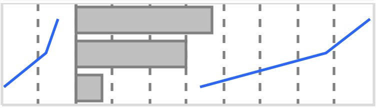

#### Fit Group

##### Oneway Analysis of ACTH By IBS-subtype Weight=0

###### Means Comparisons

###### Comparisons for each pair using Student's t

###### Detailed Comparisons Report

###### Comparing 1 with 0

| | | | |
|---|---|---|---|
| Difference | -7.909 | t Ratio | -1.58799 |
| Std Err Dif | 4.980 | DF | 49 |
| Upper CL Dif | 2.100 | Prob > \|t\| | 0.1187 |
| Lower CL Dif | -17.917 | Prob > t | 0.9406 |
| Confidence | 0.95 | Prob < t | 0.0594 |

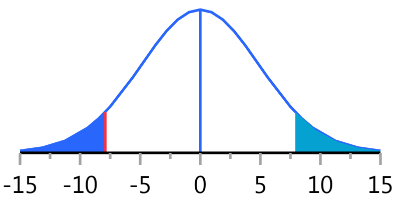


###### Comparing 2 with 0

| | | | |
|---|---|---|---|
| Difference | 1.9248 | t Ratio | 0.552348 |
| Std Err Dif | 3.4847 | DF | 49 |
| Upper CL Dif | 8.9275 | Prob > \|t\| | 0.5832 |
| Lower CL Dif | -5.0780 | Prob > t | 0.2916 |
| Confidence | 0.95 | Prob < t | 0.7084 |

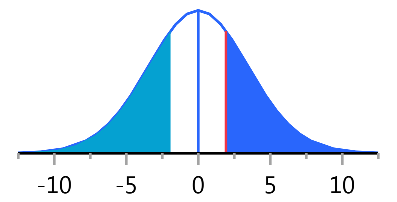


###### Comparing 2 with 1

| | | | |
|---|---|---|---|
| Difference | 9.833 | t Ratio | 1.773422 |
| Std Err Dif | 5.545 | DF | 49 |
| Upper CL Dif | 20.976 | Prob > \|t\| | 0.0824 |
| Lower CL Dif | -1.309 | Prob > t | 0.0412* |
| Confidence | 0.95 | Prob < t | 0.9588 |

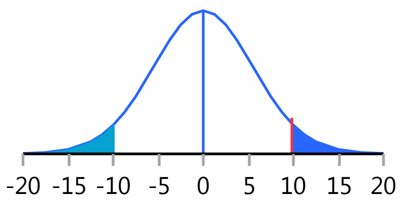


Missing Rows 3

##### Oneway Analysis of IgA By IBS-subtype Weight=0

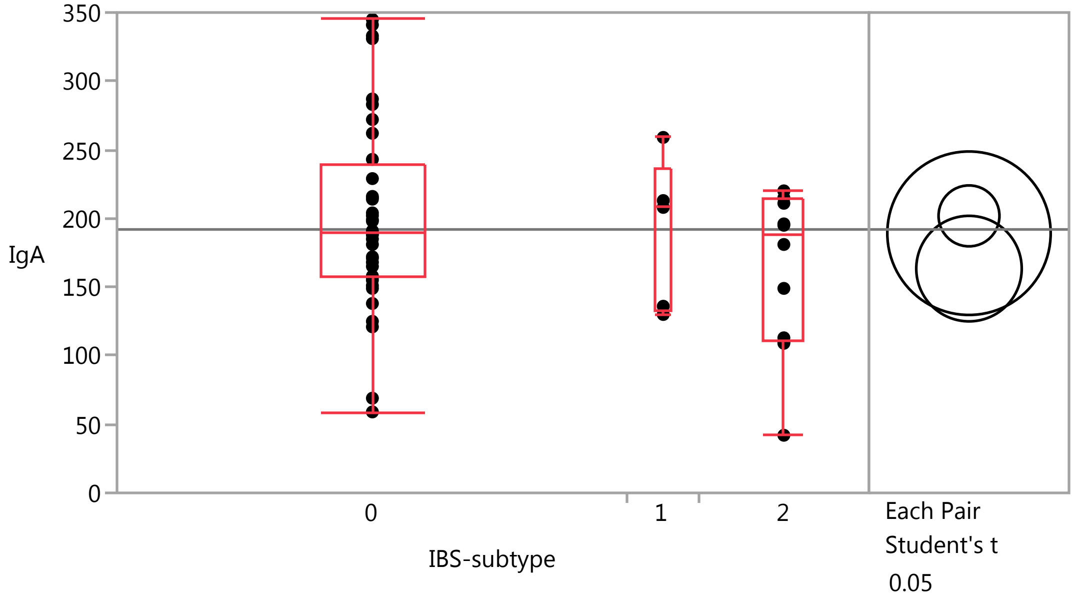

### Fit Group

#### Oneway Analysis of IgA By IBS-subtype Weight=0

##### Quantiles

| Level | Minimum | 10% | 25% | Median | 75% | 90% | Maximum |
|---|---|---|---|---|---|---|---|
| 0 | 59 | 123.8 | 157.25 | 190 | 239.5 | 331.6 | 345 |
| 1 | 130 | 130 | 133 | 208 | 236 | 259 | 259 |
| 2 | 42 | 62.1 | 110.75 | 188 | 214 | 220 | 220 |

##### Means Comparisons

###### Comparisons for each pair using Student's t

###### Confidence Quantile

| t | Alpha |
|---|---|
| 2.00856 | 0.05 |

###### LSD Threshold Matrix

Abs(Dif)-LSD

| | 0 | 1 | 2 |
|---|---|---|---|
| 0 | -31.392 | -50.930 | -5.978 |
| 1 | -50.930 | -84.233 | -45.110 |
| 2 | -5.978 | -45.110 | -54.372 |

Positive values show pairs of means that are significantly different.

###### Connecting Letters Report

| Level | | Mean |
|---|---|---|
| 0 | A | 201.83333 |
| 1 | A | 189.20000 |
| 2 | A | 163.41667 |

Levels not connected by same letter are significantly different.

###### Ordered Differences Report

| Level | - Level | Difference | Std Err Dif | Lower CL | Upper CL | p-Value |
|---|---|---|---|---|---|---|
| 0 | 2 | 38.41667 | 22.10283 | -5.9782 | 82.81150 | 0.0884 |
| 1 | 2 | 25.78333 | 35.29538 | -45.1095 | 96.67620 | 0.4685 |
| 0 | 1 | 12.63333 | 31.64643 | -50.9304 | 76.19706 | 0.6914 |

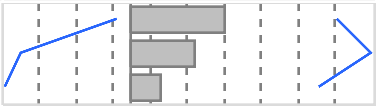

### Fit Group

#### Oneway Analysis of IgA By IBS-subtype Weight=0

##### Means Comparisons

###### Comparisons for each pair using Student's t

###### Detailed Comparisons Report

###### Comparing 1 with 0

| | | | |
|---|---|---|---|
| Difference | -12.633 | t Ratio | -0.3992 |
| Std Err Dif | 31.646 | DF | 50 |
| Upper CL Dif | 50.930 | Prob > \|t\| | 0.6914 |
| Lower CL Dif | -76.197 | Prob > t | 0.6543 |
| Confidence | 0.95 | Prob < t | 0.3457 |

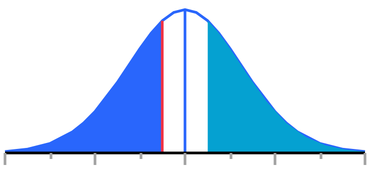


###### Comparing 2 with 0

| | | | |
|---|---|---|---|
| Difference | -38.417 | t Ratio | -1.73809 |
| Std Err Dif | 22.103 | DF | 50 |
| Upper CL Dif | 5.978 | Prob > \|t\| | 0.0884 |
| Lower CL Dif | -82.811 | Prob > t | 0.9558 |
| Confidence | 0.95 | Prob < t | 0.0442* |

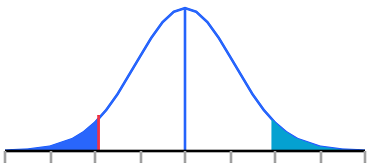


###### Comparing 2 with 1

| | | | |
|---|---|---|---|
| Difference | -25.783 | t Ratio | -0.7305 |
| Std Err Dif | 35.295 | DF | 50 |
| Upper CL Dif | 45.110 | Prob > \|t\| | 0.4685 |
| Lower CL Dif | -96.676 | Prob > t | 0.7658 |
| Confidence | 0.95 | Prob < t | 0.2342 |

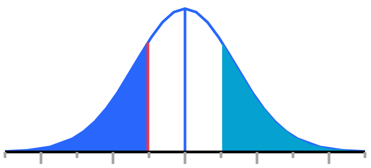


Missing Rows 2

#### Oneway Analysis of sCD14 By IBS-subtype Weight=0

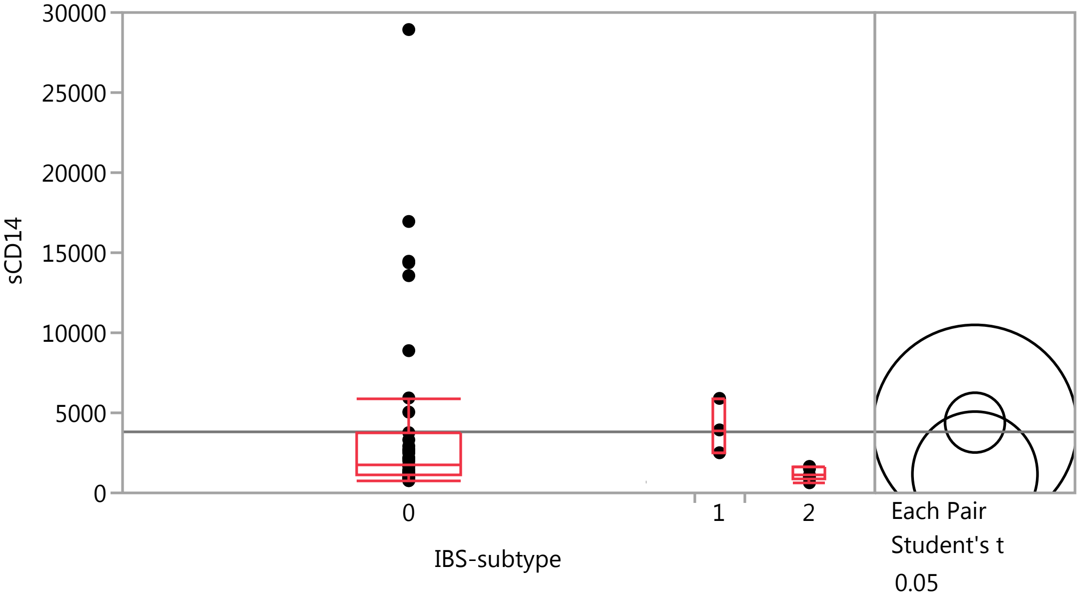

### Fit Group

#### Oneway Analysis of sCD14 By IBS-subtype Weight=0

##### Quantiles

| Level | Minimum | 10% | 25% | Median | 75% | 90% | Maximum |
|---|---|---|---|---|---|---|---|
| 0 | 767 | 878.72 | 1129.8 | 1754.3 | 3763.1 | 14411.1 | 28936 |
| 1 | 2508.3 | 2508.3 | 2508.3 | 3934.1 | 5902.4 | 5902.4 | 5902.4 |
| 2 | 634.7 | 634.7 | 896.375 | 1065.75 | 1606.525 | 1653.1 | 1653.1 |

##### Means Comparisons

###### Comparisons for each pair using Student's t

###### Confidence Quantile

| t | Alpha |
|---|---|
| 2.01669 | 0.05 |

###### LSD Threshold Matrix

Abs(Dif)-LSD

| | 0 | 1 | 2 |
|---|---|---|---|
| 0 | -2641.5 | -6372.4 | -1113.0 |
| 1 | -6372.4 | -9022.3 | -4538.7 |
| 2 | -1113.0 | -4538.7 | -5525.0 |

Positive values show pairs of means that are significantly different.

###### Connecting Letters Report

| Level | | Mean |
|---|---|---|
| 0 | A | 4390.0286 |
| 1 | A | 4114.9333 |
| 2 | A | 1172.7125 |

Levels not connected by same letter are significantly different.

###### Ordered Differences Report

| Level | - Level | Difference | Std Err Dif | Lower CL | Upper CL | p-Value |
|---|---|---|---|---|---|---|
| 0 | 2 | 3217.316 | 2147.228 | -1112.98 | 7547.61 | 0.1413 |
| 1 | 2 | 2942.221 | 3709.486 | -4538.67 | 10423.11 | 0.4320 |
| 0 | 1 | 275.095 | 3296.250 | -6372.43 | 6922.62 | 0.9339 |

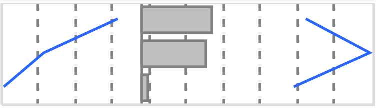

### Fit Group

#### Oneway Analysis of sCD14 By IBS-subtype Weight=0

##### Means Comparisons

###### Comparisons for each pair using Student's t

###### Detailed Comparisons Report

###### Comparing 1 with 0

| | | | |
|---|---|---|---|
| Difference | -275.1 | t Ratio | -0.08346 |
| Std Err Dif | 3296.3 | DF | 43 |
| Upper CL Dif | 6372.4 | Prob > \|t\| | 0.9339 |
| Lower CL Dif | -6922.6 | Prob > t | 0.5331 |
| Confidence | 0.95 | Prob < t | 0.4669 |

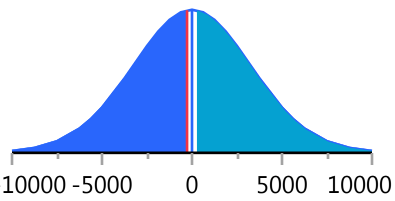


###### Comparing 2 with 0

| | | | |
|---|---|---|---|
| Difference | -3217.3 | t Ratio | -1.49836 |
| Std Err Dif | 2147.2 | DF | 43 |
| Upper CL Dif | 1113.0 | Prob > \|t\| | 0.1413 |
| Lower CL Dif | -7547.6 | Prob > t | 0.9293 |
| Confidence | 0.95 | Prob < t | 0.0707 |

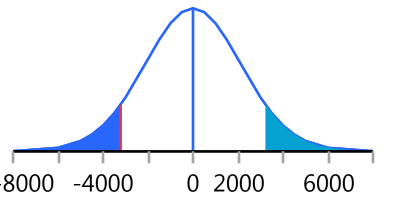


###### Comparing 2 with 1

| | | | |
|---|---|---|---|
| Difference | -2942 | t Ratio | -0.79316 |
| Std Err Dif | 3709 | DF | 43 |
| Upper CL Dif | 4539 | Prob > \|t\| | 0.4320 |
| Lower CL Dif | -10423 | Prob > t | 0.7840 |
| Confidence | 0.95 | Prob < t | 0.2160 |

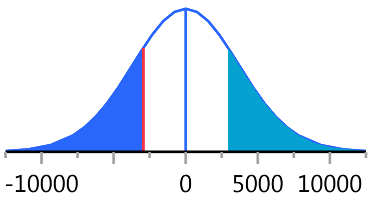


Missing Rows 9

#### Oneway Analysis of Monocytes_PCT By IBS-subtype Weight=0

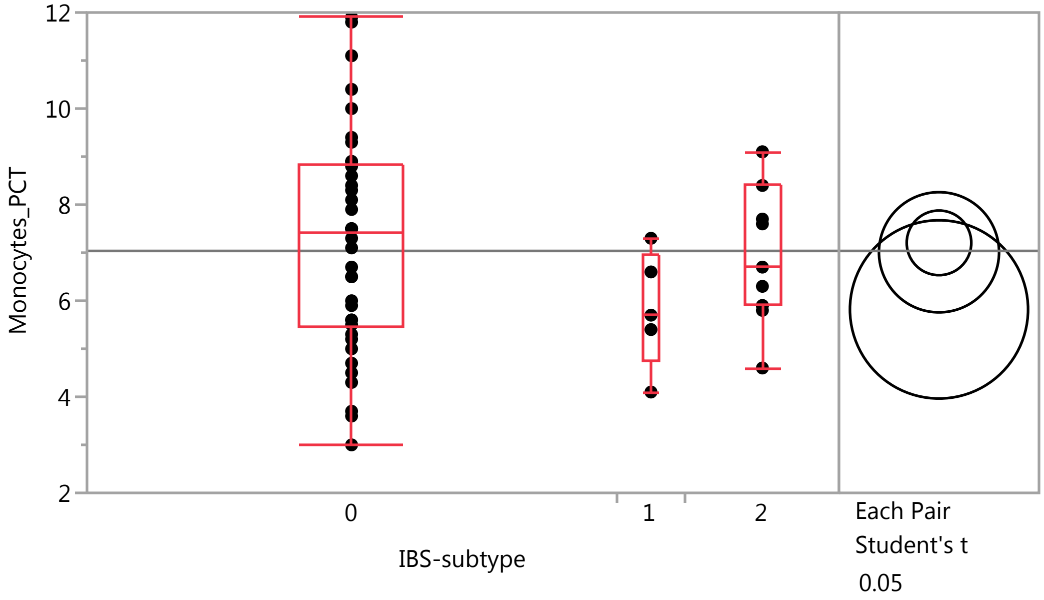

### Fit Group

#### Oneway Analysis of Monocytes_PCT By IBS-subtype Weight=0

##### Quantiles

| Level | Minimum | 10% | 25% | Median | 75% | 90% | Maximum |
|---|---|---|---|---|---|---|---|
| 0 | 3 | 4.24 | 5.45 | 7.4 | 8.825 | 10.47 | 11.9 |
| 1 | 4.1 | 4.1 | 4.75 | 5.7 | 6.95 | 7.3 | 7.3 |
| 2 | 4.6 | 4.84 | 5.9 | 6.7 | 8.4 | 9.1 | 9.1 |

##### Means Comparisons

###### Comparisons for each pair using Student's t

###### Confidence Quantile

| t | Alpha |
|---|---|
| 2.00758 | 0.05 |

###### LSD Threshold Matrix

Abs(Dif)-LSD

| | 0 | 2 | 1 |
|---|---|---|---|
| 0 | -0.9518 | -1.2243 | -0.5885 |
| 2 | -1.2243 | -1.7691 | -1.0487 |
| 1 | -0.5885 | -1.0487 | -2.6240 |

Positive values show pairs of means that are significantly different.

###### Connecting Letters Report

| Level | | Mean |
|---|---|---|
| 0 | A | 7.2052632 |
| 2 | A | 7.0090909 |
| 1 | A | 5.8200000 |

Levels not connected by same letter are significantly different.

###### Ordered Differences Report

| Level | - Level | Difference | Std Err Dif | Lower CL | Upper CL | p-Value |
|---|---|---|---|---|---|---|
| 0 | 1 | 1.385263 | 0.983144 | -0.58848 | 3.359008 | 0.1649 |
| 2 | 1 | 1.189091 | 1.114650 | -1.04866 | 3.426844 | 0.2911 |
| 0 | 2 | 0.196172 | 0.707571 | -1.22433 | 1.616679 | 0.7827 |

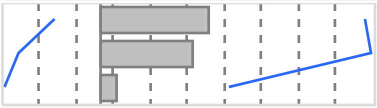

Missing Rows 1

#### Oneway Analysis of Lymphocytes_PCT By IBS-subtype Weight=0

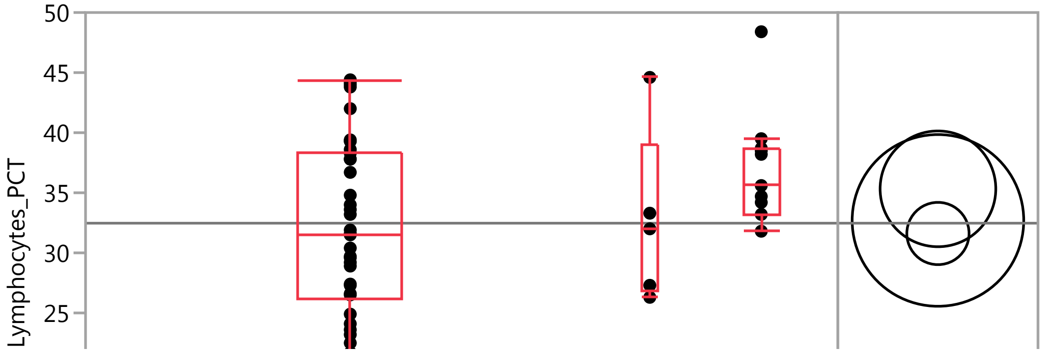

### Fit Group

#### Oneway Analysis of Lymphocytes_PCT By IBS-subtype Weight=0

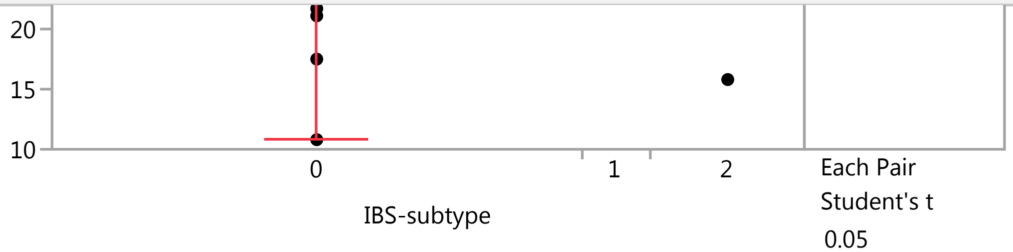


##### Quantiles

| Level | Minimum | 10% | 25% | Median | 75% | 90% | Maximum |
|---|---|---|---|---|---|---|---|
| 0 | 10.8 | 21.64 | 26.1 | 31.55 | 38.375 | 43.82 | 44.4 |
| 1 | 26.3 | 26.3 | 26.8 | 32 | 38.95 | 44.6 | 44.6 |
| 2 | 15.8 | 19 | 33.2 | 35.6 | 38.7 | 46.62 | 48.4 |

##### Means Comparisons

###### Comparisons for each pair using Student's t

###### Confidence Quantile

| t | Alpha |
|---|---|
| 2.00758 | 0.05 |

###### LSD Threshold Matrix

Abs(Dif)-LSD

| | 2 | 1 | 0 |
|---|---|---|---|
| 2 | -6.813 | -5.990 | -1.754 |
| 1 | -5.990 | -10.105 | -6.511 |
| 0 | -1.754 | -6.511 | -3.665 |

Positive values show pairs of means that are significantly different.

###### Connecting Letters Report

| Level | | Mean |
|---|---|---|
| 2 | A | 35.327273 |
| 1 | A | 32.700000 |
| 0 | A | 31.610526 |

Levels not connected by same letter are significantly different.

###### Ordered Differences Report

| Level | - Level | Difference | Std Err Dif | Lower CL | Upper CL | p-Value |
|---|---|---|---|---|---|---|
| 2 | 0 | 3.716746 | 2.724863 | -1.75365 | 9.18714 | 0.1786 |
| 2 | 1 | 2.627273 | 4.292531 | -5.99034 | 11.24489 | 0.5432 |
| 1 | 0 | 1.089474 | 3.786102 | -6.51144 | 8.69039 | 0.7747 |

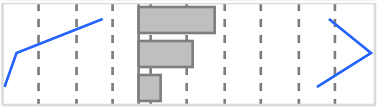

Missing Rows 1

#### Fit Group

##### Oneway Analysis of Neutrophil_PCT By IBS-subtype Weight=0

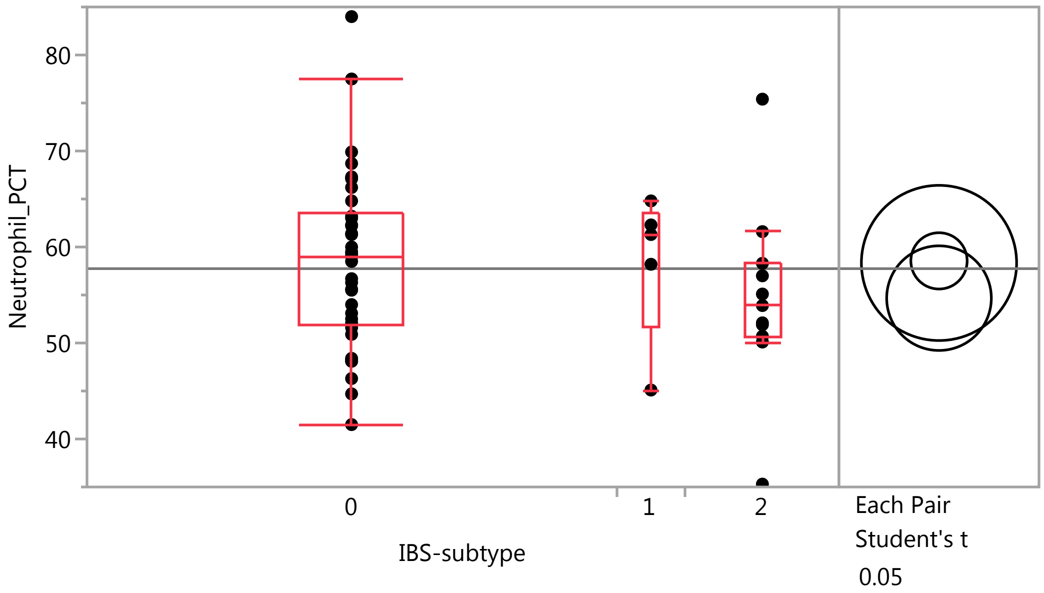


##### Quantiles

| Level | Minimum | 10% | 25% | Median | 75% | 90% | Maximum |
|---|---|---|---|---|---|---|---|
| 0 | 41.5 | 47.92 | 51.975 | 59 | 63.6 | 68.82 | 84 |
| 1 | 45.1 | 45.1 | 51.65 | 61.3 | 63.55 | 64.8 | 64.8 |
| 2 | 35.3 | 38.26 | 50.7 | 53.9 | 58.3 | 72.64 | 75.4 |

##### Means Comparisons

###### Comparisons for each pair using Student's t

###### Confidence Quantile

| t | Alpha |
|---|---|
| 2.00758 | 0.05 |

###### LSD Threshold Matrix

Abs(Dif)-LSD

| | 0 | 1 | 2 |
|---|---|---|---|
| 0 | -4.147 | -8.384 | -2.307 |
| 1 | -8.384 | -11.433 | -6.083 |
| 2 | -2.307 | -6.083 | -7.708 |

Positive values show pairs of means that are significantly different.

###### Connecting Letters Report

| Level | | Mean |
|---|---|---|
| 0 | A | 58.555263 |
| 1 | A | 58.340000 |
| 2 | A | 54.672727 |

Levels not connected by same letter are significantly different.

### Fit Group

#### Oneway Analysis of Neutrophil_PCT By IBS-subtype Weight=0

##### Means Comparisons

###### Comparisons for each pair using Student's t

###### Ordered Differences Report

| Level | - Level | Difference | Std Err Dif | Lower CL | Upper CL | p-Value |
|---|---|---|---|---|---|---|
| 0 | 2 | 3.882536 | 3.082903 | -2.30665 | 10.07172 | 0.2136 |
| 1 | 2 | 3.667273 | 4.856558 | -6.08267 | 13.41722 | 0.4537 |
| 0 | 1 | 0.215263 | 4.283585 | -8.38439 | 8.81492 | 0.9601 |

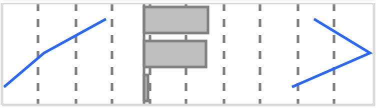

Missing Rows 1

#### Oneway Analysis of Neutrophils By IBS-subtype Weight=0

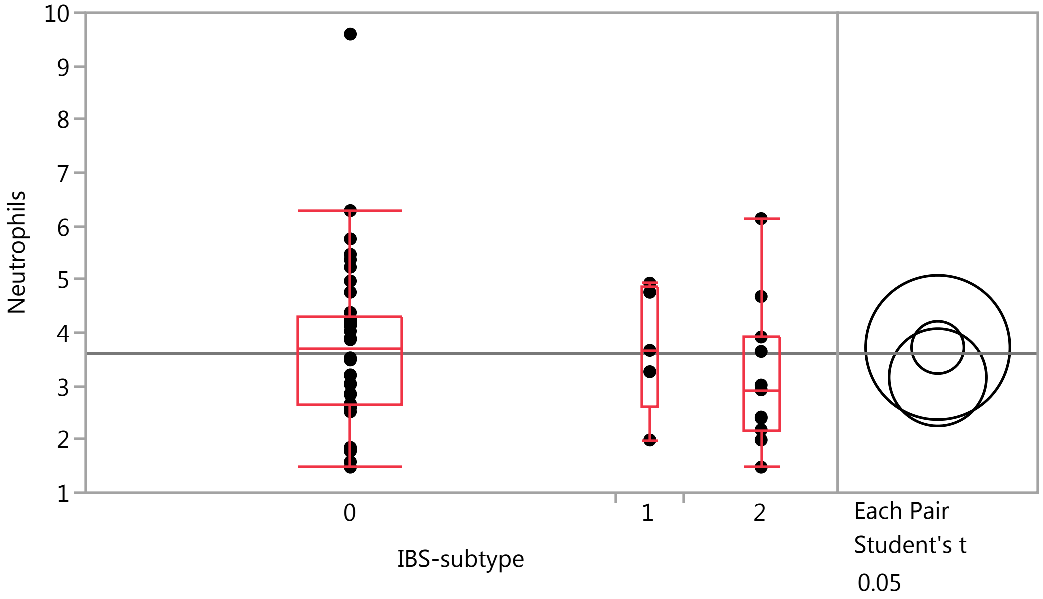


##### Quantiles

| Level | Minimum | 10% | 25% | Median | 75% | 90% | Maximum |
|---|---|---|---|---|---|---|---|
| 0 | 1.48 | 1.789 | 2.64 | 3.7 | 4.29 | 5.499 | 9.6 |
| 1 | 1.99 | 1.99 | 2.63 | 3.67 | 4.845 | 4.93 | 4.93 |
| 2 | 1.48 | 1.582 | 2.18 | 2.93 | 3.92 | 5.848 | 6.14 |

##### Means Comparisons

###### Comparisons for each pair using Student's t

###### Confidence Quantile

| t | Alpha |
|---|---|
| 2.00758 | 0.05 |

### Fit Group

#### Oneway Analysis of Neutrophils By IBS-subtype Weight=0

##### Means Comparisons

###### Comparisons for each pair using Student's t

###### LSD Threshold Matrix

Abs(Dif)-LSD

| | 0 | 1 | 2 |
|---|---|---|---|
| 0 | -0.6939 | -1.4379 | -0.4751 |
| 1 | -1.4379 | -1.9130 | -1.0719 |
| 2 | -0.4751 | -1.0719 | -1.2897 |

Positive values show pairs of means that are significantly different.

###### Connecting Letters Report

| Level | | Mean |
|---|---|---|
| 0 | A | 3.7250000 |
| 1 | A | 3.7240000 |
| 2 | A | 3.1645455 |

Levels not connected by same letter are significantly different.

###### Ordered Differences Report

| Level | - Level | Difference | Std Err Dif | Lower CL | Upper CL | p-Value |
|---|---|---|---|---|---|---|
| 0 | 2 | 0.5604545 | 0.5158340 | -0.47513 | 1.596035 | 0.2824 |
| 1 | 2 | 0.5594545 | 0.8126035 | -1.07192 | 2.190824 | 0.4943 |
| 0 | 1 | 0.0010000 | 0.7167332 | -1.43790 | 1.439902 | 0.9989 |

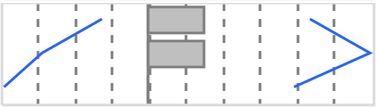

Missing Rows 1

#### Oneway Analysis of IgM By IBS-subtype Weight=0

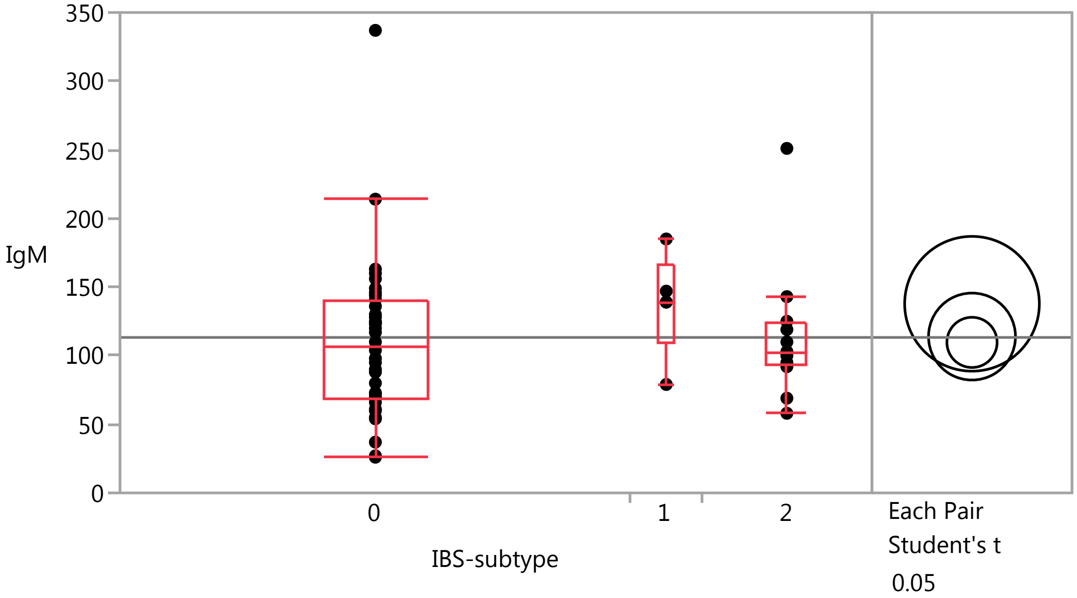

### Fit Group

#### Oneway Analysis of IgM By IBS-subtype Weight=0

##### Quantiles

| Level | Minimum | 10% | 25% | Median | 75% | 90% | Maximum |
|---|---|---|---|---|---|---|---|
| 0 | 26 | 48.9 | 69.25 | 107 | 139.75 | 160.9 | 337 |
| 1 | 79 | 79 | 109 | 139 | 166 | 185 | 185 |
| 2 | 58 | 61.3 | 92.75 | 102.5 | 123.5 | 218.6 | 251 |

##### Means Comparisons

###### Comparisons for each pair using Student's t

###### Confidence Quantile

| t | Alpha |
|---|---|
| 2.00856 | 0.05 |

###### LSD Threshold Matrix

Abs(Dif)-LSD

| | 1 | 2 | 0 |
|---|---|---|---|
| 1 | -69.441 | -34.560 | -24.268 |
| 2 | -34.560 | -44.824 | -32.348 |
| 0 | -24.268 | -32.348 | -25.879 |

Positive values show pairs of means that are significantly different.

###### Connecting Letters Report

| Level | | Mean |
|---|---|---|
| 1 | A | 137.80000 |
| 2 | A | 113.91667 |
| 0 | A | 109.66667 |

Levels not connected by same letter are significantly different.

###### Ordered Differences Report

| Level | - Level | Difference | Std Err Dif | Lower CL | Upper CL | p-Value |
|---|---|---|---|---|---|---|
| 1 | 0 | 28.13333 | 26.08881 | -24.2676 | 80.53425 | 0.2860 |
| 1 | 2 | 23.88333 | 29.09695 | -34.5596 | 82.32627 | 0.4156 |
| 2 | 0 | 4.25000 | 18.22122 | -32.3484 | 40.84839 | 0.8165 |

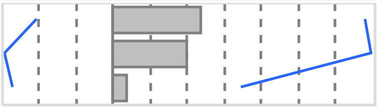

Missing Rows 2

#### Oneway Analysis of Monocytes By IBS-subtype Weight=0

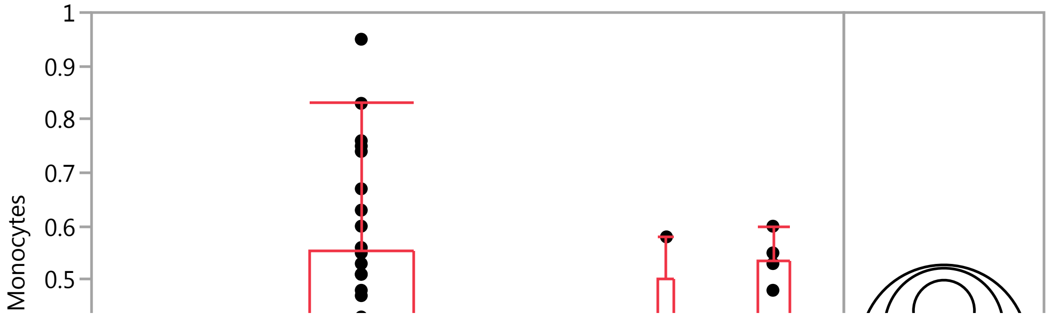

#### Fit Group

##### Oneway Analysis of Monocytes By IBS-subtype Weight=0

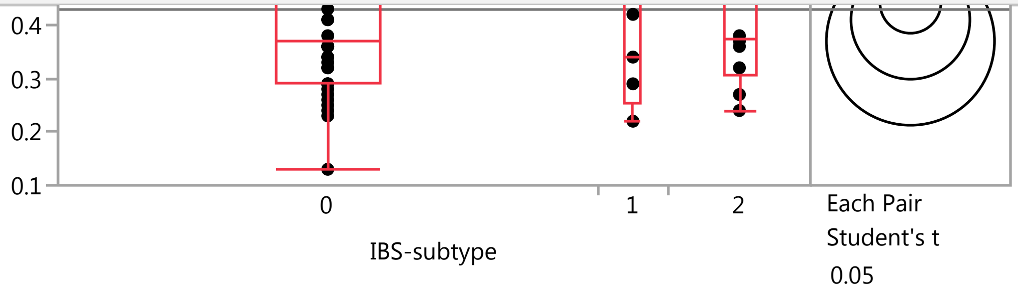


##### Quantiles

| Level | Minimum | 10% | 25% | Median | 75% | 90% | Maximum |
|---|---|---|---|---|---|---|---|
| 0 | 0.13 | 0.249 | 0.29 | 0.37 | 0.5525 | 0.751 | 0.95 |
| 1 | 0.22 | 0.22 | 0.255 | 0.34 | 0.5 | 0.58 | 0.58 |
| 2 | 0.24 | 0.243 | 0.3075 | 0.375 | 0.535 | 0.595 | 0.6 |

##### Means Comparisons

###### Comparisons for each pair using Student's t

###### Confidence Quantile

| t | Alpha |
|---|---|
| 2.00856 | 0.05 |

###### LSD Threshold Matrix

Abs(Dif)-LSD

| | 0 | 2 | 1 |
|---|---|---|---|
| 0 | -0.08081 | -0.09335 | -0.09573 |
| 2 | -0.09335 | -0.15753 | -0.15293 |
| 1 | -0.09573 | -0.15293 | -0.22278 |

Positive values show pairs of means that are significantly different.

###### Connecting Letters Report

| Level | | Mean |
|---|---|---|
| 0 | A | 0.44184211 |
| 2 | A | 0.41000000 |
| 1 | A | 0.37000000 |

Levels not connected by same letter are significantly different.

###### Ordered Differences Report

| Level | - Level | Difference | Std Err Dif | Lower CL | Upper CL | p-Value |
|---|---|---|---|---|---|---|
| 0 | 1 | 0.0718421 | 0.0834292 | -0.095730 | 0.2394147 | 0.3933 |
| 2 | 1 | 0.0400000 | 0.0960553 | -0.152933 | 0.2329328 | 0.6789 |
| 0 | 2 | 0.0318421 | 0.0623289 | -0.093349 | 0.1570334 | 0.6117 |

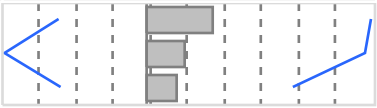

### Fit Group

#### Oneway Analysis of Monocytes By IBS-subtype Weight=0

##### Means Comparisons

###### Comparisons for each pair using Student's t

###### Detailed Comparisons Report

###### Comparing 1 with 0

| | | | |
|---|---|---|---|
| Difference | -0.07184 | t Ratio | -0.86111 |
| Std Err Dif | 0.08343 | DF | 50 |
| Upper CL Dif | 0.09573 | Prob > \|t\| | 0.3933 |
| Lower CL Dif | -0.23941 | Prob > t | 0.8034 |
| Confidence | 0.95 | Prob < t | 0.1966 |

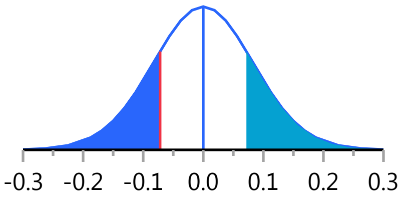


###### Comparing 2 with 0

| | | | |
|---|---|---|---|
| Difference | -0.03184 | t Ratio | -0.51087 |
| Std Err Dif | 0.06233 | DF | 50 |
| Upper CL Dif | 0.09335 | Prob > \|t\| | 0.6117 |
| Lower CL Dif | -0.15703 | Prob > t | 0.6942 |
| Confidence | 0.95 | Prob < t | 0.3058 |

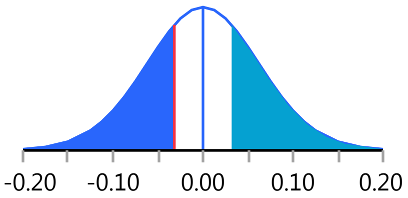


###### Comparing 2 with 1

| | | | |
|---|---|---|---|
| Difference | 0.04000 | t Ratio | 0.416427 |
| Std Err Dif | 0.09606 | DF | 50 |
| Upper CL Dif | 0.23293 | Prob > \|t\| | 0.6789 |
| Lower CL Dif | -0.15293 | Prob > t | 0.3394 |
| Confidence | 0.95 | Prob < t | 0.6606 |

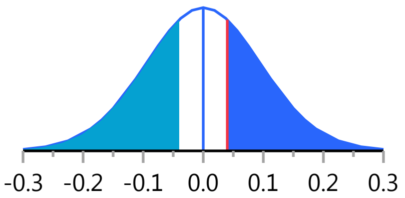


Missing Rows 2

#### Oneway Analysis of WBC By IBS-subtype Weight=0

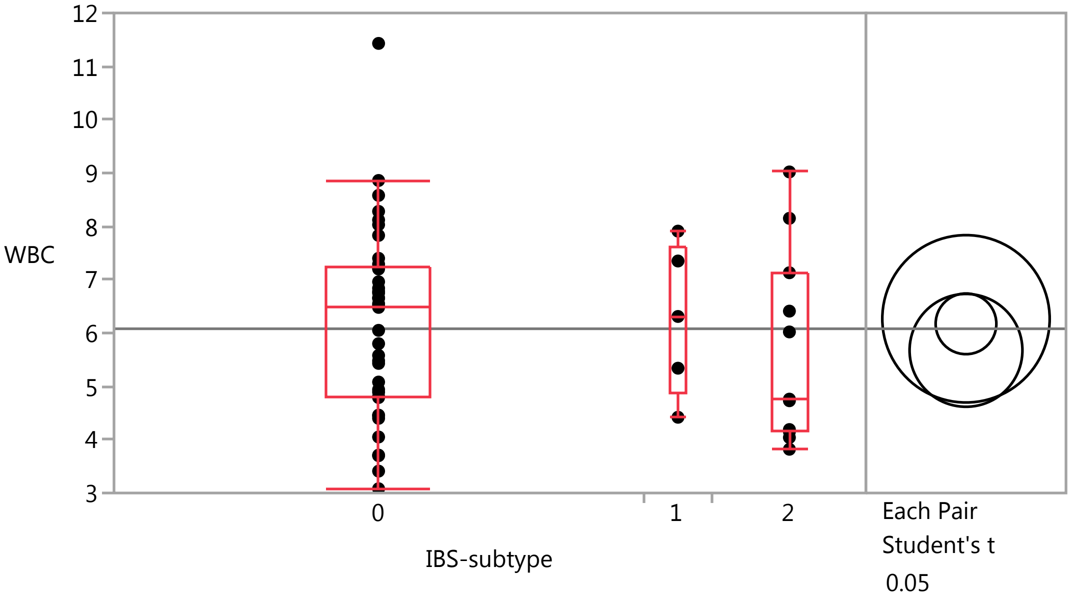

### Fit Group

#### Oneway Analysis of WBC By IBS-subtype Weight=0

##### Quantiles

| Level | Minimum | 10% | 25% | Median | 75% | 90% | Maximum |
|---|---|---|---|---|---|---|---|
| 0 | 3.08 | 3.709 | 4.805 | 6.495 | 7.245 | 8.31 | 11.43 |
| 1 | 4.42 | 4.42 | 4.88 | 6.31 | 7.63 | 7.91 | 7.91 |
| 2 | 3.82 | 3.864 | 4.15 | 4.76 | 7.13 | 8.846 | 9.02 |

##### Means Comparisons

###### Comparisons for each pair using Student's t

###### Confidence Quantile

| t | Alpha |
|---|---|
| 2.00758 | 0.05 |

###### LSD Threshold Matrix

Abs(Dif)-LSD

| | 1 | 0 | 2 |
|---|---|---|---|
| 1 | -2.2192 | -1.5754 | -1.3011 |
| 0 | -1.5754 | -0.8050 | -0.7038 |
| 2 | -1.3011 | -0.7038 | -1.4962 |

Positive values show pairs of means that are significantly different.

###### Connecting Letters Report

| Level | | Mean |
|---|---|---|
| 1 | A | 6.2660000 |
| 0 | A | 6.1721053 |
| 2 | A | 5.6745455 |

Levels not connected by same letter are significantly different.

###### Ordered Differences Report

| Level | - Level | Difference | Std Err Dif | Lower CL | Upper CL | p-Value |
|---|---|---|---|---|---|---|
| 1 | 2 | 0.5914545 | 0.9426992 | -1.30109 | 2.484002 | 0.5332 |
| 0 | 2 | 0.4975598 | 0.5984176 | -0.70381 | 1.698933 | 0.4096 |
| 1 | 0 | 0.0938947 | 0.8314802 | -1.57537 | 1.763161 | 0.9105 |

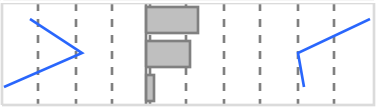

Missing Rows 1

#### Fit Group

##### Oneway Analysis of Eosinophils By IBS-subtype Weight=0

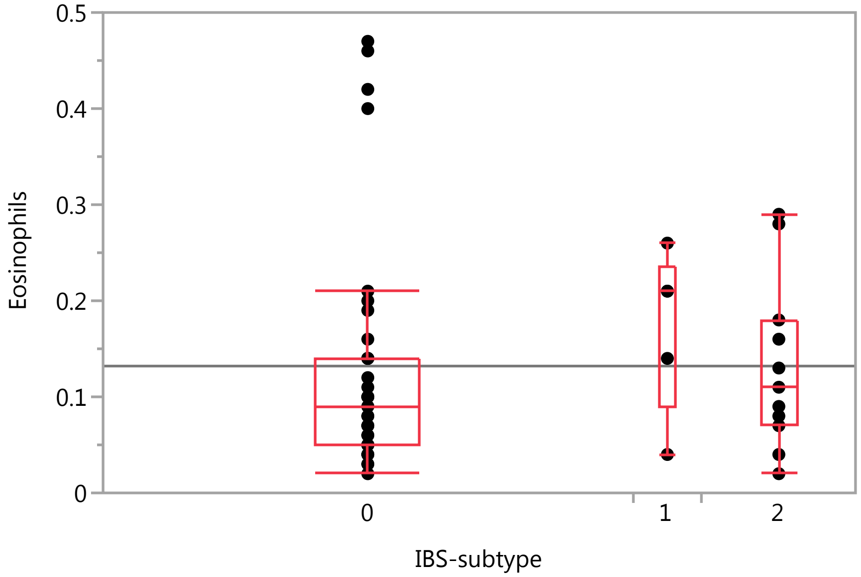


##### Quantiles

| Level | Minimum | 10% | 25% | Median | 75% | 90% | Maximum |
|---|---|---|---|---|---|---|---|
| 0 | 0.02 | 0.039 | 0.05 | 0.09 | 0.14 | 0.402 | 0.47 |
| 1 | 0.04 | 0.04 | 0.09 | 0.21 | 0.235 | 0.26 | 0.26 |
| 2 | 0.02 | 0.024 | 0.07 | 0.11 | 0.18 | 0.288 | 0.29 |

Missing Rows 1

#### Fit Group

##### Oneway Analysis of LDH By IBS-subtype Weight=0

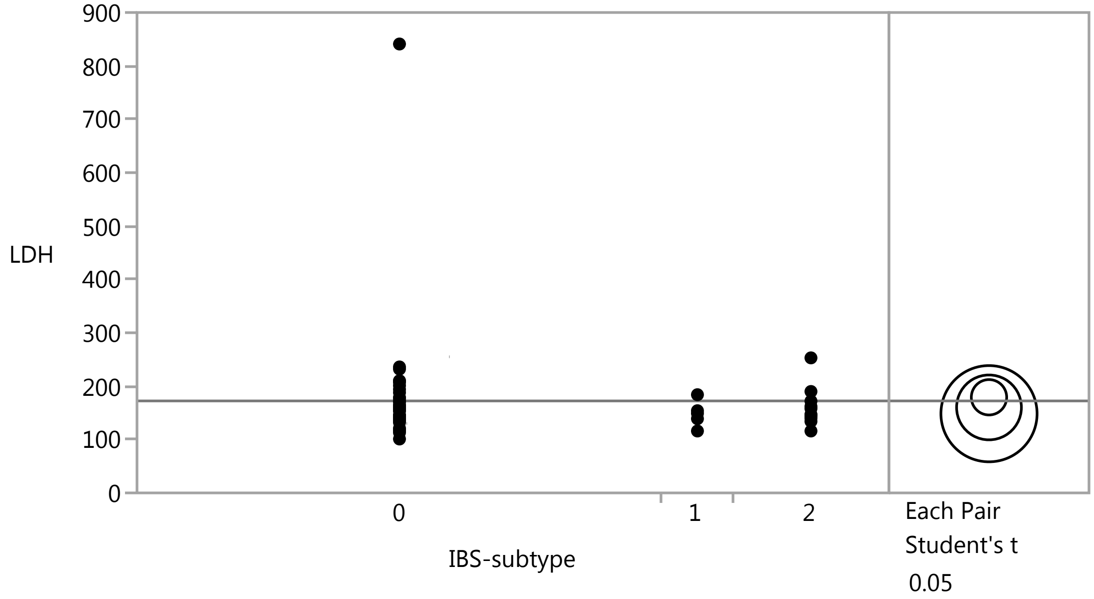


##### Means Comparisons

###### Comparisons for each pair using Student's t

###### Confidence Quantile

| t | Alpha |
|---|---|
| 2.00856 | 0.05 |

###### LSD Threshold Matrix

Abs(Dif)-LSD

| | 0 | 2 | 1 |
|---|---|---|---|
| 0 | -46.86 | -50.24 | -65.28 |
| 2 | -50.24 | -85.95 | -96.94 |
| 1 | -65.28 | -96.94 | -127.48 |

Positive values show pairs of means that are significantly different.

###### Connecting Letters Report

| Level | | Mean |
|---|---|---|
| 0 | A | 179.16216 |
| 2 | A | 160.18182 |
| 1 | A | 148.40000 |

Levels not connected by same letter are significantly different.

###### Ordered Differences Report

| Level | - Level | Difference | Std Err Dif | Lower CL | Upper CL | p-Value |
|---|---|---|---|---|---|---|
| 0 | 1 | 30.76216 | 47.81601 | -65.2791 | 126.8034 | 0.5229 |
| 0 | 2 | 18.98034 | 34.46338 | -50.2414 | 88.2021 | 0.5843 |
| 2 | 1 | 11.78182 | 54.12690 | -96.9353 | 120.4989 | 0.8286 |

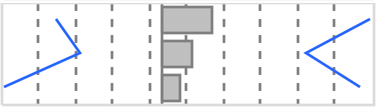

Missing Rows 2

#### Fit Group

##### Oneway Analysis of Lymphocytes By IBS-subtype Weight=0

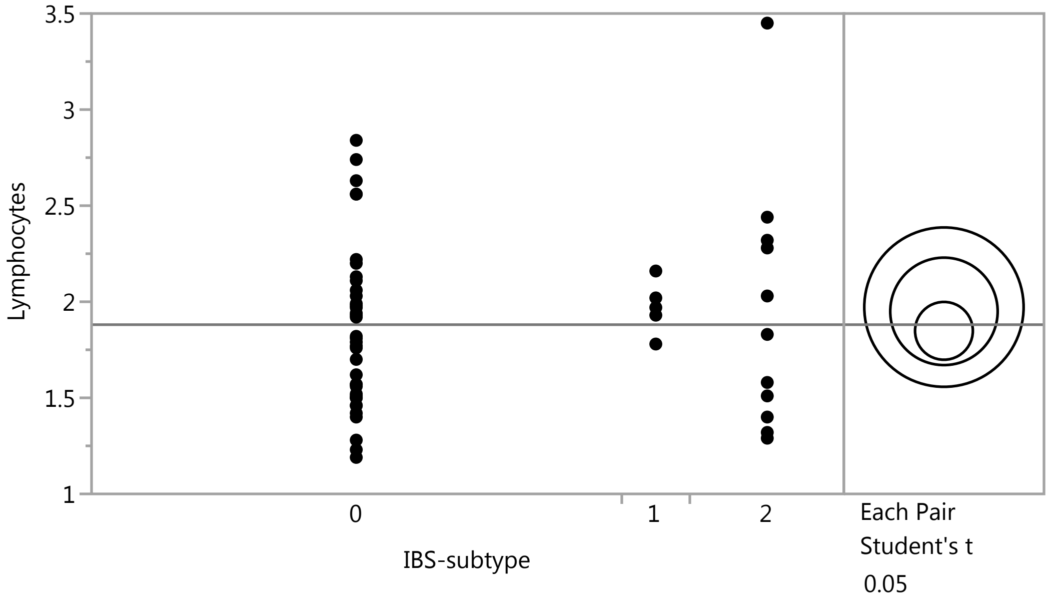


##### Means Comparisons

###### Comparisons for each pair using Student's t

**Confidence Quantile**

| t | Alpha |
|---|---|
| 2.00758 | 0.05 |

**LSD Threshold Matrix**

Abs(Dif)-LSD

| | 1 | 2 | 0 |
|---|---|---|---|
| 1 | -0.58652 | -0.47818 | -0.31785 |
| 2 | -0.47818 | -0.39543 | -0.21620 |
| 0 | -0.31785 | -0.21620 | -0.21275 |

Positive values show pairs of means that are significantly different.

**Connecting Letters Report**

| Level | | Mean |
|---|---|---|
| 1 | A | 1.9720000 |
| 2 | A | 1.9500000 |
| 0 | A | 1.8486842 |

Levels not connected by same letter are significantly different.

**Ordered Differences Report**

| Level | - Level | Difference | Std Err Dif | Lower CL | Upper CL | p-Value |
|---|---|---|---|---|---|---|
| 1 | 0 | 0.1233158 | 0.2197521 | -0.317855 | 0.5644865 | 0.5771 |
| 2 | 0 | 0.1013158 | 0.1581559 | -0.216195 | 0.4188270 | 0.5246 |
| 1 | 2 | 0.0220000 | 0.2491462 | -0.478182 | 0.5221818 | 0.9300 |

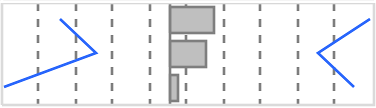

Missing Rows 1

### Fit Group

#### Oneway Analysis of Basophil_PCT By IBS-subtype Weight=0

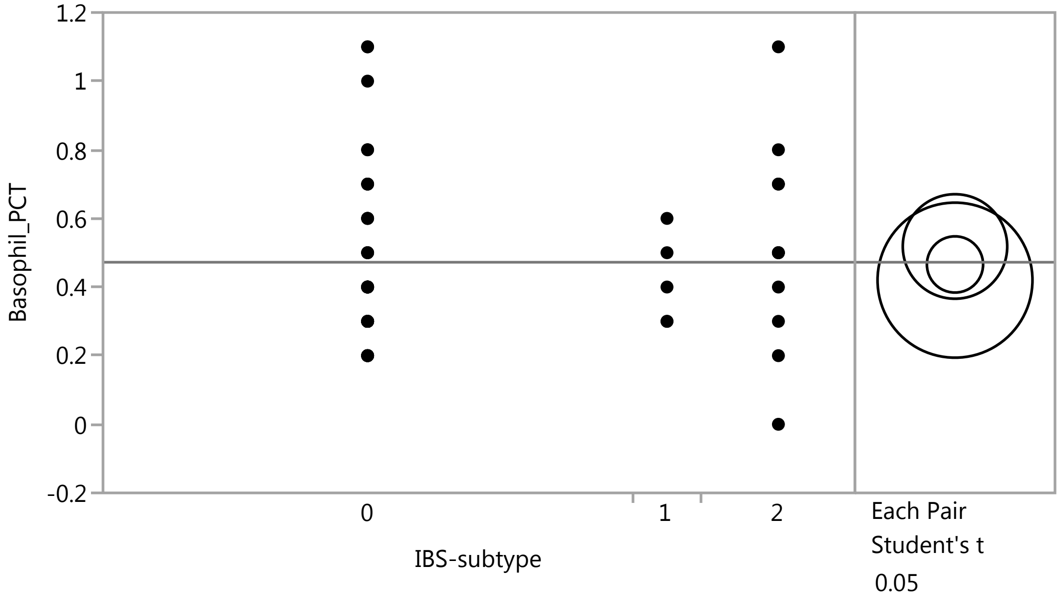


#### Means Comparisons

##### Comparisons for each pair using Student's t

###### Confidence Quantile

| t | Alpha |
|---|---|
| 2.00758 | 0.05 |

###### LSD Threshold Matrix

Abs(Dif)-LSD

| | 2 | 0 | 1 |
|---|---|---|---|
| 2 | -0.21543 | -0.12059 | -0.17431 |
| 0 | -0.12059 | -0.11591 | -0.19456 |
| 1 | -0.17431 | -0.19456 | -0.31953 |

Positive values show pairs of means that are significantly different.

###### Connecting Letters Report

| Level | | Mean |
|---|---|---|
| 2 | A | 0.51818182 |
| 0 | A | 0.46578947 |
| 1 | A | 0.42000000 |

Levels not connected by same letter are significantly different.

###### Ordered Differences Report

| Level | - Level | Difference | Std Err Dif | Lower CL | Upper CL | p-Value |
|---|---|---|---|---|---|---|
| 2 | 1 | 0.0981818 | 0.1357336 | -0.174315 | 0.3706784 | 0.4728 |
| 2 | 0 | 0.0523923 | 0.0861626 | -0.120586 | 0.2253709 | 0.5458 |
| 0 | 1 | 0.0457895 | 0.1197199 | -0.194558 | 0.2861371 | 0.7037 |

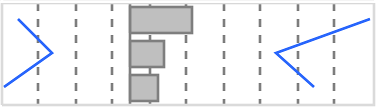

Missing Rows 1

### Fit Group

#### Oneway Analysis of Basophils By IBS-subtype Weight=0

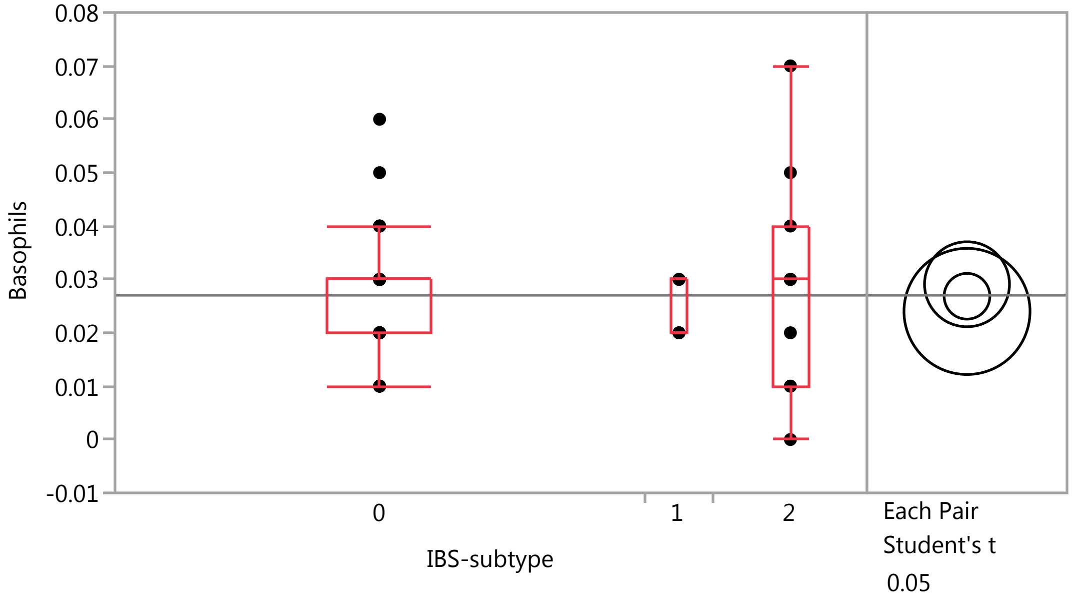


##### Quantiles

| Level | Minimum | 10% | 25% | Median | 75% | 90% | Maximum |
|---|---|---|---|---|---|---|---|
| 0 | 0.01 | 0.01 | 0.02 | 0.03 | 0.03 | 0.04 | 0.06 |
| 1 | 0.02 | 0.02 | 0.02 | 0.02 | 0.03 | 0.03 | 0.03 |
| 2 | 0 | 0.002 | 0.01 | 0.03 | 0.04 | 0.066 | 0.07 |

##### Means Comparisons

###### Comparisons for each pair using Student's t

###### Confidence Quantile

| t | Alpha |
|---|---|
| 2.00758 | 0.05 |

###### LSD Threshold Matrix

Abs(Dif)-LSD

| | 2 | 0 | 1 |
|---|---|---|---|
| 2 | -0.01127 | -0.00680 | -0.00916 |
| 0 | -0.00680 | -0.00606 | -0.00973 |
| 1 | -0.00916 | -0.00973 | -0.01671 |

Positive values show pairs of means that are significantly different.

###### Connecting Letters Report

| Level | | Mean |
|---|---|---|
| 2 | A | 0.02909091 |
| 0 | A | 0.02684211 |
| 1 | A | 0.02400000 |

Levels not connected by same letter are significantly different.

### Fit Group

#### Oneway Analysis of Basophils By IBS-subtype Weight=0

##### Means Comparisons

###### Comparisons for each pair using Student's t

###### Ordered Differences Report

| Level | - Level | Difference | Std Err Dif | Lower CL | Upper CL | p-Value |
|---|---|---|---|---|---|---|
| 2 | 1 | 0.0050909 | 0.0070978 | -0.009158 | 0.0193403 | 0.4765 |
| 0 | 1 | 0.0028421 | 0.0062604 | -0.009726 | 0.0154103 | 0.6518 |
| 2 | 0 | 0.0022488 | 0.0045056 | -0.006797 | 0.0112942 | 0.6198 |

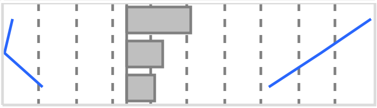

Missing Rows 1

#### Oneway Analysis of Eosinophil_PCT By IBS-subtype Weight=0

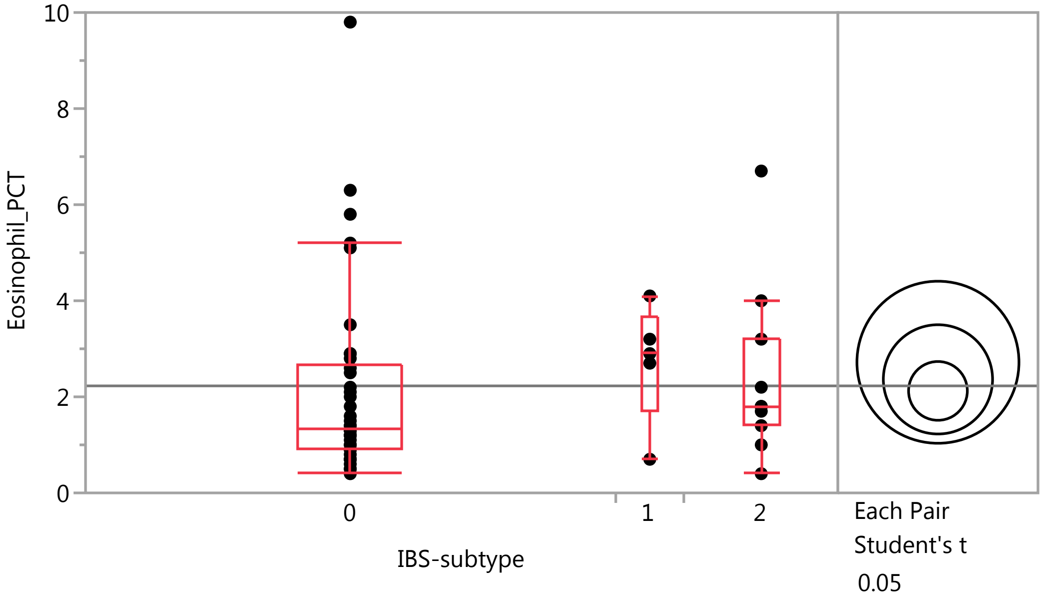


##### Quantiles

| Level | Minimum | 10% | 25% | Median | 75% | 90% | Maximum |
|---|---|---|---|---|---|---|---|
| 0 | 0.4 | 0.59 | 0.9 | 1.35 | 2.65 | 5.26 | 9.8 |
| 1 | 0.7 | 0.7 | 1.7 | 2.9 | 3.65 | 4.1 | 4.1 |
| 2 | 0.4 | 0.52 | 1.4 | 1.8 | 3.2 | 6.16 | 6.7 |

##### Means Comparisons

###### Comparisons for each pair using Student's t

###### Confidence Quantile

| t | Alpha |
|---|---|
| 2.00758 | 0.05 |

### Fit Group

#### Oneway Analysis of Eosinophil_PCT By IBS-subtype Weight=0

##### Means Comparisons

###### Comparisons for each pair using Student's t

###### LSD Threshold Matrix

Abs(Dif)-LSD

| | 1 | 2 | 0 |
|---|---|---|---|
| 1 | -2.3855 | -1.6780 | -1.1980 |
| 2 | -1.6780 | -1.6083 | -1.0514 |
| 0 | -1.1980 | -1.0514 | -0.8653 |

Positive values show pairs of means that are significantly different.

###### Connecting Letters Report

| Level | | Mean |
|---|---|---|
| 1 | A | 2.7200000 |
| 2 | A | 2.3636364 |
| 0 | A | 2.1236842 |

Levels not connected by same letter are significantly different.

###### Ordered Differences Report

| Level | - Level | Difference | Std Err Dif | Lower CL | Upper CL | p-Value |
|---|---|---|---|---|---|---|
| 1 | 0 | 0.5963158 | 0.893790 | -1.19804 | 2.390673 | 0.5077 |
| 1 | 2 | 0.3563636 | 1.013343 | -1.67801 | 2.390735 | 0.7265 |
| 2 | 0 | 0.2399522 | 0.643262 | -1.05145 | 1.531354 | 0.7107 |

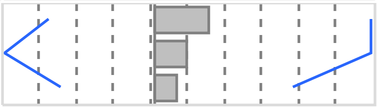

Missing Rows 1

#### Oneway Analysis of PlateletCount By IBS-subtype Weight=0

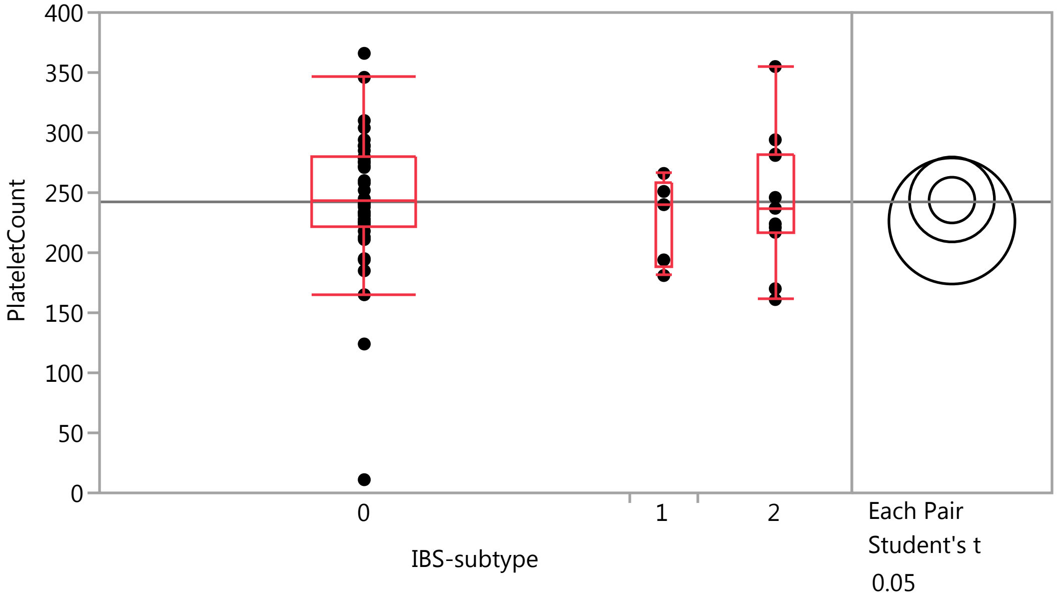

### Fit Group

#### Oneway Analysis of PlateletCount By IBS-subtype Weight=0

##### Quantiles

| Level | Minimum | 10% | 25% | Median | 75% | 90% | Maximum |
|---|---|---|---|---|---|---|---|
| 0 | 11 | 183 | 221.75 | 243 | 279.25 | 304.6 | 366 |
| 1 | 181 | 181 | 187.5 | 240 | 258.5 | 266 | 266 |
| 2 | 161 | 162.8 | 217 | 237 | 282 | 342.8 | 355 |

##### Means Comparisons

###### Comparisons for each pair using Student's t

###### Confidence Quantile

| t | Alpha |
|---|---|
| 2.00758 | 0.05 |

###### LSD Threshold Matrix

Abs(Dif)-LSD

| | 2 | 0 | 1 |
|---|---|---|---|
| 2 | -50.042 | -39.634 | -45.335 |
| 0 | -39.634 | -26.924 | -38.415 |
| 1 | -45.335 | -38.415 | -74.224 |

Positive values show pairs of means that are significantly different.

###### Connecting Letters Report

| Level | | Mean |
|---|---|---|
| 2 | A | 244.36364 |
| 0 | A | 243.81579 |
| 1 | A | 226.40000 |

Levels not connected by same letter are significantly different.

###### Ordered Differences Report

| Level | - Level | Difference | Std Err Dif | Lower CL | Upper CL | p-Value |
|---|---|---|---|---|---|---|
| 2 | 1 | 17.96364 | 31.52983 | -45.3351 | 81.26241 | 0.5714 |
| 0 | 1 | 17.41579 | 27.80996 | -38.4150 | 73.24662 | 0.5339 |
| 2 | 0 | 0.54785 | 20.01488 | -39.6337 | 40.72939 | 0.9783 |

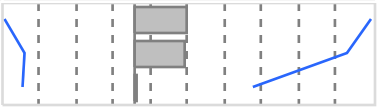

Missing Rows 1

#### Oneway Analysis of IgE By IBS-subtype Weight=0

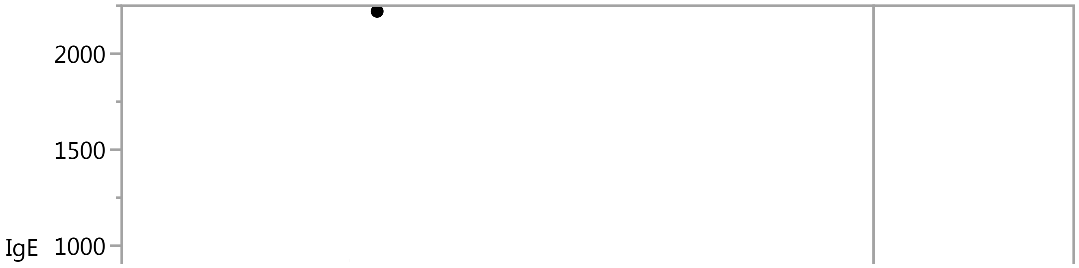

#### Fit Group

##### Oneway Analysis of IgE By IBS-subtype Weight=0

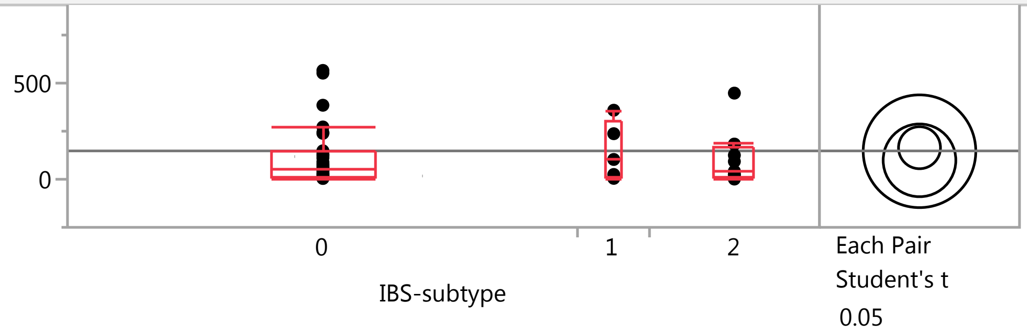


###### Quantiles

| Level | Minimum | 10% | 25% | Median | 75% | 90% | Maximum |
|---|---|---|---|---|---|---|---|
| 0 | 4.2 | 7.23 | 10.65 | 53.95 | 142 | 435.1 | 2221 |
| 1 | 4.8 | 4.8 | 14.85 | 103 | 298 | 359 | 359 |
| 2 | 1.1 | 3.17 | 14.975 | 40.7 | 167.75 | 368.5 | 448 |

###### Means Comparisons

###### Comparisons for each pair using Student's t

###### Confidence Quantile

| t | Alpha |
|---|---|
| 2.00856 | 0.05 |

###### LSD Threshold Matrix

Abs(Dif)-LSD

| | 0 | 1 | 2 |
|---|---|---|---|
| 0 | -154.51 | -294.55 | -152.96 |
| 1 | -294.55 | -414.61 | -301.70 |
| 2 | -152.96 | -301.70 | -267.63 |

Positive values show pairs of means that are significantly different.

###### Connecting Letters Report

| Level | | Mean |
|---|---|---|
| 0 | A | 164.05556 |
| 1 | A | 145.74000 |
| 2 | A | 98.50000 |

Levels not connected by same letter are significantly different.

###### Ordered Differences Report

| Level | - Level | Difference | Std Err Dif | Lower CL | Upper CL | p-Value |
|---|---|---|---|---|---|---|
| 0 | 2 | 65.55556 | 108.7926 | -152.961 | 284.0720 | 0.5495 |
| 1 | 2 | 47.24000 | 173.7279 | -301.703 | 396.1827 | 0.7868 |
| 0 | 1 | 18.31556 | 155.7673 | -294.552 | 331.1835 | 0.9069 |

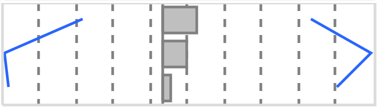

Missing Rows 2

#### Fit Group

##### Oneway Analysis of ESR By IBS-subtype Weight=0

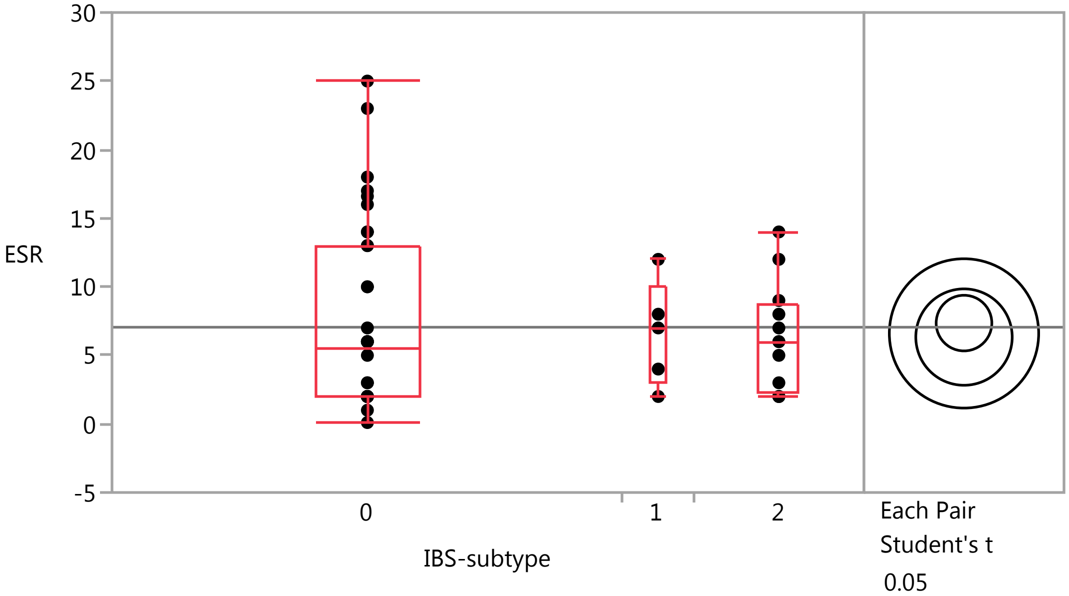


###### Quantiles

| Level | Minimum | 10% | 25% | Median | 75% | 90% | Maximum |
|---|---|---|---|---|---|---|---|
| 0 | 0.1 | 2 | 2 | 5.5 | 13 | 17.3 | 25 |
| 1 | 2 | 2 | 3 | 7 | 10 | 12 | 12 |
| 2 | 2 | 2 | 2.25 | 6 | 8.75 | 13.4 | 14 |

###### Means Comparisons

###### Comparisons for each pair using Student's t

###### Confidence Quantile

| t | Alpha |
|---|---|
| 2.00856 | 0.05 |

###### LSD Threshold Matrix

Abs(Dif)-LSD

| | 0 | 1 | 2 |
|---|---|---|---|
| 0 | -2.8697 | -5.0579 | -3.0389 |
| 1 | -5.0579 | -7.7002 | -6.2140 |
| 2 | -3.0389 | -6.2140 | -4.9705 |

Positive values show pairs of means that are significantly different.

###### Connecting Letters Report

| Level | | Mean |
|---|---|---|
| 0 | A | 7.3527778 |
| 1 | A | 6.6000000 |
| 2 | A | 6.3333333 |

Levels not connected by same letter are significantly different.

### Fit Group

#### Oneway Analysis of ESR By IBS-subtype Weight=0

##### Means Comparisons

###### Comparisons for each pair using Student's t

###### Ordered Differences Report

| Level | - Level | Difference | Std Err Dif | Lower CL | Upper CL | p-Value |
|---|---|---|---|---|---|---|
| 0 | 2 | 1.019444 | 2.020537 | -3.03892 | 5.077812 | 0.6161 |
| 0 | 1 | 0.752778 | 2.892969 | -5.05792 | 6.563476 | 0.7958 |
| 1 | 2 | 0.266667 | 3.226539 | -6.21403 | 6.747361 | 0.9345 |

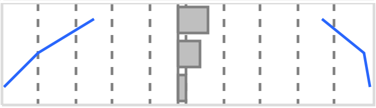

Missing Rows 2

#### Oneway Analysis of IgG By IBS-subtype Weight=0

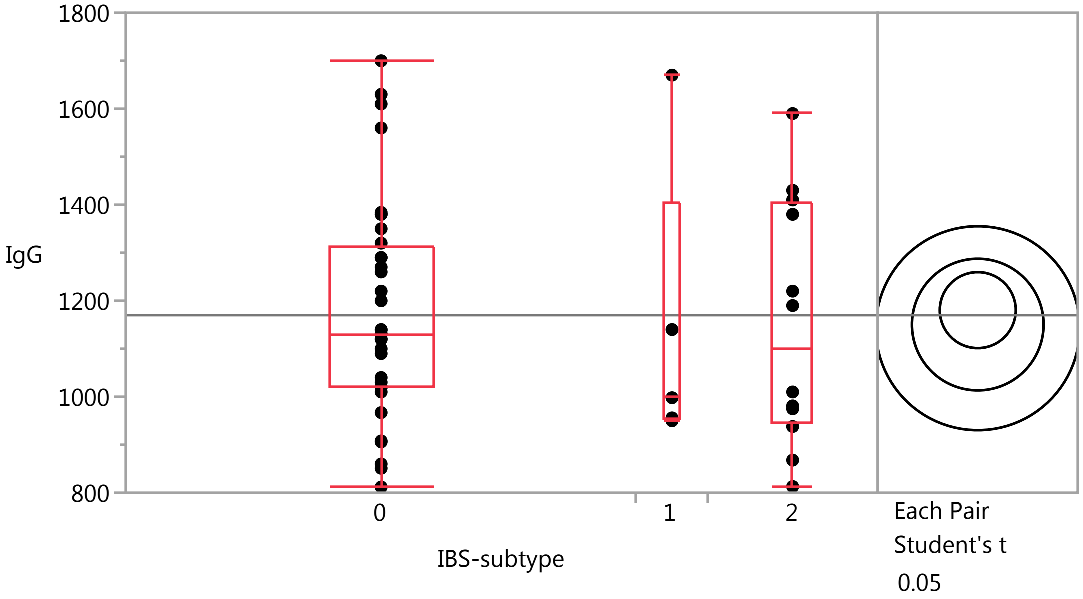


##### Quantiles

| Level | Minimum | 10% | 25% | Median | 75% | 90% | Maximum |
|---|---|---|---|---|---|---|---|
| 0 | 812 | 892.2 | 1022.5 | 1130 | 1312.5 | 1575 | 1700 |
| 1 | 950 | 950 | 953 | 998 | 1405 | 1670 | 1670 |
| 2 | 813 | 829.5 | 947.25 | 1100 | 1402.5 | 1542 | 1590 |

##### Means Comparisons

###### Comparisons for each pair using Student's t

###### Confidence Quantile

| t | Alpha |
|---|---|
| 2.00856 | 0.05 |

### Fit Group

#### Oneway Analysis of IgG By IBS-subtype Weight=0

##### Means Comparisons

###### Comparisons for each pair using Student's t

###### LSD Threshold Matrix

Abs(Dif)-LSD

| | 0 | 2 | 1 |
|---|---|---|---|
| 0 | -111.98 | -128.31 | -189.07 |
| 2 | -128.31 | -193.96 | -245.27 |
| 1 | -189.07 | -245.27 | -300.48 |

Positive values show pairs of means that are significantly different.

###### Connecting Letters Report

| Level | | Mean |
|---|---|---|
| 0 | A | 1180.4722 |
| 2 | A | 1150.4167 |
| 1 | A | 1142.8000 |

Levels not connected by same letter are significantly different.

###### Ordered Differences Report

| Level | - Level | Difference | Std Err Dif | Lower CL | Upper CL | p-Value |
|---|---|---|---|---|---|---|
| 0 | 1 | 37.67222 | 112.8898 | -189.074 | 264.4180 | 0.7400 |
| 0 | 2 | 30.05556 | 78.8456 | -128.311 | 188.4217 | 0.7047 |
| 2 | 1 | 7.61667 | 125.9064 | -245.274 | 260.5070 | 0.9520 |

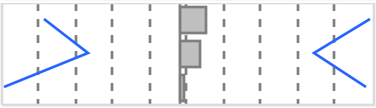

Missing Rows 2

#### Oneway Analysis of CRP By IBS-subtype Weight=0

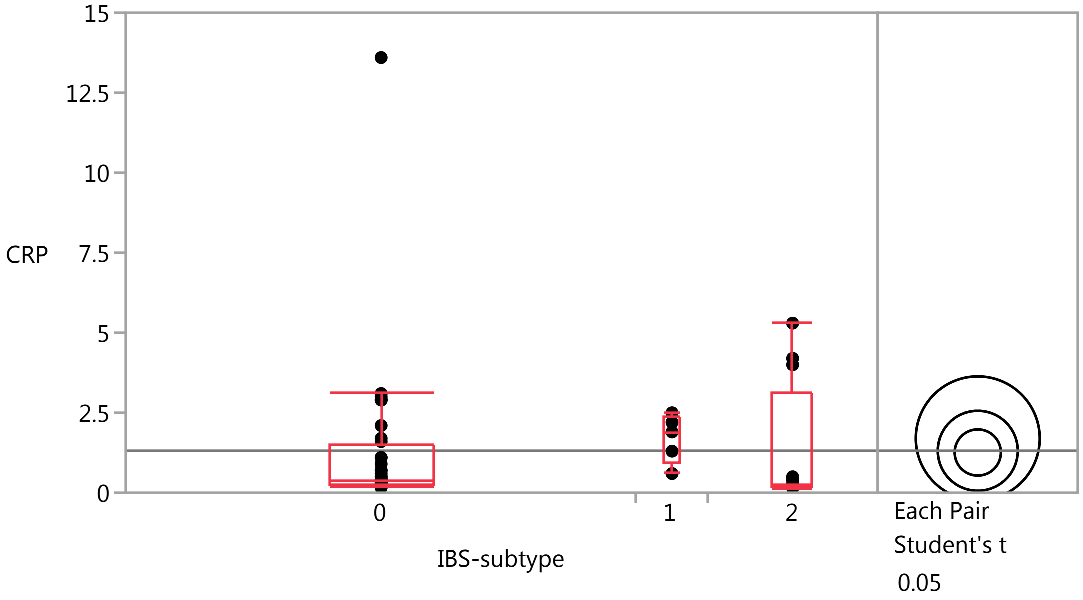

### Fit Group

#### Oneway Analysis of CRP By IBS-subtype Weight=0

##### Quantiles

| Level | Minimum | 10% | 25% | Median | 75% | 90% | Maximum |
|---|---|---|---|---|---|---|---|
| 0 | 0.16 | 0.2 | 0.255 | 0.4 | 1.475 | 2.93 | 13.6 |
| 1 | 0.6 | 0.6 | 0.95 | 1.9 | 2.35 | 2.5 | 2.5 |
| 2 | 0.15 | 0.153 | 0.17 | 0.24 | 3.125 | 4.97 | 5.3 |

##### Means Comparisons

###### Comparisons for each pair using Student's t

###### Confidence Quantile

| t | Alpha |
|---|---|
| 2.00856 | 0.05 |

###### LSD Threshold Matrix

Abs(Dif)-LSD

| | 1 | 2 | 0 |
|---|---|---|---|
| 1 | -2.7390 | -1.9177 | -1.6252 |
| 2 | -1.9177 | -1.7680 | -1.3894 |
| 0 | -1.6252 | -1.3894 | -1.0208 |

Positive values show pairs of means that are significantly different.

###### Connecting Letters Report

| Level | | Mean |
|---|---|---|
| 1 | A | 1.7000000 |
| 2 | A | 1.3125000 |
| 0 | A | 1.2583333 |

Levels not connected by same letter are significantly different.

###### Ordered Differences Report

| Level | - Level | Difference | Std Err Dif | Lower CL | Upper CL | p-Value |
|---|---|---|---|---|---|---|
| 1 | 0 | 0.4416667 | 1.029050 | -1.62524 | 2.508574 | 0.6696 |
| 1 | 2 | 0.3875000 | 1.147703 | -1.91773 | 2.692729 | 0.7371 |
| 2 | 0 | 0.0541667 | 0.718719 | -1.38942 | 1.497757 | 0.9402 |

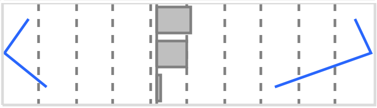

Missing Rows 2

#### Oneway Analysis of RBC By IBS-subtype Weight=0

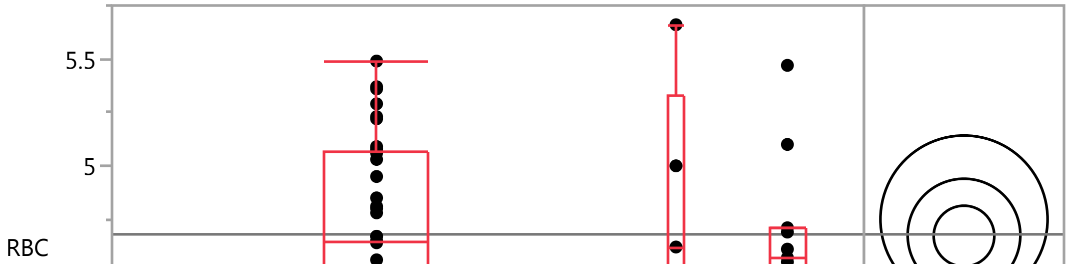

#### Fit Group

##### Oneway Analysis of RBC By IBS-subtype Weight=0

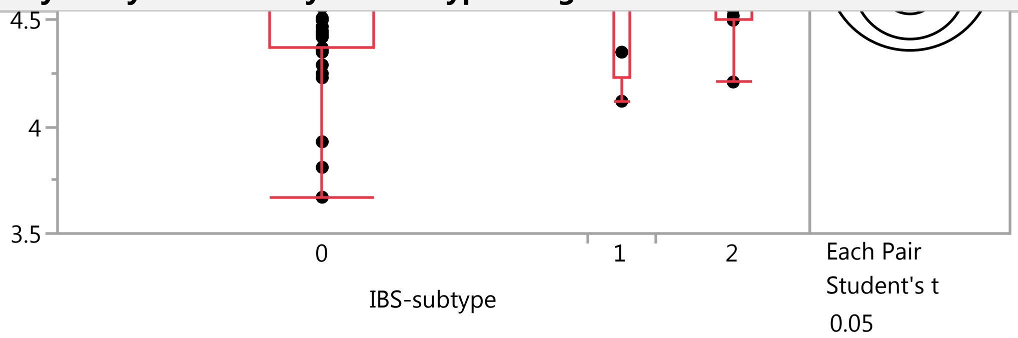


##### Quantiles

| Level | Minimum | 10% | 25% | Median | 75% | 90% | Maximum |
|---|---|---|---|---|---|---|---|
| 0 | 3.67 | 4.2 | 4.3675 | 4.645 | 5.0625 | 5.297 | 5.49 |
| 1 | 4.12 | 4.12 | 4.235 | 4.62 | 5.33 | 5.66 | 5.66 |
| 2 | 4.21 | 4.268 | 4.5 | 4.57 | 4.71 | 5.396 | 5.47 |

##### Means Comparisons

###### Comparisons for each pair using Student's t

###### Confidence Quantile

| t | Alpha |
|---|---|
| 2.00758 | 0.05 |

###### LSD Threshold Matrix

Abs(Dif)-LSD

| | 1 | 2 | 0 |
|---|---|---|---|
| 1 | -0.55405 | -0.39795 | -0.33807 |
| 2 | -0.39795 | -0.37354 | -0.29580 |
| 0 | -0.33807 | -0.29580 | -0.20097 |

Positive values show pairs of means that are significantly different.

###### Connecting Letters Report

| Level | | Mean |
|---|---|---|
| 1 | A | 4.7500000 |
| 2 | A | 4.6754545 |
| 0 | A | 4.6713158 |

Levels not connected by same letter are significantly different.

###### Ordered Differences Report

| Level | - Level | Difference | Std Err Dif | Lower CL | Upper CL | p-Value |
|---|---|---|---|---|---|---|
| 1 | 0 | 0.0786842 | 0.2075880 | -0.338066 | 0.4954345 | 0.7062 |
| 1 | 2 | 0.0745455 | 0.2353550 | -0.397949 | 0.5470404 | 0.7527 |
| 2 | 0 | 0.0041388 | 0.1494014 | -0.295797 | 0.3040746 | 0.9780 |

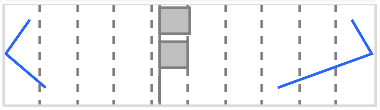

Missing Rows 1

#### Fit Group

##### Oneway Analysis of HCT By IBS-subtype Weight=0

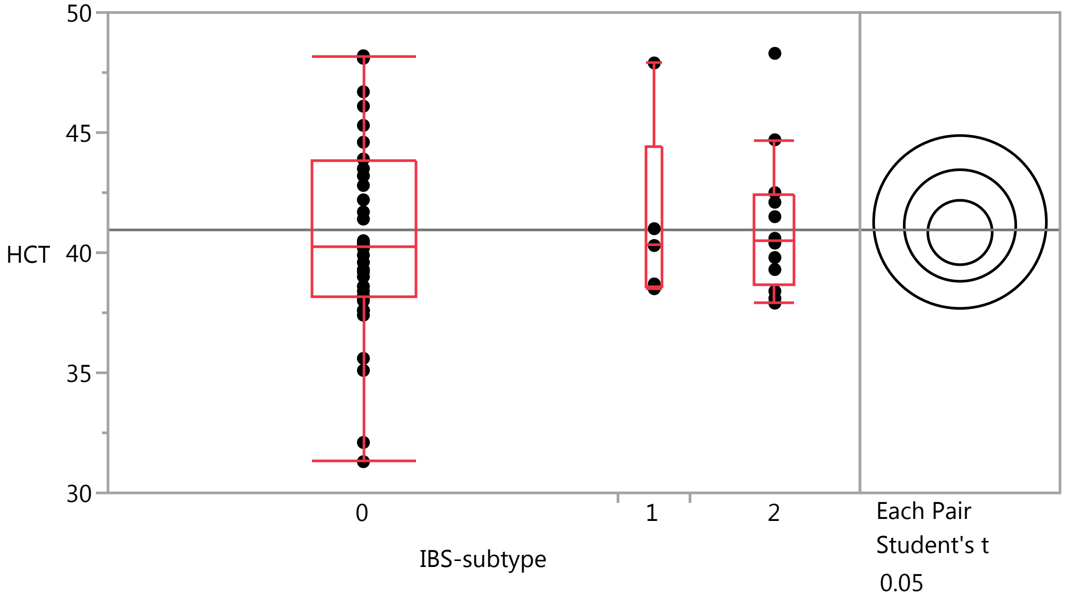


##### Quantiles

| Level | Minimum | 10% | 25% | Median | 75% | 90% | Maximum |
|---|---|---|---|---|---|---|---|
| 0 | 31.3 | 35.45 | 38.125 | 40.25 | 43.8 | 48.1 | 48.2 |
| 1 | 38.5 | 38.5 | 38.6 | 40.3 | 44.45 | 47.9 | 47.9 |
| 2 | 37.9 | 37.96 | 38.625 | 40.5 | 42.4 | 47.22 | 48.3 |

##### Means Comparisons

###### Comparisons for each pair using Student's t

###### Confidence Quantile

| t | Alpha |
|---|---|
| 2.00856 | 0.05 |

###### LSD Threshold Matrix

Abs(Dif)-LSD

| | 1 | 2 | 0 |
|---|---|---|---|
| 1 | -5.0891 | -4.1365 | -3.4020 |
| 2 | -4.1365 | -3.2850 | -2.3905 |
| 0 | -3.4020 | -2.3905 | -1.8966 |

Positive values show pairs of means that are significantly different.

###### Connecting Letters Report

| Level | | Mean |
|---|---|---|
| 1 | A | 41.280000 |
| 2 | A | 41.133333 |
| 0 | A | 40.841667 |

Levels not connected by same letter are significantly different.

#### Fit Group

##### Oneway Analysis of HCT By IBS-subtype Weight=0

###### Means Comparisons

###### Comparisons for each pair using Student's t

###### Ordered Differences Report

| Level | - Level | Difference | Std Err Dif | Lower CL | Upper CL | p-Value |
|---|---|---|---|---|---|---|
| 1 | 0 | 0.4383333 | 1.911978 | -3.40199 | 4.278655 | 0.8196 |
| 2 | 0 | 0.2916667 | 1.335384 | -2.39053 | 2.973863 | 0.8280 |
| 1 | 2 | 0.1466667 | 2.132437 | -4.13646 | 4.429792 | 0.9454 |

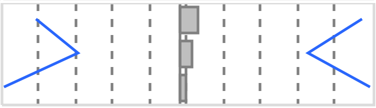

Missing Rows 2

#### Fit Group

##### Oneway Analysis of MCH By IBS-subtype Weight=1

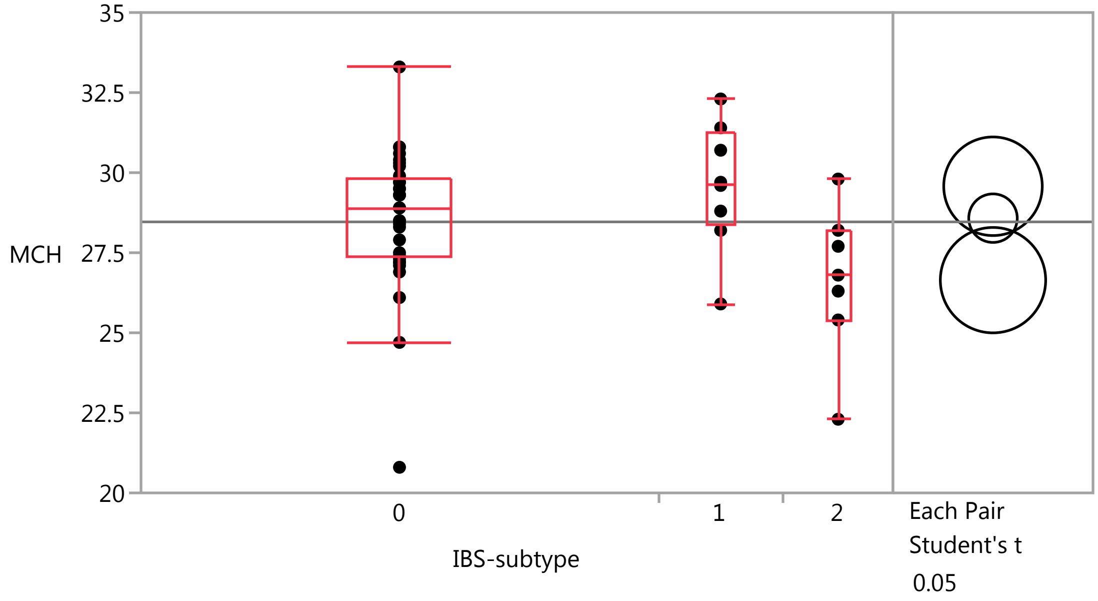


###### Quantiles

| Level | Minimum | 10% | 25% | Median | 75% | 90% | Maximum |
|---|---|---|---|---|---|---|---|
| 0 | 20.8 | 26.42 | 27.35 | 28.9 | 29.8 | 30.72 | 33.3 |
| 1 | 25.9 | 25.9 | 28.35 | 29.65 | 31.225 | 32.3 | 32.3 |
| 2 | 22.3 | 22.3 | 25.4 | 26.8 | 28.2 | 29.8 | 29.8 |

###### Means Comparisons

###### Comparisons for each pair using Student's t

###### Confidence Quantile

| t | Alpha |
|---|---|
| 2.01410 | 0.05 |

### Fit Group

#### Oneway Analysis of MCH By IBS-subtype Weight=1

##### Means Comparisons

###### Comparisons for each pair using Student's t

###### LSD Threshold Matrix

Abs(Dif)-LSD

| | 1 | 0 | 2 |
|---|---|---|---|
| 1 | -2.1773 | -0.7199 | 0.6784 |
| 0 | -0.7199 | -1.0720 | 0.1239 |
| 2 | 0.6784 | 0.1239 | -2.3276 |

Positive values show pairs of means that are significantly different.

###### Connecting Letters Report

| Level | | | Mean |
|---|---|---|---|
| 1 | A | | 29.575000 |
| 0 | A | | 28.578788 |
| 2 | | B | 26.642857 |

Levels not connected by same letter are significantly different.

###### Ordered Differences Report

| Level | - Level | Difference | Std Err Dif | Lower CL | Upper CL | p-Value |
|---|---|---|---|---|---|---|
| 1 | 2 | 2.932143 | 1.118958 | 0.678446 | 5.185840 | 0.0119* |
| 0 | 2 | 1.935931 | 0.899676 | 0.123890 | 3.747972 | 0.0368* |
| 1 | 0 | 0.996212 | 0.852025 | -0.719854 | 2.712278 | 0.2485 |

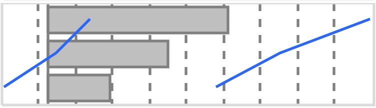

Missing Rows 8

#### Oneway Analysis of LBP By IBS-subtype Weight=1

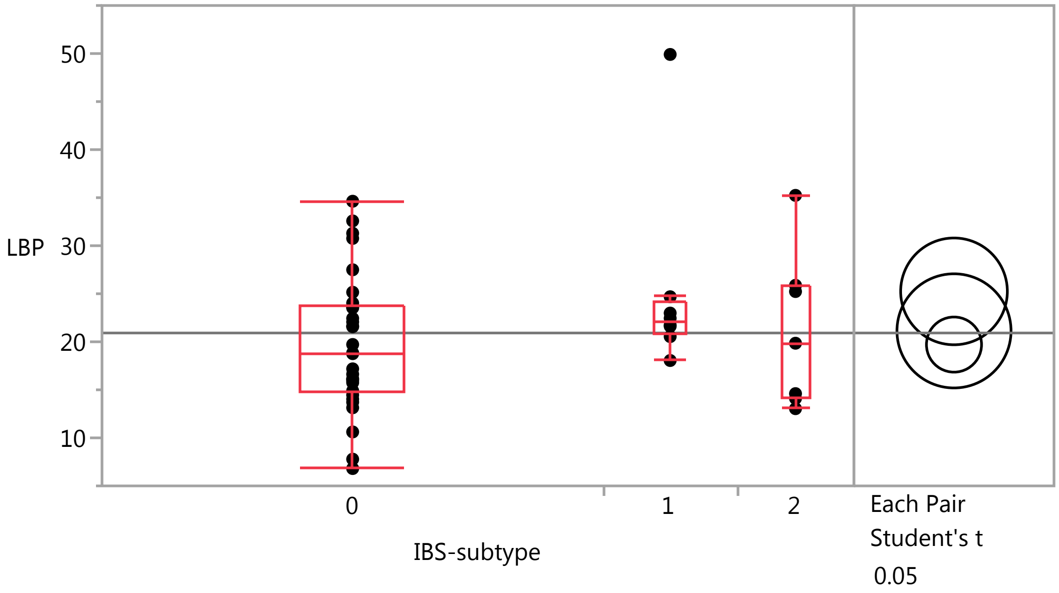

#### Fit Group

##### Oneway Analysis of LBP By IBS-subtype Weight=1

###### Quantiles

| Level | Minimum | 10% | 25% | Median | 75% | 90% | Maximum |
|---|---|---|---|---|---|---|---|
| 0 | 6.808 | 10.8687 | 14.7905 | 18.802 | 23.7075 | 31.2388 | 34.628 |
| 1 | 18.052 | 18.052 | 20.81 | 22.0925 | 24.26275 | 49.906 | 49.906 |
| 2 | 13.039 | 13.039 | 14.11 | 19.85 | 25.877 | 35.239 | 35.239 |

###### Means Comparisons

###### Comparisons for each pair using Student's t

###### Confidence Quantile

| t | Alpha |
|---|---|
| 2.01808 | 0.05 |

###### LSD Threshold Matrix

Abs(Dif)-LSD

| | 1 | 2 | 0 |
|---|---|---|---|
| 1 | -7.8592 | -4.0241 | -0.7175 |
| 2 | -4.0241 | -8.4018 | -5.1717 |
| 0 | -0.7175 | -5.1717 | -4.0585 |

Positive values show pairs of means that are significantly different.

###### Connecting Letters Report

| Level | | Mean |
|---|---|---|
| 1 | A | 25.246750 |
| 2 | A | 21.135857 |
| 0 | A | 19.709733 |

Levels not connected by same letter are significantly different.

###### Ordered Differences Report

| Level | - Level | Difference | Std Err Dif | Lower CL | Upper CL | p-Value |
|---|---|---|---|---|---|---|
| 1 | 0 | 5.537017 | 3.099230 | -0.71748 | 11.79152 | 0.0812 |
| 1 | 2 | 4.110893 | 4.031060 | -4.02412 | 12.24590 | 0.3137 |
| 2 | 0 | 1.426124 | 3.269331 | -5.17165 | 8.02390 | 0.6649 |

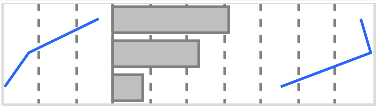

Missing Rows 11

##### Oneway Analysis of Lymphocytes_PCT By IBS-subtype Weight=1

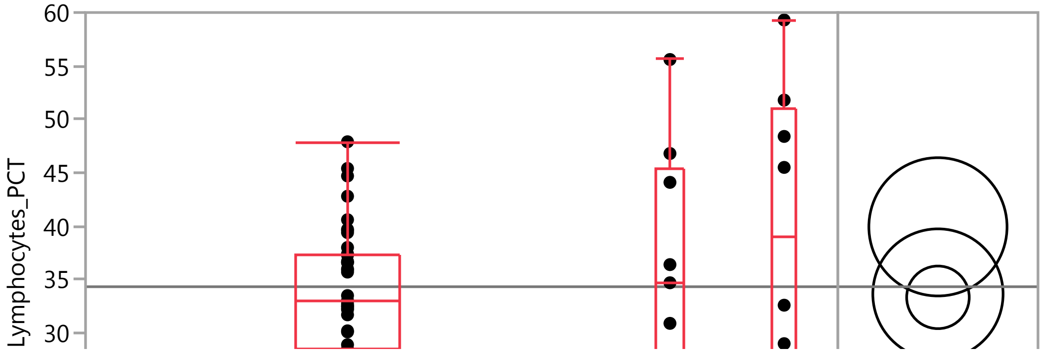

#### Fit Group

##### Oneway Analysis of Lymphocytes_PCT By IBS-subtype Weight=1

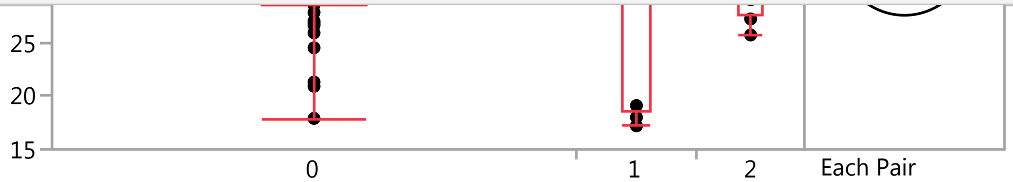


##### Quantiles

| Level | Minimum | 10% | 25% | Median | 75% | 90% | Maximum |
|---|---|---|---|---|---|---|---|
| 0 | 17.9 | 24.5 | 28.5 | 33 | 37.4 | 42.8 | 47.9 |
| 1 | 17.2 | 17.2 | 18.55 | 34.7 | 45.45 | 55.6 | 55.6 |
| 2 | 25.7 | 25.7 | 27.65 | 39.05 | 50.95 | 59.3 | 59.3 |

##### Means Comparisons

###### Comparisons for each pair using Student's t

###### Confidence Quantile

| t | Alpha |
|---|---|
| 2.00575 | 0.05 |

###### LSD Threshold Matrix

Abs(Dif)-LSD

| | 2 | 1 | 0 |
|---|---|---|---|
| 2 | -9.1542 | -2.6033 | -0.5044 |
| 1 | -2.6033 | -8.6307 | -6.4619 |
| 0 | -0.5044 | -6.4619 | -4.1461 |

Positive values show pairs of means that are significantly different.

###### Connecting Letters Report

| Level | | Mean |
|---|---|---|
| 2 | A | 39.937500 |
| 1 | A | 33.644444 |
| 0 | A | 33.335897 |

Levels not connected by same letter are significantly different.

###### Ordered Differences Report

| Level | - Level | Difference | Std Err Dif | Lower CL | Upper CL | p-Value |
|---|---|---|---|---|---|---|
| 2 | 0 | 6.601603 | 3.542809 | -0.50437 | 13.70758 | 0.0680 |
| 2 | 1 | 6.293056 | 4.435416 | -2.60326 | 15.18937 | 0.1618 |
| 1 | 0 | 0.308547 | 3.375540 | -6.46193 | 7.07902 | 0.9275 |

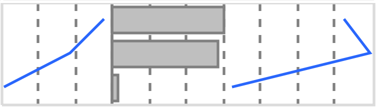

#### Fit Group

##### Oneway Analysis of Neutrophil_PCT By IBS-subtype Weight=1

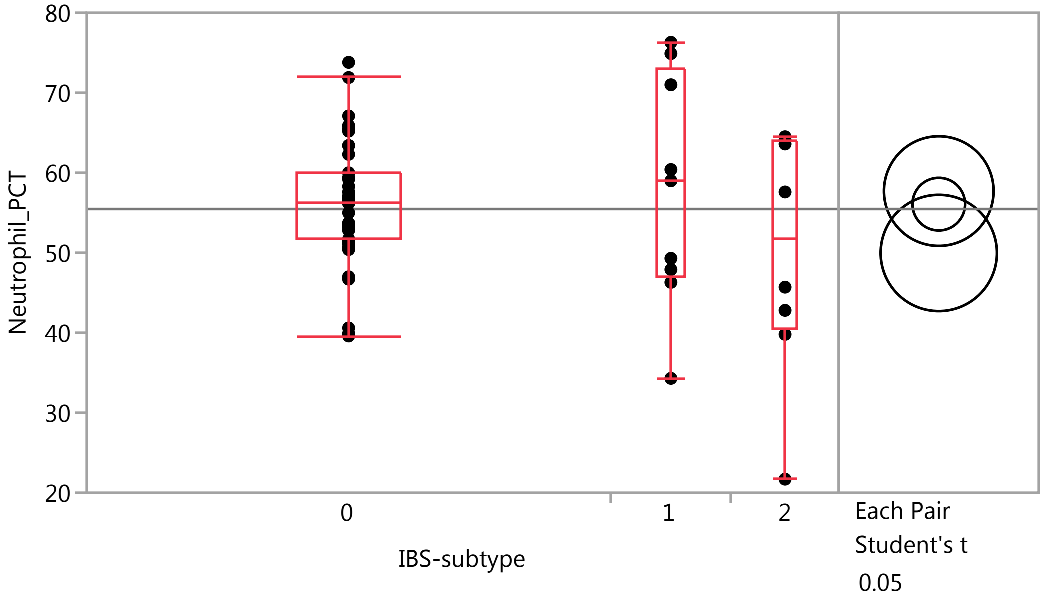


##### Quantiles

| Level | Minimum | 10% | 25% | Median | 75% | 90% | Maximum |
|---|---|---|---|---|---|---|---|
| 0 | 39.6 | 46.7 | 51.7 | 56.2 | 60 | 65.9 | 73.8 |
| 1 | 34.3 | 34.3 | 47.1 | 59 | 72.95 | 76.3 | 76.3 |
| 2 | 21.7 | 21.7 | 40.55 | 51.65 | 63.975 | 64.5 | 64.5 |

##### Means Comparisons

###### Comparisons for each pair using Student's t

###### Confidence Quantile

| t | Alpha |
|---|---|
| 2.00575 | 0.05 |

###### LSD Threshold Matrix

Abs(Dif)-LSD

| | 1 | 0 | 2 |
|---|---|---|---|
| 1 | -9.692 | -5.969 | -2.254 |
| 0 | -5.969 | -4.656 | -1.878 |
| 2 | -2.254 | -1.878 | -10.280 |

Positive values show pairs of means that are significantly different.

###### Connecting Letters Report

| Level | | Mean |
|---|---|---|
| 1 | A | 57.711111 |
| 0 | A | 56.076923 |
| 2 | A | 49.975000 |

Levels not connected by same letter are significantly different.

#### Fit Group

##### Oneway Analysis of Neutrophil_PCT By IBS-subtype Weight=1

###### Means Comparisons

###### Comparisons for each pair using Student's t

###### Ordered Differences Report

| Level | - Level | Difference | Std Err Dif | Lower CL | Upper CL | p-Value |
|---|---|---|---|---|---|---|
| 1 | 2 | 7.736111 | 4.980686 | -2.25388 | 17.72610 | 0.1263 |
| 0 | 2 | 6.101923 | 3.978346 | -1.87763 | 14.08147 | 0.1310 |
| 1 | 0 | 1.634188 | 3.790513 | -5.96862 | 9.23699 | 0.6681 |

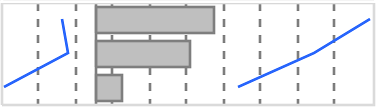

##### Oneway Analysis of Neutrophils By IBS-subtype Weight=1

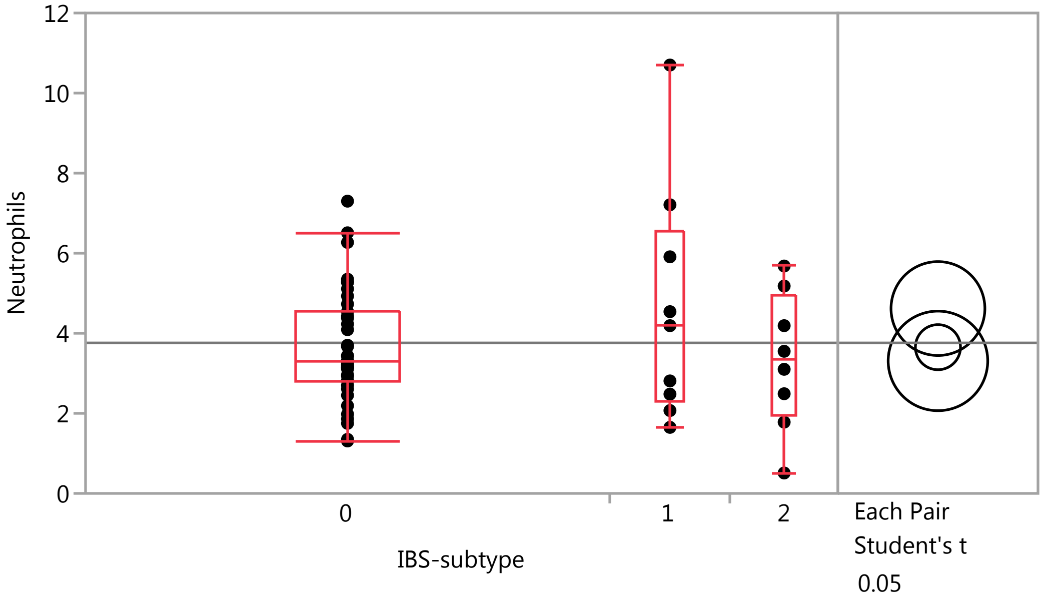


###### Quantiles

| Level | Minimum | 10% | 25% | Median | 75% | 90% | Maximum |
|---|---|---|---|---|---|---|---|
| 0 | 1.31 | 1.86 | 2.82 | 3.3 | 4.54 | 5.35 | 7.3 |
| 1 | 1.65 | 1.65 | 2.275 | 4.19 | 6.56 | 10.7 | 10.7 |
| 2 | 0.51 | 0.51 | 1.9575 | 3.325 | 4.9325 | 5.68 | 5.68 |

###### Means Comparisons

###### Comparisons for each pair using Student's t

###### Confidence Quantile

| t | Alpha |
|---|---|
| 2.00575 | 0.05 |

### Fit Group

#### Oneway Analysis of Neutrophils By IBS-subtype Weight=1

##### Means Comparisons

###### Comparisons for each pair using Student's t

###### LSD Threshold Matrix

Abs(Dif)-LSD

| | 1 | 0 | 2 |
|---|---|---|---|
| 1 | -1.6583 | -0.3357 | -0.4016 |
| 0 | -0.3357 | -0.7966 | -1.0228 |
| 2 | -0.4016 | -1.0228 | -1.7589 |

Positive values show pairs of means that are significantly different.

###### Connecting Letters Report

| Level | | Mean |
|---|---|---|
| 1 | A | 4.6177778 |
| 0 | A | 3.6525641 |
| 2 | A | 3.3100000 |

Levels not connected by same letter are significantly different.

###### Ordered Differences Report

| Level | - Level | Difference | Std Err Dif | Lower CL | Upper CL | p-Value |
|---|---|---|---|---|---|---|
| 1 | 2 | 1.307778 | 0.8522253 | -0.40157 | 3.017125 | 0.1308 |
| 1 | 0 | 0.965214 | 0.6485796 | -0.33567 | 2.266100 | 0.1426 |
| 0 | 2 | 0.342564 | 0.6807190 | -1.02279 | 1.707913 | 0.6169 |

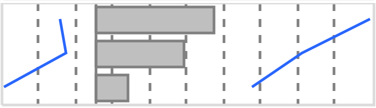

#### Oneway Analysis of IgM By IBS-subtype Weight=1

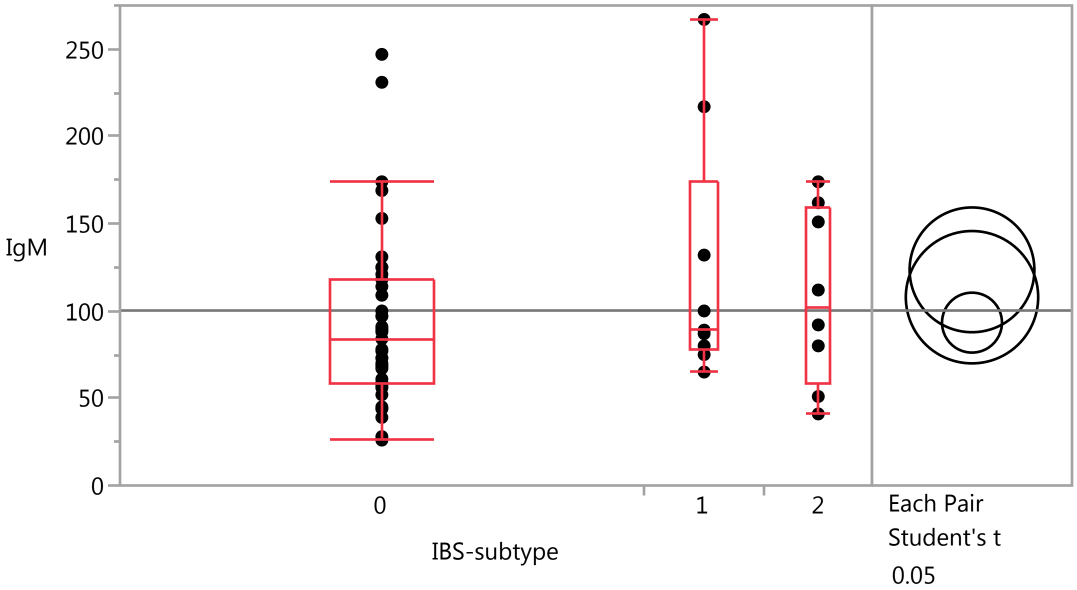

### Fit Group

#### Oneway Analysis of IgM By IBS-subtype Weight=1

##### Quantiles

| Level | Minimum | 10% | 25% | Median | 75% | 90% | Maximum |
|---|---|---|---|---|---|---|---|
| 0 | 26 | 44 | 59 | 84 | 118 | 169 | 247 |
| 1 | 65 | 65 | 77.5 | 89 | 174.5 | 267 | 267 |
| 2 | 41 | 41 | 58.25 | 102 | 159.25 | 174 | 174 |

##### Means Comparisons

###### Comparisons for each pair using Student's t

###### Confidence Quantile

| t | Alpha |
|---|---|
| 2.00575 | 0.05 |

###### LSD Threshold Matrix

Abs(Dif)-LSD

| | 1 | 2 | 0 |
|---|---|---|---|
| 1 | -50.555 | -36.431 | -9.411 |
| 2 | -36.431 | -53.622 | -27.057 |
| 0 | -9.411 | -27.057 | -24.286 |

Positive values show pairs of means that are significantly different.

###### Connecting Letters Report

| Level | | Mean |
|---|---|---|
| 1 | A | 123.55556 |
| 2 | A | 107.87500 |
| 0 | A | 93.30769 |

Levels not connected by same letter are significantly different.

###### Ordered Differences Report

| Level | - Level | Difference | Std Err Dif | Lower CL | Upper CL | p-Value |
|---|---|---|---|---|---|---|
| 1 | 0 | 30.24786 | 19.77258 | -9.4109 | 69.90663 | 0.1320 |
| 1 | 2 | 15.68056 | 25.98091 | -36.4306 | 67.79167 | 0.5487 |
| 2 | 0 | 14.56731 | 20.75238 | -27.0567 | 56.19130 | 0.4858 |

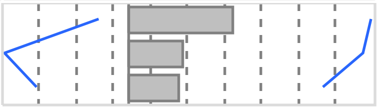

#### Oneway Analysis of SerumCortisol By IBS-subtype Weight=1

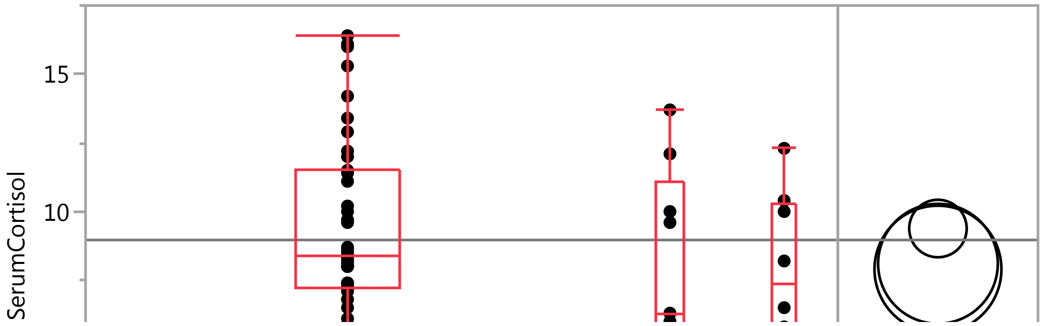

#### Fit Group

##### Oneway Analysis of SerumCortisol By IBS-subtype Weight=1

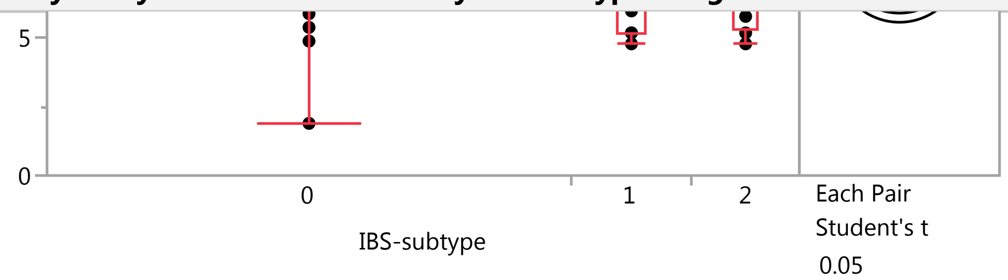


##### Quantiles

| Level | Minimum | 10% | 25% | Median | 75% | 90% | Maximum |
|---|---|---|---|---|---|---|---|
| 0 | 1.9 | 5.9 | 7.2 | 8.4 | 11.5 | 15.3 | 16.4 |
| 1 | 4.8 | 4.8 | 5.2 | 6.3 | 11.05 | 13.7 | 13.7 |
| 2 | 4.8 | 4.8 | 5.35 | 7.35 | 10.3 | 12.3 | 12.3 |

##### Means Comparisons

###### Comparisons for each pair using Student's t

###### Confidence Quantile

| t | Alpha |
|---|---|
| 2.00575 | 0.05 |

###### LSD Threshold Matrix

Abs(Dif)-LSD

| | 0 | 1 | 2 |
|---|---|---|---|
| 0 | -1.4815 | -1.1373 | -1.0572 |
| 1 | -1.1373 | -3.0841 | -2.9790 |
| 2 | -1.0572 | -2.9790 | -3.2711 |

Positive values show pairs of means that are significantly different.

###### Connecting Letters Report

| Level | | Mean |
|---|---|---|
| 0 | A | 9.3820513 |
| 1 | A | 8.1000000 |
| 2 | A | 7.9000000 |

Levels not connected by same letter are significantly different.

###### Ordered Differences Report

| Level | - Level | Difference | Std Err Dif | Lower CL | Upper CL | p-Value |
|---|---|---|---|---|---|---|
| 0 | 2 | 1.482051 | 1.265978 | -1.05718 | 4.021281 | 0.2470 |
| 0 | 1 | 1.282051 | 1.206206 | -1.13729 | 3.701394 | 0.2927 |
| 1 | 2 | 0.200000 | 1.584939 | -2.97899 | 3.378986 | 0.9001 |

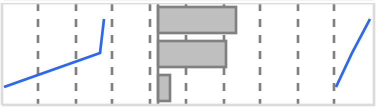

### Fit Group

#### Oneway Analysis of WBC By IBS-subtype Weight=1

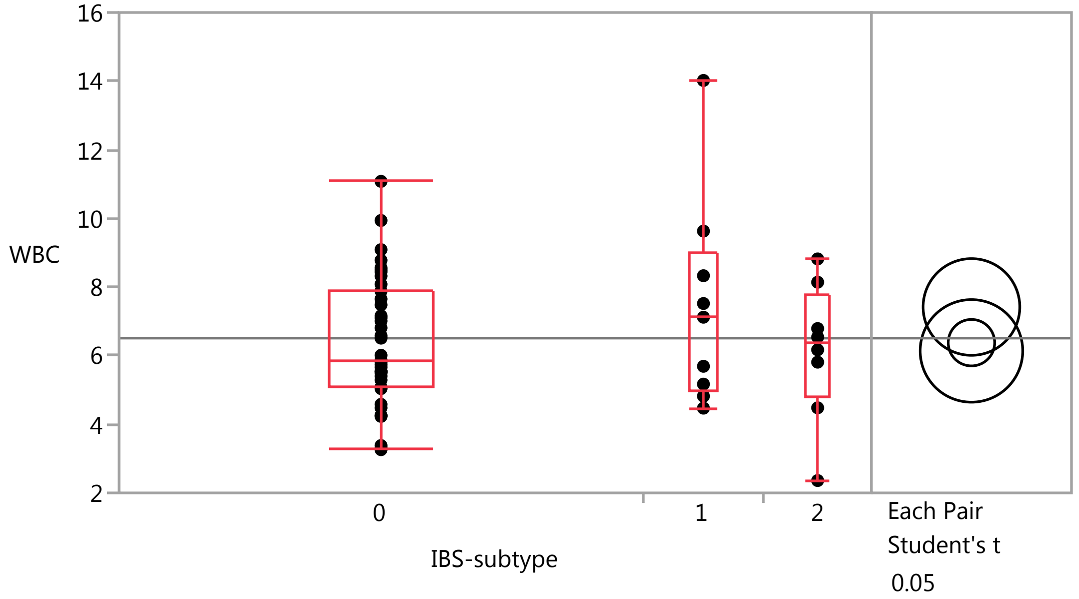


##### Quantiles

| Level | Minimum | 10% | 25% | Median | 75% | 90% | Maximum |
|---|---|---|---|---|---|---|---|
| 0 | 3.26 | 4.23 | 5.08 | 5.87 | 7.88 | 8.78 | 11.08 |
| 1 | 4.47 | 4.47 | 4.995 | 7.12 | 8.98 | 14.02 | 14.02 |
| 2 | 2.36 | 2.36 | 4.8125 | 6.355 | 7.8025 | 8.82 | 8.82 |

##### Means Comparisons

###### Comparisons for each pair using Student's t

###### Confidence Quantile

| t | Alpha |
|---|---|
| 2.00575 | 0.05 |

###### LSD Threshold Matrix

Abs(Dif)-LSD

| | 1 | 0 | 2 |
|---|---|---|---|
| 1 | -1.9931 | -0.5226 | -0.7743 |
| 0 | -0.5226 | -0.9575 | -1.4018 |
| 2 | -0.7743 | -1.4018 | -2.1140 |

Positive values show pairs of means that are significantly different.

###### Connecting Letters Report

| Level | | Mean |
|---|---|---|
| 1 | A | 7.4188889 |
| 0 | A | 6.3779487 |
| 2 | A | 6.1387500 |

Levels not connected by same letter are significantly different.

#### Fit Group

##### Oneway Analysis of WBC By IBS-subtype Weight=1

###### Means Comparisons

###### Comparisons for each pair using Student's t

###### Ordered Differences Report

| Level | - Level | Difference | Std Err Dif | Lower CL | Upper CL | p-Value |
|---|---|---|---|---|---|---|
| 1 | 2 | 1.280139 | 1.024281 | -0.77431 | 3.334586 | 0.2169 |
| 1 | 0 | 1.040940 | 0.779521 | -0.52258 | 2.604462 | 0.1875 |
| 0 | 2 | 0.239199 | 0.818149 | -1.40180 | 1.880198 | 0.7711 |

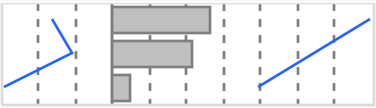

##### Oneway Analysis of MPV By IBS-subtype Weight=1

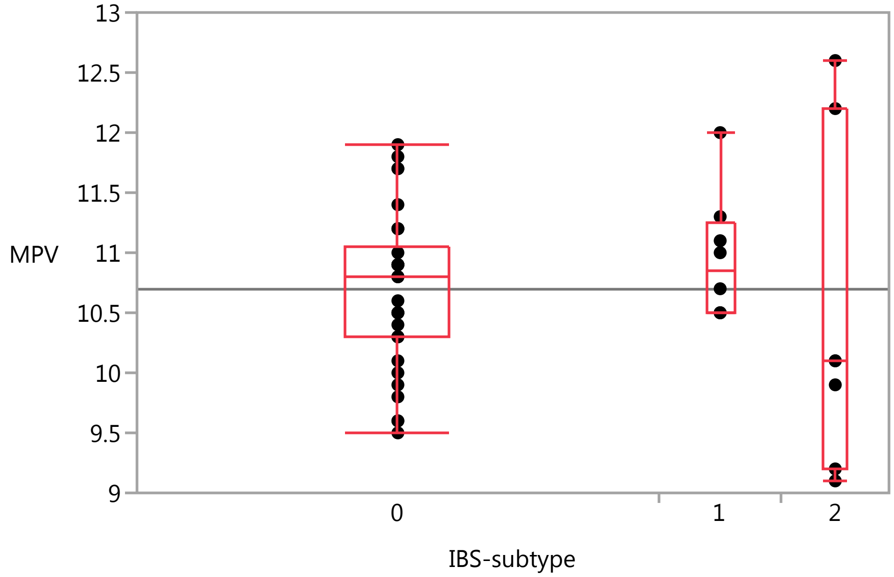


###### Quantiles

| Level | Minimum | 10% | 25% | Median | 75% | 90% | Maximum |
|---|---|---|---|---|---|---|---|
| 0 | 9.5 | 9.7 | 10.3 | 10.8 | 11.05 | 11.7 | 11.9 |
| 1 | 10.5 | 10.5 | 10.5 | 10.85 | 11.25 | 12 | 12 |
| 2 | 9.1 | 9.1 | 9.2 | 10.1 | 12.2 | 12.6 | 12.6 |

Missing Rows 7

#### Fit Group

##### Oneway Analysis of Monocytes By IBS-subtype Weight=1

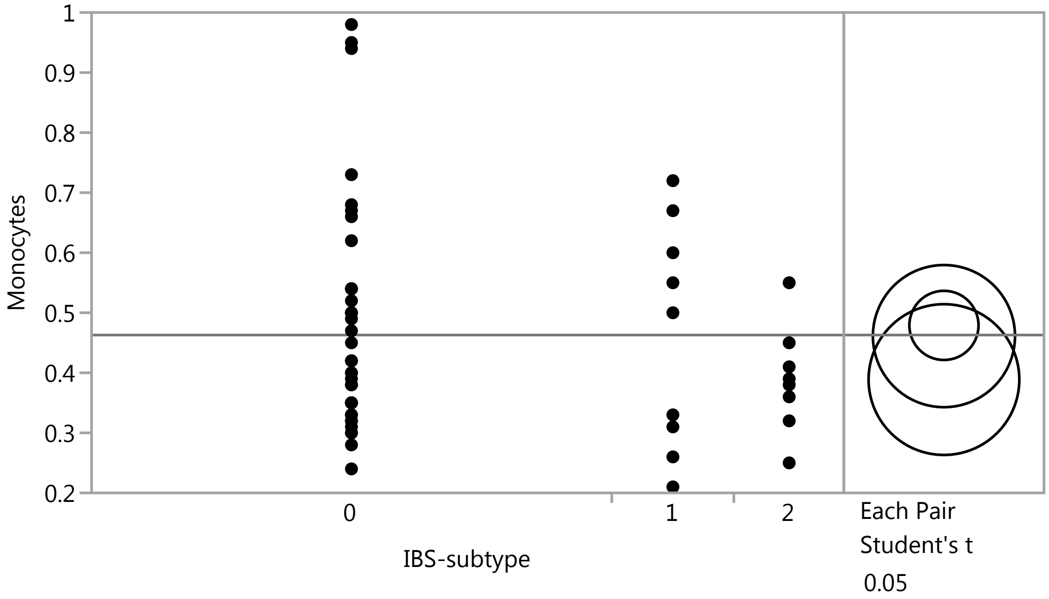


#### Means Comparisons

##### Comparisons for each pair using Student's t

###### Confidence Quantile

| t | Alpha |
|---|---|
| 2.00665 | 0.05 |

###### LSD Threshold Matrix

Abs(Dif)-LSD

| | 0 | 1 | 2 |
|---|---|---|---|
| 0 | -0.08148 | -0.11382 | -0.04796 |
| 1 | -0.11382 | -0.16742 | -0.10021 |
| 2 | -0.04796 | -0.10021 | -0.17758 |

Positive values show pairs of means that are significantly different.

###### Connecting Letters Report

| Level | | Mean |
|---|---|---|
| 0 | A | 0.47894737 |
| 1 | A | 0.46111111 |
| 2 | A | 0.38875000 |

Levels not connected by same letter are significantly different.

###### Ordered Differences Report

| Level | - Level | Difference | Std Err Dif | Lower CL | Upper CL | p-Value |
|---|---|---|---|---|---|---|
| 0 | 2 | 0.0901974 | 0.0688481 | -0.047957 | 0.2283513 | 0.1959 |
| 1 | 2 | 0.0723611 | 0.0860020 | -0.100214 | 0.2449367 | 0.4040 |
| 0 | 1 | 0.0178363 | 0.0656124 | -0.113825 | 0.1494972 | 0.7868 |

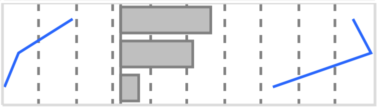

Missing Rows 1

#### Fit Group

##### Oneway Analysis of Monocytes_PCT By IBS-subtype Weight=1

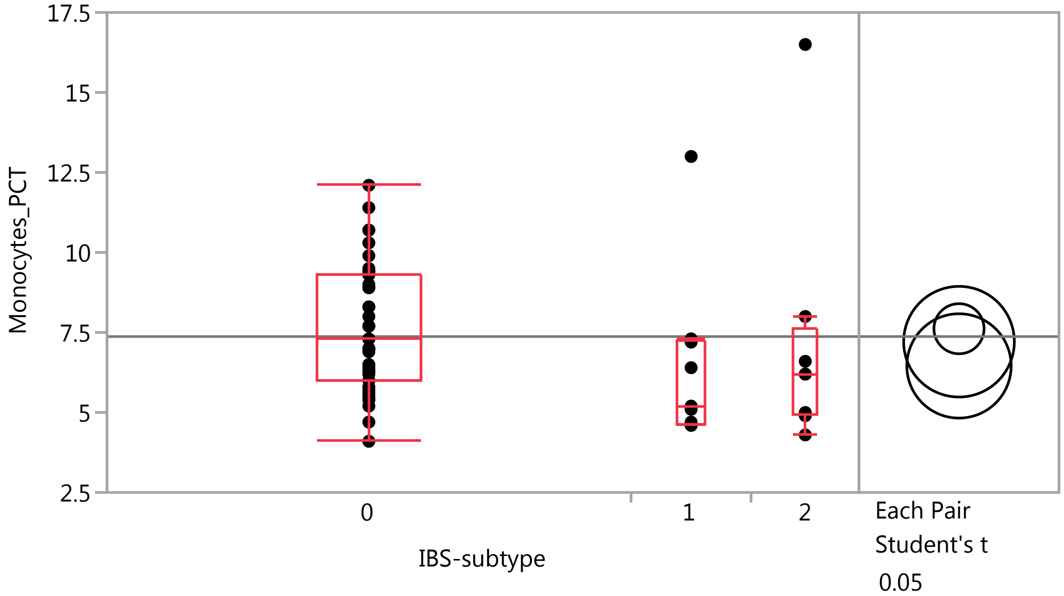


##### Quantiles

| Level | Minimum | 10% | 25% | Median | 75% | 90% | Maximum |
|---|---|---|---|---|---|---|---|
| 0 | 4.1 | 5.4 | 6 | 7.3 | 9.3 | 10.7 | 12.1 |
| 1 | 4.6 | 4.6 | 4.65 | 5.2 | 7.25 | 13 | 13 |
| 2 | 4.3 | 4.3 | 4.925 | 6.2 | 7.65 | 16.5 | 16.5 |

##### Means Comparisons

###### Comparisons for each pair using Student's t

###### Confidence Quantile

| t | Alpha |
|---|---|
| 2.00575 | 0.05 |

###### LSD Threshold Matrix

Abs(Dif)-LSD

| | 0 | 2 | 1 |
|---|---|---|---|
| 0 | -1.1076 | -1.4928 | -0.6462 |
| 2 | -1.4928 | -2.4454 | -1.6196 |
| 1 | -0.6462 | -1.6196 | -2.3056 |

Positive values show pairs of means that are significantly different.

###### Connecting Letters Report

| Level | | Mean |
|---|---|---|
| 0 | A | 7.6179487 |
| 2 | A | 7.2125000 |
| 1 | A | 6.4555556 |

Levels not connected by same letter are significantly different.

### Fit Group

#### Oneway Analysis of Monocytes_PCT By IBS-subtype Weight=1

##### Means Comparisons

###### Comparisons for each pair using Student's t

###### Ordered Differences Report

| Level | - Level | Difference | Std Err Dif | Lower CL | Upper CL | p-Value |
|---|---|---|---|---|---|---|
| 0 | 1 | 1.162393 | 0.901722 | -0.64623 | 2.971019 | 0.2030 |
| 2 | 1 | 0.756944 | 1.184852 | -1.61957 | 3.133456 | 0.5257 |
| 0 | 2 | 0.405449 | 0.946406 | -1.49280 | 2.303699 | 0.6701 |

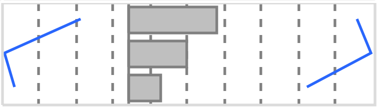

#### Oneway Analysis of sCD14 By IBS-subtype Weight=1

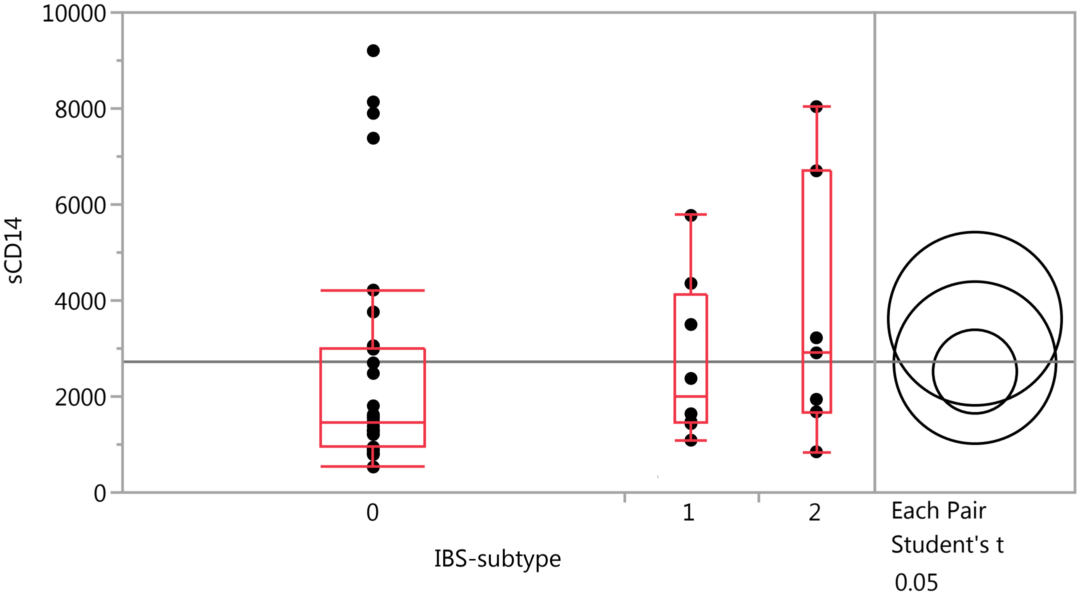


##### Quantiles

| Level | Minimum | 10% | 25% | Median | 75% | 90% | Maximum |
|---|---|---|---|---|---|---|---|
| 0 | 531.3 | 810.49 | 941.3 | 1471.25 | 3002.975 | 7847.13 | 9204.5 |
| 1 | 1089.9 | 1089.9 | 1445.325 | 2008 | 4142.425 | 5772 | 5772 |
| 2 | 847.6 | 847.6 | 1680.2 | 2909 | 6701.5 | 8038.3 | 8038.3 |

##### Means Comparisons

###### Comparisons for each pair using Student's t

###### Confidence Quantile

| t | Alpha |
|---|---|
| 2.01808 | 0.05 |

### Fit Group

#### Oneway Analysis of sCD14 By IBS-subtype Weight=1

##### Means Comparisons

###### Comparisons for each pair using Student's t

###### LSD Threshold Matrix

Abs(Dif)-LSD

| | 2 | 1 | 0 |
|---|---|---|---|
| 2 | -2552.8 | -1557.3 | -903.5 |
| 1 | -1557.3 | -2388.0 | -1713.7 |
| 0 | -903.5 | -1713.7 | -1233.1 |

Positive values show pairs of means that are significantly different.

###### Connecting Letters Report

| Level | | Mean |
|---|---|---|
| 2 | A | 3620.0571 |
| 1 | A | 2705.6125 |
| 0 | A | 2518.9033 |

Levels not connected by same letter are significantly different.

###### Ordered Differences Report

| Level | - Level | Difference | Std Err Dif | Lower CL | Upper CL | p-Value |
|---|---|---|---|---|---|---|
| 2 | 0 | 1101.154 | 993.366 | -903.54 | 3105.847 | 0.2740 |
| 2 | 1 | 914.445 | 1224.812 | -1557.33 | 3386.216 | 0.4595 |
| 1 | 0 | 186.709 | 941.682 | -1713.68 | 2087.100 | 0.8438 |

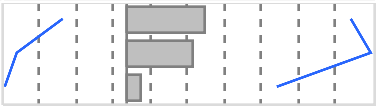

Missing Rows 11

#### Oneway Analysis of Basophil_PCT By IBS-subtype Weight=1

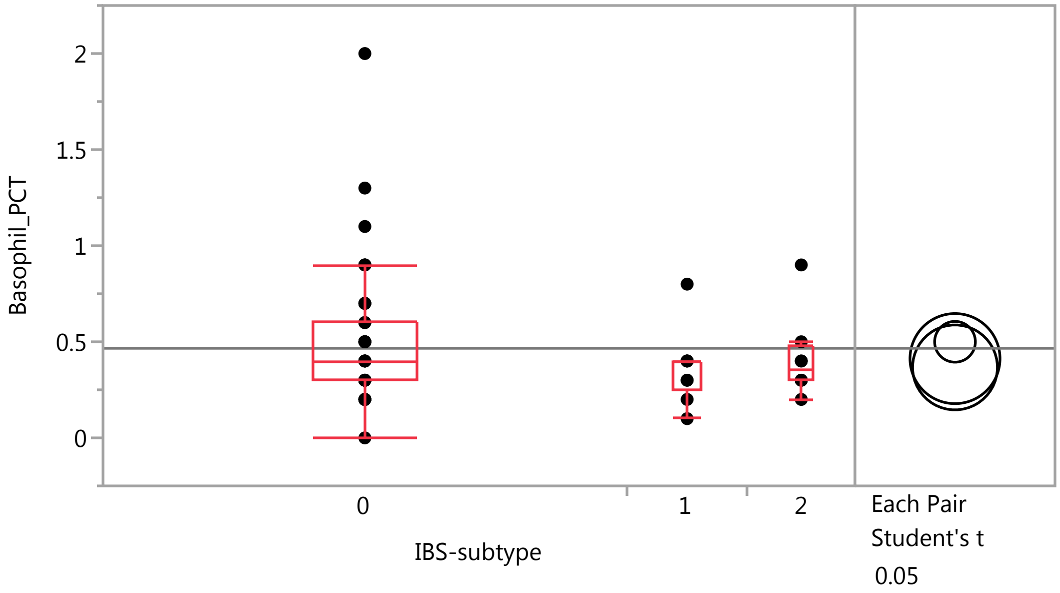

### Fit Group

#### Oneway Analysis of Basophil_PCT By IBS-subtype Weight=1

##### Quantiles

| Level | Minimum | 10% | 25% | Median | 75% | 90% | Maximum |
|---|---|---|---|---|---|---|---|
| 0 | 0 | 0.2 | 0.3 | 0.4 | 0.6 | 0.9 | 2 |
| 1 | 0.1 | 0.1 | 0.25 | 0.4 | 0.4 | 0.8 | 0.8 |
| 2 | 0.2 | 0.2 | 0.3 | 0.35 | 0.475 | 0.9 | 0.9 |

##### Means Comparisons

###### Comparisons for each pair using Student's t

###### Confidence Quantile

| t | Alpha |
|---|---|
| 2.00575 | 0.05 |

###### LSD Threshold Matrix

Abs(Dif)-LSD

| | 0 | 2 | 1 |
|---|---|---|---|
| 0 | -0.15011 | -0.16978 | -0.11180 |
| 2 | -0.16978 | -0.33144 | -0.27627 |
| 1 | -0.11180 | -0.27627 | -0.31248 |

Positive values show pairs of means that are significantly different.

###### Connecting Letters Report

| Level | | Mean |
|---|---|---|
| 0 | A | 0.50000000 |
| 2 | A | 0.41250000 |
| 1 | A | 0.36666667 |

Levels not connected by same letter are significantly different.

###### Ordered Differences Report

| Level | - Level | Difference | Std Err Dif | Lower CL | Upper CL | p-Value |
|---|---|---|---|---|---|---|
| 0 | 1 | 0.1333333 | 0.1222142 | -0.111797 | 0.3784639 | 0.2802 |
| 0 | 2 | 0.0875000 | 0.1282703 | -0.169778 | 0.3447776 | 0.4981 |
| 2 | 1 | 0.0458333 | 0.1605878 | -0.276265 | 0.3679318 | 0.7764 |

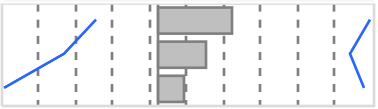

#### Oneway Analysis of Lymphocytes By IBS-subtype Weight=1

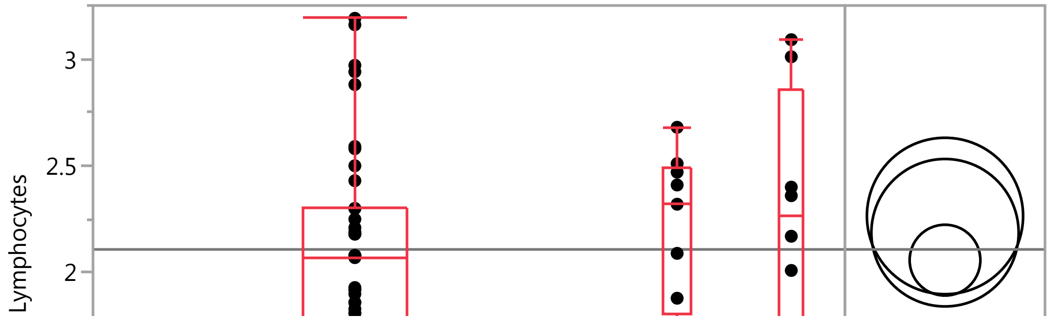

#### Fit Group

##### Oneway Analysis of Lymphocytes By IBS-subtype Weight=1

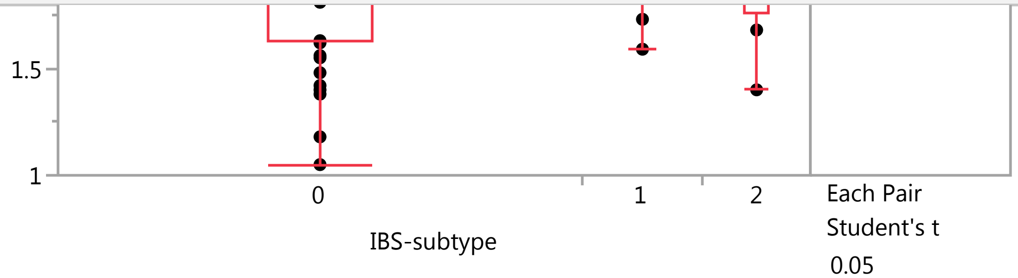


##### Quantiles

| Level | Minimum | 10% | 25% | Median | 75% | 90% | Maximum |
|---|---|---|---|---|---|---|---|
| 0 | 1.05 | 1.4 | 1.63 | 2.07 | 2.3 | 2.94 | 3.19 |
| 1 | 1.59 | 1.59 | 1.805 | 2.32 | 2.49 | 2.68 | 2.68 |
| 2 | 1.4 | 1.4 | 1.7625 | 2.265 | 2.8575 | 3.09 | 3.09 |

##### Means Comparisons

###### Comparisons for each pair using Student's t

###### Confidence Quantile

| t | Alpha |
|---|---|
| 2.00575 | 0.05 |

###### LSD Threshold Matrix

Abs(Dif)-LSD

| | 2 | 1 | 0 |
|---|---|---|---|
| 2 | -0.51768 | -0.42476 | -0.19505 |
| 1 | -0.42476 | -0.48807 | -0.25441 |
| 0 | -0.19505 | -0.25441 | -0.23446 |

Positive values show pairs of means that are significantly different.

###### Connecting Letters Report

| Level | | Mean |
|---|---|---|
| 2 | A | 2.2650000 |
| 1 | A | 2.1866667 |
| 0 | A | 2.0582051 |

Levels not connected by same letter are significantly different.

###### Ordered Differences Report

| Level | - Level | Difference | Std Err Dif | Lower CL | Upper CL | p-Value |
|---|---|---|---|---|---|---|
| 2 | 0 | 0.2067949 | 0.2003478 | -0.195052 | 0.6086416 | 0.3067 |
| 1 | 0 | 0.1284615 | 0.1908886 | -0.254412 | 0.5113355 | 0.5039 |
| 2 | 1 | 0.0783333 | 0.2508251 | -0.424758 | 0.5814249 | 0.7560 |

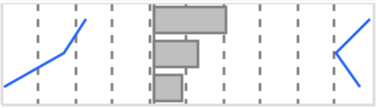

#### Fit Group

##### Oneway Analysis of Basophils By IBS-subtype Weight=1

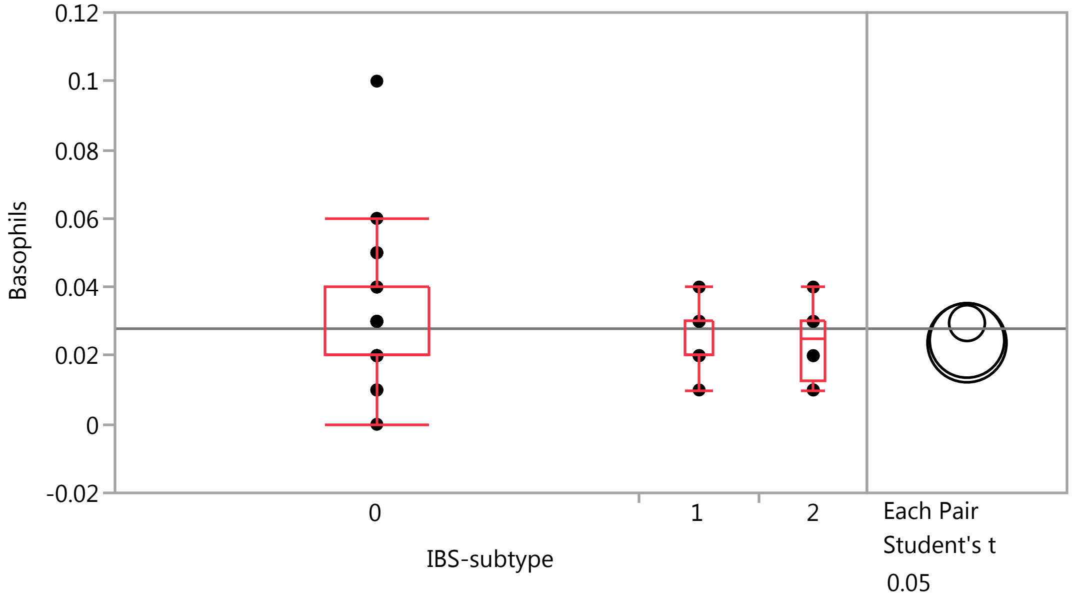


##### Quantiles

| Level | Minimum | 10% | 25% | Median | 75% | 90% | Maximum |
|---|---|---|---|---|---|---|---|
| 0 | 0 | 0.01 | 0.02 | 0.02 | 0.04 | 0.05 | 0.1 |
| 1 | 0.01 | 0.01 | 0.02 | 0.02 | 0.03 | 0.04 | 0.04 |
| 2 | 0.01 | 0.01 | 0.0125 | 0.025 | 0.03 | 0.04 | 0.04 |

##### Means Comparisons

###### Comparisons for each pair using Student's t

###### Confidence Quantile

| t | Alpha |
|---|---|
| 2.00575 | 0.05 |

###### LSD Threshold Matrix

Abs(Dif)-LSD

| | 0 | 1 | 2 |
|---|---|---|---|
| 0 | -0.00738 | -0.00701 | -0.00692 |
| 1 | -0.00701 | -0.01537 | -0.01515 |
| 2 | -0.00692 | -0.01515 | -0.01630 |

Positive values show pairs of means that are significantly different.

###### Connecting Letters Report

| Level | | Mean |
|---|---|---|
| 0 | A | 0.02948718 |
| 1 | A | 0.02444444 |
| 2 | A | 0.02375000 |

Levels not connected by same letter are significantly different.

### Fit Group

#### Oneway Analysis of Basophils By IBS-subtype Weight=1

##### Means Comparisons

###### Comparisons for each pair using Student's t

###### Ordered Differences Report

| Level | - Level | Difference | Std Err Dif | Lower CL | Upper CL | p-Value |
|---|---|---|---|---|---|---|
| 0 | 2 | 0.0057372 | 0.0063080 | -0.006915 | 0.0183894 | 0.3672 |
| 0 | 1 | 0.0050427 | 0.0060101 | -0.007012 | 0.0170976 | 0.4052 |
| 1 | 2 | 0.0006944 | 0.0078973 | -0.015145 | 0.0165343 | 0.9303 |

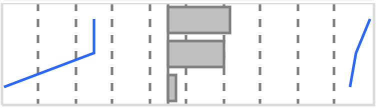

#### Oneway Analysis of Eosinophil_PCT By IBS-subtype Weight=1

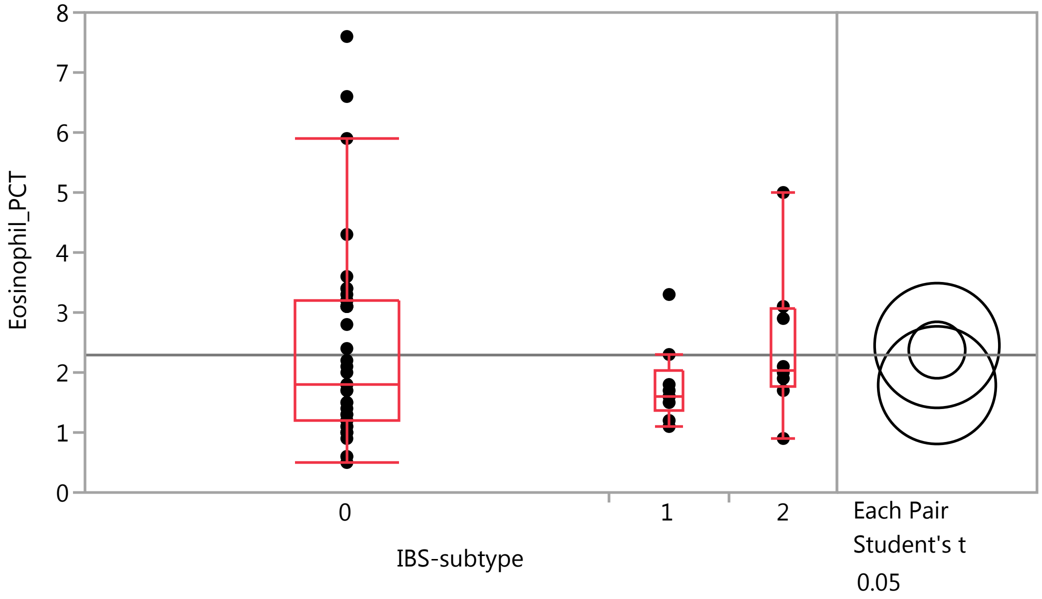


##### Quantiles

| Level | Minimum | 10% | 25% | Median | 75% | 90% | Maximum |
|---|---|---|---|---|---|---|---|
| 0 | 0.5 | 0.9 | 1.2 | 1.8 | 3.2 | 4.3 | 7.6 |
| 1 | 1.1 | 1.1 | 1.35 | 1.6 | 2.05 | 3.3 | 3.3 |
| 2 | 0.9 | 0.9 | 1.75 | 2.05 | 3.05 | 5 | 5 |

##### Means Comparisons

###### Comparisons for each pair using Student's t

###### Confidence Quantile

| t | Alpha |
|---|---|
| 2.00575 | 0.05 |

### Fit Group

#### Oneway Analysis of Eosinophil_PCT By IBS-subtype Weight=1

##### Means Comparisons

###### Comparisons for each pair using Student's t

###### LSD Threshold Matrix

Abs(Dif)-LSD

| | 2 | 0 | 1 |
|---|---|---|---|
| 2 | -1.4703 | -1.0657 | -0.7678 |
| 0 | -1.0657 | -0.6659 | -0.5020 |
| 1 | -0.7678 | -0.5020 | -1.3862 |

Positive values show pairs of means that are significantly different.

###### Connecting Letters Report

| Level | | Mean |
|---|---|---|
| 2 | A | 2.4500000 |
| 0 | A | 2.3743590 |
| 1 | A | 1.7888889 |

Levels not connected by same letter are significantly different.

###### Ordered Differences Report

| Level | - Level | Difference | Std Err Dif | Lower CL | Upper CL | p-Value |
|---|---|---|---|---|---|---|
| 2 | 1 | 0.6611111 | 0.7124042 | -0.76779 | 2.090013 | 0.3576 |
| 0 | 1 | 0.5854701 | 0.5421698 | -0.50198 | 1.672925 | 0.2851 |
| 2 | 0 | 0.0756410 | 0.5690362 | -1.06570 | 1.216983 | 0.8948 |

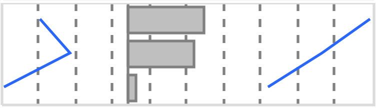

#### Oneway Analysis of HCT By IBS-subtype Weight=1

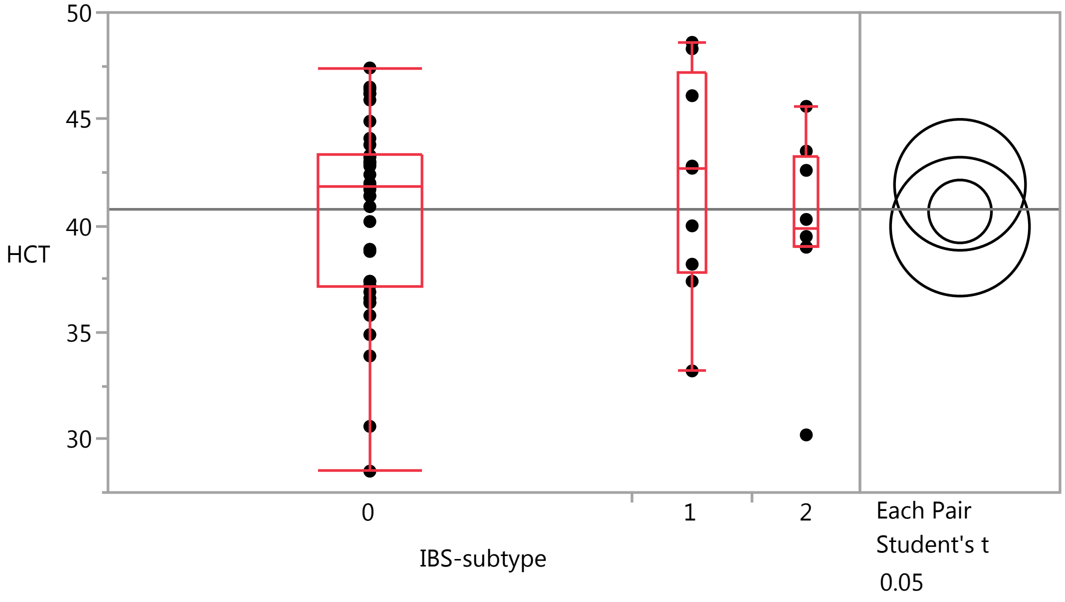

### Fit Group

#### Oneway Analysis of HCT By IBS-subtype Weight=1

##### Quantiles

| Level | Minimum | 10% | 25% | Median | 75% | 90% | Maximum |
|---|---|---|---|---|---|---|---|
| 0 | 28.5 | 34.9 | 37.2 | 41.8 | 43.3 | 46.2 | 47.4 |
| 1 | 33.2 | 33.2 | 37.8 | 42.7 | 47.2 | 48.6 | 48.6 |
| 2 | 30.2 | 30.2 | 39 | 39.9 | 43.275 | 45.6 | 45.6 |

##### Means Comparisons

###### Comparisons for each pair using Student's t

###### Confidence Quantile

| t | Alpha |
|---|---|
| 2.00575 | 0.05 |

###### LSD Threshold Matrix

Abs(Dif)-LSD

| | 1 | 0 | 2 |
|---|---|---|---|
| 1 | -4.3411 | -2.1576 | -2.5150 |
| 0 | -2.1576 | -2.0854 | -2.8623 |
| 2 | -2.5150 | -2.8623 | -4.6044 |

Positive values show pairs of means that are significantly different.

###### Connecting Letters Report

| Level | | Mean |
|---|---|---|
| 1 | A | 41.922222 |
| 0 | A | 40.674359 |
| 2 | A | 39.962500 |

Levels not connected by same letter are significantly different.

###### Ordered Differences Report

| Level | - Level | Difference | Std Err Dif | Lower CL | Upper CL | p-Value |
|---|---|---|---|---|---|---|
| 1 | 2 | 1.959722 | 2.230938 | -2.51497 | 6.434416 | 0.3837 |
| 1 | 0 | 1.247863 | 1.697838 | -2.15757 | 4.653295 | 0.4656 |
| 0 | 2 | 0.711859 | 1.781972 | -2.86232 | 4.286042 | 0.6911 |

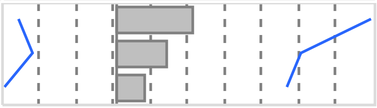

#### Oneway Analysis of RBC By IBS-subtype Weight=1

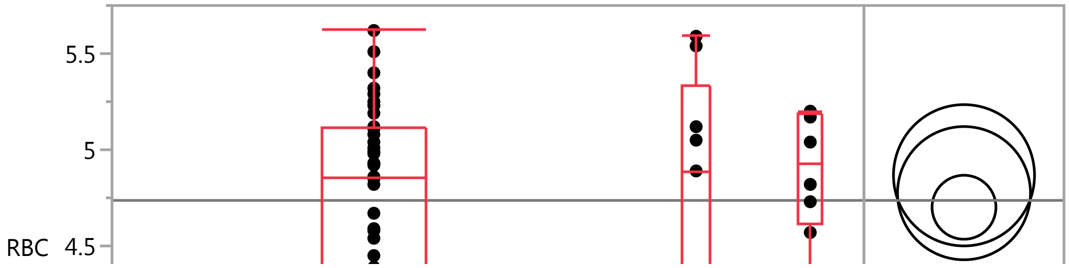

#### Fit Group

##### Oneway Analysis of RBC By IBS-subtype Weight=1

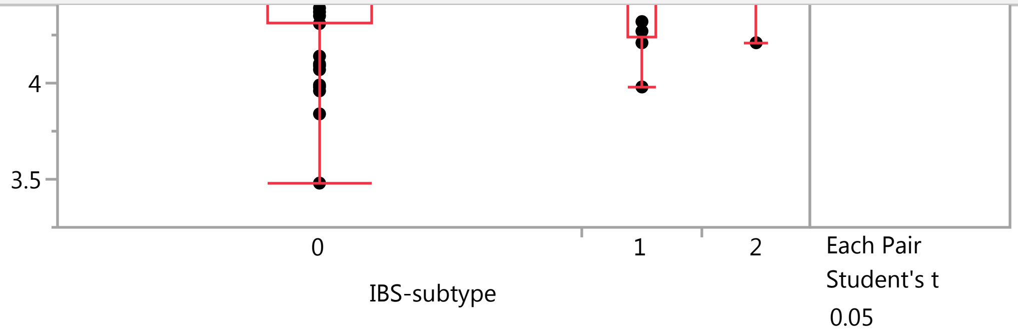


##### Quantiles

| Level | Minimum | 10% | 25% | Median | 75% | 90% | Maximum |
|---|---|---|---|---|---|---|---|
| 0 | 3.48 | 3.98 | 4.31 | 4.85 | 5.11 | 5.32 | 5.62 |
| 1 | 3.98 | 3.98 | 4.24 | 4.89 | 5.33 | 5.59 | 5.59 |
| 2 | 4.21 | 4.21 | 4.61 | 4.93 | 5.1925 | 5.2 | 5.2 |

##### Means Comparisons

###### Comparisons for each pair using Student's t

###### Confidence Quantile

| t | Alpha |
|---|---|
| 2.00575 | 0.05 |

###### LSD Threshold Matrix

Abs(Dif)-LSD

| | 2 | 1 | 0 |
|---|---|---|---|
| 2 | -0.51945 | -0.41176 | -0.23700 |
| 1 | -0.41176 | -0.48974 | -0.31102 |
| 0 | -0.23700 | -0.31102 | -0.23526 |

Positive values show pairs of means that are significantly different.

###### Connecting Letters Report

| Level | | Mean |
|---|---|---|
| 2 | A | 4.8675000 |
| 1 | A | 4.7744444 |
| 0 | A | 4.7012821 |

Levels not connected by same letter are significantly different.

###### Ordered Differences Report

| Level | - Level | Difference | Std Err Dif | Lower CL | Upper CL | p-Value |
|---|---|---|---|---|---|---|
| 2 | 0 | 0.1662179 | 0.2010335 | -0.237004 | 0.5694400 | 0.4120 |
| 2 | 1 | 0.0930556 | 0.2516836 | -0.411758 | 0.5978689 | 0.7131 |
| 1 | 0 | 0.0731624 | 0.1915419 | -0.311022 | 0.4573468 | 0.7040 |

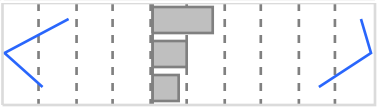

#### Fit Group

##### Oneway Analysis of PlateletCount By IBS-subtype Weight=1

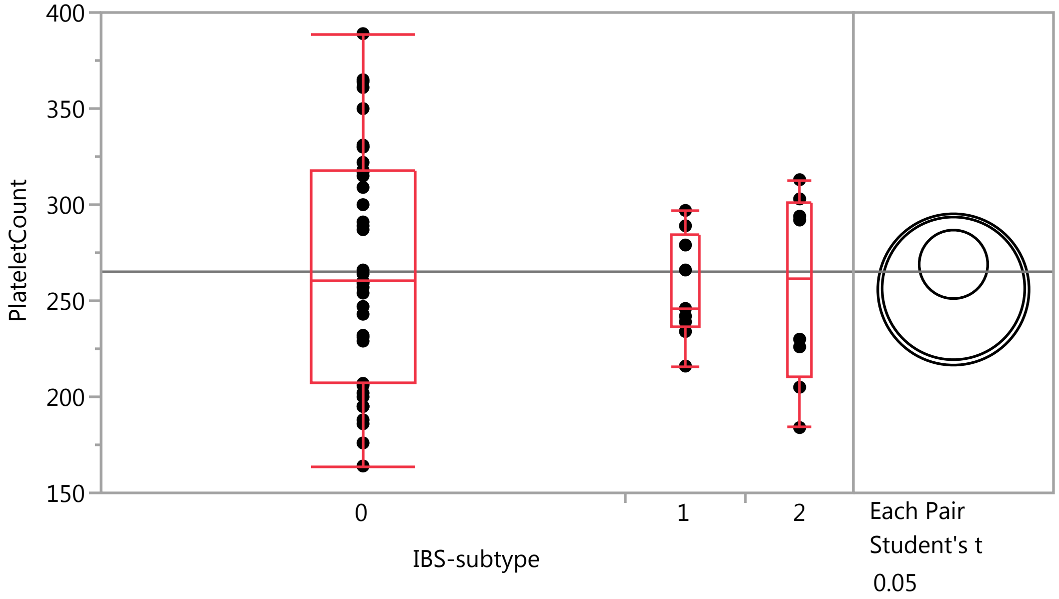


##### Quantiles

| Level | Minimum | 10% | 25% | Median | 75% | 90% | Maximum |
|---|---|---|---|---|---|---|---|
| 0 | 164 | 188 | 207 | 260 | 318 | 361 | 389 |
| 1 | 216 | 216 | 236.5 | 246 | 284 | 297 | 297 |
| 2 | 184 | 184 | 210.25 | 261 | 300.75 | 313 | 313 |

##### Means Comparisons

###### Comparisons for each pair using Student's t

###### Confidence Quantile

| t | Alpha |
|---|---|
| 2.00575 | 0.05 |

###### LSD Threshold Matrix

Abs(Dif)-LSD

| | 0 | 1 | 2 |
|---|---|---|---|
| 0 | -25.216 | -28.674 | -30.145 |
| 1 | -28.674 | -52.492 | -53.538 |
| 2 | -30.145 | -53.538 | -55.676 |

Positive values show pairs of means that are significantly different.

###### Connecting Letters Report

| Level | | Mean |
|---|---|---|
| 0 | A | 268.94872 |
| 1 | A | 256.44444 |
| 2 | A | 255.87500 |

Levels not connected by same letter are significantly different.

### Fit Group

#### Oneway Analysis of PlateletCount By IBS-subtype Weight=1

##### Means Comparisons

###### Comparisons for each pair using Student's t

###### Ordered Differences Report

| Level | - Level | Difference | Std Err Dif | Lower CL | Upper CL | p-Value |
|---|---|---|---|---|---|---|
| 0 | 2 | 13.07372 | 21.54752 | -30.1451 | 56.29257 | 0.5466 |
| 0 | 1 | 12.50427 | 20.53018 | -28.6740 | 53.68259 | 0.5451 |
| 1 | 2 | 0.56944 | 26.97639 | -53.5383 | 54.67723 | 0.9832 |

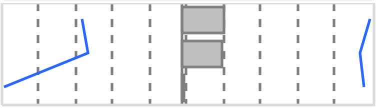

#### Oneway Analysis of IgE By IBS-subtype Weight=1

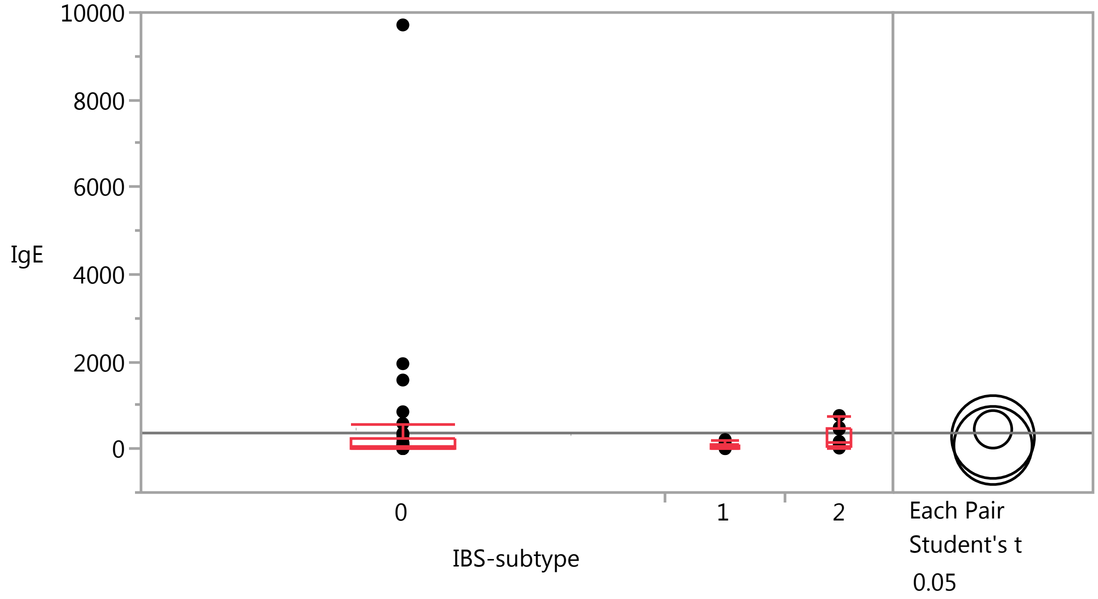


##### Quantiles

| Level | Minimum | 10% | 25% | Median | 75% | 90% | Maximum |
|---|---|---|---|---|---|---|---|
| 0 | 5.6 | 7 | 20.7 | 46.6 | 245 | 850 | 9715 |
| 1 | 4.8 | 4.8 | 16.8 | 68.9 | 115.75 | 213 | 213 |
| 2 | 18.5 | 18.5 | 44.1 | 164.5 | 479.25 | 764 | 764 |

##### Means Comparisons

###### Comparisons for each pair using Student's t

###### Confidence Quantile

| t | Alpha |
|---|---|
| 2.00575 | 0.05 |

### Fit Group

#### Oneway Analysis of IgE By IBS-subtype Weight=1

##### Means Comparisons

###### Comparisons for each pair using Student's t

###### LSD Threshold Matrix

Abs(Dif)-LSD

| | 0 | 2 | 1 |
|---|---|---|---|
| 0 | -608.2 | -866.5 | -623.9 |
| 2 | -866.5 | -1342.9 | -1111.6 |
| 1 | -623.9 | -1111.6 | -1266.1 |

Positive values show pairs of means that are significantly different.

###### Connecting Letters Report

| Level | | Mean |
|---|---|---|
| 0 | A | 447.45897 |
| 2 | A | 271.56250 |
| 1 | A | 78.13333 |

Levels not connected by same letter are significantly different.

###### Ordered Differences Report

| Level | - Level | Difference | Std Err Dif | Lower CL | Upper CL | p-Value |
|---|---|---|---|---|---|---|
| 0 | 1 | 369.3256 | 495.1796 | -623.88 | 1362.530 | 0.4591 |
| 2 | 1 | 193.4292 | 650.6597 | -1111.63 | 1498.487 | 0.7674 |
| 0 | 2 | 175.8965 | 519.7175 | -866.52 | 1218.318 | 0.7364 |

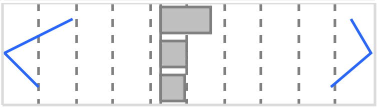

#### Oneway Analysis of ACTH By IBS-subtype Weight=1

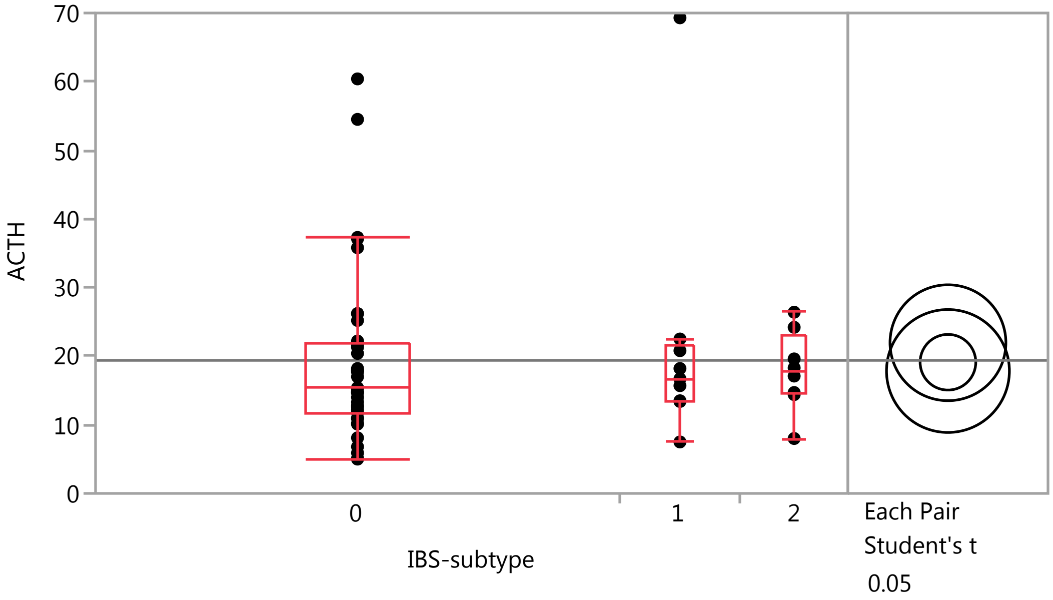

#### Fit Group

##### Oneway Analysis of ACTH By IBS-subtype Weight=1

###### Quantiles

| Level | Minimum | 10% | 25% | Median | 75% | 90% | Maximum |
|---|---|---|---|---|---|---|---|
| 0 | 5 | 6.8 | 11.8 | 15.4 | 22 | 37.1 | 60.4 |
| 1 | 7.5 | 7.5 | 13.45 | 16.7 | 21.65 | 69.3 | 69.3 |
| 2 | 8 | 8 | 14.475 | 17.7 | 23.05 | 26.4 | 26.4 |

###### Means Comparisons

###### Comparisons for each pair using Student's t

###### Confidence Quantile

| t | Alpha |
|---|---|
| 2.00575 | 0.05 |

###### LSD Threshold Matrix

Abs(Dif)-LSD

| | 1 | 0 | 2 |
|---|---|---|---|
| 1 | -11.942 | -6.526 | -8.192 |
| 0 | -6.526 | -5.737 | -8.557 |
| 2 | -8.192 | -8.557 | -12.667 |

Positive values show pairs of means that are significantly different.

###### Connecting Letters Report

| Level | | Mean |
|---|---|---|
| 1 | A | 21.955556 |
| 0 | A | 19.113287 |
| 2 | A | 17.837500 |

Levels not connected by same letter are significantly different.

###### Ordered Differences Report

| Level | - Level | Difference | Std Err Dif | Lower CL | Upper CL | p-Value |
|---|---|---|---|---|---|---|
| 1 | 2 | 4.118056 | 6.137291 | -8.19179 | 16.42790 | 0.5051 |
| 1 | 0 | 2.842268 | 4.670739 | -6.52605 | 12.21058 | 0.5454 |
| 0 | 2 | 1.275787 | 4.902190 | -8.55676 | 11.10834 | 0.7957 |

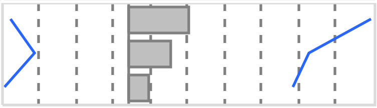

##### Oneway Analysis of Eosinophils By IBS-subtype Weight=1

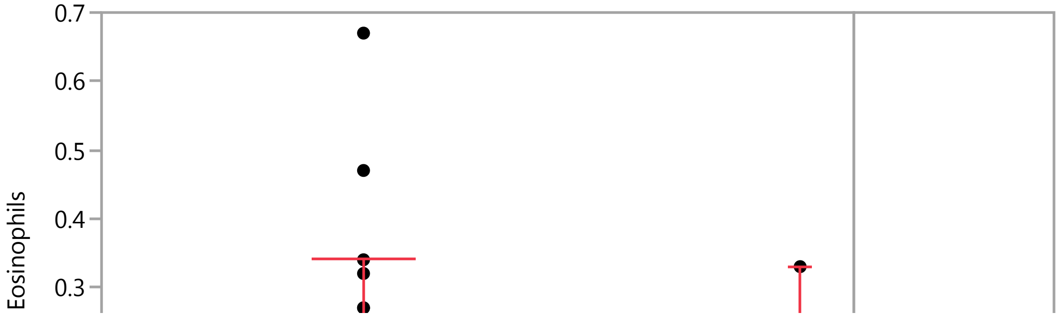

#### Fit Group

##### Oneway Analysis of Eosinophils By IBS-subtype Weight=1

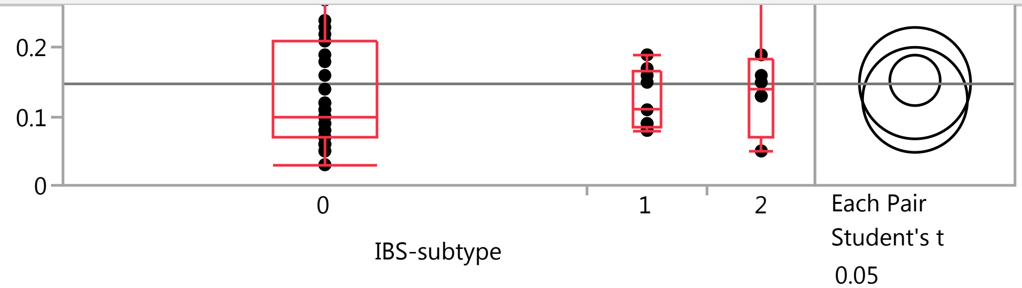


##### Quantiles

| Level | Minimum | 10% | 25% | Median | 75% | 90% | Maximum |
|---|---|---|---|---|---|---|---|
| 0 | 0.03 | 0.05 | 0.07 | 0.1 | 0.21 | 0.32 | 0.67 |
| 1 | 0.08 | 0.08 | 0.085 | 0.11 | 0.165 | 0.19 | 0.19 |
| 2 | 0.05 | 0.05 | 0.07 | 0.14 | 0.1825 | 0.33 | 0.33 |

##### Means Comparisons

###### Comparisons for each pair using Student's t

###### Confidence Quantile

| t | Alpha |
|---|---|
| 2.00575 | 0.05 |

###### LSD Threshold Matrix

Abs(Dif)-LSD

| | 0 | 2 | 1 |
|---|---|---|---|
| 0 | -0.05196 | -0.08498 | -0.05647 |
| 2 | -0.08498 | -0.11472 | -0.08718 |
| 1 | -0.05647 | -0.08718 | -0.10816 |

Positive values show pairs of means that are significantly different.

###### Connecting Letters Report

| Level | | Mean |
|---|---|---|
| 0 | A | 0.15282051 |
| 2 | A | 0.14875000 |
| 1 | A | 0.12444444 |

Levels not connected by same letter are significantly different.

###### Ordered Differences Report

| Level | - Level | Difference | Std Err Dif | Lower CL | Upper CL | p-Value | |
|---|---|---|---|---|---|---|---|
| 0 | 1 | 0.0283761 | 0.0423012 | -0.056469 | 0.1132214 | 0.5053 | 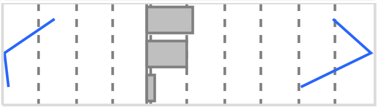 |
| 2 | 1 | 0.0243056 | 0.0555832 | -0.087180 | 0.1357913 | 0.6637 | |
| 0 | 2 | 0.0040705 | 0.0443973 | -0.084979 | 0.0931203 | 0.9273 | |

#### Fit Group

##### Oneway Analysis of IgG By IBS-subtype Weight=1

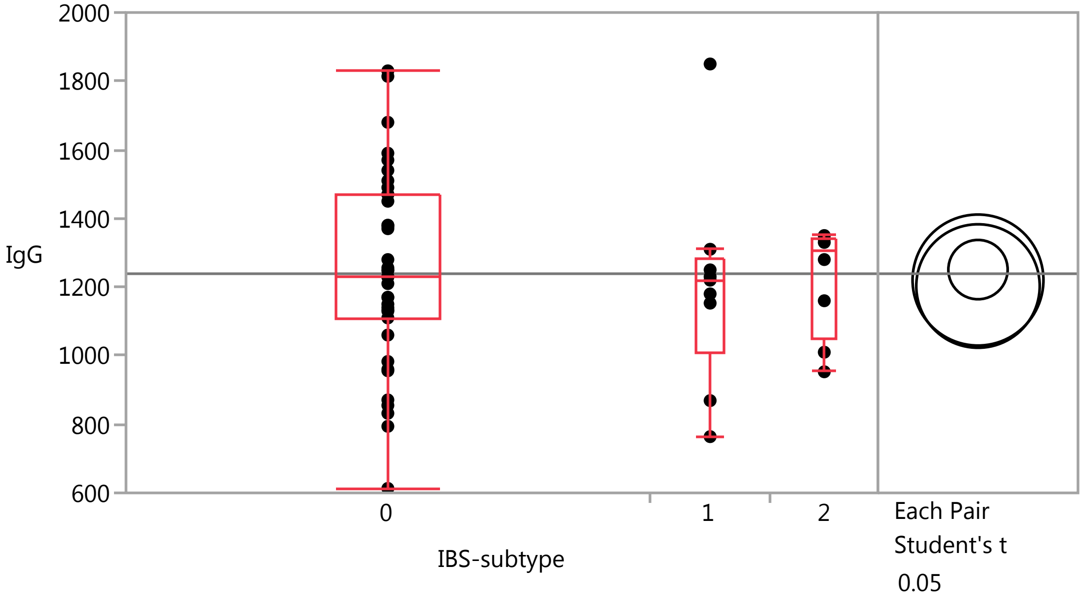


##### Quantiles

| Level | Minimum | 10% | 25% | Median | 75% | 90% | Maximum |
|---|---|---|---|---|---|---|---|
| 0 | 613 | 855 | 1110 | 1230 | 1470 | 1590 | 1830 |
| 1 | 764 | 764 | 1011 | 1220 | 1280 | 1850 | 1850 |
| 2 | 953 | 953 | 1047.5 | 1305 | 1340 | 1350 | 1350 |

##### Means Comparisons

###### Comparisons for each pair using Student's t

###### Confidence Quantile

| t | Alpha |
|---|---|
| 2.00575 | 0.05 |

###### LSD Threshold Matrix

Abs(Dif)-LSD

| | 0 | 2 | 1 |
|---|---|---|---|
| 0 | -122.27 | -179.15 | -151.77 |
| 2 | -179.15 | -269.97 | -244.88 |
| 1 | -151.77 | -244.88 | -254.53 |

Positive values show pairs of means that are significantly different.

###### Connecting Letters Report

| Level | | Mean |
|---|---|---|
| 0 | A | 1250.7949 |
| 2 | A | 1220.3750 |
| 1 | A | 1202.8889 |

Levels not connected by same letter are significantly different.

### Fit Group

#### Oneway Analysis of IgG By IBS-subtype Weight=1

##### Means Comparisons

###### Comparisons for each pair using Student's t

###### Ordered Differences Report

| Level | - Level | Difference | Std Err Dif | Lower CL | Upper CL | p-Value |
|---|---|---|---|---|---|---|
| 0 | 1 | 47.90598 | 99.5507 | -151.767 | 247.5794 | 0.6323 |
| 0 | 2 | 30.41987 | 104.4838 | -179.148 | 239.9878 | 0.7721 |
| 2 | 1 | 17.48611 | 130.8083 | -244.882 | 279.8544 | 0.8942 |

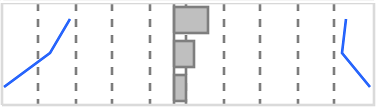

#### Oneway Analysis of ESR By IBS-subtype Weight=1

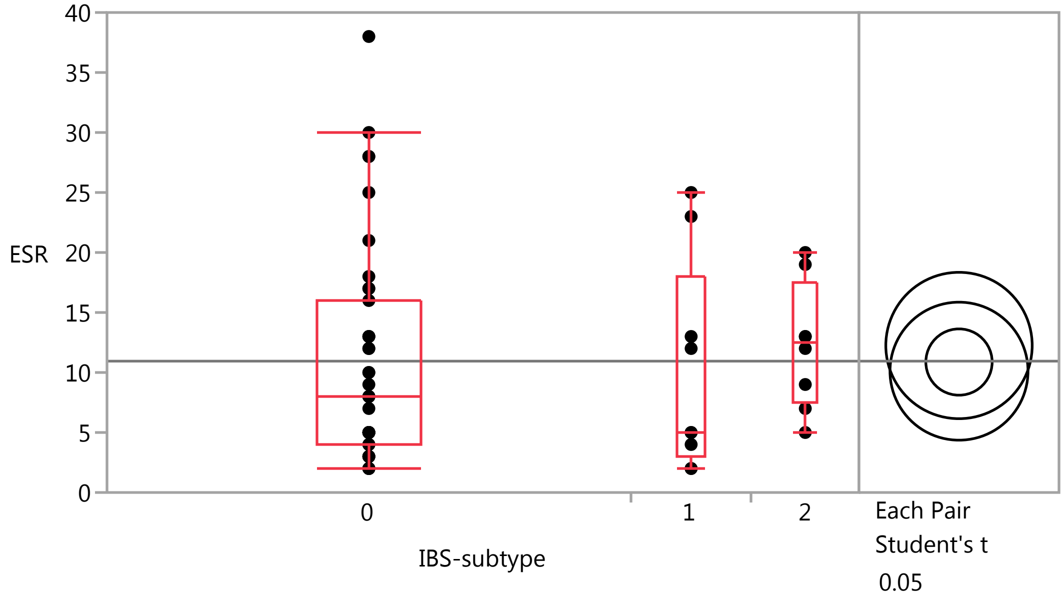


##### Quantiles

| Level | Minimum | 10% | 25% | Median | 75% | 90% | Maximum |
|---|---|---|---|---|---|---|---|
| 0 | 2 | 2 | 4 | 8 | 16 | 28 | 38 |
| 1 | 2 | 2 | 3 | 5 | 18 | 25 | 25 |
| 2 | 5 | 5 | 7.5 | 12.5 | 17.5 | 20 | 20 |

##### Means Comparisons

###### Comparisons for each pair using Student's t

###### Confidence Quantile

| t | Alpha |
|---|---|
| 2.00575 | 0.05 |

### Fit Group

#### Oneway Analysis of ESR By IBS-subtype Weight=1

##### Means Comparisons

###### Comparisons for each pair using Student's t

###### LSD Threshold Matrix

Abs(Dif)-LSD

| | 2 | 0 | 1 |
|---|---|---|---|
| 2 | -8.6257 | -5.3175 | -6.2437 |
| 0 | -5.3175 | -3.9067 | -5.6189 |
| 1 | -6.2437 | -5.6189 | -8.1324 |

Positive values show pairs of means that are significantly different.

###### Connecting Letters Report

| Level | | Mean |
|---|---|---|
| 2 | A | 12.250000 |
| 0 | A | 10.871795 |
| 1 | A | 10.111111 |

Levels not connected by same letter are significantly different.

###### Ordered Differences Report

| Level | - Level | Difference | Std Err Dif | Lower CL | Upper CL | p-Value |
|---|---|---|---|---|---|---|
| 2 | 1 | 2.138889 | 4.179311 | -6.24375 | 10.52153 | 0.6109 |
| 2 | 0 | 1.378205 | 3.338245 | -5.31747 | 8.07388 | 0.6814 |
| 0 | 1 | 0.760684 | 3.180633 | -5.61886 | 7.14023 | 0.8119 |

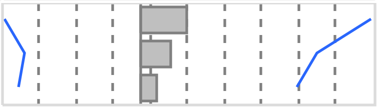

#### Oneway Analysis of CRP By IBS-subtype Weight=1

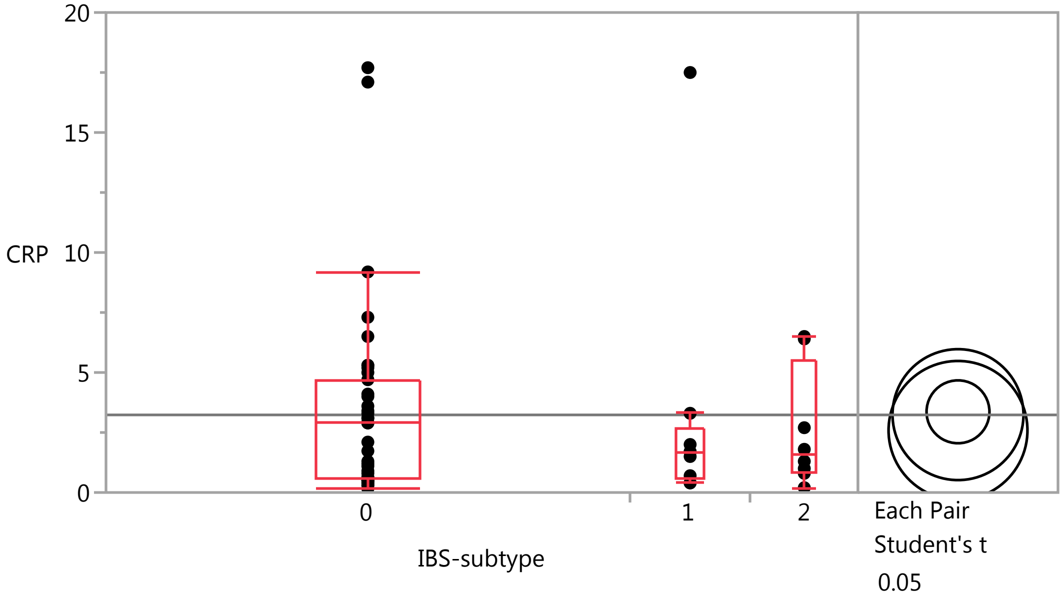

### Fit Group

#### Oneway Analysis of CRP By IBS-subtype Weight=1

##### Quantiles

| Level | Minimum | 10% | 25% | Median | 75% | 90% | Maximum |
|---|---|---|---|---|---|---|---|
| 0 | 0.16 | 0.4 | 0.6 | 2.9 | 4.7 | 7.3 | 17.7 |
| 1 | 0.4 | 0.4 | 0.55 | 1.7 | 2.65 | 17.5 | 17.5 |
| 2 | 0.2 | 0.2 | 0.85 | 1.55 | 5.475 | 6.5 | 6.5 |

##### Means Comparisons

###### Comparisons for each pair using Student's t

###### Confidence Quantile

| t | Alpha |
|---|---|
| 2.00575 | 0.05 |

###### LSD Threshold Matrix

Abs(Dif)-LSD

| | 0 | 1 | 2 |
|---|---|---|---|
| 0 | -1.8534 | -2.9087 | -2.4017 |
| 1 | -2.9087 | -3.8581 | -3.3199 |
| 2 | -2.4017 | -3.3199 | -4.0921 |

Positive values show pairs of means that are significantly different.

###### Connecting Letters Report

| Level | | Mean |
|---|---|---|
| 0 | A | 3.3623077 |
| 1 | A | 3.2444444 |
| 2 | A | 2.5875000 |

Levels not connected by same letter are significantly different.

###### Ordered Differences Report

| Level | - Level | Difference | Std Err Dif | Lower CL | Upper CL | p-Value |
|---|---|---|---|---|---|---|
| 0 | 2 | 0.7748077 | 1.583702 | -2.40170 | 3.951312 | 0.6267 |
| 1 | 2 | 0.6569444 | 1.982714 | -3.31988 | 4.633765 | 0.7417 |
| 0 | 1 | 0.1178632 | 1.508929 | -2.90867 | 3.144392 | 0.9380 |

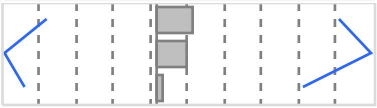

#### Oneway Analysis of LDH By IBS-subtype Weight=1

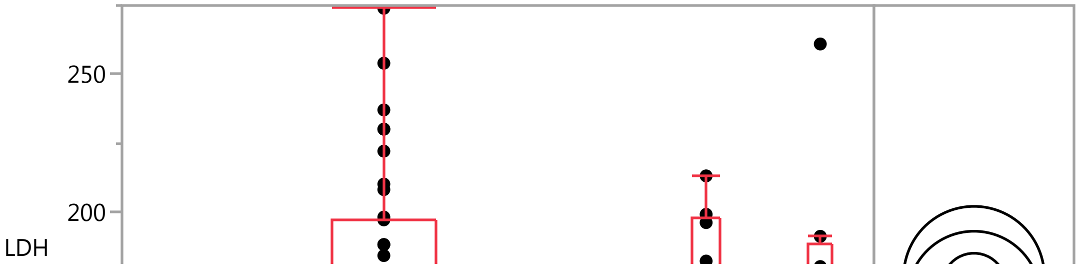

#### Fit Group

##### Oneway Analysis of LDH By IBS-subtype Weight=1

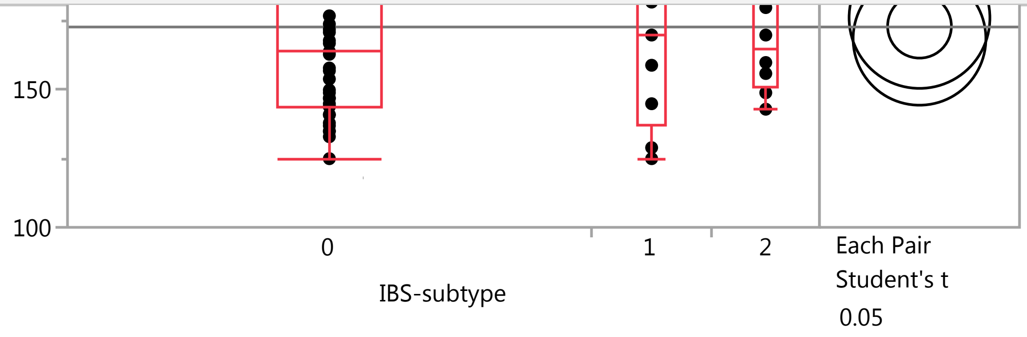


##### Quantiles

| Level | Minimum | 10% | 25% | Median | 75% | 90% | Maximum |
|---|---|---|---|---|---|---|---|
| 0 | 125 | 135 | 144 | 164 | 197 | 230 | 274 |
| 1 | 125 | 125 | 137 | 170 | 197.5 | 213 | 213 |
| 2 | 143 | 143 | 150.75 | 165 | 188.25 | 261 | 261 |

##### Means Comparisons

###### Comparisons for each pair using Student's t

###### Confidence Quantile

| t | Alpha |
|---|---|
| 2.00575 | 0.05 |

###### LSD Threshold Matrix

Abs(Dif)-LSD

| | 2 | 0 | 1 |
|---|---|---|---|
| 2 | -36.299 | -25.158 | -27.693 |
| 0 | -25.158 | -16.440 | -22.282 |
| 1 | -27.693 | -22.282 | -34.223 |

Positive values show pairs of means that are significantly different.

###### Connecting Letters Report

| Level | | Mean |
|---|---|---|
| 2 | A | 176.25000 |
| 0 | A | 173.23077 |
| 1 | A | 168.66667 |

Levels not connected by same letter are significantly different.

###### Ordered Differences Report

| Level | - Level | Difference | Std Err Dif | Lower CL | Upper CL | p-Value |
|---|---|---|---|---|---|---|
| 2 | 1 | 7.583333 | 17.58739 | -27.6925 | 42.85917 | 0.6681 |
| 0 | 1 | 4.564103 | 13.38475 | -22.2823 | 31.41051 | 0.7345 |
| 2 | 0 | 3.019231 | 14.04801 | -25.1575 | 31.19597 | 0.8307 |

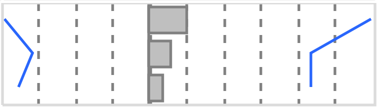

#### Fit Group

##### Oneway Analysis of IgA By IBS-subtype Weight=1

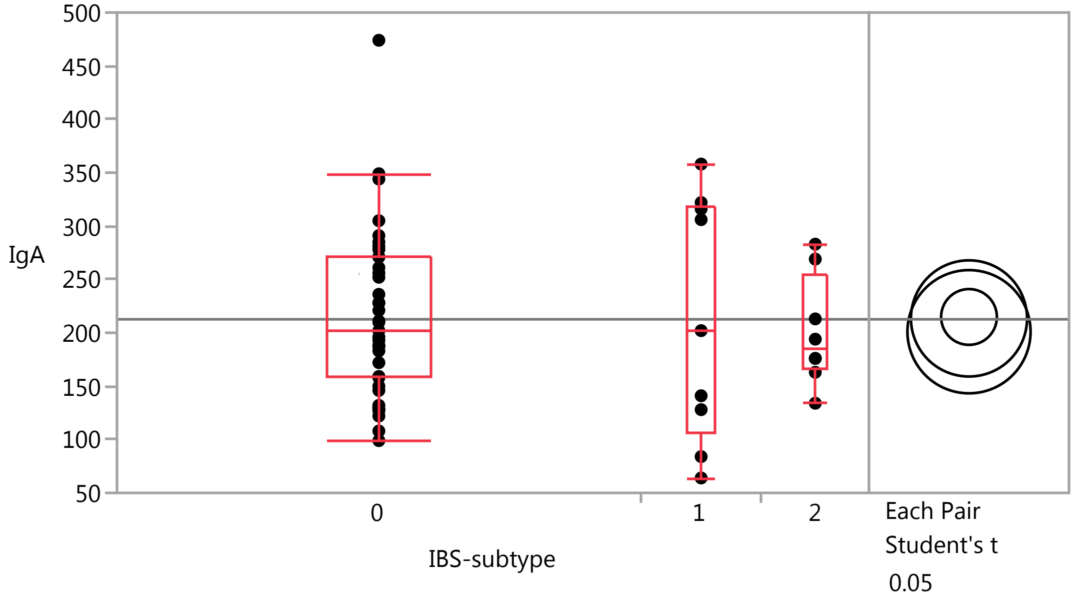


##### Quantiles

| Level | Minimum | 10% | 25% | Median | 75% | 90% | Maximum |
|---|---|---|---|---|---|---|---|
| 0 | 99 | 127 | 158 | 202 | 271 | 305 | 474 |
| 1 | 64 | 64 | 106 | 202 | 319 | 358 | 358 |
| 2 | 134 | 134 | 166.25 | 185 | 255 | 283 | 283 |

##### Means Comparisons

###### Comparisons for each pair using Student's t

###### Confidence Quantile

| t | Alpha |
|---|---|
| 2.00575 | 0.05 |

###### LSD Threshold Matrix

Abs(Dif)-LSD

| | 0 | 1 | 2 |
|---|---|---|---|
| 0 | -36.970 | -58.995 | -49.543 |
| 1 | -58.995 | -76.959 | -66.883 |
| 2 | -49.543 | -66.883 | -81.627 |

Positive values show pairs of means that are significantly different.

###### Connecting Letters Report

| Level | | Mean |
|---|---|---|
| 0 | A | 214.82051 |
| 1 | A | 213.44444 |
| 2 | A | 201.00000 |

Levels not connected by same letter are significantly different.

### Fit Group

#### Oneway Analysis of IgA By IBS-subtype Weight=1

##### Means Comparisons

###### Comparisons for each pair using Student's t

###### Ordered Differences Report

| Level | - Level | Difference | Std Err Dif | Lower CL | Upper CL | p-Value |
|---|---|---|---|---|---|---|
| 0 | 2 | 13.82051 | 31.59077 | -49.5425 | 77.18357 | 0.6635 |
| 1 | 2 | 12.44444 | 39.55003 | -66.8829 | 91.77176 | 0.7543 |
| 0 | 1 | 1.37607 | 30.09925 | -58.9954 | 61.74752 | 0.9637 |

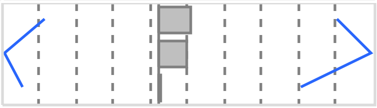
