## Supplementary Data 3. Weight by IBS for "Complete blood count with differential: An effective diagnostic for IBS subtype in the context of BMI?"

### Fit Group

#### Oneway Analysis of CRP By Weight IBS-subtype=0

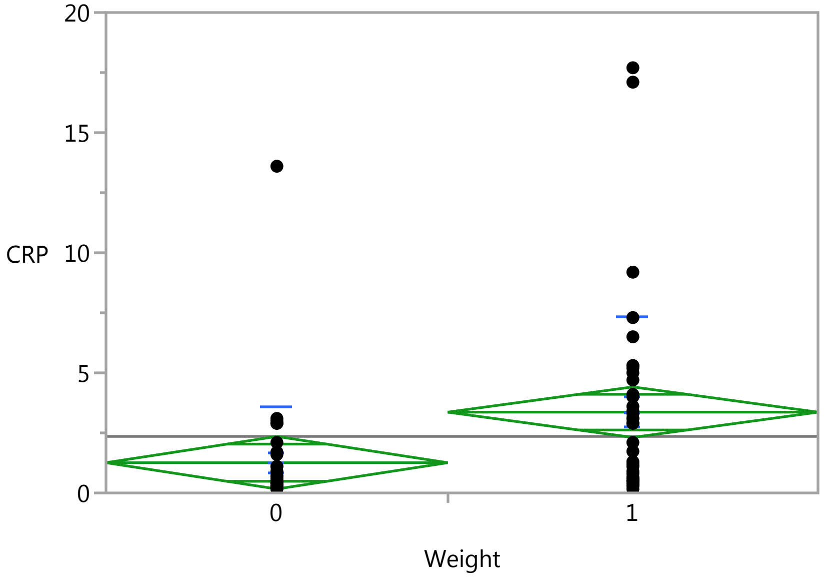


#### Oneway Anova

##### Summary of Fit

| | |
|---|---|
| Rsquare | 0.094715 |
| Adj Rsquare | 0.082314 |
| Root Mean Square Error | 3.29393 |
| Mean of Response | 2.3524 |
| Observations (or Sum Wgts) | 75 |

##### t Test

1-0

Assuming equal variances

| | | | |
|---|---|---|---|
| Difference | 2.10397 | t Ratio | 2.763624 |
| Std Err Dif | 0.76131 | DF | 73 |
| Upper CL Dif | 3.62126 | Prob > \|t\| | 0.0072* |
| Lower CL Dif | 0.58669 | Prob > t | 0.0036* |
| Confidence | 0.95 | Prob < t | 0.9964 |

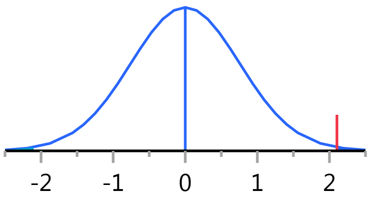


##### Analysis of Variance

| Source | DF | Sum of Squares | Mean Square | F Ratio | Prob > F |
|---|---|---|---|---|---|
| Weight | 1 | 82.86798 | 82.8680 | 7.6376 | 0.0072* |
| Error | 73 | 792.04839 | 10.8500 | | |
| C. Total | 74 | 874.91637 | | | |

##### Means for Oneway Anova

| Level | Number | Mean | Std Error | Lower 95% | Upper 95% |
|---|---|---|---|---|---|
| 0 | 36 | 1.25833 | 0.54899 | 0.1642 | 2.3525 |
| 1 | 39 | 3.36231 | 0.52745 | 2.3111 | 4.4135 |

Std Error uses a pooled estimate of error variance

### Fit Group

#### Oneway Analysis of CRP By Weight IBS-subtype=0

##### Means and Std Deviations

| Level | Number | Mean | Std Dev | Std Err Mean | Lower 95% | Upper 95% |
|---|---|---|---|---|---|---|
| 0 | 36 | 1.25833 | 2.32016 | 0.38669 | 0.4733 | 2.0434 |
| 1 | 39 | 3.36231 | 3.98562 | 0.63821 | 2.0703 | 4.6543 |

##### t Test

1-0

Assuming unequal variances

| | | | |
|---|---|---|---|
| Difference | 2.10397 | t Ratio | 2.819507 |
| Std Err Dif | 0.74622 | DF | 61.95653 |
| Upper CL Dif | 3.59567 | Prob > \|t\| | 0.0065* |
| Lower CL Dif | 0.61228 | Prob > t | 0.0032* |
| Confidence | 0.95 | Prob < t | 0.9968 |

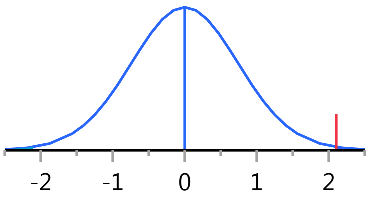


Missing Rows 2

#### Oneway Analysis of SerumCortisol By Weight IBS-subtype=0

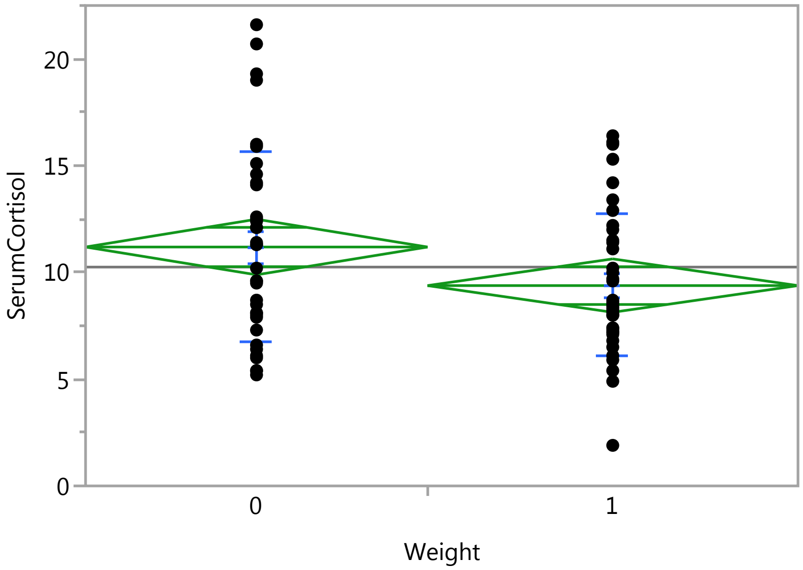


##### Oneway Anova

###### Summary of Fit

| | |
|---|---|
| Rsquare | 0.051678 |
| Adj Rsquare | 0.038687 |
| Root Mean Square Error | 3.919556 |
| Mean of Response | 10.24933 |
| Observations (or Sum Wgts) | 75 |

###### t Test

1-0

### Fit Group

#### Oneway Analysis of SerumCortisol By Weight IBS-subtype=0

##### Oneway Anova

###### t Test

Assuming equal variances

| | | | |
|---|---|---|---|
| Difference | -1.8068 | t Ratio | -1.99451 |
| Std Err Dif | 0.9059 | DF | 73 |
| Upper CL Dif | -0.0014 | Prob > \|t\| | 0.0498* |
| Lower CL Dif | -3.6123 | Prob > t | 0.9751 |
| Confidence | 0.95 | Prob < t | 0.0249* |

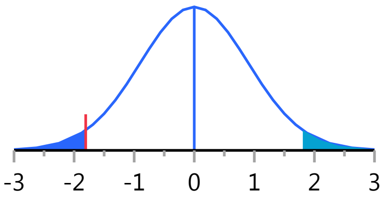


###### Analysis of Variance

| Source | DF | Sum of Squares | Mean Square | F Ratio | Prob > F |
|---|---|---|---|---|---|
| Weight | 1 | 61.1145 | 61.1145 | 3.9781 | 0.0498* |
| Error | 73 | 1121.4930 | 15.3629 | | |
| C. Total | 74 | 1182.6075 | | | |

###### Means for Oneway Anova

| Level | Number | Mean | Std Error | Lower 95% | Upper 95% |
|---|---|---|---|---|---|
| 0 | 36 | 11.1889 | 0.65326 | 9.8869 | 12.491 |
| 1 | 39 | 9.3821 | 0.62763 | 8.1312 | 10.633 |

Std Error uses a pooled estimate of error variance

##### Means and Std Deviations

| Level | Number | Mean | Std Dev | Std Err Mean | Lower 95% | Upper 95% |
|---|---|---|---|---|---|---|
| 0 | 36 | 11.1889 | 4.47078 | 0.74513 | 9.6762 | 12.702 |
| 1 | 39 | 9.3821 | 3.33213 | 0.53357 | 8.3019 | 10.462 |

##### t Test

1-0

Assuming unequal variances

| | | | |
|---|---|---|---|
| Difference | -1.8068 | t Ratio | -1.97152 |
| Std Err Dif | 0.9165 | DF | 64.48057 |
| Upper CL Dif | 0.0238 | Prob > \|t\| | 0.0530 |
| Lower CL Dif | -3.6374 | Prob > t | 0.9735 |
| Confidence | 0.95 | Prob < t | 0.0265* |

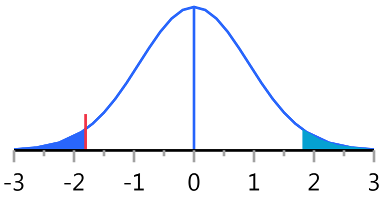


Missing Rows 2

#### Oneway Analysis of Lymphocytes By Weight IBS-subtype=0

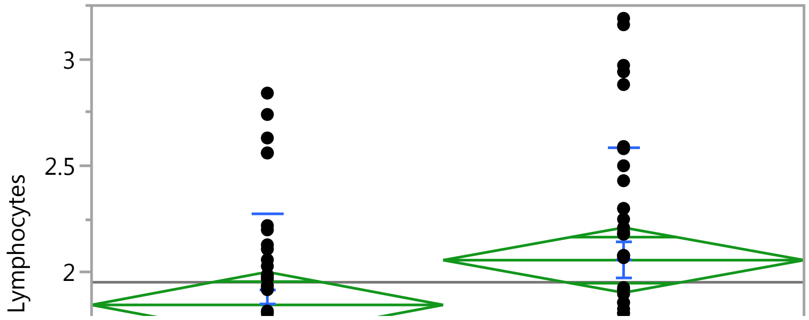

#### Fit Group

##### Oneway Analysis of Lymphocytes By Weight IBS-subtype=0

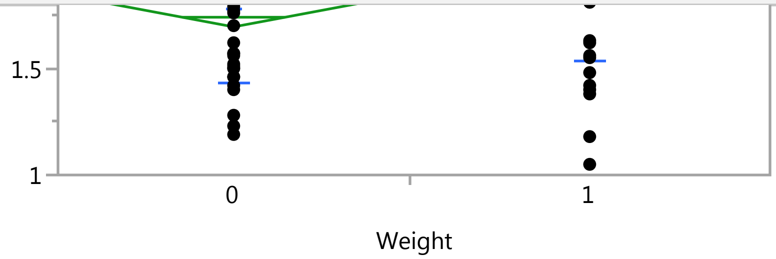


##### Oneway Anova

###### Summary of Fit

| | |
|---|---|
| Rsquare | 0.047045 |
| Adj Rsquare | 0.034339 |
| Root Mean Square Error | 0.4777 |
| Mean of Response | 1.954805 |
| Observations (or Sum Wgts) | 77 |

###### t Test

1-0

Assuming equal variances

| | | | |
|---|---|---|---|
| Difference | 0.20952 | t Ratio | 1.924202 |
| Std Err Dif | 0.10889 | DF | 75 |
| Upper CL Dif | 0.42644 | Prob > \|t\| | 0.0581 |
| Lower CL Dif | -0.00739 | Prob > t | 0.0291* |
| Confidence | 0.95 | Prob < t | 0.9709 |

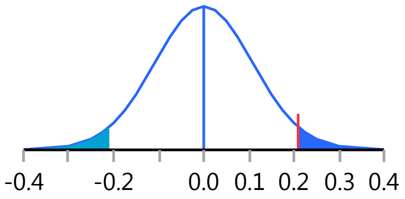


###### Analysis of Variance

| Source | DF | Sum of Squares | Mean Square | F Ratio | Prob > F |
|---|---|---|---|---|---|
| Weight | 1 | 0.844914 | 0.844914 | 3.7026 | 0.0581 |
| Error | 75 | 17.114809 | 0.228197 | | |
| C. Total | 76 | 17.959722 | | | |

###### Means for Oneway Anova

| Level | Number | Mean | Std Error | Lower 95% | Upper 95% |
|---|---|---|---|---|---|
| 0 | 38 | 1.84868 | 0.07749 | 1.6943 | 2.0031 |
| 1 | 39 | 2.05821 | 0.07649 | 1.9058 | 2.2106 |

Std Error uses a pooled estimate of error variance

##### Means and Std Deviations

| Level | Number | Mean | Std Dev | Std Err Mean | Lower 95% | Upper 95% |
|---|---|---|---|---|---|---|
| 0 | 38 | 1.84868 | 0.421831 | 0.06843 | 1.7100 | 1.9873 |
| 1 | 39 | 2.05821 | 0.526432 | 0.08430 | 1.8876 | 2.2289 |

##### t Test

1-0

#### Fit Group

##### Oneway Analysis of Lymphocytes By Weight IBS-subtype=0

###### t Test

Assuming unequal variances

| | | | |
|---|---|---|---|
| Difference | 0.20952 | t Ratio | 1.92973 |
| Std Err Dif | 0.10858 | DF | 72.32699 |
| Upper CL Dif | 0.42595 | Prob > \|t\| | 0.0576 |
| Lower CL Dif | -0.00690 | Prob > t | 0.0288* |
| Confidence | 0.95 | Prob < t | 0.9712 |

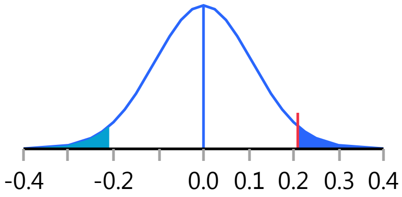


##### Oneway Analysis of ESR By Weight IBS-subtype=0

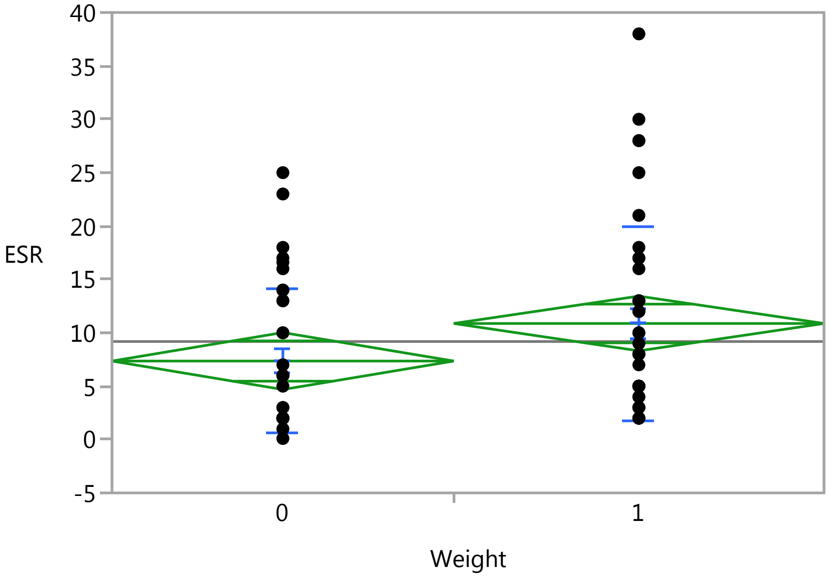


##### Oneway Anova

###### Summary of Fit

| | |
|---|---|
| Rsquare | 0.046913 |
| Adj Rsquare | 0.033857 |
| Root Mean Square Error | 8.032169 |
| Mean of Response | 9.182667 |
| Observations (or Sum Wgts) | 75 |

###### t Test

1-0

Assuming equal variances

| | | | |
|---|---|---|---|
| Difference | 3.5190 | t Ratio | 1.895577 |
| Std Err Dif | 1.8564 | DF | 73 |
| Upper CL Dif | 7.2189 | Prob > \|t\| | 0.0620 |
| Lower CL Dif | -0.1809 | Prob > t | 0.0310* |
| Confidence | 0.95 | Prob < t | 0.9690 |

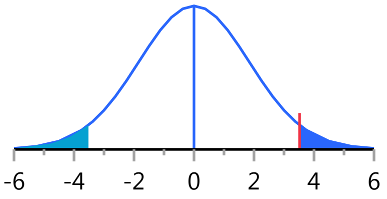

### Fit Group

#### Oneway Analysis of ESR By Weight IBS-subtype=0

##### Oneway Anova

###### Analysis of Variance

| Source | DF | Sum of Squares | Mean Square | F Ratio | Prob > F |
|---|---|---|---|---|---|
| Weight | 1 | 231.8188 | 231.819 | 3.5932 | 0.0620 |
| Error | 73 | 4709.6487 | 64.516 | | |
| C. Total | 74 | 4941.4675 | | | |

###### Means for Oneway Anova

| Level | Number | Mean | Std Error | Lower 95% | Upper 95% |
|---|---|---|---|---|---|
| 0 | 36 | 7.3528 | 1.3387 | 4.6848 | 10.021 |
| 1 | 39 | 10.8718 | 1.2862 | 8.3084 | 13.435 |

Std Error uses a pooled estimate of error variance

##### Means and Std Deviations

| Level | Number | Mean | Std Dev | Std Err Mean | Lower 95% | Upper 95% |
|---|---|---|---|---|---|---|
| 0 | 36 | 7.3528 | 6.77662 | 1.1294 | 5.0599 | 9.646 |
| 1 | 39 | 10.8718 | 9.03554 | 1.4468 | 7.9428 | 13.801 |

##### t Test

1-0

Assuming unequal variances

| | | | |
|---|---|---|---|
| Difference | 3.5190 | t Ratio | 1.917219 |
| Std Err Dif | 1.8355 | DF | 70.14357 |
| Upper CL Dif | 7.1796 | Prob > \|t\| | 0.0593 |
| Lower CL Dif | -0.1416 | Prob > t | 0.0296* |
| Confidence | 0.95 | Prob < t | 0.9704 |

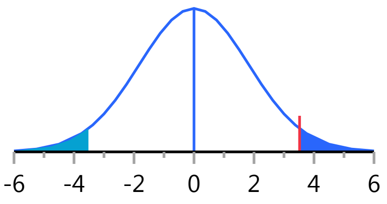


Missing Rows 2

#### Oneway Analysis of PlateletCount By Weight IBS-subtype=0

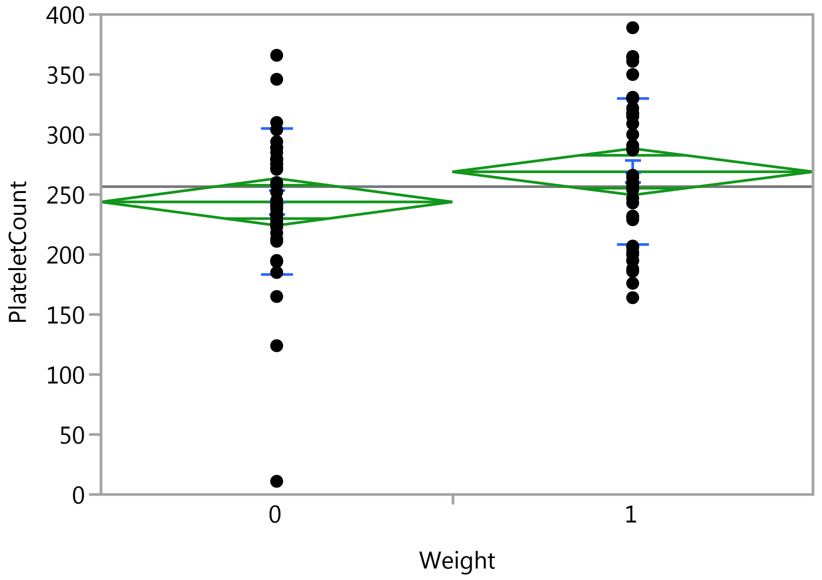

### Fit Group

#### Oneway Analysis of PlateletCount By Weight IBS-subtype=0

##### Oneway Anova

###### Summary of Fit

| | |
|---|---|
| Rsquare | 0.042069 |
| Adj Rsquare | 0.029296 |
| Root Mean Square Error | 60.75455 |
| Mean of Response | 256.5455 |
| Observations (or Sum Wgts) | 77 |

###### t Test

1-0

Assuming equal variances

| | | | |
|---|---|---|---|
| Difference | 25.133 | t Ratio | 1.81486 |
| Std Err Dif | 13.848 | DF | 75 |
| Upper CL Dif | 52.720 | Prob > \|t\| | 0.0735 |
| Lower CL Dif | -2.455 | Prob > t | 0.0368* |
| Confidence | 0.95 | Prob < t | 0.9632 |

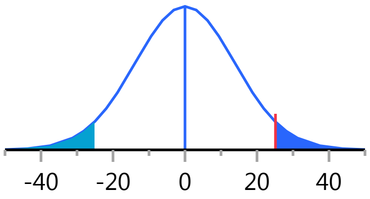


###### Analysis of Variance

| Source | DF | Sum of Squares | Mean Square | F Ratio | Prob > F |
|---|---|---|---|---|---|
| Weight | 1 | 12157.48 | 12157.5 | 3.2937 | 0.0735 |
| Error | 75 | 276833.61 | 3691.1 | | |
| C. Total | 76 | 288991.09 | | | |

###### Means for Oneway Anova

| Level | Number | Mean | Std Error | Lower 95% | Upper 95% |
|---|---|---|---|---|---|
| 0 | 38 | 243.816 | 9.8557 | 224.18 | 263.45 |
| 1 | 39 | 268.949 | 9.7285 | 249.57 | 288.33 |

Std Error uses a pooled estimate of error variance

##### Means and Std Deviations

| Level | Number | Mean | Std Dev | Std Err Mean | Lower 95% | Upper 95% |
|---|---|---|---|---|---|---|
| 0 | 38 | 243.816 | 60.8733 | 9.8750 | 223.81 | 263.82 |
| 1 | 39 | 268.949 | 60.6387 | 9.7100 | 249.29 | 288.61 |

##### t Test

1-0

Assuming unequal variances

| | | | |
|---|---|---|---|
| Difference | 25.133 | t Ratio | 1.814767 |
| Std Err Dif | 13.849 | DF | 74.93173 |
| Upper CL Dif | 52.722 | Prob > \|t\| | 0.0736 |
| Lower CL Dif | -2.456 | Prob > t | 0.0368* |
| Confidence | 0.95 | Prob < t | 0.9632 |

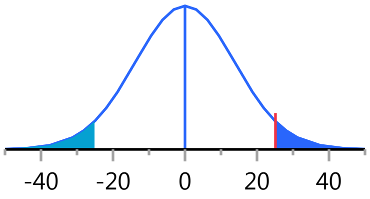

#### Fit Group

##### Oneway Analysis of sCD14 By Weight IBS-subtype=0

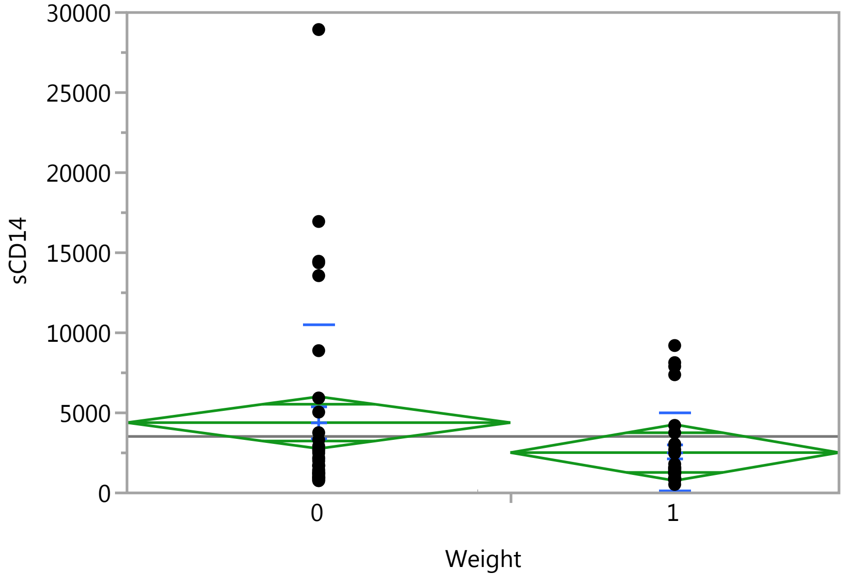


##### Oneway Anova

###### Summary of Fit

| | |
|---|---|
| Rsquare | 0.037399 |
| Adj Rsquare | 0.02212 |
| Root Mean Square Error | 4806.857 |
| Mean of Response | 3526.432 |
| Observations (or Sum Wgts) | 65 |

###### t Test

1-0

Assuming equal variances

| | | | |
|---|---|---|---|
| Difference | -1871.1 | t Ratio | -1.56451 |
| Std Err Dif | 1196.0 | DF | 63 |
| Upper CL Dif | 518.8 | Prob > \|t\| | 0.1227 |
| Lower CL Dif | -4261.1 | Prob > t | 0.9386 |
| Confidence | 0.95 | Prob < t | 0.0614 |

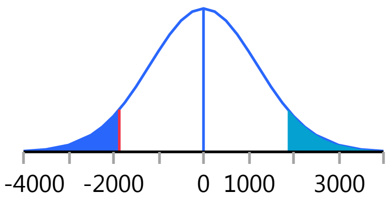


###### Analysis of Variance

| Source | DF | Sum of Squares | Mean Square | F Ratio | Prob > F |
|---|---|---|---|---|---|
| Weight | 1 | 56556386.8 | 56556387 | 2.4477 | 0.1227 |
| Error | 63 | 1455670122 | 23105875 | | |
| C. Total | 64 | 1512226509 | | | |

###### Means for Oneway Anova

| Level | Number | Mean | Std Error | Lower 95% | Upper 95% |
|---|---|---|---|---|---|
| 0 | 35 | 4390.03 | 812.51 | 2766.4 | 6013.7 |
| 1 | 30 | 2518.90 | 877.61 | 765.1 | 4272.7 |

Std Error uses a pooled estimate of error variance

#### Fit Group

##### Oneway Analysis of sCD14 By Weight IBS-subtype=0

###### Means and Std Deviations

| Level | Number | Mean | Std Dev | Std Err Mean | Lower 95% | Upper 95% |
|---|---|---|---|---|---|---|
| 0 | 35 | 4390.03 | 6145.62 | 1038.8 | 2278.9 | 6501.1 |
| 1 | 30 | 2518.90 | 2432.08 | 444.0 | 1610.7 | 3427.1 |

###### t Test

1-0

Assuming unequal variances

| | | | |
|---|---|---|---|
| Difference | -1871.1 | t Ratio | -1.65627 |
| Std Err Dif | 1129.7 | DF | 45.76822 |
| Upper CL Dif | 403.2 | Prob > \|t\| | 0.1045 |
| Lower CL Dif | -4145.4 | Prob > t | 0.9477 |
| Confidence | 0.95 | Prob < t | 0.0523 |

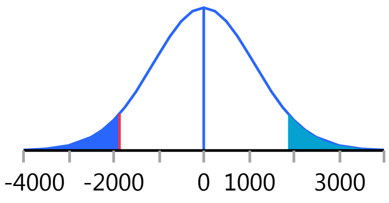


Missing Rows 12

##### Oneway Analysis of MPV By Weight IBS-subtype=0

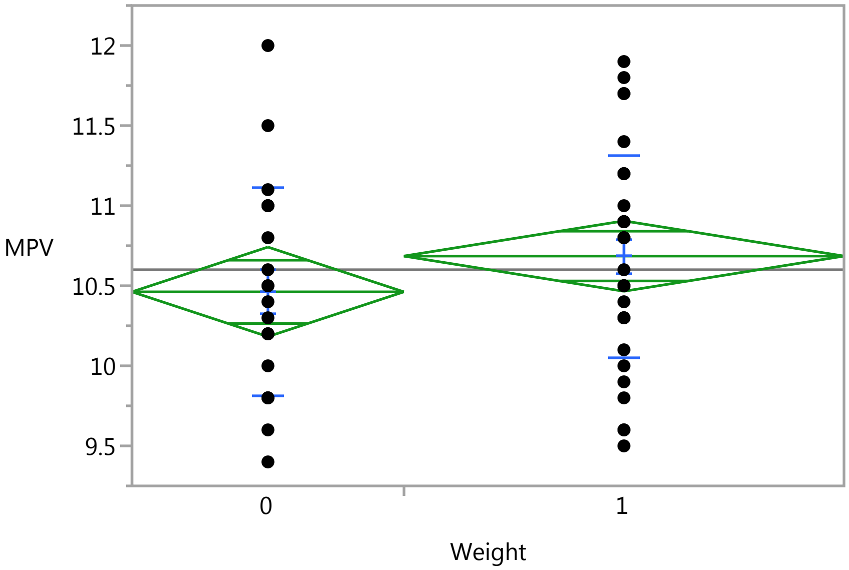


###### Oneway Anova

###### Summary of Fit

| | |
|---|---|
| Rsquare | 0.029077 |
| Adj Rsquare | 0.010757 |
| Root Mean Square Error | 0.638869 |
| Mean of Response | 10.6 |
| Observations (or Sum Wgts) | 55 |

###### t Test

1-0

### Fit Group

#### Oneway Analysis of MPV By Weight IBS-subtype=0

##### Oneway Anova

###### t Test

Assuming equal variances

| | | | |
|---|---|---|---|
| Difference | 0.22339 | t Ratio | 1.259848 |
| Std Err Dif | 0.17731 | DF | 53 |
| Upper CL Dif | 0.57904 | Prob > \|t\| | 0.2132 |
| Lower CL Dif | -0.13226 | Prob > t | 0.1066 |
| Confidence | 0.95 | Prob < t | 0.8934 |

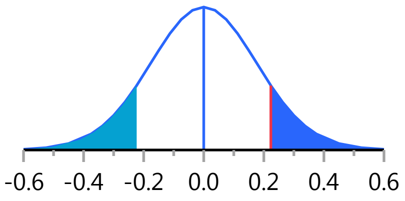


###### Analysis of Variance

| Source | DF | Sum of Squares | Mean Square | F Ratio | Prob > F |
|---|---|---|---|---|---|
| Weight | 1 | 0.647829 | 0.647829 | 1.5872 | 0.2132 |
| Error | 53 | 21.632171 | 0.408154 | | |
| C. Total | 54 | 22.280000 | | | |

###### Means for Oneway Anova

| Level | Number | Mean | Std Error | Lower 95% | Upper 95% |
|---|---|---|---|---|---|
| 0 | 21 | 10.4619 | 0.13941 | 10.182 | 10.742 |
| 1 | 34 | 10.6853 | 0.10957 | 10.466 | 10.905 |

Std Error uses a pooled estimate of error variance

##### Means and Std Deviations

| Level | Number | Mean | Std Dev | Std Err Mean | Lower 95% | Upper 95% |
|---|---|---|---|---|---|---|
| 0 | 21 | 10.4619 | 0.651518 | 0.14217 | 10.165 | 10.758 |
| 1 | 34 | 10.6853 | 0.631080 | 0.10823 | 10.465 | 10.905 |

##### t Test

1-0

Assuming unequal variances

| | | | |
|---|---|---|---|
| Difference | 0.22339 | t Ratio | 1.250216 |
| Std Err Dif | 0.17868 | DF | 41.45856 |
| Upper CL Dif | 0.58412 | Prob > \|t\| | 0.2182 |
| Lower CL Dif | -0.13734 | Prob > t | 0.1091 |
| Confidence | 0.95 | Prob < t | 0.8909 |

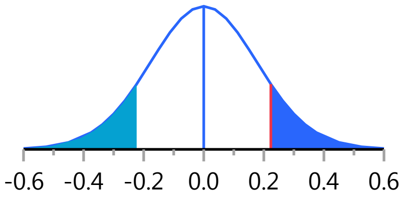


Missing Rows 22

#### Oneway Analysis of IgM By Weight IBS-subtype=0

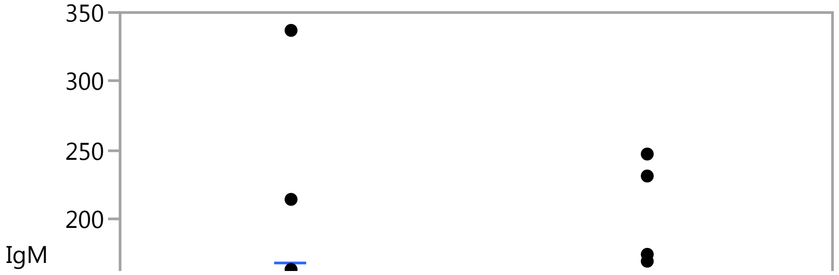

### Fit Group

#### Oneway Analysis of IgM By Weight IBS-subtype=0

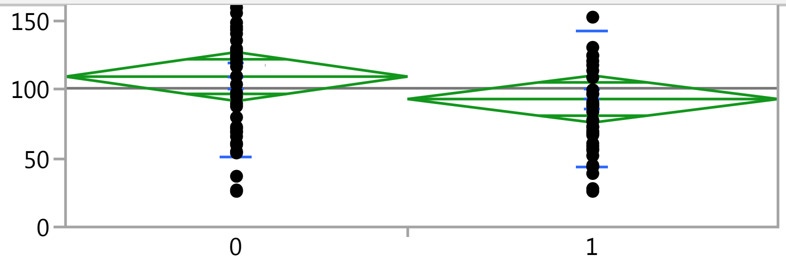


#### Oneway Anova

##### Summary of Fit

| | |
|---|---|
| Rsquare | 0.023208 |
| Adj Rsquare | 0.009828 |
| Root Mean Square Error | 53.74347 |
| Mean of Response | 101.16 |
| Observations (or Sum Wgts) | 75 |

##### t Test

1-0

Assuming equal variances

| | | | |
|---|---|---|---|
| Difference | -16.359 | t Ratio | -1.31699 |
| Std Err Dif | 12.421 | DF | 73 |
| Upper CL Dif | 8.397 | Prob > \|t\| | 0.1920 |
| Lower CL Dif | -41.115 | Prob > t | 0.9040 |
| Confidence | 0.95 | Prob < t | 0.0960 |

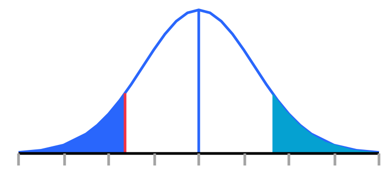


##### Analysis of Variance

| Source | DF | Sum of Squares | Mean Square | F Ratio | Prob > F |
|---|---|---|---|---|---|
| Weight | 1 | 5009.77 | 5009.77 | 1.7345 | 0.1920 |
| Error | 73 | 210850.31 | 2888.36 | | |
| C. Total | 74 | 215860.08 | | | |

##### Means for Oneway Anova

| Level | Number | Mean | Std Error | Lower 95% | Upper 95% |
|---|---|---|---|---|---|
| 0 | 36 | 109.667 | 8.9572 | 91.815 | 127.52 |
| 1 | 39 | 93.308 | 8.6058 | 76.156 | 110.46 |

Std Error uses a pooled estimate of error variance

#### Means and Std Deviations

| Level | Number | Mean | Std Dev | Std Err Mean | Lower 95% | Upper 95% |
|---|---|---|---|---|---|---|
| 0 | 36 | 109.667 | 57.9147 | 9.6525 | 90.071 | 129.26 |
| 1 | 39 | 93.308 | 49.5921 | 7.9411 | 77.232 | 109.38 |

#### t Test

1-0

#### Fit Group

##### Oneway Analysis of IgM By Weight IBS-subtype=0

###### t Test

Assuming unequal variances

| | | | |
|---|---|---|---|
| Difference | -16.359 | t Ratio | -1.3088 |
| Std Err Dif | 12.499 | DF | 69.20988 |
| Upper CL Dif | 8.575 | Prob > \|t\| | 0.1949 |
| Lower CL Dif | -41.293 | Prob > t | 0.9025 |
| Confidence | 0.95 | Prob < t | 0.0975 |

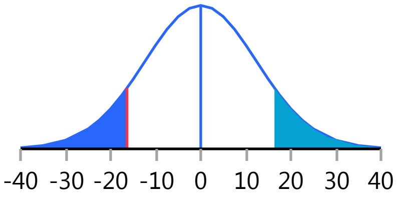


Missing Rows 2

##### Oneway Analysis of Neutrophil_PCT By Weight IBS-subtype=0

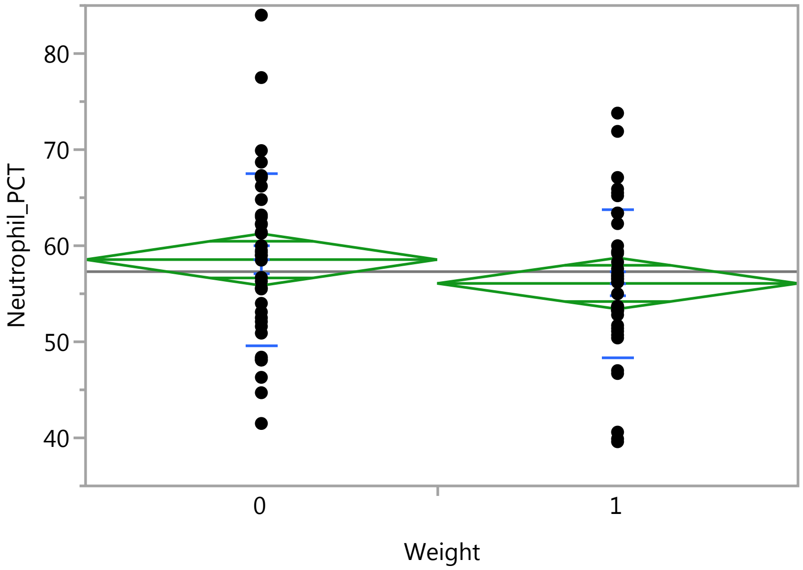


###### Oneway Anova

###### Summary of Fit

| | |
|---|---|
| Rsquare | 0.022063 |
| Adj Rsquare | 0.009024 |
| Root Mean Square Error | 8.358519 |
| Mean of Response | 57.3 |
| Observations (or Sum Wgts) | 77 |

###### t Test

1-0

Assuming equal variances

| | | | |
|---|---|---|---|
| Difference | -2.4783 | t Ratio | -1.3008 |
| Std Err Dif | 1.9052 | DF | 75 |
| Upper CL Dif | 1.3171 | Prob > \|t\| | 0.1973 |
| Lower CL Dif | -6.2738 | Prob > t | 0.9013 |
| Confidence | 0.95 | Prob < t | 0.0987 |

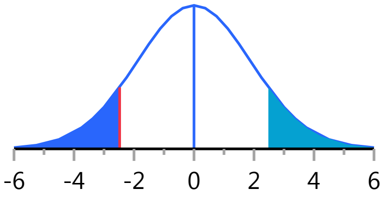

### Fit Group

#### Oneway Analysis of Neutrophil_PCT By Weight IBS-subtype=0

##### Oneway Anova

###### Analysis of Variance

| Source | DF | Sum of Squares | Mean Square | F Ratio | Prob > F |
|---|---|---|---|---|---|
| Weight | 1 | 118.2168 | 118.217 | 1.6921 | 0.1973 |
| Error | 75 | 5239.8632 | 69.865 | | |
| C. Total | 76 | 5358.0800 | | | |

###### Means for Oneway Anova

| Level | Number | Mean | Std Error | Lower 95% | Upper 95% |
|---|---|---|---|---|---|
| 0 | 38 | 58.5553 | 1.3559 | 55.854 | 61.256 |
| 1 | 39 | 56.0769 | 1.3384 | 53.411 | 58.743 |

Std Error uses a pooled estimate of error variance

##### Means and Std Deviations

| Level | Number | Mean | Std Dev | Std Err Mean | Lower 95% | Upper 95% |
|---|---|---|---|---|---|---|
| 0 | 38 | 58.5553 | 8.95966 | 1.4534 | 55.610 | 61.500 |
| 1 | 39 | 56.0769 | 7.72840 | 1.2375 | 53.572 | 58.582 |

##### t Test

1-0

Assuming unequal variances

| | | | |
|---|---|---|---|
| Difference | -2.4783 | t Ratio | -1.29829 |
| Std Err Dif | 1.9089 | DF | 72.82557 |
| Upper CL Dif | 1.3263 | Prob > \|t\| | 0.1983 |
| Lower CL Dif | -6.2830 | Prob > t | 0.9009 |
| Confidence | 0.95 | Prob < t | 0.0991 |

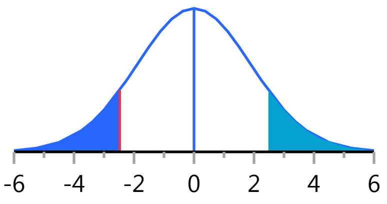


#### Oneway Analysis of IgG By Weight IBS-subtype=0

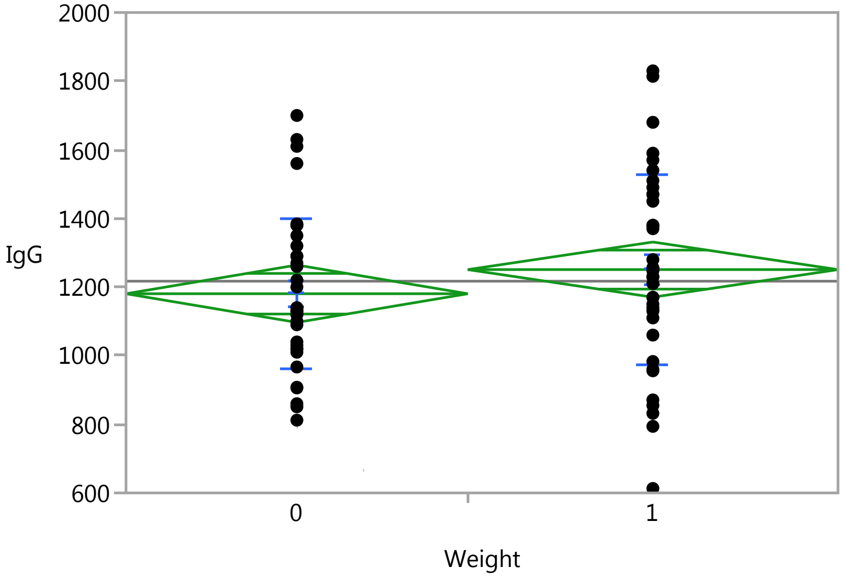

### Fit Group

#### Oneway Analysis of IgG By Weight IBS-subtype=0

##### Oneway Anova

###### Summary of Fit

| | |
|---|---|
| Rsquare | 0.0196 |
| Adj Rsquare | 0.00617 |
| Root Mean Square Error | 251.8583 |
| Mean of Response | 1217.04 |
| Observations (or Sum Wgts) | 75 |

###### t Test

1-0

Assuming equal variances

| | | | |
|---|---|---|---|
| Difference | 70.32 | t Ratio | 1.208069 |
| Std Err Dif | 58.21 | DF | 73 |
| Upper CL Dif | 186.34 | Prob > \|t\| | 0.2309 |
| Lower CL Dif | -45.69 | Prob > t | 0.1155 |
| Confidence | 0.95 | Prob < t | 0.8845 |

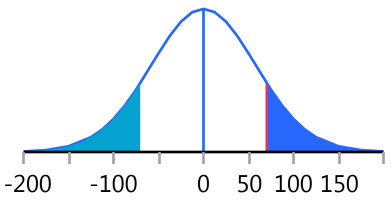


###### Analysis of Variance

| Source | DF | Sum of Squares | Mean Square | F Ratio | Prob > F |
|---|---|---|---|---|---|
| Weight | 1 | 92575.5 | 92575.5 | 1.4594 | 0.2309 |
| Error | 73 | 4630579.3 | 63432.6 | | |
| C. Total | 74 | 4723154.9 | | | |

###### Means for Oneway Anova

| Level | Number | Mean | Std Error | Lower 95% | Upper 95% |
|---|---|---|---|---|---|
| 0 | 36 | 1180.47 | 41.976 | 1096.8 | 1264.1 |
| 1 | 39 | 1250.79 | 40.330 | 1170.4 | 1331.2 |

Std Error uses a pooled estimate of error variance

##### Means and Std Deviations

| Level | Number | Mean | Std Dev | Std Err Mean | Lower 95% | Upper 95% |
|---|---|---|---|---|---|---|
| 0 | 36 | 1180.47 | 221.081 | 36.847 | 1105.7 | 1255.3 |
| 1 | 39 | 1250.79 | 277.199 | 44.387 | 1160.9 | 1340.7 |

##### t Test

1-0

Assuming unequal variances

| | | | |
|---|---|---|---|
| Difference | 70.32 | t Ratio | 1.219013 |
| Std Err Dif | 57.69 | DF | 71.53533 |
| Upper CL Dif | 185.33 | Prob > \|t\| | 0.2268 |
| Lower CL Dif | -44.69 | Prob > t | 0.1134 |
| Confidence | 0.95 | Prob < t | 0.8866 |

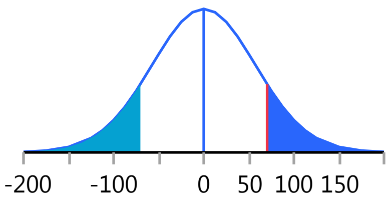


Missing Rows 2

#### Fit Group

##### Oneway Analysis of IgE By Weight IBS-subtype=0

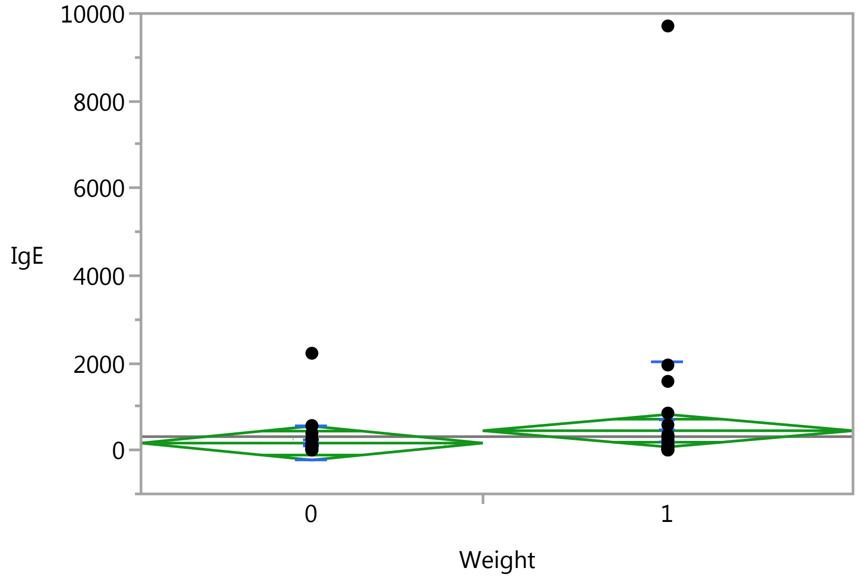


##### Oneway Anova

###### Summary of Fit

| | |
|---|---|
| Rsquare | 0.014878 |
| Adj Rsquare | 0.001384 |
| Root Mean Square Error | 1167.788 |
| Mean of Response | 311.4253 |
| Observations (or Sum Wgts) | 75 |

###### t Test

1-0

Assuming equal variances

| | | | |
|---|---|---|---|
| Difference | 283.40 | t Ratio | 1.050011 |
| Std Err Dif | 269.91 | DF | 73 |
| Upper CL Dif | 821.32 | Prob > \|t\| | 0.2972 |
| Lower CL Dif | -254.52 | Prob > t | 0.1486 |
| Confidence | 0.95 | Prob < t | 0.8514 |

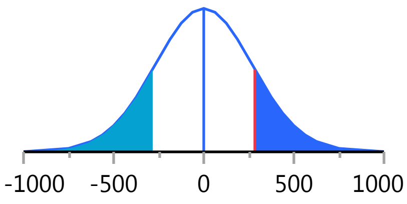


###### Analysis of Variance

| Source | DF | Sum of Squares | Mean Square | F Ratio | Prob > F |
|---|---|---|---|---|---|
| Weight | 1 | 1503544 | 1503544 | 1.1025 | 0.2972 |
| Error | 73 | 99552262 | 1363730 | | |
| C. Total | 74 | 101055806 | | | |

###### Means for Oneway Anova

| Level | Number | Mean | Std Error | Lower 95% | Upper 95% |
|---|---|---|---|---|---|
| 0 | 36 | 164.056 | 194.63 | -223.8 | 551.96 |
| 1 | 39 | 447.459 | 187.00 | 74.8 | 820.14 |

Std Error uses a pooled estimate of error variance

### Fit Group

#### Oneway Analysis of IgE By Weight IBS-subtype=0

##### Means and Std Deviations

| Level | Number | Mean | Std Dev | Std Err Mean | Lower 95% | Upper 95% |
|---|---|---|---|---|---|---|
| 0 | 36 | 164.056 | 380.08 | 63.35 | 35.45 | 292.66 |
| 1 | 39 | 447.459 | 1576.94 | 252.51 | -63.73 | 958.64 |

##### t Test

1-0

Assuming unequal variances

| | | | |
|---|---|---|---|
| Difference | 283.40 | t Ratio | 1.088602 |
| Std Err Dif | 260.34 | DF | 42.74972 |
| Upper CL Dif | 808.51 | Prob > \|t\| | 0.2824 |
| Lower CL Dif | -241.71 | Prob > t | 0.1412 |
| Confidence | 0.95 | Prob < t | 0.8588 |

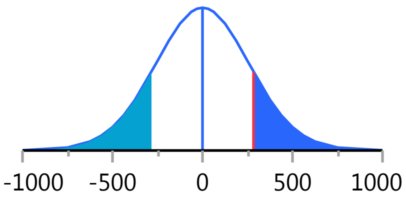


Missing Rows 2

#### Oneway Analysis of Lymphocytes_PCT By Weight IBS-subtype=0

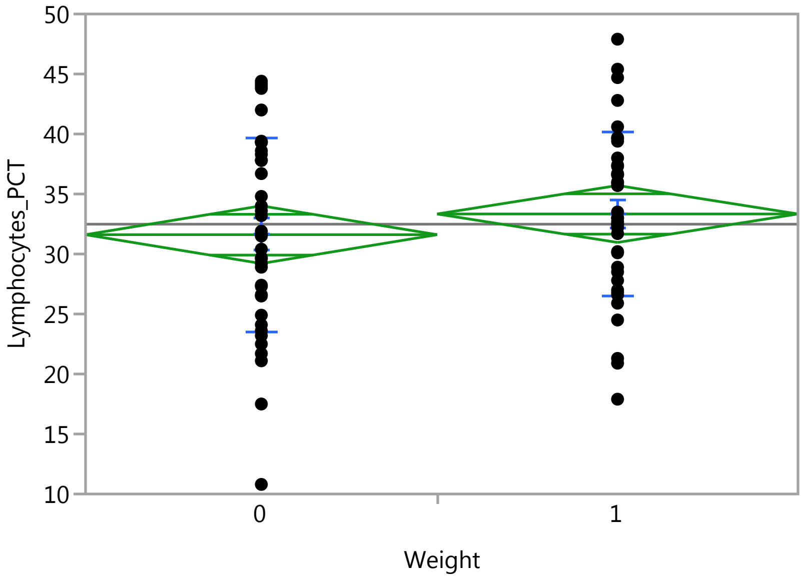


##### Oneway Anova

###### Summary of Fit

| | |
|---|---|
| Rsquare | 0.013613 |
| Adj Rsquare | 0.000461 |
| Root Mean Square Error | 7.440148 |
| Mean of Response | 32.48442 |
| Observations (or Sum Wgts) | 77 |

###### t Test

1-0

### Fit Group

#### Oneway Analysis of Lymphocytes_PCT By Weight IBS-subtype=0

##### Oneway Anova

###### t Test

Assuming equal variances

| | | | |
|---|---|---|---|
| Difference | 1.7254 | t Ratio | 1.017372 |
| Std Err Dif | 1.6959 | DF | 75 |
| Upper CL Dif | 5.1038 | Prob > \|t\| | 0.3122 |
| Lower CL Dif | -1.6531 | Prob > t | 0.1561 |
| Confidence | 0.95 | Prob < t | 0.8439 |

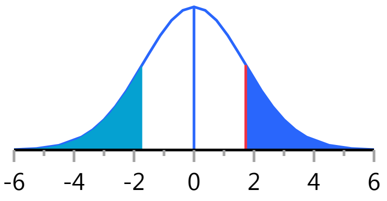


###### Analysis of Variance

| Source | DF | Sum of Squares | Mean Square | F Ratio | Prob > F |
|---|---|---|---|---|---|
| Weight | 1 | 57.2958 | 57.2958 | 1.0350 | 0.3122 |
| Error | 75 | 4151.6855 | 55.3558 | | |
| C. Total | 76 | 4208.9813 | | | |

###### Means for Oneway Anova

| Level | Number | Mean | Std Error | Lower 95% | Upper 95% |
|---|---|---|---|---|---|
| 0 | 38 | 31.6105 | 1.2070 | 29.206 | 34.015 |
| 1 | 39 | 33.3359 | 1.1914 | 30.963 | 35.709 |

Std Error uses a pooled estimate of error variance

##### Means and Std Deviations

| Level | Number | Mean | Std Dev | Std Err Mean | Lower 95% | Upper 95% |
|---|---|---|---|---|---|---|
| 0 | 38 | 31.6105 | 8.05333 | 1.3064 | 28.963 | 34.258 |
| 1 | 39 | 33.3359 | 6.79010 | 1.0873 | 31.135 | 35.537 |

##### t Test

1-0

Assuming unequal variances

| | | | |
|---|---|---|---|
| Difference | 1.7254 | t Ratio | 1.015112 |
| Std Err Dif | 1.6997 | DF | 72.25472 |
| Upper CL Dif | 5.1134 | Prob > \|t\| | 0.3134 |
| Lower CL Dif | -1.6627 | Prob > t | 0.1567 |
| Confidence | 0.95 | Prob < t | 0.8433 |

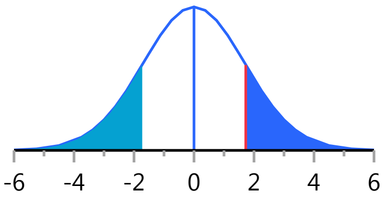


#### Oneway Analysis of Eosinophils By Weight IBS-subtype=0

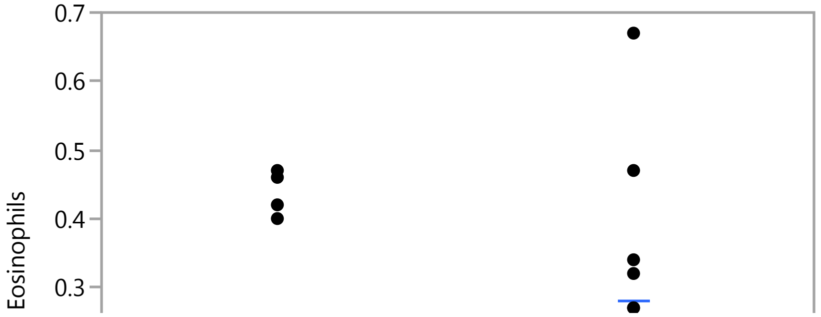

#### Fit Group

##### Oneway Analysis of Eosinophils By Weight IBS-subtype=0

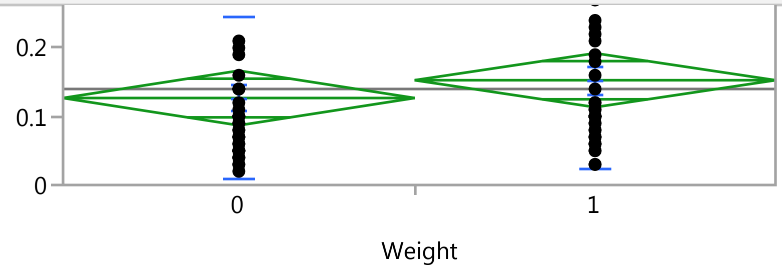


#### Oneway Anova

##### Summary of Fit

| | |
|---|---|
| Rsquare | 0.011242 |
| Adj Rsquare | -0.00194 |
| Root Mean Square Error | 0.123419 |
| Mean of Response | 0.14 |
| Observations (or Sum Wgts) | 77 |

##### t Test

1-0

Assuming equal variances

| | | | |
|---|---|---|---|
| Difference | 0.02598 | t Ratio | 0.923444 |
| Std Err Dif | 0.02813 | DF | 75 |
| Upper CL Dif | 0.08202 | Prob > \|t\| | 0.3587 |
| Lower CL Dif | -0.03006 | Prob > t | 0.1794 |
| Confidence | 0.95 | Prob < t | 0.8206 |

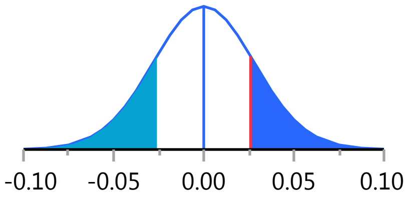


##### Analysis of Variance

| Source | DF | Sum of Squares | Mean Square | F Ratio | Prob > F |
|---|---|---|---|---|---|
| Weight | 1 | 0.0129892 | 0.012989 | 0.8527 | 0.3587 |
| Error | 75 | 1.1424108 | 0.015232 | | |
| C. Total | 76 | 1.1554000 | | | |

##### Means for Oneway Anova

| Level | Number | Mean | Std Error | Lower 95% | Upper 95% |
|---|---|---|---|---|---|
| 0 | 38 | 0.126842 | 0.02002 | 0.08696 | 0.16673 |
| 1 | 39 | 0.152821 | 0.01976 | 0.11345 | 0.19219 |

Std Error uses a pooled estimate of error variance

#### Means and Std Deviations

| Level | Number | Mean | Std Dev | Std Err Mean | Lower 95% | Upper 95% |
|---|---|---|---|---|---|---|
| 0 | 38 | 0.126842 | 0.118393 | 0.01921 | 0.08793 | 0.16576 |
| 1 | 39 | 0.152821 | 0.128123 | 0.02052 | 0.11129 | 0.19435 |

#### t Test

1-0

### Fit Group

#### Oneway Analysis of Eosinophils By Weight IBS-subtype=0

##### t Test

Assuming unequal variances

| | | | |
|---|---|---|---|
| Difference | 0.02598 | t Ratio | 0.924403 |
| Std Err Dif | 0.02810 | DF | 74.79327 |
| Upper CL Dif | 0.08196 | Prob > \|t\| | 0.3582 |
| Lower CL Dif | -0.03001 | Prob > t | 0.1791 |
| Confidence | 0.95 | Prob < t | 0.8209 |

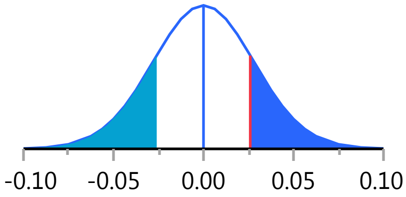


#### Oneway Analysis of LBP By Weight IBS-subtype=0

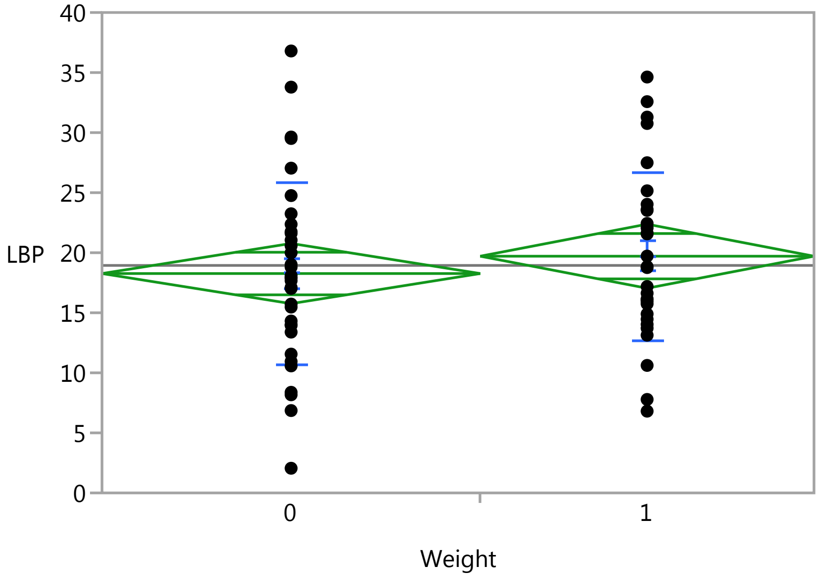


##### Oneway Anova

###### Summary of Fit

| | |
|---|---|
| Rsquare | 0.009921 |
| Adj Rsquare | -0.00605 |
| Root Mean Square Error | 7.312225 |
| Mean of Response | 18.94275 |
| Observations (or Sum Wgts) | 64 |

###### t Test

1-0

Assuming equal variances

| | | | |
|---|---|---|---|
| Difference | 1.4437 | t Ratio | 0.78822 |
| Std Err Dif | 1.8316 | DF | 62 |
| Upper CL Dif | 5.1051 | Prob > \|t\| | 0.4336 |
| Lower CL Dif | -2.2177 | Prob > t | 0.2168 |
| Confidence | 0.95 | Prob < t | 0.7832 |

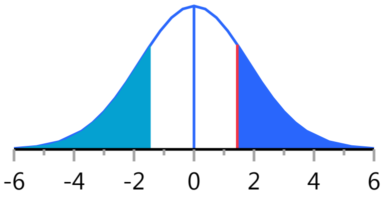

### Fit Group

#### Oneway Analysis of LBP By Weight IBS-subtype=0

##### Oneway Anova

###### Analysis of Variance

| Source | DF | Sum of Squares | Mean Square | F Ratio | Prob > F |
|---|---|---|---|---|---|
| Weight | 1 | 33.2196 | 33.2196 | 0.6213 | 0.4336 |
| Error | 62 | 3315.0552 | 53.4686 | | |
| C. Total | 63 | 3348.2748 | | | |

###### Means for Oneway Anova

| Level | Number | Mean | Std Error | Lower 95% | Upper 95% |
|---|---|---|---|---|---|
| 0 | 34 | 18.2660 | 1.2540 | 15.759 | 20.773 |
| 1 | 30 | 19.7097 | 1.3350 | 17.041 | 22.378 |

Std Error uses a pooled estimate of error variance

##### Means and Std Deviations

| Level | Number | Mean | Std Dev | Std Err Mean | Lower 95% | Upper 95% |
|---|---|---|---|---|---|---|
| 0 | 34 | 18.2660 | 7.55834 | 1.2962 | 15.629 | 20.903 |
| 1 | 30 | 19.7097 | 7.02168 | 1.2820 | 17.088 | 22.332 |

##### t Test

1-0

Assuming unequal variances

| | | | |
|---|---|---|---|
| Difference | 1.4437 | t Ratio | 0.791909 |
| Std Err Dif | 1.8231 | DF | 61.82265 |
| Upper CL Dif | 5.0883 | Prob > \|t\| | 0.4314 |
| Lower CL Dif | -2.2008 | Prob > t | 0.2157 |
| Confidence | 0.95 | Prob < t | 0.7843 |

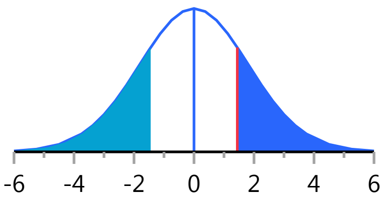


Missing Rows 13

#### Oneway Analysis of Monocytes By Weight IBS-subtype=0

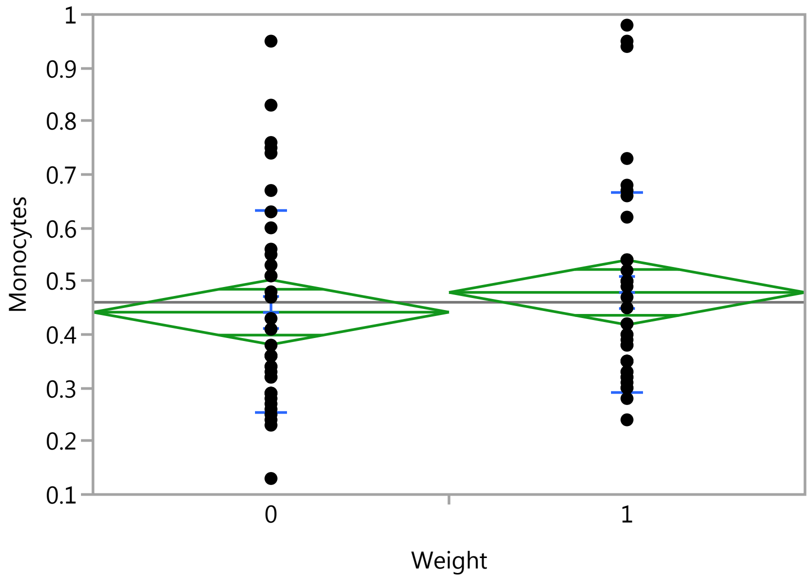

### Fit Group

#### Oneway Analysis of Monocytes By Weight IBS-subtype=0

##### Oneway Anova

###### Summary of Fit

| | |
|---|---|
| Rsquare | 0.009902 |
| Adj Rsquare | -0.00348 |
| Root Mean Square Error | 0.18801 |
| Mean of Response | 0.460395 |
| Observations (or Sum Wgts) | 76 |

###### t Test

1-0

Assuming equal variances

| | | | |
|---|---|---|---|
| Difference | 0.03711 | t Ratio | 0.860264 |
| Std Err Dif | 0.04313 | DF | 74 |
| Upper CL Dif | 0.12305 | Prob > \|t\| | 0.3924 |
| Lower CL Dif | -0.04884 | Prob > t | 0.1962 |
| Confidence | 0.95 | Prob < t | 0.8038 |

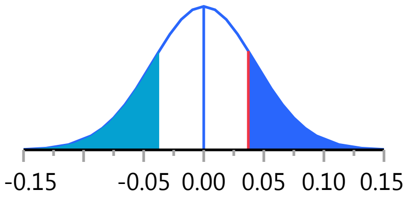


###### Analysis of Variance

| Source | DF | Sum of Squares | Mean Square | F Ratio | Prob > F |
|---|---|---|---|---|---|
| Weight | 1 | 0.0261592 | 0.026159 | 0.7401 | 0.3924 |
| Error | 74 | 2.6157289 | 0.035348 | | |
| C. Total | 75 | 2.6418882 | | | |

###### Means for Oneway Anova

| Level | Number | Mean | Std Error | Lower 95% | Upper 95% |
|---|---|---|---|---|---|
| 0 | 38 | 0.441842 | 0.03050 | 0.38107 | 0.50261 |
| 1 | 38 | 0.478947 | 0.03050 | 0.41818 | 0.53972 |

Std Error uses a pooled estimate of error variance

##### Means and Std Deviations

| Level | Number | Mean | Std Dev | Std Err Mean | Lower 95% | Upper 95% |
|---|---|---|---|---|---|---|
| 0 | 38 | 0.441842 | 0.189221 | 0.03070 | 0.37965 | 0.50404 |
| 1 | 38 | 0.478947 | 0.186791 | 0.03030 | 0.41755 | 0.54034 |

##### t Test

1-0

Assuming unequal variances

| | | | |
|---|---|---|---|
| Difference | 0.03711 | t Ratio | 0.860264 |
| Std Err Dif | 0.04313 | DF | 73.98764 |
| Upper CL Dif | 0.12305 | Prob > \|t\| | 0.3924 |
| Lower CL Dif | -0.04884 | Prob > t | 0.1962 |
| Confidence | 0.95 | Prob < t | 0.8038 |

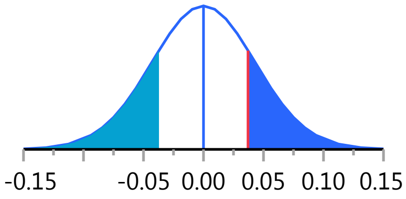


Missing Rows 1

#### Fit Group

##### Oneway Analysis of Monocytes_PCT By Weight IBS-subtype=0

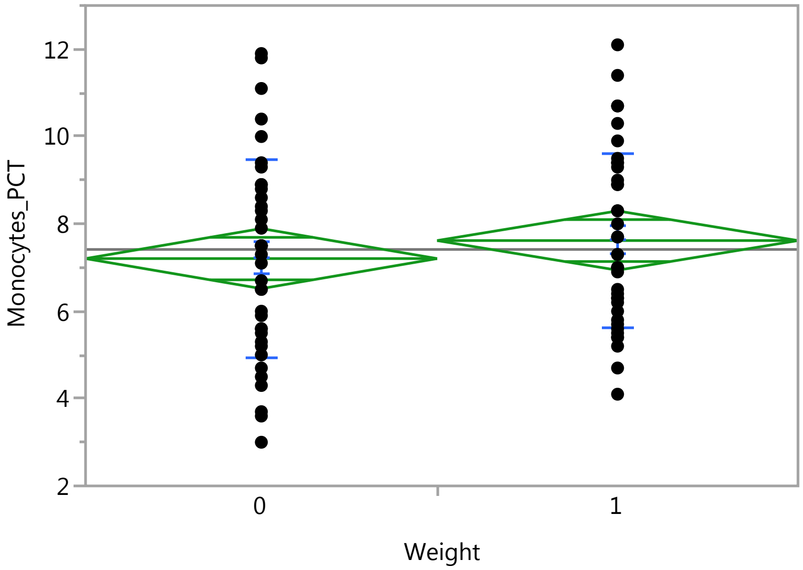


##### Oneway Anova

###### Summary of Fit

| | |
|---|---|
| Rsquare | 0.009558 |
| Adj Rsquare | -0.00365 |
| Root Mean Square Error | 2.128086 |
| Mean of Response | 7.414286 |
| Observations (or Sum Wgts) | 77 |

###### t Test

1-0

Assuming equal variances

| | | | |
|---|---|---|---|
| Difference | 0.4127 | t Ratio | 0.850763 |
| Std Err Dif | 0.4851 | DF | 75 |
| Upper CL Dif | 1.3790 | Prob > \|t\| | 0.3976 |
| Lower CL Dif | -0.5536 | Prob > t | 0.1988 |
| Confidence | 0.95 | Prob < t | 0.8012 |

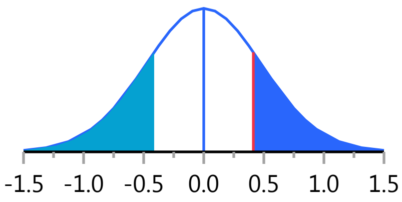


###### Analysis of Variance

| Source | DF | Sum of Squares | Mean Square | F Ratio | Prob > F |
|---|---|---|---|---|---|
| Weight | 1 | 3.27790 | 3.27790 | 0.7238 | 0.3976 |
| Error | 75 | 339.65638 | 4.52875 | | |
| C. Total | 76 | 342.93429 | | | |

###### Means for Oneway Anova

| Level | Number | Mean | Std Error | Lower 95% | Upper 95% |
|---|---|---|---|---|---|
| 0 | 38 | 7.20526 | 0.34522 | 6.5175 | 7.8930 |
| 1 | 39 | 7.61795 | 0.34077 | 6.9391 | 8.2968 |

Std Error uses a pooled estimate of error variance

### Fit Group

#### Oneway Analysis of Monocytes_PCT By Weight IBS-subtype=0

##### Means and Std Deviations

| Level | Number | Mean | Std Dev | Std Err Mean | Lower 95% | Upper 95% |
|---|---|---|---|---|---|---|
| 0 | 38 | 7.20526 | 2.26632 | 0.36764 | 6.4603 | 7.9502 |
| 1 | 39 | 7.61795 | 1.98426 | 0.31774 | 6.9747 | 8.2612 |

##### t Test

1-0

Assuming unequal variances

| | | | |
|---|---|---|---|
| Difference | 0.4127 | t Ratio | 0.849285 |
| Std Err Dif | 0.4859 | DF | 73.16841 |
| Upper CL Dif | 1.3811 | Prob > \|t\| | 0.3985 |
| Lower CL Dif | -0.5557 | Prob > t | 0.1992 |
| Confidence | 0.95 | Prob < t | 0.8008 |

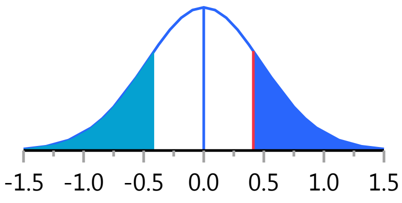


#### Oneway Analysis of MCH By Weight IBS-subtype=0

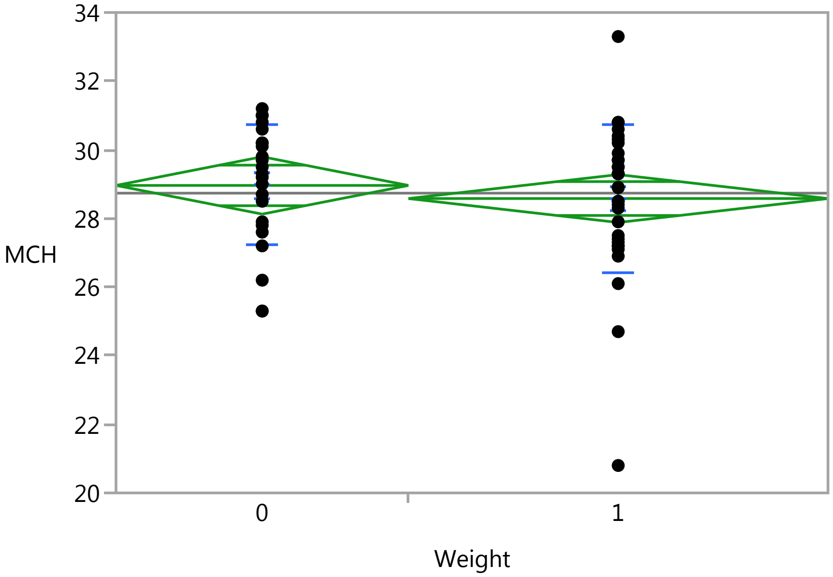


##### Oneway Anova

###### Summary of Fit

| | |
|---|---|
| Rsquare | 0.009083 |
| Adj Rsquare | -0.00927 |
| Root Mean Square Error | 1.999398 |
| Mean of Response | 28.73571 |
| Observations (or Sum Wgts) | 56 |

###### t Test

1-0

### Fit Group

#### Oneway Analysis of MCH By Weight IBS-subtype=0

##### Oneway Anova

###### t Test

Assuming equal variances

| | | | |
|---|---|---|---|
| Difference | -0.3821 | t Ratio | -0.70353 |
| Std Err Dif | 0.5431 | DF | 54 |
| Upper CL Dif | 0.7067 | Prob > \|t\| | 0.4847 |
| Lower CL Dif | -1.4709 | Prob > t | 0.7576 |
| Confidence | 0.95 | Prob < t | 0.2424 |

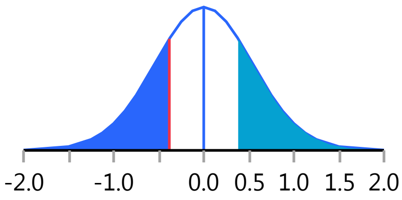


###### Analysis of Variance

| Source | DF | Sum of Squares | Mean Square | F Ratio | Prob > F |
|---|---|---|---|---|---|
| Weight | 1 | 1.97864 | 1.97864 | 0.4950 | 0.4847 |
| Error | 54 | 215.86993 | 3.99759 | | |
| C. Total | 55 | 217.84857 | | | |

###### Means for Oneway Anova

| Level | Number | Mean | Std Error | Lower 95% | Upper 95% |
|---|---|---|---|---|---|
| 0 | 23 | 28.9609 | 0.41690 | 28.125 | 29.797 |
| 1 | 33 | 28.5788 | 0.34805 | 27.881 | 29.277 |

Std Error uses a pooled estimate of error variance

##### Means and Std Deviations

| Level | Number | Mean | Std Dev | Std Err Mean | Lower 95% | Upper 95% |
|---|---|---|---|---|---|---|
| 0 | 23 | 28.9609 | 1.75311 | 0.36555 | 28.203 | 29.719 |
| 1 | 33 | 28.5788 | 2.15243 | 0.37469 | 27.816 | 29.342 |

##### t Test

1-0

Assuming unequal variances

| | | | |
|---|---|---|---|
| Difference | -0.3821 | t Ratio | -0.7299 |
| Std Err Dif | 0.5235 | DF | 52.59715 |
| Upper CL Dif | 0.6681 | Prob > \|t\| | 0.4687 |
| Lower CL Dif | -1.4322 | Prob > t | 0.7657 |
| Confidence | 0.95 | Prob < t | 0.2343 |

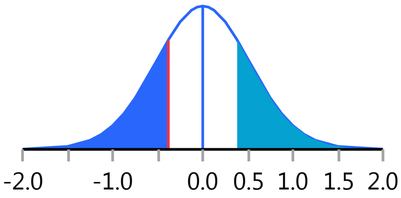


Missing Rows 21

#### Oneway Analysis of IgA By Weight IBS-subtype=0

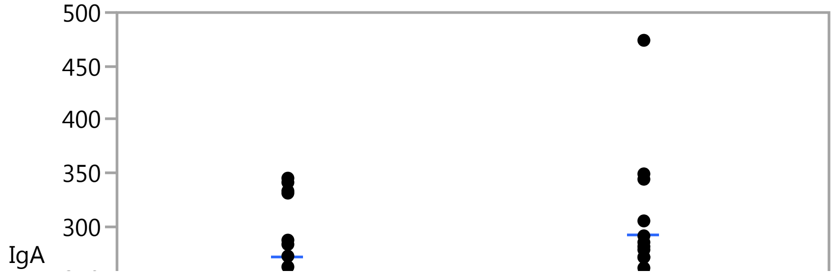

### Fit Group

#### Oneway Analysis of IgA By Weight IBS-subtype=0

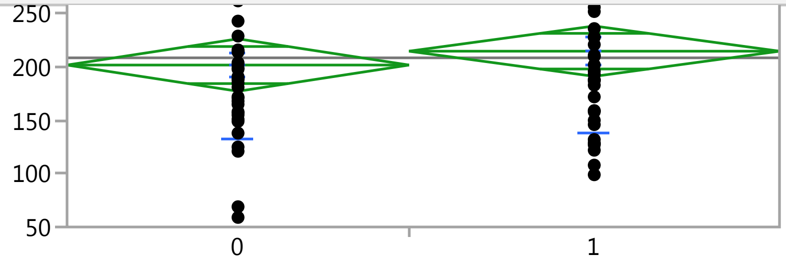


#### Oneway Anova

##### Summary of Fit

| | |
|---|---|
| Rsquare | 0.00784 |
| Adj Rsquare | -0.00575 |
| Root Mean Square Error | 73.98359 |
| Mean of Response | 208.5867 |
| Observations (or Sum Wgts) | 75 |

##### t Test

1-0

Assuming equal variances

| | | | |
|---|---|---|---|
| Difference | 12.987 | t Ratio | 0.759508 |
| Std Err Dif | 17.099 | DF | 73 |
| Upper CL Dif | 47.066 | Prob > \|t\| | 0.4500 |
| Lower CL Dif | -21.092 | Prob > t | 0.2250 |
| Confidence | 0.95 | Prob < t | 0.7750 |

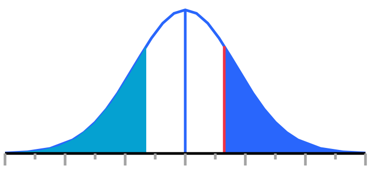


##### Analysis of Variance

| Source | DF | Sum of Squares | Mean Square | F Ratio | Prob > F |
|---|---|---|---|---|---|
| Weight | 1 | 3157.44 | 3157.44 | 0.5769 | 0.4500 |
| Error | 73 | 399570.74 | 5473.57 | | |
| C. Total | 74 | 402728.19 | | | |

##### Means for Oneway Anova

| Level | Number | Mean | Std Error | Lower 95% | Upper 95% |
|---|---|---|---|---|---|
| 0 | 36 | 201.833 | 12.331 | 177.26 | 226.41 |
| 1 | 39 | 214.821 | 11.847 | 191.21 | 238.43 |

Std Error uses a pooled estimate of error variance

#### Means and Std Deviations

| Level | Number | Mean | Std Dev | Std Err Mean | Lower 95% | Upper 95% |
|---|---|---|---|---|---|---|
| 0 | 36 | 201.833 | 69.8388 | 11.640 | 178.20 | 225.46 |
| 1 | 39 | 214.821 | 77.6056 | 12.427 | 189.66 | 239.98 |

#### t Test

1-0

### Fit Group

#### Oneway Analysis of IgA By Weight IBS-subtype=0

##### t Test

Assuming unequal variances

| | | | |
|---|---|---|---|
| Difference | 12.987 | t Ratio | 0.76275 |
| Std Err Dif | 17.027 | DF | 72.95706 |
| Upper CL Dif | 46.922 | Prob > \|t\| | 0.4481 |
| Lower CL Dif | -20.947 | Prob > t | 0.2240 |
| Confidence | 0.95 | Prob < t | 0.7760 |

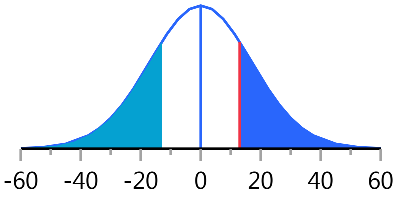


Missing Rows 2

#### Oneway Analysis of Basophils By Weight IBS-subtype=0

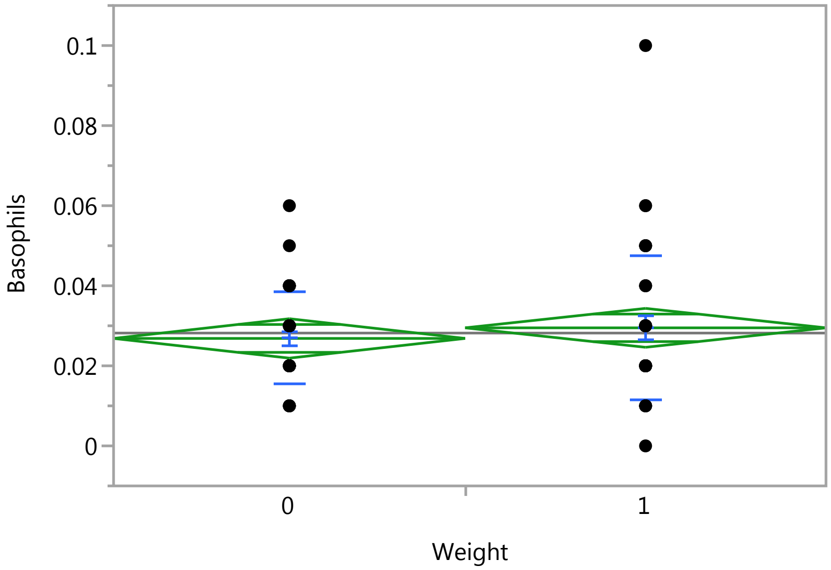


##### Oneway Anova

###### Summary of Fit

| | |
|---|---|
| Rsquare | 0.007675 |
| Adj Rsquare | -0.00556 |
| Root Mean Square Error | 0.015236 |
| Mean of Response | 0.028182 |
| Observations (or Sum Wgts) | 77 |

###### t Test

1-0

Assuming equal variances

| | | | |
|---|---|---|---|
| Difference | 0.00265 | t Ratio | 0.761619 |
| Std Err Dif | 0.00347 | DF | 75 |
| Upper CL Dif | 0.00956 | Prob > \|t\| | 0.4487 |
| Lower CL Dif | -0.00427 | Prob > t | 0.2243 |
| Confidence | 0.95 | Prob < t | 0.7757 |

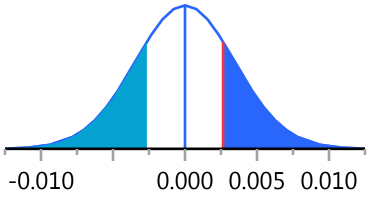

### Fit Group

#### Oneway Analysis of Basophils By Weight IBS-subtype=0

##### Oneway Anova

###### Analysis of Variance

| Source | DF | Sum of Squares | Mean Square | F Ratio | Prob > F |
|---|---|---|---|---|---|
| Weight | 1 | 0.00013466 | 0.000135 | 0.5801 | 0.4487 |
| Error | 75 | 0.01741080 | 0.000232 | | |
| C. Total | 76 | 0.01754545 | | | |

###### Means for Oneway Anova

| Level | Number | Mean | Std Error | Lower 95% | Upper 95% |
|---|---|---|---|---|---|
| 0 | 38 | 0.026842 | 0.00247 | 0.02192 | 0.03177 |
| 1 | 39 | 0.029487 | 0.00244 | 0.02463 | 0.03435 |

Std Error uses a pooled estimate of error variance

##### Means and Std Deviations

| Level | Number | Mean | Std Dev | Std Err Mean | Lower 95% | Upper 95% |
|---|---|---|---|---|---|---|
| 0 | 38 | 0.026842 | 0.011415 | 0.00185 | 0.02309 | 0.03059 |
| 1 | 39 | 0.029487 | 0.018202 | 0.00291 | 0.02359 | 0.03539 |

##### t Test

1-0

Assuming unequal variances

| | | | |
|---|---|---|---|
| Difference | 0.00265 | t Ratio | 0.765996 |
| Std Err Dif | 0.00345 | DF | 64.13574 |
| Upper CL Dif | 0.00954 | Prob > \|t\| | 0.4465 |
| Lower CL Dif | -0.00425 | Prob > t | 0.2232 |
| Confidence | 0.95 | Prob < t | 0.7768 |

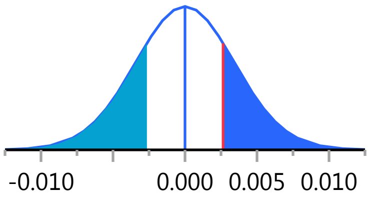


#### Oneway Analysis of Eosinophil_PCT By Weight IBS-subtype=0

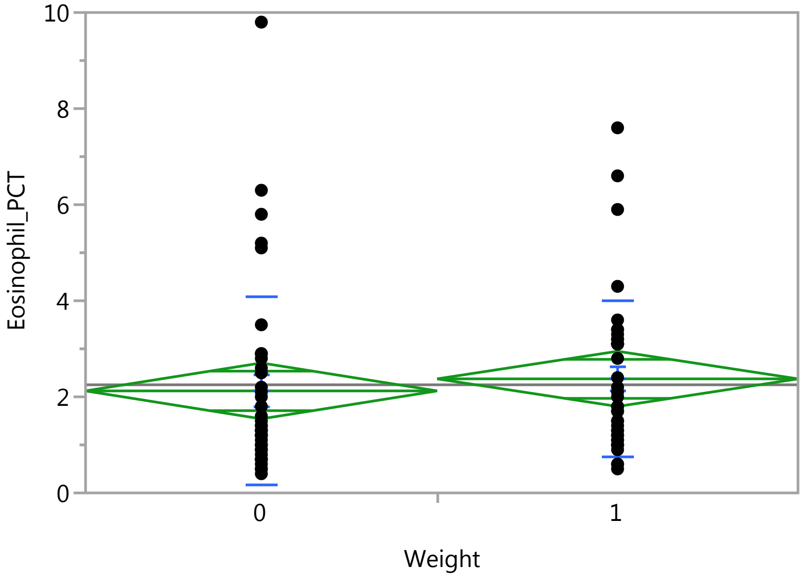

### Fit Group

#### Oneway Analysis of Eosinophil_PCT By Weight IBS-subtype=0

##### Oneway Anova

###### Summary of Fit

| | |
|---|---|
| Rsquare | 0.004948 |
| Adj Rsquare | -0.00832 |
| Root Mean Square Error | 1.800752 |
| Mean of Response | 2.250649 |
| Observations (or Sum Wgts) | 77 |

###### t Test

1-0

Assuming equal variances

| | | | |
|---|---|---|---|
| Difference | 0.2507 | t Ratio | 0.610711 |
| Std Err Dif | 0.4105 | DF | 75 |
| Upper CL Dif | 1.0684 | Prob > \|t\| | 0.5432 |
| Lower CL Dif | -0.5670 | Prob > t | 0.2716 |
| Confidence | 0.95 | Prob < t | 0.7284 |

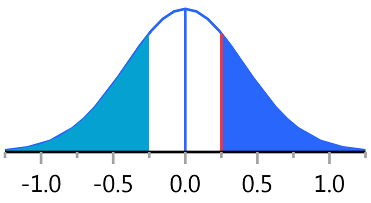


###### Analysis of Variance

| Source | DF | Sum of Squares | Mean Square | F Ratio | Prob > F |
|---|---|---|---|---|---|
| Weight | 1 | 1.20942 | 1.20942 | 0.3730 | 0.5432 |
| Error | 75 | 243.20304 | 3.24271 | | |
| C. Total | 76 | 244.41247 | | | |

###### Means for Oneway Anova

| Level | Number | Mean | Std Error | Lower 95% | Upper 95% |
|---|---|---|---|---|---|
| 0 | 38 | 2.12368 | 0.29212 | 1.5418 | 2.7056 |
| 1 | 39 | 2.37436 | 0.28835 | 1.7999 | 2.9488 |

Std Error uses a pooled estimate of error variance

##### Means and Std Deviations

| Level | Number | Mean | Std Dev | Std Err Mean | Lower 95% | Upper 95% |
|---|---|---|---|---|---|---|
| 0 | 38 | 2.12368 | 1.96956 | 0.31950 | 1.4763 | 2.7711 |
| 1 | 39 | 2.37436 | 1.61957 | 0.25934 | 1.8494 | 2.8994 |

##### t Test

1-0

Assuming unequal variances

| | | | |
|---|---|---|---|
| Difference | 0.2507 | t Ratio | 0.60916 |
| Std Err Dif | 0.4115 | DF | 71.56718 |
| Upper CL Dif | 1.0711 | Prob > \|t\| | 0.5443 |
| Lower CL Dif | -0.5697 | Prob > t | 0.2722 |
| Confidence | 0.95 | Prob < t | 0.7278 |

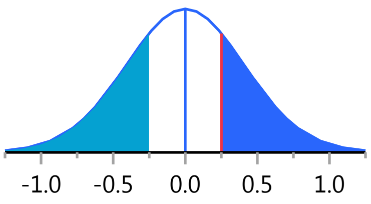

#### Fit Group

##### Oneway Analysis of WBC By Weight IBS-subtype=0

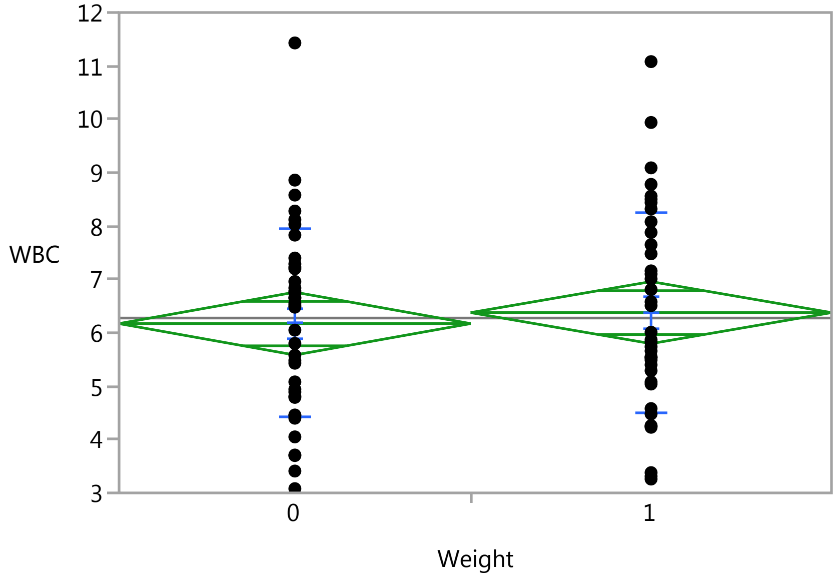


##### Oneway Anova

###### Summary of Fit

| | |
|---|---|
| Rsquare | 0.003266 |
| Adj Rsquare | -0.01002 |
| Root Mean Square Error | 1.82165 |
| Mean of Response | 6.276364 |
| Observations (or Sum Wgts) | 77 |

###### t Test

1-0

Assuming equal variances

| | | | |
|---|---|---|---|
| Difference | 0.2058 | t Ratio | 0.495736 |
| Std Err Dif | 0.4152 | DF | 75 |
| Upper CL Dif | 1.0330 | Prob > \|t\| | 0.6215 |
| Lower CL Dif | -0.6213 | Prob > t | 0.3108 |
| Confidence | 0.95 | Prob < t | 0.6892 |

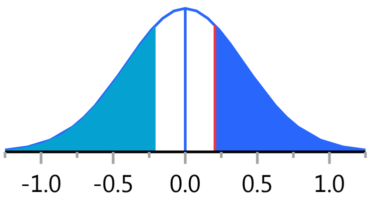


###### Analysis of Variance

| Source | DF | Sum of Squares | Mean Square | F Ratio | Prob > F |
|---|---|---|---|---|---|
| Weight | 1 | 0.81551 | 0.81551 | 0.2458 | 0.6215 |
| Error | 75 | 248.88067 | 3.31841 | | |
| C. Total | 76 | 249.69618 | | | |

###### Means for Oneway Anova

| Level | Number | Mean | Std Error | Lower 95% | Upper 95% |
|---|---|---|---|---|---|
| 0 | 38 | 6.17211 | 0.29551 | 5.5834 | 6.7608 |
| 1 | 39 | 6.37795 | 0.29170 | 5.7969 | 6.9590 |

Std Error uses a pooled estimate of error variance

### Fit Group

#### Oneway Analysis of WBC By Weight IBS-subtype=0

##### Means and Std Deviations

| Level | Number | Mean | Std Dev | Std Err Mean | Lower 95% | Upper 95% |
|---|---|---|---|---|---|---|
| 0 | 38 | 6.17211 | 1.76441 | 0.28622 | 5.5922 | 6.7521 |
| 1 | 39 | 6.37795 | 1.87571 | 0.30035 | 5.7699 | 6.9860 |

##### t Test

1-0

Assuming unequal variances

| | | | |
|---|---|---|---|
| Difference | 0.2058 | t Ratio | 0.496135 |
| Std Err Dif | 0.4149 | DF | 74.90919 |
| Upper CL Dif | 1.0324 | Prob > \|t\| | 0.6213 |
| Lower CL Dif | -0.6207 | Prob > t | 0.3106 |
| Confidence | 0.95 | Prob < t | 0.6894 |

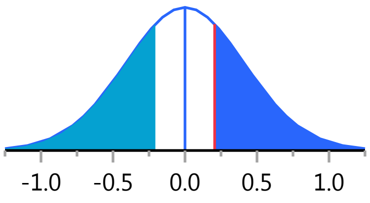


#### Oneway Analysis of Basophil_PCT By Weight IBS-subtype=0

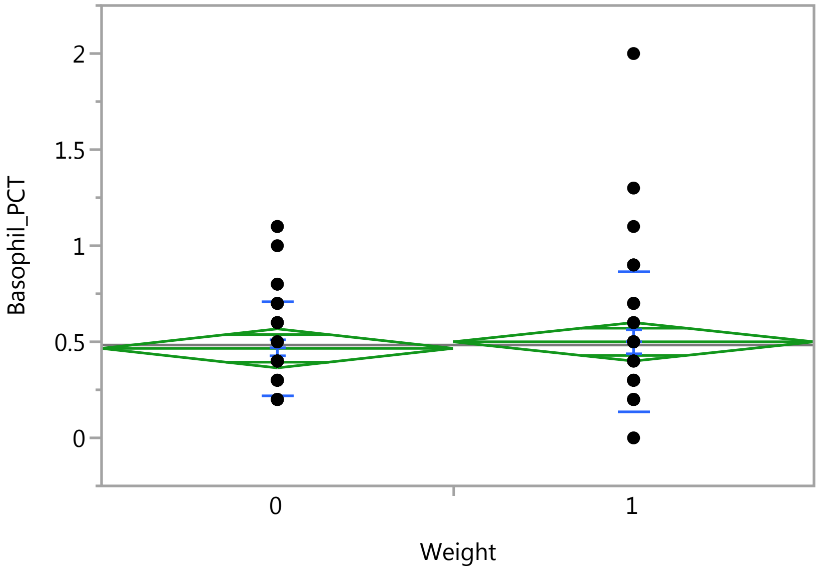


##### Oneway Anova

###### Summary of Fit

| | |
|---|---|
| Rsquare | 0.003033 |
| Adj Rsquare | -0.01026 |
| Root Mean Square Error | 0.31423 |
| Mean of Response | 0.483117 |
| Observations (or Sum Wgts) | 77 |

###### t Test

1-0

### Fit Group

#### Oneway Analysis of Basophil_PCT By Weight IBS-subtype=0

##### Oneway Anova

###### t Test

Assuming equal variances

| | | | |
|---|---|---|---|
| Difference | 0.03421 | t Ratio | 0.47763 |
| Std Err Dif | 0.07163 | DF | 75 |
| Upper CL Dif | 0.17690 | Prob > \|t\| | 0.6343 |
| Lower CL Dif | -0.10848 | Prob > t | 0.3172 |
| Confidence | 0.95 | Prob < t | 0.6828 |

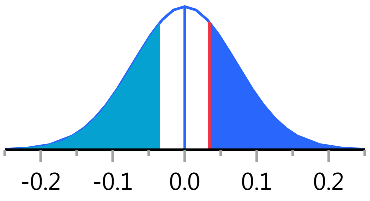


###### Analysis of Variance

| Source | DF | Sum of Squares | Mean Square | F Ratio | Prob > F |
|---|---|---|---|---|---|
| Weight | 1 | 0.0225256 | 0.022526 | 0.2281 | 0.6343 |
| Error | 75 | 7.4055263 | 0.098740 | | |
| C. Total | 76 | 7.4280519 | | | |

###### Means for Oneway Anova

| Level | Number | Mean | Std Error | Lower 95% | Upper 95% |
|---|---|---|---|---|---|
| 0 | 38 | 0.465789 | 0.05097 | 0.36424 | 0.56734 |
| 1 | 39 | 0.500000 | 0.05032 | 0.39976 | 0.60024 |

Std Error uses a pooled estimate of error variance

##### Means and Std Deviations

| Level | Number | Mean | Std Dev | Std Err Mean | Lower 95% | Upper 95% |
|---|---|---|---|---|---|---|
| 0 | 38 | 0.465789 | 0.246353 | 0.03996 | 0.38482 | 0.54676 |
| 1 | 39 | 0.500000 | 0.368496 | 0.05901 | 0.38055 | 0.61945 |

##### t Test

1-0

Assuming unequal variances

| | | | |
|---|---|---|---|
| Difference | 0.03421 | t Ratio | 0.480038 |
| Std Err Dif | 0.07127 | DF | 66.48899 |
| Upper CL Dif | 0.17648 | Prob > \|t\| | 0.6328 |
| Lower CL Dif | -0.10806 | Prob > t | 0.3164 |
| Confidence | 0.95 | Prob < t | 0.6836 |

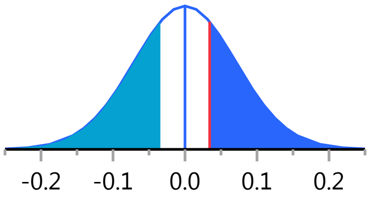


#### Oneway Analysis of LDH By Weight IBS-subtype=0

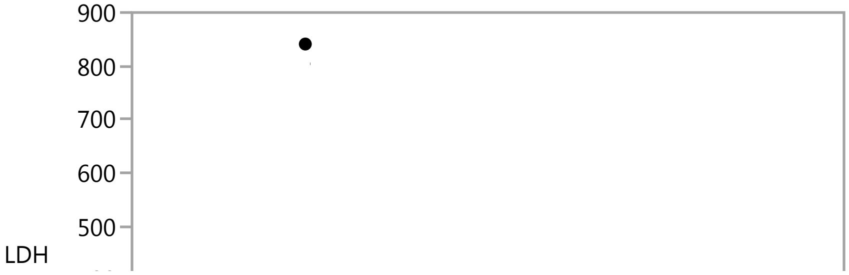

#### Fit Group

##### Oneway Analysis of LDH By Weight IBS-subtype=0

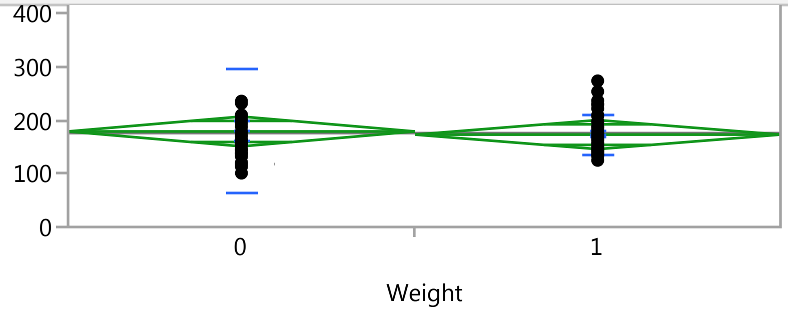


#### Oneway Anova

##### Summary of Fit

| | |
|---|---|
| Rsquare | 0.001237 |
| Adj Rsquare | -0.01226 |
| Root Mean Square Error | 85.36772 |
| Mean of Response | 176.1184 |
| Observations (or Sum Wgts) | 76 |

##### t Test

1-0

Assuming equal variances

| | | | |
|---|---|---|---|
| Difference | -5.931 | t Ratio | -0.30275 |
| Std Err Dif | 19.591 | DF | 74 |
| Upper CL Dif | 33.105 | Prob > \|t\| | 0.7629 |
| Lower CL Dif | -44.968 | Prob > t | 0.6185 |
| Confidence | 0.95 | Prob < t | 0.3815 |

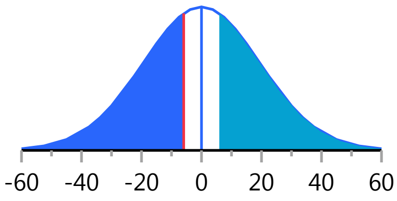


##### Analysis of Variance

| Source | DF | Sum of Squares | Mean Square | F Ratio | Prob > F |
|---|---|---|---|---|---|
| Weight | 1 | 667.98 | 667.98 | 0.0917 | 0.7629 |
| Error | 74 | 539285.95 | 7287.65 | | |
| C. Total | 75 | 539953.93 | | | |

##### Means for Oneway Anova

| Level | Number | Mean | Std Error | Lower 95% | Upper 95% |
|---|---|---|---|---|---|
| 0 | 37 | 179.162 | 14.034 | 151.20 | 207.13 |
| 1 | 39 | 173.231 | 13.670 | 145.99 | 200.47 |

Std Error uses a pooled estimate of error variance

#### Means and Std Deviations

| Level | Number | Mean | Std Dev | Std Err Mean | Lower 95% | Upper 95% |
|---|---|---|---|---|---|---|
| 0 | 37 | 179.162 | 116.401 | 19.136 | 140.35 | 217.97 |
| 1 | 39 | 173.231 | 36.818 | 5.896 | 161.30 | 185.17 |

#### t Test

1-0

### Fit Group

#### Oneway Analysis of LDH By Weight IBS-subtype=0

##### t Test

Assuming unequal variances

| | | | |
|---|---|---|---|
| Difference | -5.931 | t Ratio | -0.29622 |
| Std Err Dif | 20.024 | DF | 42.79299 |
| Upper CL Dif | 34.456 | Prob > \|t\| | 0.7685 |
| Lower CL Dif | -46.319 | Prob > t | 0.6158 |
| Confidence | 0.95 | Prob < t | 0.3842 |

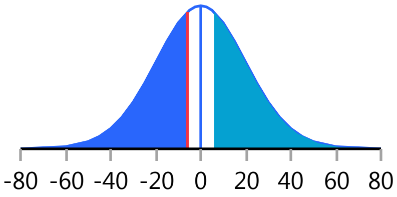


Missing Rows 1

#### Oneway Analysis of ACTH By Weight IBS-subtype=0

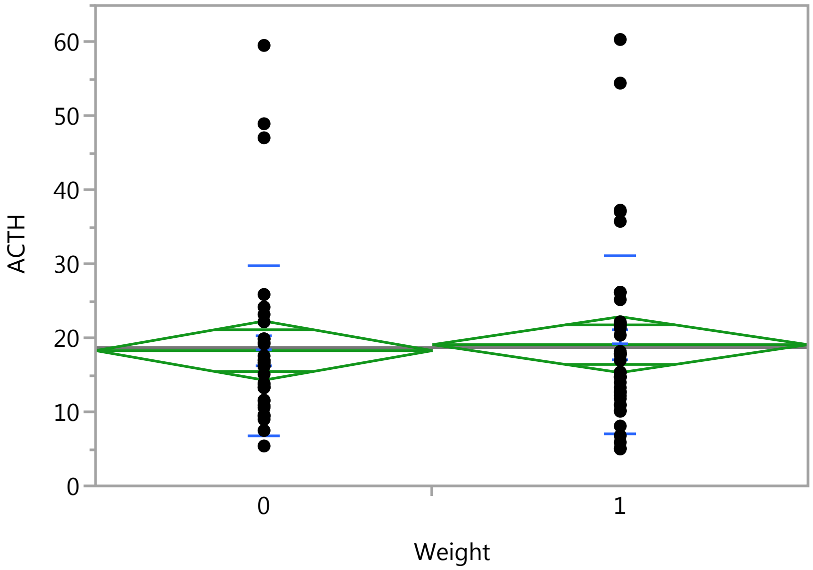


##### Oneway Anova

###### Summary of Fit

| | |
|---|---|
| Rsquare | 0.00118 |
| Adj Rsquare | -0.01269 |
| Root Mean Square Error | 11.85103 |
| Mean of Response | 18.73268 |
| Observations (or Sum Wgts) | 74 |

###### t Test

1-0

Assuming equal variances

| | | | |
|---|---|---|---|
| Difference | 0.8047 | t Ratio | 0.291633 |
| Std Err Dif | 2.7593 | DF | 72 |
| Upper CL Dif | 6.3054 | Prob > \|t\| | 0.7714 |
| Lower CL Dif | -4.6959 | Prob > t | 0.3857 |
| Confidence | 0.95 | Prob < t | 0.6143 |

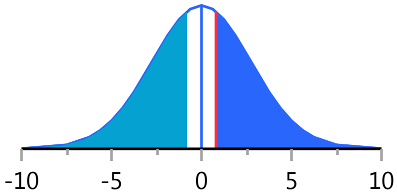

#### Fit Group

##### Oneway Analysis of ACTH By Weight IBS-subtype=0

###### Oneway Anova

###### Analysis of Variance

| Source | DF | Sum of Squares | Mean Square | F Ratio | Prob > F |
|---|---|---|---|---|---|
| Weight | 1 | 11.945 | 11.945 | 0.0850 | 0.7714 |
| Error | 72 | 10112.173 | 140.447 | | |
| C. Total | 73 | 10124.118 | | | |

###### Means for Oneway Anova

| Level | Number | Mean | Std Error | Lower 95% | Upper 95% |
|---|---|---|---|---|---|
| 0 | 35 | 18.3086 | 2.0032 | 14.315 | 22.302 |
| 1 | 39 | 19.1133 | 1.8977 | 15.330 | 22.896 |

Std Error uses a pooled estimate of error variance

###### Means and Std Deviations

| Level | Number | Mean | Std Dev | Std Err Mean | Lower 95% | Upper 95% |
|---|---|---|---|---|---|---|
| 0 | 35 | 18.3086 | 11.5948 | 1.9599 | 14.326 | 22.292 |
| 1 | 39 | 19.1133 | 12.0757 | 1.9337 | 15.199 | 23.028 |

###### t Test

1-0

Assuming unequal variances

| | | | |
|---|---|---|---|
| Difference | 0.8047 | t Ratio | 0.292283 |
| Std Err Dif | 2.7532 | DF | 71.65756 |
| Upper CL Dif | 6.2936 | Prob > \|t\| | 0.7709 |
| Lower CL Dif | -4.6841 | Prob > t | 0.3855 |
| Confidence | 0.95 | Prob < t | 0.6145 |

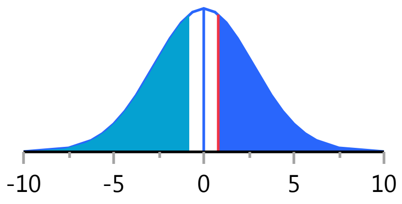


Missing Rows 3

##### Oneway Analysis of RBC By Weight IBS-subtype=0

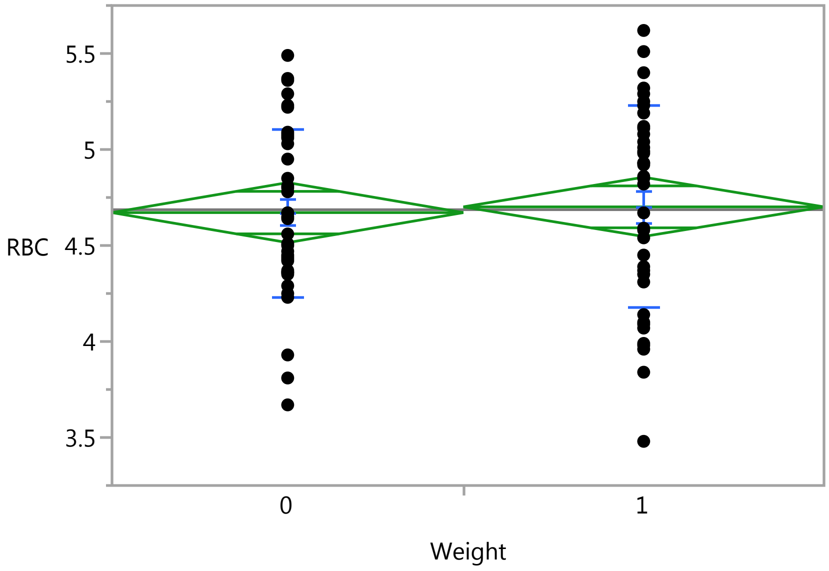

### Fit Group

#### Oneway Analysis of RBC By Weight IBS-subtype=0

##### Oneway Anova

###### Summary of Fit

| | |
|---|---|
| Rsquare | 0.000983 |
| Adj Rsquare | -0.01234 |
| Root Mean Square Error | 0.483929 |
| Mean of Response | 4.686494 |
| Observations (or Sum Wgts) | 77 |

###### t Test

1-0

Assuming equal variances

| | | | |
|---|---|---|---|
| Difference | 0.02997 | t Ratio | 0.271662 |
| Std Err Dif | 0.11031 | DF | 75 |
| Upper CL Dif | 0.24971 | Prob > \|t\| | 0.7866 |
| Lower CL Dif | -0.18978 | Prob > t | 0.3933 |
| Confidence | 0.95 | Prob < t | 0.6067 |

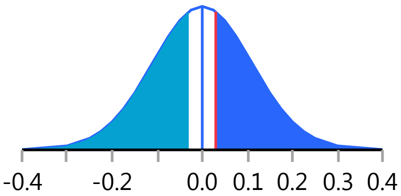


###### Analysis of Variance

| Source | DF | Sum of Squares | Mean Square | F Ratio | Prob > F |
|---|---|---|---|---|---|
| Weight | 1 | 0.017283 | 0.017283 | 0.0738 | 0.7866 |
| Error | 75 | 17.564070 | 0.234188 | | |
| C. Total | 76 | 17.581353 | | | |

###### Means for Oneway Anova

| Level | Number | Mean | Std Error | Lower 95% | Upper 95% |
|---|---|---|---|---|---|
| 0 | 38 | 4.67132 | 0.07850 | 4.5149 | 4.8277 |
| 1 | 39 | 4.70128 | 0.07749 | 4.5469 | 4.8557 |

Std Error uses a pooled estimate of error variance

##### Means and Std Deviations

| Level | Number | Mean | Std Dev | Std Err Mean | Lower 95% | Upper 95% |
|---|---|---|---|---|---|---|
| 0 | 38 | 4.67132 | 0.438056 | 0.07106 | 4.5273 | 4.8153 |
| 1 | 39 | 4.70128 | 0.524756 | 0.08403 | 4.5312 | 4.8714 |

##### t Test

1-0

Assuming unequal variances

| | | | |
|---|---|---|---|
| Difference | 0.02997 | t Ratio | 0.272302 |
| Std Err Dif | 0.11005 | DF | 73.29042 |
| Upper CL Dif | 0.24928 | Prob > \|t\| | 0.7862 |
| Lower CL Dif | -0.18934 | Prob > t | 0.3931 |
| Confidence | 0.95 | Prob < t | 0.6069 |

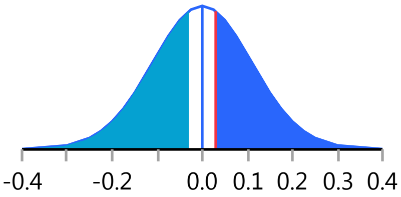

#### Fit Group

##### Oneway Analysis of Neutrophils By Weight IBS-subtype=0

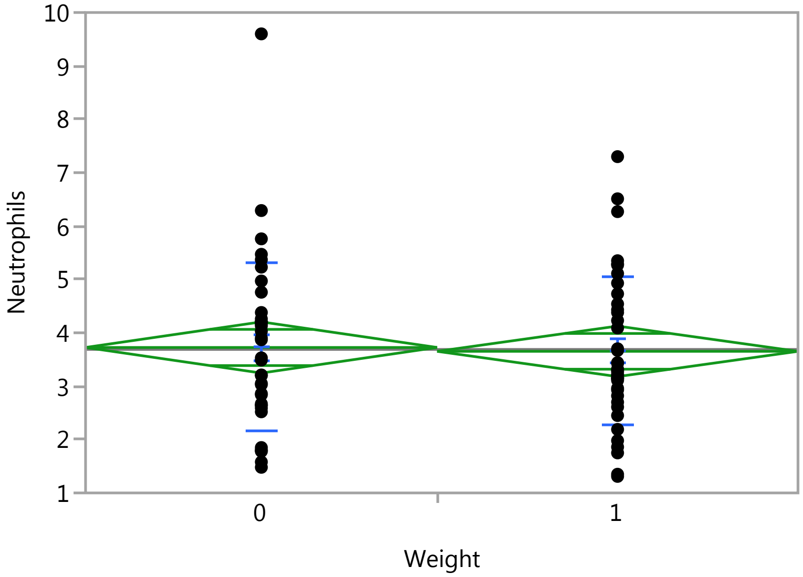


#### Oneway Anova

##### Summary of Fit

| | |
|---|---|
| Rsquare | 0.000611 |
| Adj Rsquare | -0.01271 |
| Root Mean Square Error | 1.483869 |
| Mean of Response | 3.688312 |
| Observations (or Sum Wgts) | 77 |

##### t Test

1-0

Assuming equal variances

| | | | |
|---|---|---|---|
| Difference | -0.07244 | t Ratio | -0.21416 |
| Std Err Dif | 0.33823 | DF | 75 |
| Upper CL Dif | 0.60136 | Prob > \|t\| | 0.8310 |
| Lower CL Dif | -0.74623 | Prob > t | 0.5845 |
| Confidence | 0.95 | Prob < t | 0.4155 |

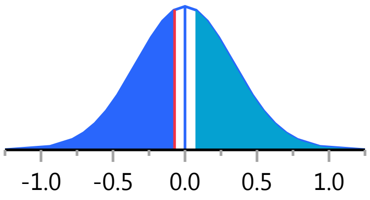


##### Analysis of Variance

| Source | DF | Sum of Squares | Mean Square | F Ratio | Prob > F |
|---|---|---|---|---|---|
| Weight | 1 | 0.10099 | 0.10099 | 0.0459 | 0.8310 |
| Error | 75 | 165.14009 | 2.20187 | | |
| C. Total | 76 | 165.24108 | | | |

##### Means for Oneway Anova

| Level | Number | Mean | Std Error | Lower 95% | Upper 95% |
|---|---|---|---|---|---|
| 0 | 38 | 3.72500 | 0.24072 | 3.2455 | 4.2045 |
| 1 | 39 | 3.65256 | 0.23761 | 3.1792 | 4.1259 |

Std Error uses a pooled estimate of error variance

#### Fit Group

##### Oneway Analysis of Neutrophils By Weight IBS-subtype=0

###### Means and Std Deviations

| Level | Number | Mean | Std Dev | Std Err Mean | Lower 95% | Upper 95% |
|---|---|---|---|---|---|---|
| 0 | 38 | 3.72500 | 1.57396 | 0.25533 | 3.2077 | 4.2423 |
| 1 | 39 | 3.65256 | 1.39055 | 0.22267 | 3.2018 | 4.1033 |

###### t Test

1-0

Assuming unequal variances

| | | | |
|---|---|---|---|
| Difference | -0.07244 | t Ratio | -0.21381 |
| Std Err Dif | 0.33878 | DF | 73.36296 |
| Upper CL Dif | 0.60270 | Prob > \|t\| | 0.8313 |
| Lower CL Dif | -0.74757 | Prob > t | 0.5844 |
| Confidence | 0.95 | Prob < t | 0.4156 |

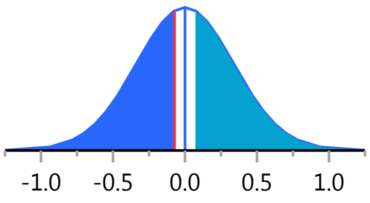


##### Oneway Analysis of HCT By Weight IBS-subtype=0

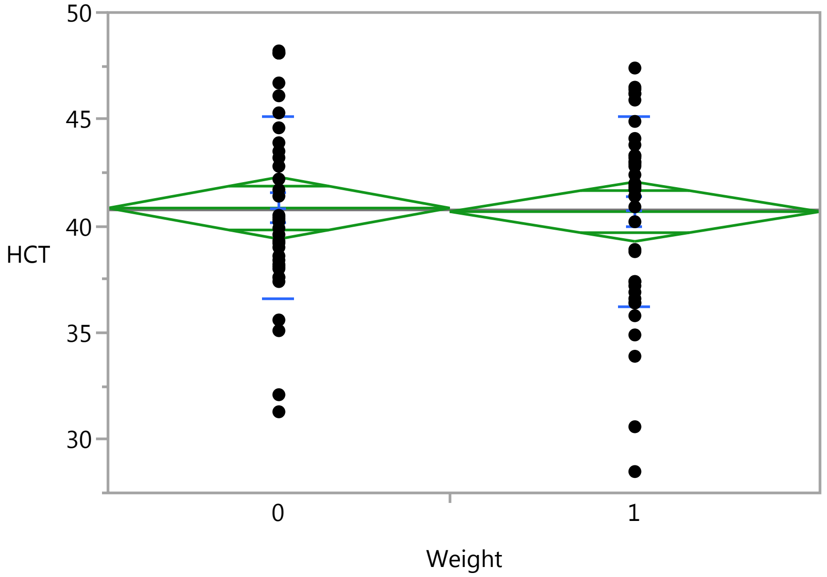


###### Oneway Anova

###### Summary of Fit

| | |
|---|---|
| Rsquare | 0.000376 |
| Adj Rsquare | -0.01332 |
| Root Mean Square Error | 4.368947 |
| Mean of Response | 40.75467 |
| Observations (or Sum Wgts) | 75 |

###### t Test

1-0

### Fit Group

#### Oneway Analysis of HCT By Weight IBS-subtype=0

##### Oneway Anova

###### t Test

Assuming equal variances

| | | | |
|---|---|---|---|
| Difference | -0.1673 | t Ratio | -0.16569 |
| Std Err Dif | 1.0098 | DF | 73 |
| Upper CL Dif | 1.8452 | Prob > \|t\| | 0.8689 |
| Lower CL Dif | -2.1798 | Prob > t | 0.5656 |
| Confidence | 0.95 | Prob < t | 0.4344 |

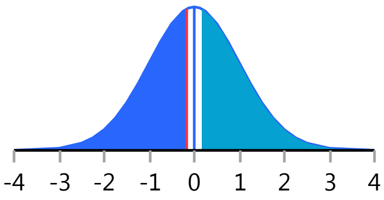


###### Analysis of Variance

| Source | DF | Sum of Squares | Mean Square | F Ratio | Prob > F |
|---|---|---|---|---|---|
| Weight | 1 | 0.5240 | 0.5240 | 0.0275 | 0.8689 |
| Error | 73 | 1393.4019 | 19.0877 | | |
| C. Total | 74 | 1393.9259 | | | |

###### Means for Oneway Anova

| Level | Number | Mean | Std Error | Lower 95% | Upper 95% |
|---|---|---|---|---|---|
| 0 | 36 | 40.8417 | 0.72816 | 39.390 | 42.293 |
| 1 | 39 | 40.6744 | 0.69959 | 39.280 | 42.069 |

Std Error uses a pooled estimate of error variance

##### Means and Std Deviations

| Level | Number | Mean | Std Dev | Std Err Mean | Lower 95% | Upper 95% |
|---|---|---|---|---|---|---|
| 0 | 36 | 40.8417 | 4.28655 | 0.71442 | 39.391 | 42.292 |
| 1 | 39 | 40.6744 | 4.44349 | 0.71153 | 39.234 | 42.115 |

##### t Test

1-0

Assuming unequal variances

| | | | |
|---|---|---|---|
| Difference | -0.1673 | t Ratio | -0.16593 |
| Std Err Dif | 1.0083 | DF | 72.85118 |
| Upper CL Dif | 1.8423 | Prob > \|t\| | 0.8687 |
| Lower CL Dif | -2.1769 | Prob > t | 0.5657 |
| Confidence | 0.95 | Prob < t | 0.4343 |

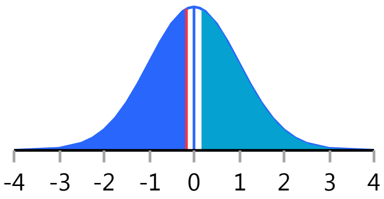


Missing Rows 2

#### Fit Group

##### Oneway Analysis of SerumCortisol By Weight IBS-subtype=1

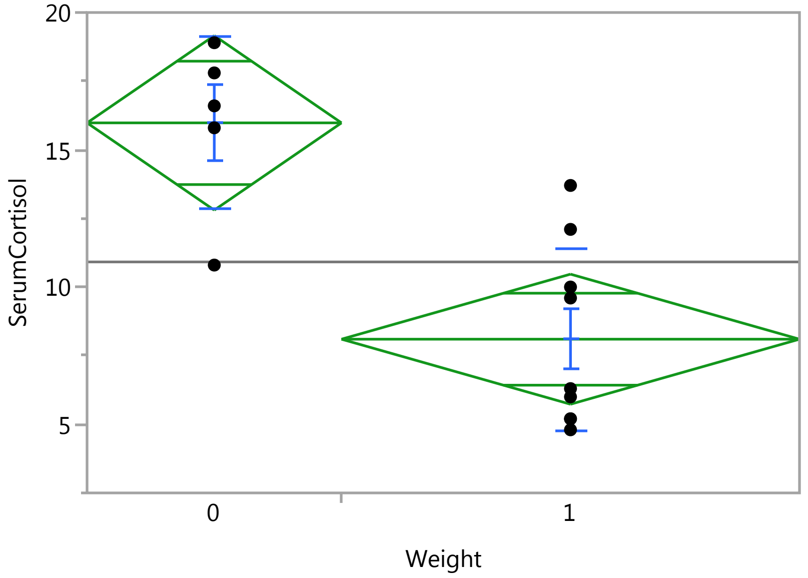


##### Oneway Anova

###### Summary of Fit

| | |
|---|---|
| Rsquare | 0.609885 |
| Adj Rsquare | 0.577375 |
| Root Mean Square Error | 3.261748 |
| Mean of Response | 10.91429 |
| Observations (or Sum Wgts) | 14 |

###### t Test

1-0

Assuming equal variances

| | | | |
|---|---|---|---|
| Difference | -7.880 | t Ratio | -4.3313 |
| Std Err Dif | 1.819 | DF | 12 |
| Upper CL Dif | -3.916 | Prob > \|t\| | 0.0010* |
| Lower CL Dif | -11.844 | Prob > t | 0.9995 |
| Confidence | 0.95 | Prob < t | 0.0005* |

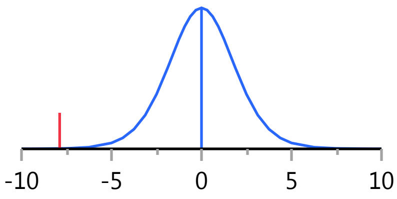


###### Analysis of Variance

| Source | DF | Sum of Squares | Mean Square | F Ratio | Prob > F |
|---|---|---|---|---|---|
| Weight | 1 | 199.58914 | 199.589 | 18.7601 | 0.0010* |
| Error | 12 | 127.66800 | 10.639 | | |
| C. Total | 13 | 327.25714 | | | |

###### Means for Oneway Anova

| Level | Number | Mean | Std Error | Lower 95% | Upper 95% |
|---|---|---|---|---|---|
| 0 | 5 | 15.9800 | 1.4587 | 12.802 | 19.158 |
| 1 | 9 | 8.1000 | 1.0872 | 5.731 | 10.469 |

Std Error uses a pooled estimate of error variance

### Fit Group

#### Oneway Analysis of SerumCortisol By Weight IBS-subtype=1

##### Means and Std Deviations

| Level | Number | Mean | Std Dev | Std Err Mean | Lower 95% | Upper 95% |
|---|---|---|---|---|---|---|
| 0 | 5 | 15.9800 | 3.12602 | 1.3980 | 12.099 | 19.861 |
| 1 | 9 | 8.1000 | 3.32754 | 1.1092 | 5.542 | 10.658 |

##### t Test

1-0

Assuming unequal variances

| | | | |
|---|---|---|---|
| Difference | -7.880 | t Ratio | -4.41564 |
| Std Err Dif | 1.785 | DF | 8.864623 |
| Upper CL Dif | -3.834 | Prob > \|t\| | 0.0017* |
| Lower CL Dif | -11.926 | Prob > t | 0.9991 |
| Confidence | 0.95 | Prob < t | 0.0009* |

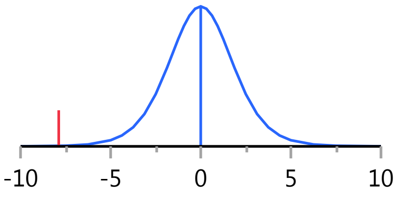


#### Oneway Analysis of MPV By Weight IBS-subtype=1

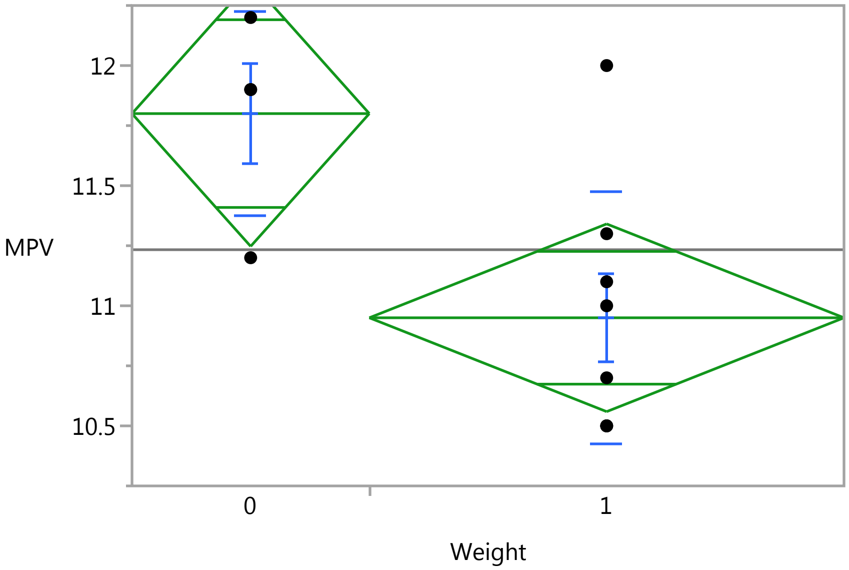


##### Oneway Anova

###### Summary of Fit

| | |
|---|---|
| Rsquare | 0.43921 |
| Adj Rsquare | 0.383131 |
| Root Mean Square Error | 0.495984 |
| Mean of Response | 11.23333 |
| Observations (or Sum Wgts) | 12 |

###### t Test

1-0

### Fit Group

#### Oneway Analysis of MPV By Weight IBS-subtype=1

##### Oneway Anova

###### t Test

Assuming equal variances

| | | | |
|---|---|---|---|
| Difference | -0.8500 | t Ratio | -2.79857 |
| Std Err Dif | 0.3037 | DF | 10 |
| Upper CL Dif | -0.1733 | Prob > \|t\| | 0.0188* |
| Lower CL Dif | -1.5267 | Prob > t | 0.9906 |
| Confidence | 0.95 | Prob < t | 0.0094* |

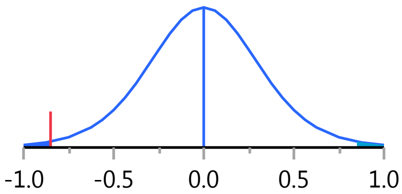


###### Analysis of Variance

| Source | DF | Sum of Squares | Mean Square | F Ratio | Prob > F |
|---|---|---|---|---|---|
| Weight | 1 | 1.9266667 | 1.92667 | 7.8320 | 0.0188* |
| Error | 10 | 2.4600000 | 0.24600 | | |
| C. Total | 11 | 4.3866667 | | | |

###### Means for Oneway Anova

| Level | Number | Mean | Std Error | Lower 95% | Upper 95% |
|---|---|---|---|---|---|
| 0 | 4 | 11.8000 | 0.24799 | 11.247 | 12.353 |
| 1 | 8 | 10.9500 | 0.17536 | 10.559 | 11.341 |

Std Error uses a pooled estimate of error variance

##### Means and Std Deviations

| Level | Number | Mean | Std Dev | Std Err Mean | Lower 95% | Upper 95% |
|---|---|---|---|---|---|---|
| 0 | 4 | 11.8000 | 0.424264 | 0.21213 | 11.125 | 12.475 |
| 1 | 8 | 10.9500 | 0.523723 | 0.18516 | 10.512 | 11.388 |

##### t Test

1-0

Assuming unequal variances

| | | | |
|---|---|---|---|
| Difference | -0.8500 | t Ratio | -3.01871 |
| Std Err Dif | 0.2816 | DF | 7.457588 |
| Upper CL Dif | -0.1924 | Prob > \|t\| | 0.0180* |
| Lower CL Dif | -1.5076 | Prob > t | 0.9910 |
| Confidence | 0.95 | Prob < t | 0.0090* |

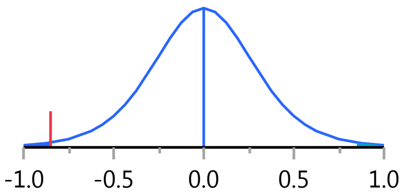


Missing Rows 2

#### Fit Group

##### Oneway Analysis of Eosinophil_PCT By Weight IBS-subtype=1

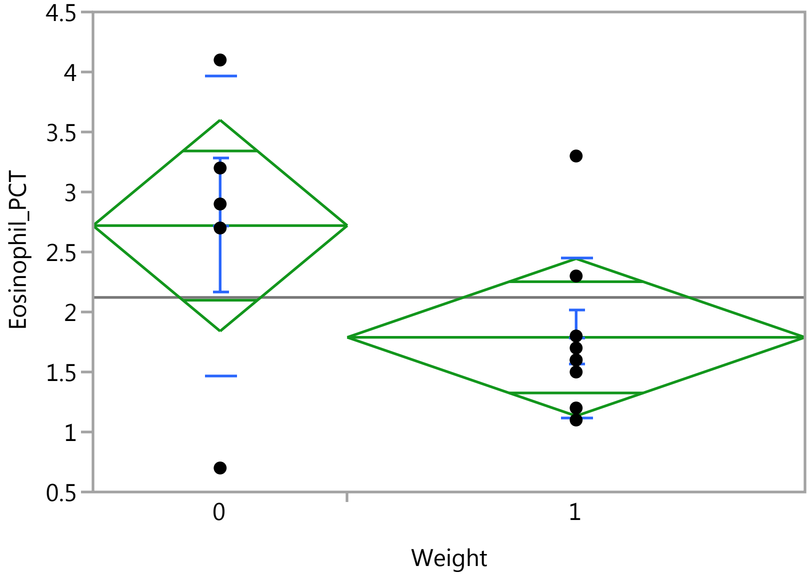


#### Oneway Anova

##### Summary of Fit

| | |
|---|---|
| Rsquare | 0.221807 |
| Adj Rsquare | 0.156957 |
| Root Mean Square Error | 0.90263 |
| Mean of Response | 2.121429 |
| Observations (or Sum Wgts) | 14 |

##### t Test

1-0

Assuming equal variances

| | | | |
|---|---|---|---|
| Difference | -0.9311 | t Ratio | -1.84941 |
| Std Err Dif | 0.5035 | DF | 12 |
| Upper CL Dif | 0.1658 | Prob > \|t\| | 0.0892 |
| Lower CL Dif | -2.0281 | Prob > t | 0.9554 |
| Confidence | 0.95 | Prob < t | 0.0446* |

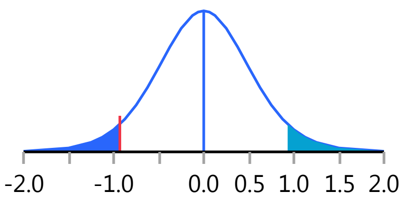


##### Analysis of Variance

| Source | DF | Sum of Squares | Mean Square | F Ratio | Prob > F |
|---|---|---|---|---|---|
| Weight | 1 | 2.786683 | 2.78668 | 3.4203 | 0.0892 |
| Error | 12 | 9.776889 | 0.81474 | | |
| C. Total | 13 | 12.563571 | | | |

##### Means for Oneway Anova

| Level | Number | Mean | Std Error | Lower 95% | Upper 95% |
|---|---|---|---|---|---|
| 0 | 5 | 2.72000 | 0.40367 | 1.8405 | 3.5995 |
| 1 | 9 | 1.78889 | 0.30088 | 1.1333 | 2.4444 |

Std Error uses a pooled estimate of error variance

### Fit Group

#### Oneway Analysis of Eosinophil_PCT By Weight IBS-subtype=1

##### Means and Std Deviations

| Level | Number | Mean | Std Dev | Std Err Mean | Lower 95% | Upper 95% |
|---|---|---|---|---|---|---|
| 0 | 5 | 2.72000 | 1.24980 | 0.55893 | 1.1682 | 4.2718 |
| 1 | 9 | 1.78889 | 0.66416 | 0.22139 | 1.2784 | 2.2994 |

##### t Test

1-0

Assuming unequal variances

| | | | |
|---|---|---|---|
| Difference | -0.9311 | t Ratio | -1.54882 |
| Std Err Dif | 0.6012 | DF | 5.288489 |
| Upper CL Dif | 0.5892 | Prob > \|t\| | 0.1789 |
| Lower CL Dif | -2.4515 | Prob > t | 0.9105 |
| Confidence | 0.95 | Prob < t | 0.0895 |

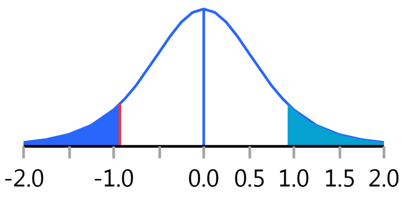


#### Oneway Analysis of PlateletCount By Weight IBS-subtype=1

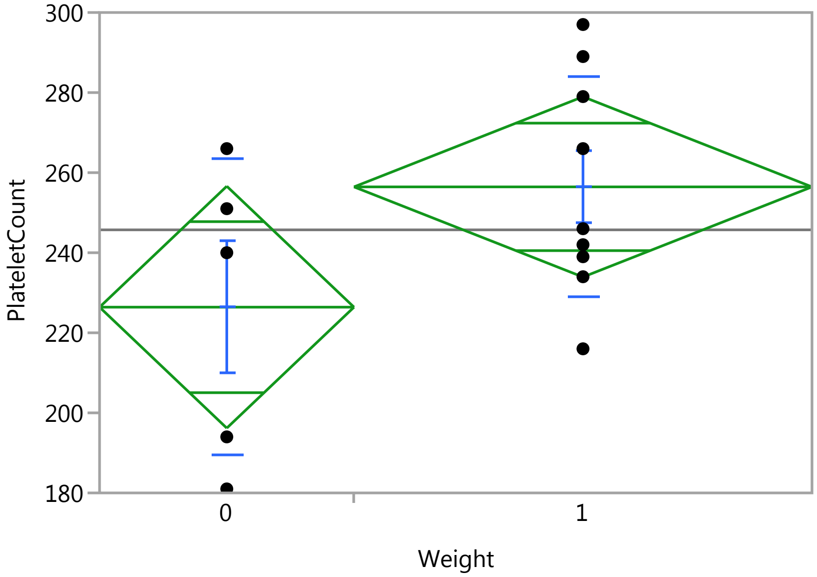


##### Oneway Anova

###### Summary of Fit

| | |
|---|---|
| Rsquare | 0.200974 |
| Adj Rsquare | 0.134389 |
| Root Mean Square Error | 31.0046 |
| Mean of Response | 245.7143 |
| Observations (or Sum Wgts) | 14 |

###### t Test

1-0

### Fit Group

#### Oneway Analysis of PlateletCount By Weight IBS-subtype=1

##### Oneway Anova

###### t Test

Assuming equal variances

| | | | |
|---|---|---|---|
| Difference | 30.044 | t Ratio | 1.737322 |
| Std Err Dif | 17.294 | DF | 12 |
| Upper CL Dif | 67.724 | Prob > \|t\| | 0.1079 |
| Lower CL Dif | -7.635 | Prob > t | 0.0540 |
| Confidence | 0.95 | Prob < t | 0.9460 |

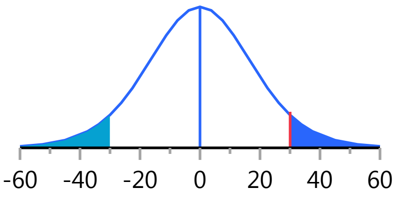


###### Analysis of Variance

| Source | DF | Sum of Squares | Mean Square | F Ratio | Prob > F |
|---|---|---|---|---|---|
| Weight | 1 | 2901.435 | 2901.43 | 3.0183 | 0.1079 |
| Error | 12 | 11535.422 | 961.29 | | |
| C. Total | 13 | 14436.857 | | | |

###### Means for Oneway Anova

| Level | Number | Mean | Std Error | Lower 95% | Upper 95% |
|---|---|---|---|---|---|
| 0 | 5 | 226.400 | 13.866 | 196.19 | 256.61 |
| 1 | 9 | 256.444 | 10.335 | 233.93 | 278.96 |

Std Error uses a pooled estimate of error variance

##### Means and Std Deviations

| Level | Number | Mean | Std Dev | Std Err Mean | Lower 95% | Upper 95% |
|---|---|---|---|---|---|---|
| 0 | 5 | 226.400 | 36.9770 | 16.537 | 180.49 | 272.31 |
| 1 | 9 | 256.444 | 27.5368 | 9.179 | 235.28 | 277.61 |

##### t Test

1-0

Assuming unequal variances

| | | | |
|---|---|---|---|
| Difference | 30.044 | t Ratio | 1.588535 |
| Std Err Dif | 18.913 | DF | 6.534365 |
| Upper CL Dif | 75.422 | Prob > \|t\| | 0.1592 |
| Lower CL Dif | -15.333 | Prob > t | 0.0796 |
| Confidence | 0.95 | Prob < t | 0.9204 |

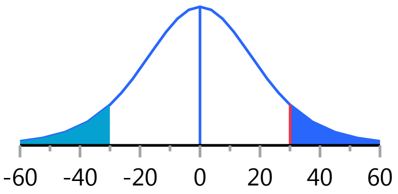


#### Oneway Analysis of sCD14 By Weight IBS-subtype=1

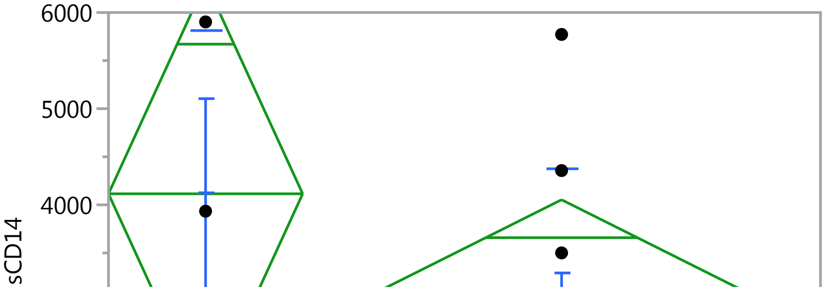

### Fit Group

#### Oneway Analysis of sCD14 By Weight IBS-subtype=1

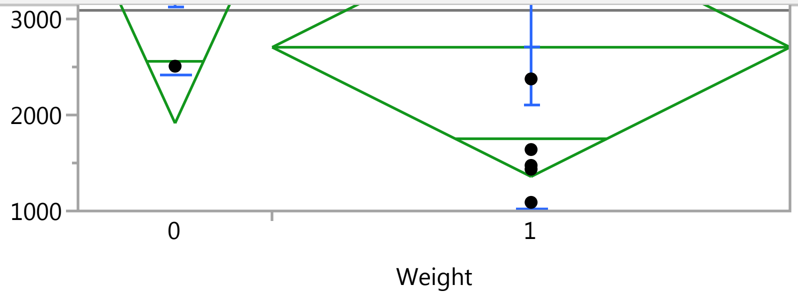


#### Oneway Anova

##### Summary of Fit

| | |
|---|---|
| Rsquare | 0.145008 |
| Adj Rsquare | 0.050009 |
| Root Mean Square Error | 1684.934 |
| Mean of Response | 3089.973 |
| Observations (or Sum Wgts) | 11 |

##### t Test

1-0

Assuming equal variances

| | | | |
|---|---|---|---|
| Difference | -1409.3 | t Ratio | -1.23548 |
| Std Err Dif | 1140.7 | DF | 9 |
| Upper CL Dif | 1171.1 | Prob > \|t\| | 0.2479 |
| Lower CL Dif | -3989.8 | Prob > t | 0.8760 |
| Confidence | 0.95 | Prob < t | 0.1240 |

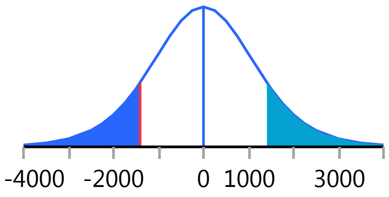


##### Analysis of Variance

| Source | DF | Sum of Squares | Mean Square | F Ratio | Prob > F |
|---|---|---|---|---|---|
| Weight | 1 | 4333495 | 4333495 | 1.5264 | 0.2479 |
| Error | 9 | 25551032 | 2839004 | | |
| C. Total | 10 | 29884527 | | | |

##### Means for Oneway Anova

| Level | Number | Mean | Std Error | Lower 95% | Upper 95% |
|---|---|---|---|---|---|
| 0 | 3 | 4114.93 | 972.80 | 1914.3 | 6315.6 |
| 1 | 8 | 2705.61 | 595.71 | 1358.0 | 4053.2 |

Std Error uses a pooled estimate of error variance

#### Means and Std Deviations

| Level | Number | Mean | Std Dev | Std Err Mean | Lower 95% | Upper 95% |
|---|---|---|---|---|---|---|
| 0 | 3 | 4114.93 | 1704.26 | 983.96 | -119 | 8348.6 |
| 1 | 8 | 2705.61 | 1679.37 | 593.75 | 1302 | 4109.6 |

#### t Test

1-0

#### Fit Group

##### Oneway Analysis of sCD14 By Weight IBS-subtype=1

###### t Test

Assuming unequal variances

| | | | |
|---|---|---|---|
| Difference | -1409.3 | t Ratio | -1.22633 |
| Std Err Dif | 1149.2 | DF | 3.585844 |
| Upper CL Dif | 1932.2 | Prob > \|t\| | 0.2945 |
| Lower CL Dif | -4750.8 | Prob > t | 0.8528 |
| Confidence | 0.95 | Prob < t | 0.1472 |

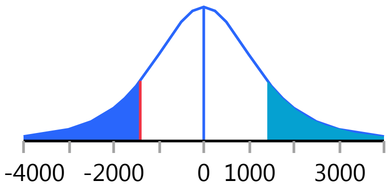


Missing Rows 3

##### Oneway Analysis of Eosinophils By Weight IBS-subtype=1

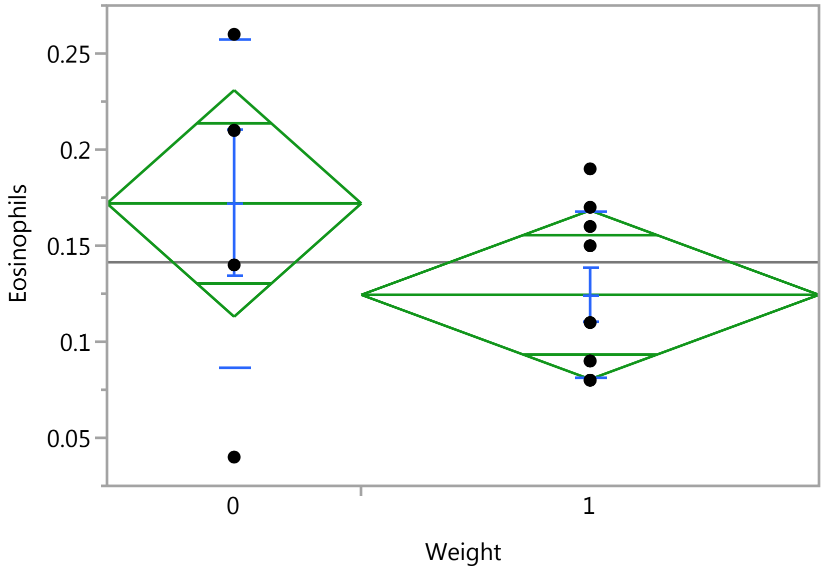


###### Oneway Anova

###### Summary of Fit

| | |
|---|---|
| Rsquare | 0.142056 |
| Adj Rsquare | 0.070561 |
| Root Mean Square Error | 0.060486 |
| Mean of Response | 0.141429 |
| Observations (or Sum Wgts) | 14 |

###### t Test

1-0

Assuming equal variances

| | | | |
|---|---|---|---|
| Difference | -0.04756 | t Ratio | -1.40958 |
| Std Err Dif | 0.03374 | DF | 12 |
| Upper CL Dif | 0.02595 | Prob > \|t\| | 0.1840 |
| Lower CL Dif | -0.12106 | Prob > t | 0.9080 |
| Confidence | 0.95 | Prob < t | 0.0920 |

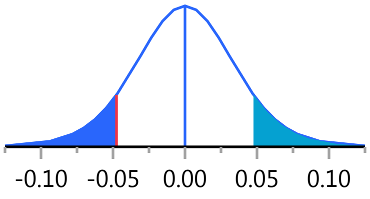

### Fit Group

#### Oneway Analysis of Eosinophils By Weight IBS-subtype=1

##### Oneway Anova

###### Analysis of Variance

| Source | DF | Sum of Squares | Mean Square | F Ratio | Prob > F |
|---|---|---|---|---|---|
| Weight | 1 | 0.00726921 | 0.007269 | 1.9869 | 0.1840 |
| Error | 12 | 0.04390222 | 0.003659 | | |
| C. Total | 13 | 0.05117143 | | | |

###### Means for Oneway Anova

| Level | Number | Mean | Std Error | Lower 95% | Upper 95% |
|---|---|---|---|---|---|
| 0 | 5 | 0.172000 | 0.02705 | 0.11306 | 0.23094 |
| 1 | 9 | 0.124444 | 0.02016 | 0.08052 | 0.16837 |

Std Error uses a pooled estimate of error variance

##### Means and Std Deviations

| Level | Number | Mean | Std Dev | Std Err Mean | Lower 95% | Upper 95% |
|---|---|---|---|---|---|---|
| 0 | 5 | 0.172000 | 0.085264 | 0.03813 | 0.06613 | 0.27787 |
| 1 | 9 | 0.124444 | 0.043044 | 0.01435 | 0.09136 | 0.15753 |

##### t Test

1-0

Assuming unequal variances

| | | | |
|---|---|---|---|
| Difference | -0.04756 | t Ratio | -1.16725 |
| Std Err Dif | 0.04074 | DF | 5.161132 |
| Upper CL Dif | 0.05620 | Prob > \|t\| | 0.2942 |
| Lower CL Dif | -0.15131 | Prob > t | 0.8529 |
| Confidence | 0.95 | Prob < t | 0.1471 |

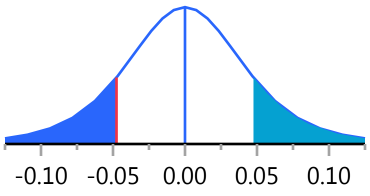


#### Oneway Analysis of ACTH By Weight IBS-subtype=1

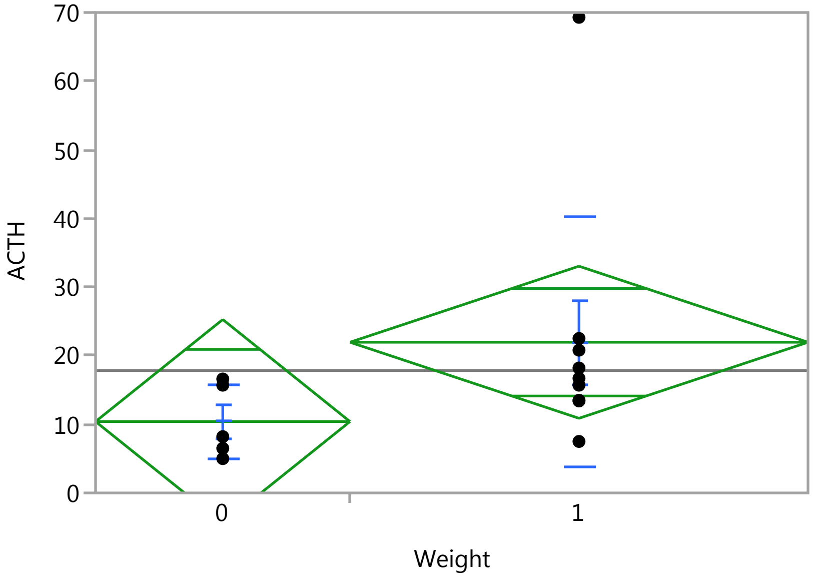

### Fit Group

#### Oneway Analysis of ACTH By Weight IBS-subtype=1

Weight

##### Oneway Anova

###### Summary of Fit

| | |
|---|---|
| Rsquare | 0.133187 |
| Adj Rsquare | 0.060952 |
| Root Mean Square Error | 15.25719 |
| Mean of Response | 17.82857 |
| Observations (or Sum Wgts) | 14 |

###### t Test

1-0

Assuming equal variances

| | | | |
|---|---|---|---|
| Difference | 11.556 | t Ratio | 1.357871 |
| Std Err Dif | 8.510 | DF | 12 |
| Upper CL Dif | 30.097 | Prob > \|t\| | 0.1995 |
| Lower CL Dif | -6.986 | Prob > t | 0.0997 |
| Confidence | 0.95 | Prob < t | 0.9003 |

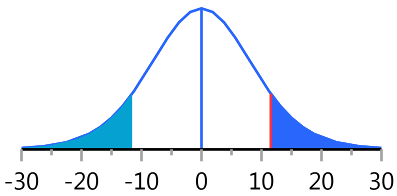


###### Analysis of Variance

| Source | DF | Sum of Squares | Mean Square | F Ratio | Prob > F |
|---|---|---|---|---|---|
| Weight | 1 | 429.2063 | 429.206 | 1.8438 | 0.1995 |
| Error | 12 | 2793.3822 | 232.782 | | |
| C. Total | 13 | 3222.5886 | | | |

###### Means for Oneway Anova

| Level | Number | Mean | Std Error | Lower 95% | Upper 95% |
|---|---|---|---|---|---|
| 0 | 5 | 10.4000 | 6.8232 | -4.47 | 25.267 |
| 1 | 9 | 21.9556 | 5.0857 | 10.87 | 33.036 |

Std Error uses a pooled estimate of error variance

##### Means and Std Deviations

| Level | Number | Mean | Std Dev | Std Err Mean | Lower 95% | Upper 95% |
|---|---|---|---|---|---|---|
| 0 | 5 | 10.4000 | 5.3791 | 2.4056 | 3.7209 | 17.079 |
| 1 | 9 | 21.9556 | 18.2950 | 6.0983 | 7.8928 | 36.018 |

##### t Test

1-0

Assuming unequal variances

| | | | |
|---|---|---|---|
| Difference | 11.556 | t Ratio | 1.762688 |
| Std Err Dif | 6.556 | DF | 10.18997 |
| Upper CL Dif | 26.126 | Prob > \|t\| | 0.1079 |
| Lower CL Dif | -3.015 | Prob > t | 0.0539 |
| Confidence | 0.95 | Prob < t | 0.9461 |

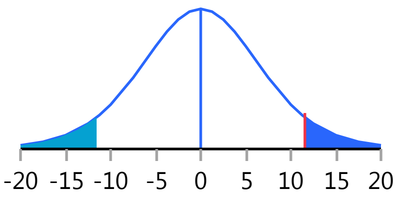

#### Fit Group

##### Oneway Analysis of LDH By Weight IBS-subtype=1

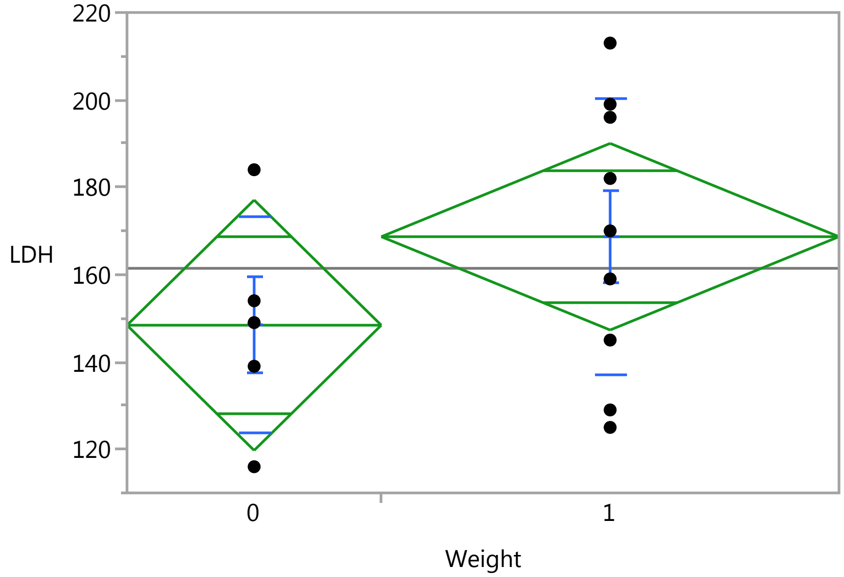


##### Oneway Anova

###### Summary of Fit

| | |
|---|---|
| Rsquare | 0.112807 |
| Adj Rsquare | 0.038874 |
| Root Mean Square Error | 29.41542 |
| Mean of Response | 161.4286 |
| Observations (or Sum Wgts) | 14 |

###### t Test

1-0

Assuming equal variances

| | | | |
|---|---|---|---|
| Difference | 20.267 | t Ratio | 1.235235 |
| Std Err Dif | 16.407 | DF | 12 |
| Upper CL Dif | 56.015 | Prob > \|t\| | 0.2404 |
| Lower CL Dif | -15.481 | Prob > t | 0.1202 |
| Confidence | 0.95 | Prob < t | 0.8798 |

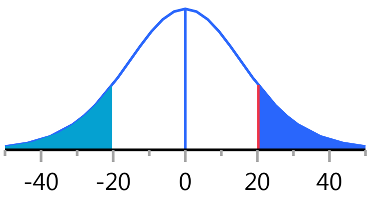


###### Analysis of Variance

| Source | DF | Sum of Squares | Mean Square | F Ratio | Prob > F |
|---|---|---|---|---|---|
| Weight | 1 | 1320.229 | 1320.23 | 1.5258 | 0.2404 |
| Error | 12 | 10383.200 | 865.27 | | |
| C. Total | 13 | 11703.429 | | | |

###### Means for Oneway Anova

| Level | Number | Mean | Std Error | Lower 95% | Upper 95% |
|---|---|---|---|---|---|
| 0 | 5 | 148.400 | 13.155 | 119.74 | 177.06 |
| 1 | 9 | 168.667 | 9.805 | 147.30 | 190.03 |

Std Error uses a pooled estimate of error variance

### Fit Group

#### Oneway Analysis of LDH By Weight IBS-subtype=1

##### Means and Std Deviations

| Level | Number | Mean | Std Dev | Std Err Mean | Lower 95% | Upper 95% |
|---|---|---|---|---|---|---|
| 0 | 5 | 148.400 | 24.6840 | 11.039 | 117.75 | 179.05 |
| 1 | 9 | 168.667 | 31.5159 | 10.505 | 144.44 | 192.89 |

##### t Test

1-0

Assuming unequal variances

| | | | |
|---|---|---|---|
| Difference | 20.267 | t Ratio | 1.329938 |
| Std Err Dif | 15.239 | DF | 10.30135 |
| Upper CL Dif | 54.087 | Prob > \|t\| | 0.2122 |
| Lower CL Dif | -13.553 | Prob > t | 0.1061 |
| Confidence | 0.95 | Prob < t | 0.8939 |

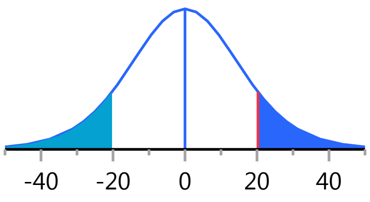


#### Oneway Analysis of Lymphocytes By Weight IBS-subtype=1

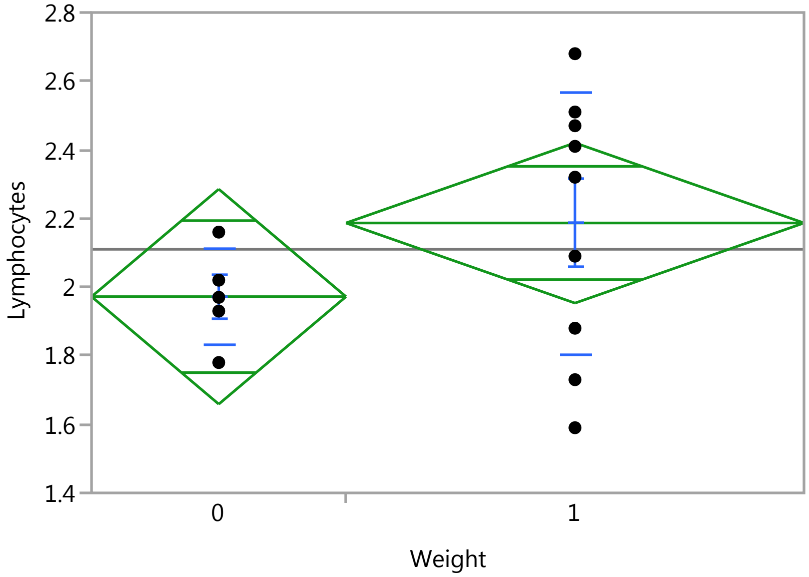


##### Oneway Anova

###### Summary of Fit

| | |
|---|---|
| Rsquare | 0.106699 |
| Adj Rsquare | 0.032258 |
| Root Mean Square Error | 0.321465 |
| Mean of Response | 2.11 |
| Observations (or Sum Wgts) | 14 |

###### t Test

1-0

### Fit Group

#### Oneway Analysis of Lymphocytes By Weight IBS-subtype=1

##### Oneway Anova

###### t Test

Assuming equal variances

| | | | |
|---|---|---|---|
| Difference | 0.21467 | t Ratio | 1.197216 |
| Std Err Dif | 0.17930 | DF | 12 |
| Upper CL Dif | 0.60534 | Prob > \|t\| | 0.2543 |
| Lower CL Dif | -0.17600 | Prob > t | 0.1272 |
| Confidence | 0.95 | Prob < t | 0.8728 |

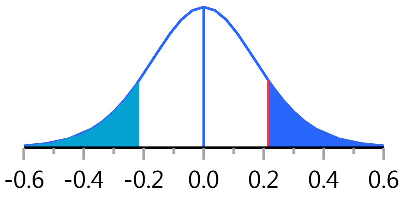


###### Analysis of Variance

| Source | DF | Sum of Squares | Mean Square | F Ratio | Prob > F |
|---|---|---|---|---|---|
| Weight | 1 | 0.1481200 | 0.148120 | 1.4333 | 0.2543 |
| Error | 12 | 1.2400800 | 0.103340 | | |
| C. Total | 13 | 1.3882000 | | | |

###### Means for Oneway Anova

| Level | Number | Mean | Std Error | Lower 95% | Upper 95% |
|---|---|---|---|---|---|
| 0 | 5 | 1.97200 | 0.14376 | 1.6588 | 2.2852 |
| 1 | 9 | 2.18667 | 0.10716 | 1.9532 | 2.4201 |

Std Error uses a pooled estimate of error variance

##### Means and Std Deviations

| Level | Number | Mean | Std Dev | Std Err Mean | Lower 95% | Upper 95% |
|---|---|---|---|---|---|---|
| 0 | 5 | 1.97200 | 0.138094 | 0.06176 | 1.8005 | 2.1435 |
| 1 | 9 | 2.18667 | 0.381412 | 0.12714 | 1.8935 | 2.4798 |

##### t Test

1-0

Assuming unequal variances

| | | | |
|---|---|---|---|
| Difference | 0.21467 | t Ratio | 1.518762 |
| Std Err Dif | 0.14134 | DF | 10.99628 |
| Upper CL Dif | 0.52577 | Prob > \|t\| | 0.1570 |
| Lower CL Dif | -0.09644 | Prob > t | 0.0785 |
| Confidence | 0.95 | Prob < t | 0.9215 |

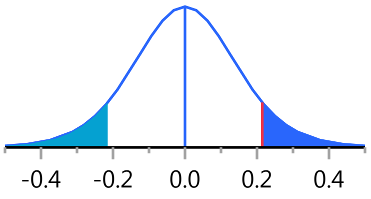


#### Oneway Analysis of IgE By Weight IBS-subtype=1

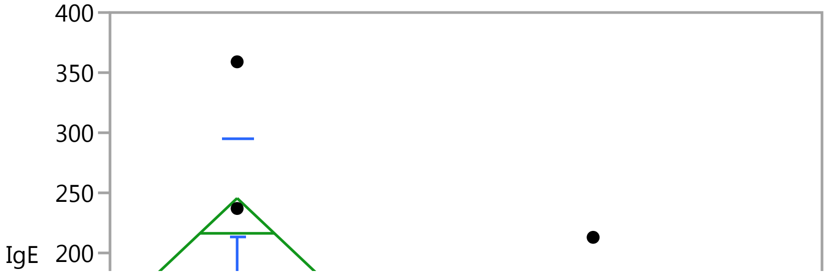

### Fit Group

#### Oneway Analysis of IgE By Weight IBS-subtype=1

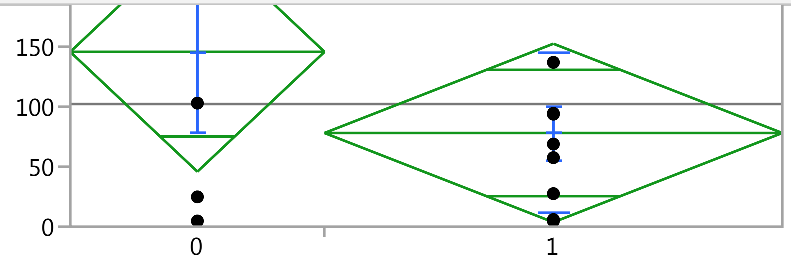


#### Oneway Anova

##### Summary of Fit

| | |
|---|---|
| Rsquare | 0.104489 |
| Adj Rsquare | 0.029864 |
| Root Mean Square Error | 102.4331 |
| Mean of Response | 102.2786 |
| Observations (or Sum Wgts) | 14 |

##### t Test

1-0

Assuming equal variances

| | | | |
|---|---|---|---|
| Difference | -67.61 | t Ratio | -1.18329 |
| Std Err Dif | 57.13 | DF | 12 |
| Upper CL Dif | 56.88 | Prob > \|t\| | 0.2596 |
| Lower CL Dif | -192.09 | Prob > t | 0.8702 |
| Confidence | 0.95 | Prob < t | 0.1298 |

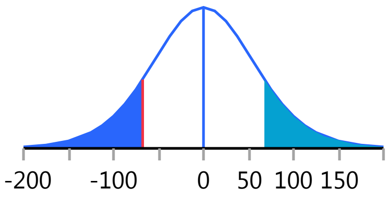


##### Analysis of Variance

| Source | DF | Sum of Squares | Mean Square | F Ratio | Prob > F |
|---|---|---|---|---|---|
| Weight | 1 | 14691.41 | 14691.4 | 1.4002 | 0.2596 |
| Error | 12 | 125910.41 | 10492.5 | | |
| C. Total | 13 | 140601.82 | | | |

##### Means for Oneway Anova

| Level | Number | Mean | Std Error | Lower 95% | Upper 95% |
|---|---|---|---|---|---|
| 0 | 5 | 145.740 | 45.809 | 45.930 | 245.55 |
| 1 | 9 | 78.133 | 34.144 | 3.739 | 152.53 |

Std Error uses a pooled estimate of error variance

#### Means and Std Deviations

| Level | Number | Mean | Std Dev | Std Err Mean | Lower 95% | Upper 95% |
|---|---|---|---|---|---|---|
| 0 | 5 | 145.740 | 150.084 | 67.120 | -40.61 | 332.09 |
| 1 | 9 | 78.133 | 66.904 | 22.301 | 26.71 | 129.56 |

#### t Test

1-0

### Fit Group

#### Oneway Analysis of IgE By Weight IBS-subtype=1

##### t Test

Assuming unequal variances

| | | | |
|---|---|---|---|
| Difference | -67.61 | t Ratio | -0.95587 |
| Std Err Dif | 70.73 | DF | 4.902059 |
| Upper CL Dif | 115.30 | Prob > \|t\| | 0.3839 |
| Lower CL Dif | -250.52 | Prob > t | 0.8081 |
| Confidence | 0.95 | Prob < t | 0.1919 |

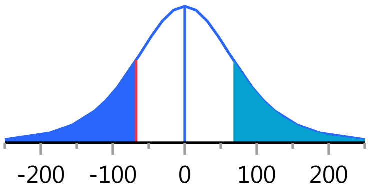


#### Oneway Analysis of Monocytes By Weight IBS-subtype=1

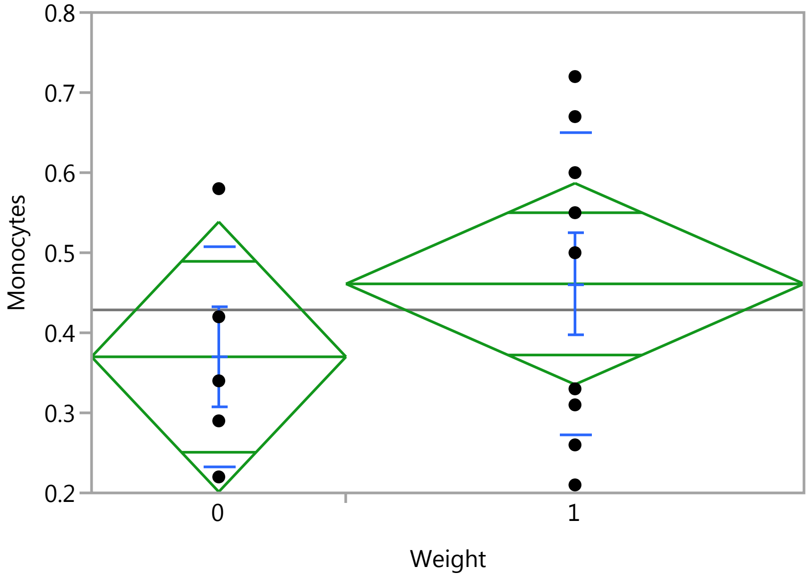


##### Oneway Anova

###### Summary of Fit

| | |
|---|---|
| Rsquare | 0.069131 |
| Adj Rsquare | -0.00844 |
| Root Mean Square Error | 0.173034 |
| Mean of Response | 0.428571 |
| Observations (or Sum Wgts) | 14 |

###### t Test

1-0

Assuming equal variances

| | | | |
|---|---|---|---|
| Difference | 0.09111 | t Ratio | 0.944022 |
| Std Err Dif | 0.09651 | DF | 12 |
| Upper CL Dif | 0.30140 | Prob > \|t\| | 0.3638 |
| Lower CL Dif | -0.11917 | Prob > t | 0.1819 |
| Confidence | 0.95 | Prob < t | 0.8181 |

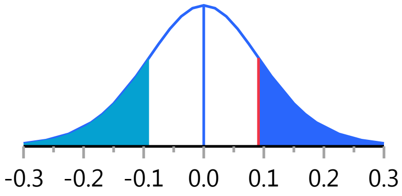

### Fit Group

#### Oneway Analysis of Monocytes By Weight IBS-subtype=1

##### Oneway Anova

###### Analysis of Variance

| Source | DF | Sum of Squares | Mean Square | F Ratio | Prob > F |
|---|---|---|---|---|---|
| Weight | 1 | 0.02668254 | 0.026683 | 0.8912 | 0.3638 |
| Error | 12 | 0.35928889 | 0.029941 | | |
| C. Total | 13 | 0.38597143 | | | |

###### Means for Oneway Anova

| Level | Number | Mean | Std Error | Lower 95% | Upper 95% |
|---|---|---|---|---|---|
| 0 | 5 | 0.370000 | 0.07738 | 0.20140 | 0.53860 |
| 1 | 9 | 0.461111 | 0.05768 | 0.33544 | 0.58678 |

Std Error uses a pooled estimate of error variance

##### Means and Std Deviations

| Level | Number | Mean | Std Dev | Std Err Mean | Lower 95% | Upper 95% |
|---|---|---|---|---|---|---|
| 0 | 5 | 0.370000 | 0.138203 | 0.06181 | 0.19840 | 0.54160 |
| 1 | 9 | 0.461111 | 0.188046 | 0.06268 | 0.31657 | 0.60566 |

##### t Test

1-0

Assuming unequal variances

| | | | |
|---|---|---|---|
| Difference | 0.09111 | t Ratio | 1.035019 |
| Std Err Dif | 0.08803 | DF | 10.7655 |
| Upper CL Dif | 0.28538 | Prob > \|t\| | 0.3234 |
| Lower CL Dif | -0.10315 | Prob > t | 0.1617 |
| Confidence | 0.95 | Prob < t | 0.8383 |

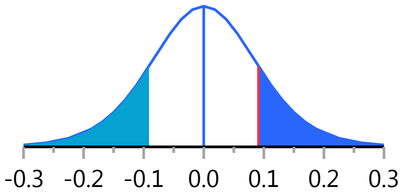


#### Oneway Analysis of ESR By Weight IBS-subtype=1

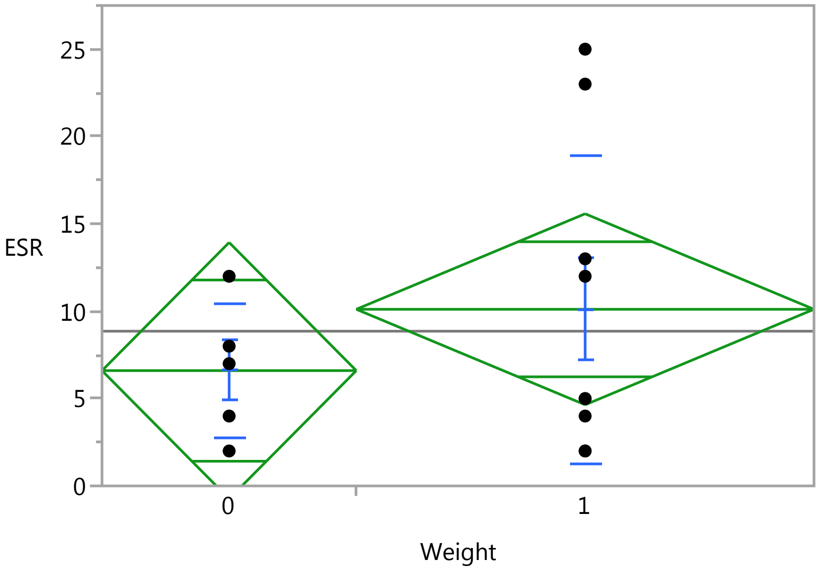

### Fit Group

#### Oneway Analysis of ESR By Weight IBS-subtype=1

Weight

##### Oneway Anova

###### Summary of Fit

| | |
|---|---|
| Rsquare | 0.055057 |
| Adj Rsquare | -0.02369 |
| Root Mean Square Error | 7.528219 |
| Mean of Response | 8.857143 |
| Observations (or Sum Wgts) | 14 |

###### t Test

1-0

Assuming equal variances

| | | | |
|---|---|---|---|
| Difference | 3.511 | t Ratio | 0.83617 |
| Std Err Dif | 4.199 | DF | 12 |
| Upper CL Dif | 12.660 | Prob > \|t\| | 0.4194 |
| Lower CL Dif | -5.638 | Prob > t | 0.2097 |
| Confidence | 0.95 | Prob < t | 0.7903 |

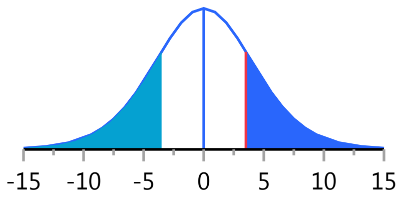


###### Analysis of Variance

| Source | DF | Sum of Squares | Mean Square | F Ratio | Prob > F |
|---|---|---|---|---|---|
| Weight | 1 | 39.62540 | 39.6254 | 0.6992 | 0.4194 |
| Error | 12 | 680.08889 | 56.6741 | | |
| C. Total | 13 | 719.71429 | | | |

###### Means for Oneway Anova

| Level | Number | Mean | Std Error | Lower 95% | Upper 95% |
|---|---|---|---|---|---|
| 0 | 5 | 6.6000 | 3.3667 | -0.735 | 13.935 |
| 1 | 9 | 10.1111 | 2.5094 | 4.644 | 15.579 |

Std Error uses a pooled estimate of error variance

##### Means and Std Deviations

| Level | Number | Mean | Std Dev | Std Err Mean | Lower 95% | Upper 95% |
|---|---|---|---|---|---|---|
| 0 | 5 | 6.6000 | 3.84708 | 1.7205 | 1.8232 | 11.377 |
| 1 | 9 | 10.1111 | 8.80972 | 2.9366 | 3.3394 | 16.883 |

##### t Test

1-0

Assuming unequal variances

| | | | |
|---|---|---|---|
| Difference | 3.511 | t Ratio | 1.031634 |
| Std Err Dif | 3.403 | DF | 11.68184 |
| Upper CL Dif | 10.949 | Prob > \|t\| | 0.3231 |
| Lower CL Dif | -3.927 | Prob > t | 0.1616 |
| Confidence | 0.95 | Prob < t | 0.8384 |

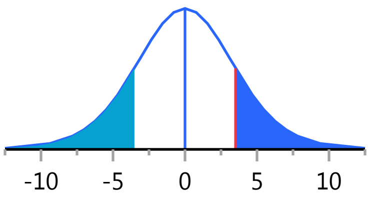

#### Fit Group

##### Oneway Analysis of WBC By Weight IBS-subtype=1

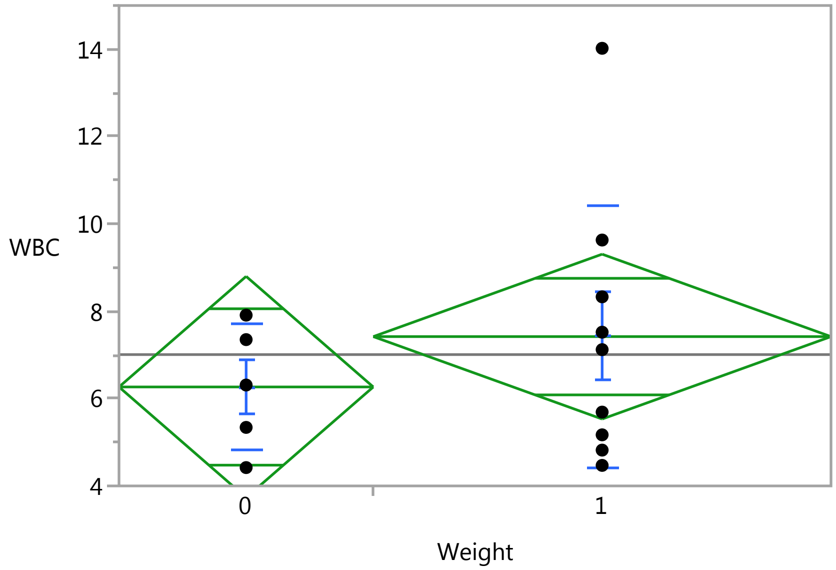


#### Oneway Anova

##### Summary of Fit

| | |
|---|---|
| Rsquare | 0.050081 |
| Adj Rsquare | -0.02908 |
| Root Mean Square Error | 2.598638 |
| Mean of Response | 7.007143 |
| Observations (or Sum Wgts) | 14 |

##### t Test

1-0

Assuming equal variances

| | | | |
|---|---|---|---|
| Difference | 1.1529 | t Ratio | 0.795397 |
| Std Err Dif | 1.4495 | DF | 12 |
| Upper CL Dif | 4.3110 | Prob > \|t\| | 0.4418 |
| Lower CL Dif | -2.0052 | Prob > t | 0.2209 |
| Confidence | 0.95 | Prob < t | 0.7791 |

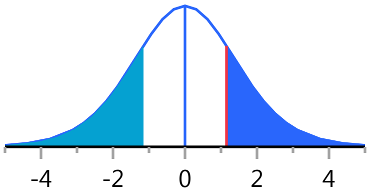


##### Analysis of Variance

| Source | DF | Sum of Squares | Mean Square | F Ratio | Prob > F |
|---|---|---|---|---|---|
| Weight | 1 | 4.272277 | 4.27228 | 0.6327 | 0.4418 |
| Error | 12 | 81.035009 | 6.75292 | | |
| C. Total | 13 | 85.307286 | | | |

##### Means for Oneway Anova

| Level | Number | Mean | Std Error | Lower 95% | Upper 95% |
|---|---|---|---|---|---|
| 0 | 5 | 6.26600 | 1.1621 | 3.7339 | 8.7981 |
| 1 | 9 | 7.41889 | 0.8662 | 5.5316 | 9.3062 |

Std Error uses a pooled estimate of error variance

### Fit Group

#### Oneway Analysis of WBC By Weight IBS-subtype=1

##### Means and Std Deviations

| Level | Number | Mean | Std Dev | Std Err Mean | Lower 95% | Upper 95% |
|---|---|---|---|---|---|---|
| 0 | 5 | 6.26600 | 1.42697 | 0.6382 | 4.4942 | 8.0378 |
| 1 | 9 | 7.41889 | 3.01849 | 1.0062 | 5.0987 | 9.7391 |

##### t Test

1-0

Assuming unequal variances

| | | | |
|---|---|---|---|
| Difference | 1.1529 | t Ratio | 0.967616 |
| Std Err Dif | 1.1915 | DF | 11.88455 |
| Upper CL Dif | 3.7517 | Prob > \|t\| | 0.3525 |
| Lower CL Dif | -1.4459 | Prob > t | 0.1763 |
| Confidence | 0.95 | Prob < t | 0.8237 |

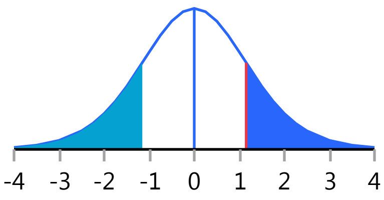


#### Oneway Analysis of Neutrophils By Weight IBS-subtype=1

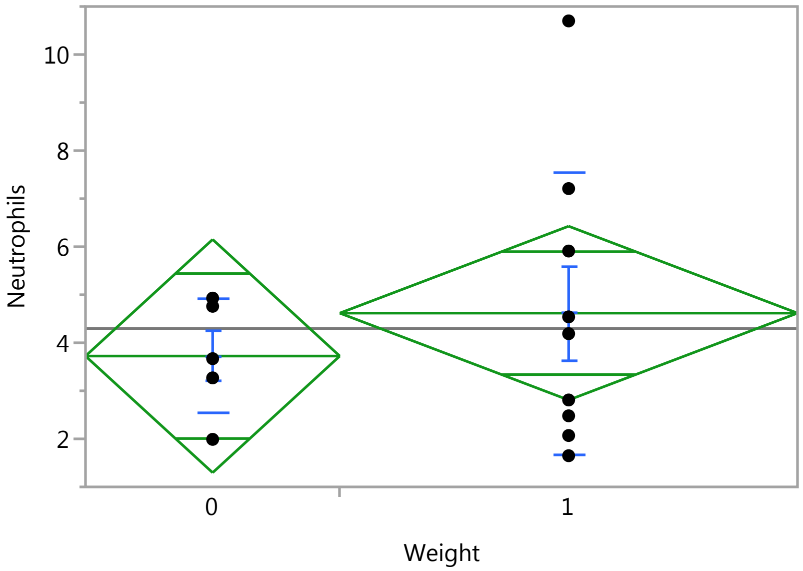


##### Oneway Anova

###### Summary of Fit

| | |
|---|---|
| Rsquare | 0.033338 |
| Adj Rsquare | -0.04722 |
| Root Mean Square Error | 2.490858 |
| Mean of Response | 4.298571 |
| Observations (or Sum Wgts) | 14 |

###### t Test

1-0

### Fit Group

#### Oneway Analysis of Neutrophils By Weight IBS-subtype=1

##### Oneway Anova

###### t Test

Assuming equal variances

| | | | |
|---|---|---|---|
| Difference | 0.8938 | t Ratio | 0.643314 |
| Std Err Dif | 1.3893 | DF | 12 |
| Upper CL Dif | 3.9209 | Prob > \|t\| | 0.5321 |
| Lower CL Dif | -2.1333 | Prob > t | 0.2661 |
| Confidence | 0.95 | Prob < t | 0.7339 |

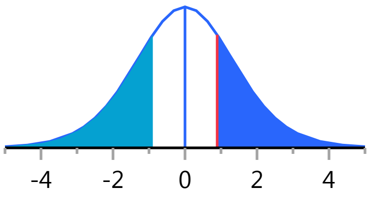


###### Analysis of Variance

| Source | DF | Sum of Squares | Mean Square | F Ratio | Prob > F |
|---|---|---|---|---|---|
| Weight | 1 | 2.567696 | 2.56770 | 0.4139 | 0.5321 |
| Error | 12 | 74.452476 | 6.20437 | | |
| C. Total | 13 | 77.020171 | | | |

###### Means for Oneway Anova

| Level | Number | Mean | Std Error | Lower 95% | Upper 95% |
|---|---|---|---|---|---|
| 0 | 5 | 3.72400 | 1.1139 | 1.2969 | 6.1511 |
| 1 | 9 | 4.61778 | 0.8303 | 2.8087 | 6.4268 |

Std Error uses a pooled estimate of error variance

##### Means and Std Deviations

| Level | Number | Mean | Std Dev | Std Err Mean | Lower 95% | Upper 95% |
|---|---|---|---|---|---|---|
| 0 | 5 | 3.72400 | 1.19828 | 0.53589 | 2.2361 | 5.2119 |
| 1 | 9 | 4.61778 | 2.93063 | 0.97688 | 2.3651 | 6.8705 |

##### t Test

1-0

Assuming unequal variances

| | | | |
|---|---|---|---|
| Difference | 0.8938 | t Ratio | 0.802162 |
| Std Err Dif | 1.1142 | DF | 11.46318 |
| Upper CL Dif | 3.3341 | Prob > \|t\| | 0.4388 |
| Lower CL Dif | -1.5465 | Prob > t | 0.2194 |
| Confidence | 0.95 | Prob < t | 0.7806 |

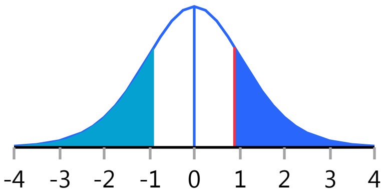


#### Oneway Analysis of CRP By Weight IBS-subtype=1

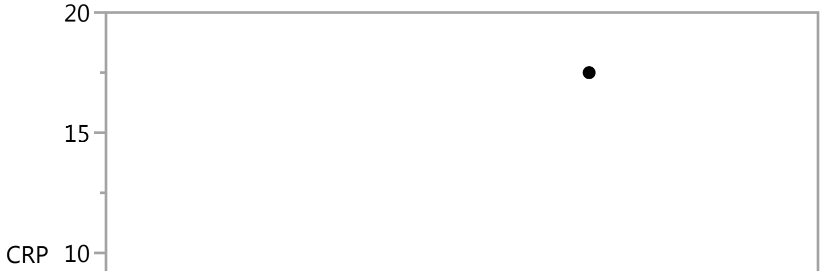

#### Fit Group

##### Oneway Analysis of CRP By Weight IBS-subtype=1

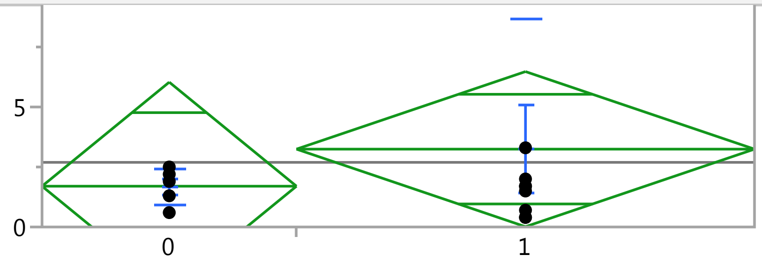


#### Oneway Anova

##### Summary of Fit

| | |
|---|---|
| Rsquare | 0.031267 |
| Adj Rsquare | -0.04946 |
| Root Mean Square Error | 4.449178 |
| Mean of Response | 2.692857 |
| Observations (or Sum Wgts) | 14 |

##### t Test

1-0

Assuming equal variances

| | | | |
|---|---|---|---|
| Difference | 1.5444 | t Ratio | 0.62235 |
| Std Err Dif | 2.4816 | DF | 12 |
| Upper CL Dif | 6.9515 | Prob > \|t\| | 0.5454 |
| Lower CL Dif | -3.8626 | Prob > t | 0.2727 |
| Confidence | 0.95 | Prob < t | 0.7273 |

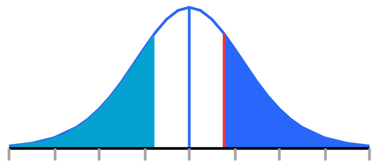


##### Analysis of Variance

| Source | DF | Sum of Squares | Mean Square | F Ratio | Prob > F |
|---|---|---|---|---|---|
| Weight | 1 | 7.66706 | 7.6671 | 0.3873 | 0.5454 |
| Error | 12 | 237.54222 | 19.7952 | | |
| C. Total | 13 | 245.20929 | | | |

##### Means for Oneway Anova

| Level | Number | Mean | Std Error | Lower 95% | Upper 95% |
|---|---|---|---|---|---|
| 0 | 5 | 1.70000 | 1.9897 | -2.635 | 6.0353 |
| 1 | 9 | 3.24444 | 1.4831 | 0.013 | 6.4758 |

Std Error uses a pooled estimate of error variance

#### Means and Std Deviations

| Level | Number | Mean | Std Dev | Std Err Mean | Lower 95% | Upper 95% |
|---|---|---|---|---|---|---|
| 0 | 5 | 1.70000 | 0.75829 | 0.3391 | 0.7585 | 2.6415 |
| 1 | 9 | 3.24444 | 5.42266 | 1.8076 | -0.9238 | 7.4127 |

#### t Test

1-0

### Fit Group

#### Oneway Analysis of CRP By Weight IBS-subtype=1

##### t Test

Assuming unequal variances

| | | | |
|---|---|---|---|
| Difference | 1.5444 | t Ratio | 0.839787 |
| Std Err Dif | 1.8391 | DF | 8.551886 |
| Upper CL Dif | 5.7382 | Prob > \|t\| | 0.4239 |
| Lower CL Dif | -2.6493 | Prob > t | 0.2119 |
| Confidence | 0.95 | Prob < t | 0.7881 |

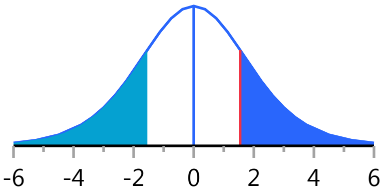


#### Oneway Analysis of Basophil_PCT By Weight IBS-subtype=1

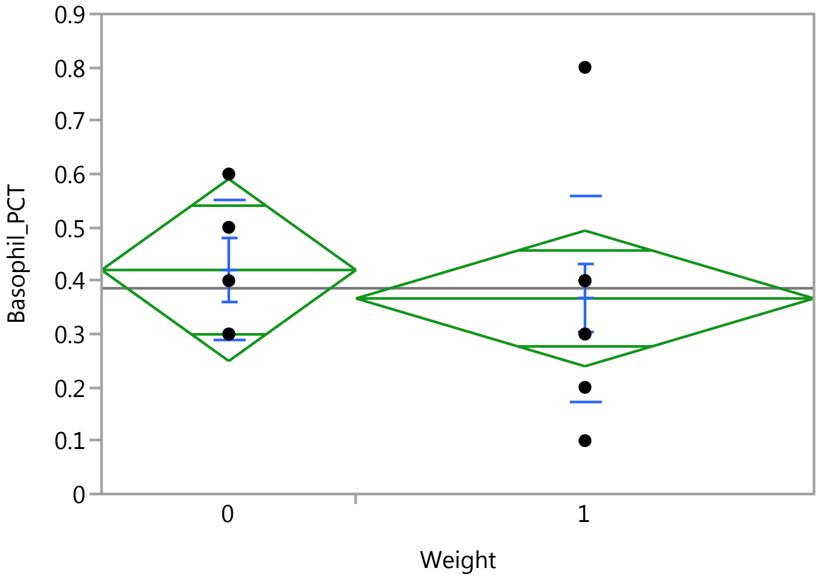


##### Oneway Anova

###### Summary of Fit

| | |
|---|---|
| Rsquare | 0.024242 |
| Adj Rsquare | -0.05707 |
| Root Mean Square Error | 0.175119 |
| Mean of Response | 0.385714 |
| Observations (or Sum Wgts) | 14 |

###### t Test

1-0

Assuming equal variances

| | | | |
|---|---|---|---|
| Difference | -0.05333 | t Ratio | -0.54602 |
| Std Err Dif | 0.09768 | DF | 12 |
| Upper CL Dif | 0.15949 | Prob > \|t\| | 0.5951 |
| Lower CL Dif | -0.26615 | Prob > t | 0.7025 |
| Confidence | 0.95 | Prob < t | 0.2975 |

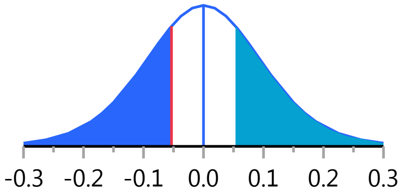

#### Fit Group

##### Oneway Analysis of Basophil_PCT By Weight IBS-subtype=1

###### Oneway Anova

###### Analysis of Variance

| Source | DF | Sum of Squares | Mean Square | F Ratio | Prob > F |
|---|---|---|---|---|---|
| Weight | 1 | 0.00914286 | 0.009143 | 0.2981 | 0.5951 |
| Error | 12 | 0.36800000 | 0.030667 | | |
| C. Total | 13 | 0.37714286 | | | |

###### Means for Oneway Anova

| Level | Number | Mean | Std Error | Lower 95% | Upper 95% |
|---|---|---|---|---|---|
| 0 | 5 | 0.420000 | 0.07832 | 0.24936 | 0.59064 |
| 1 | 9 | 0.366667 | 0.05837 | 0.23948 | 0.49385 |

Std Error uses a pooled estimate of error variance

###### Means and Std Deviations

| Level | Number | Mean | Std Dev | Std Err Mean | Lower 95% | Upper 95% |
|---|---|---|---|---|---|---|
| 0 | 5 | 0.420000 | 0.130384 | 0.05831 | 0.25811 | 0.58189 |
| 1 | 9 | 0.366667 | 0.193649 | 0.06455 | 0.21781 | 0.51552 |

###### t Test

1-0

Assuming unequal variances

| | | | |
|---|---|---|---|
| Difference | -0.05333 | t Ratio | -0.61312 |
| Std Err Dif | 0.08699 | DF | 11.3148 |
| Upper CL Dif | 0.13747 | Prob > \|t\| | 0.5519 |
| Lower CL Dif | -0.24414 | Prob > t | 0.7240 |
| Confidence | 0.95 | Prob < t | 0.2760 |

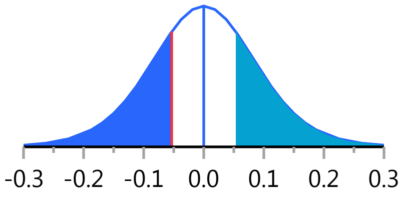


##### Oneway Analysis of Monocytes_PCT By Weight IBS-subtype=1

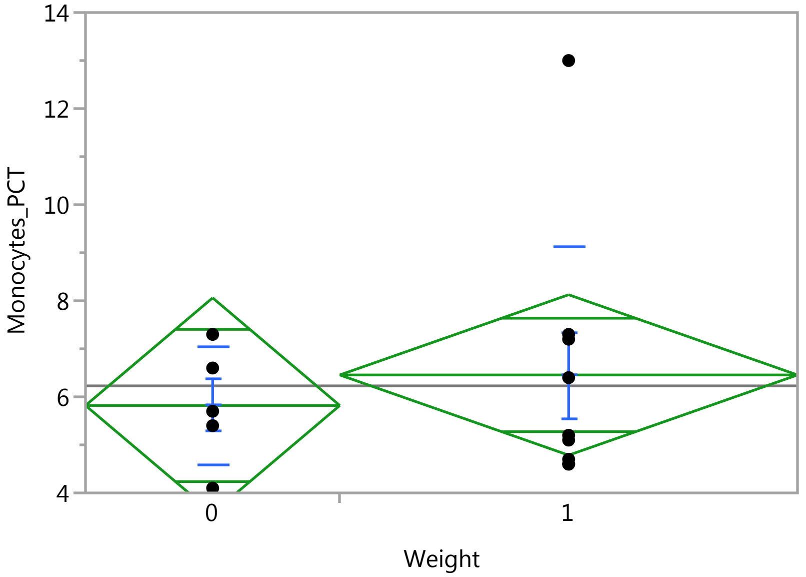

### Fit Group

#### Oneway Analysis of Monocytes_PCT By Weight IBS-subtype=1

Weight

##### Oneway Anova

###### Summary of Fit

| | |
|---|---|
| Rsquare | 0.020058 |
| Adj Rsquare | -0.0616 |
| Root Mean Square Error | 2.299098 |
| Mean of Response | 6.228571 |
| Observations (or Sum Wgts) | 14 |

###### t Test

1-0

Assuming equal variances

| | | | |
|---|---|---|---|
| Difference | 0.6356 | t Ratio | 0.495608 |
| Std Err Dif | 1.2824 | DF | 12 |
| Upper CL Dif | 3.4296 | Prob > \|t\| | 0.6291 |
| Lower CL Dif | -2.1585 | Prob > t | 0.3146 |
| Confidence | 0.95 | Prob < t | 0.6854 |

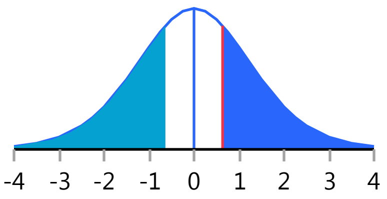


###### Analysis of Variance

| Source | DF | Sum of Squares | Mean Square | F Ratio | Prob > F |
|---|---|---|---|---|---|
| Weight | 1 | 1.298349 | 1.29835 | 0.2456 | 0.6291 |
| Error | 12 | 63.430222 | 5.28585 | | |
| C. Total | 13 | 64.728571 | | | |

###### Means for Oneway Anova

| Level | Number | Mean | Std Error | Lower 95% | Upper 95% |
|---|---|---|---|---|---|
| 0 | 5 | 5.82000 | 1.0282 | 3.5798 | 8.0602 |
| 1 | 9 | 6.45556 | 0.7664 | 4.7858 | 8.1253 |

Std Error uses a pooled estimate of error variance

##### Means and Std Deviations

| Level | Number | Mean | Std Dev | Std Err Mean | Lower 95% | Upper 95% |
|---|---|---|---|---|---|---|
| 0 | 5 | 5.82000 | 1.21943 | 0.54534 | 4.3059 | 7.3341 |
| 1 | 9 | 6.45556 | 2.68054 | 0.89351 | 4.3951 | 8.5160 |

##### t Test

1-0

Assuming unequal variances

| | | | |
|---|---|---|---|
| Difference | 0.6356 | t Ratio | 0.607149 |
| Std Err Dif | 1.0468 | DF | 11.79644 |
| Upper CL Dif | 2.9207 | Prob > \|t\| | 0.5553 |
| Lower CL Dif | -1.6496 | Prob > t | 0.2776 |
| Confidence | 0.95 | Prob < t | 0.7224 |

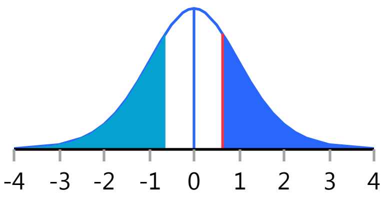

#### Fit Group

##### Oneway Analysis of IgA By Weight IBS-subtype=1

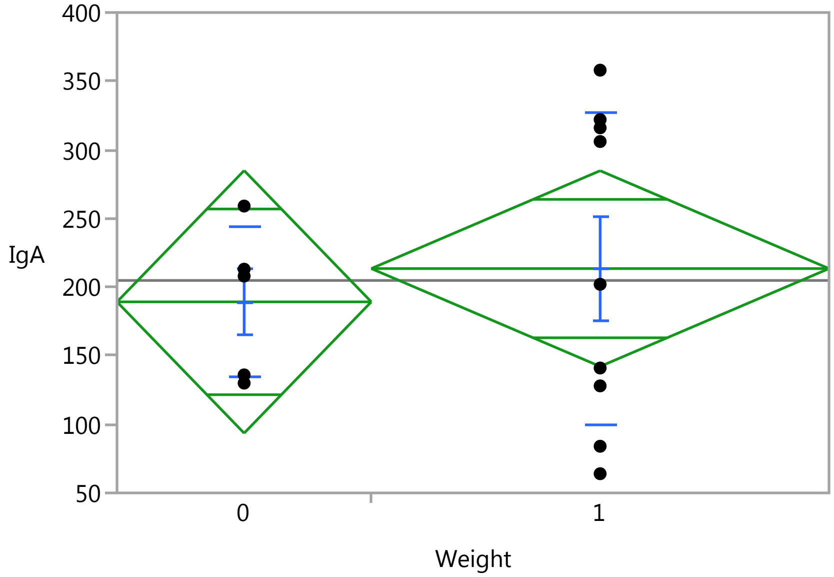


#### Oneway Anova

##### Summary of Fit

| | |
|---|---|
| Rsquare | 0.016073 |
| Adj Rsquare | -0.06592 |
| Root Mean Square Error | 98.17545 |
| Mean of Response | 204.7857 |
| Observations (or Sum Wgts) | 14 |

##### t Test

1-0

Assuming equal variances

| | | | |
|---|---|---|---|
| Difference | 24.24 | t Ratio | 0.442743 |
| Std Err Dif | 54.76 | DF | 12 |
| Upper CL Dif | 143.56 | Prob > \|t\| | 0.6658 |
| Lower CL Dif | -95.07 | Prob > t | 0.3329 |
| Confidence | 0.95 | Prob < t | 0.6671 |

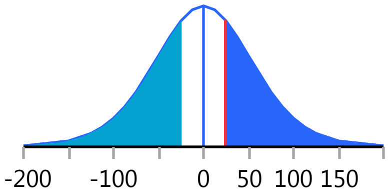


##### Analysis of Variance

| Source | DF | Sum of Squares | Mean Square | F Ratio | Prob > F |
|---|---|---|---|---|---|
| Weight | 1 | 1889.33 | 1889.33 | 0.1960 | 0.6658 |
| Error | 12 | 115661.02 | 9638.42 | | |
| C. Total | 13 | 117550.36 | | | |

##### Means for Oneway Anova

| Level | Number | Mean | Std Error | Lower 95% | Upper 95% |
|---|---|---|---|---|---|
| 0 | 5 | 189.200 | 43.905 | 93.54 | 284.86 |
| 1 | 9 | 213.444 | 32.725 | 142.14 | 284.75 |

Std Error uses a pooled estimate of error variance

### Fit Group

#### Oneway Analysis of IgA By Weight IBS-subtype=1

##### Means and Std Deviations

| Level | Number | Mean | Std Dev | Std Err Mean | Lower 95% | Upper 95% |
|---|---|---|---|---|---|---|
| 0 | 5 | 189.200 | 55.061 | 24.624 | 120.83 | 257.57 |
| 1 | 9 | 213.444 | 113.762 | 37.921 | 126.00 | 300.89 |

##### t Test

1-0

Assuming unequal variances

| | | | |
|---|---|---|---|
| Difference | 24.24 | t Ratio | 0.536214 |
| Std Err Dif | 45.21 | DF | 11.92757 |
| Upper CL Dif | 122.82 | Prob > \|t\| | 0.6017 |
| Lower CL Dif | -74.34 | Prob > t | 0.3008 |
| Confidence | 0.95 | Prob < t | 0.6992 |

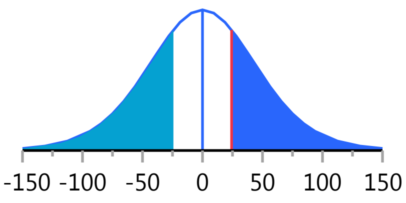


#### Oneway Analysis of IgM By Weight IBS-subtype=1

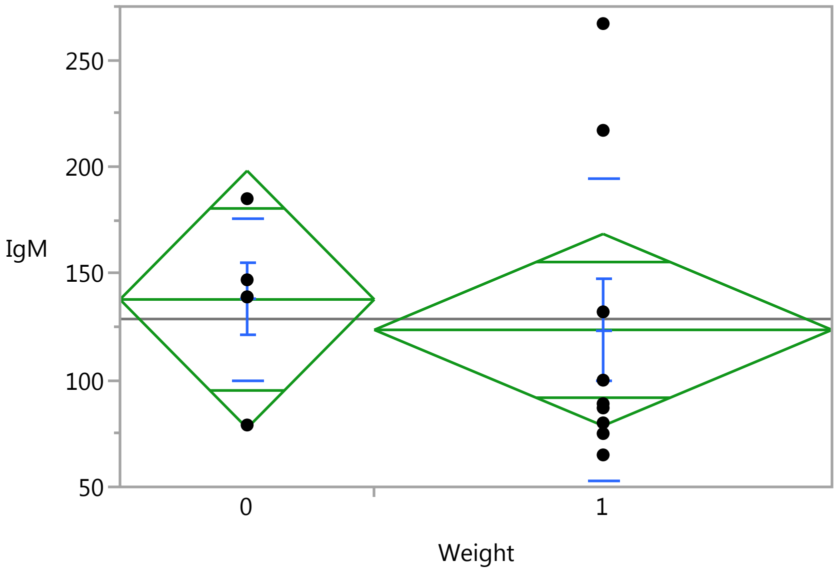


##### Oneway Anova

###### Summary of Fit

| | |
|---|---|
| Rsquare | 0.014004 |
| Adj Rsquare | -0.06816 |
| Root Mean Square Error | 61.86075 |
| Mean of Response | 128.6429 |
| Observations (or Sum Wgts) | 14 |

###### t Test

1-0

### Fit Group

#### Oneway Analysis of IgM By Weight IBS-subtype=1

##### Oneway Anova

###### t Test

Assuming equal variances

| | | | |
|---|---|---|---|
| Difference | -14.244 | t Ratio | -0.41283 |
| Std Err Dif | 34.504 | DF | 12 |
| Upper CL Dif | 60.934 | Prob > \|t\| | 0.6870 |
| Lower CL Dif | -89.423 | Prob > t | 0.6565 |
| Confidence | 0.95 | Prob < t | 0.3435 |

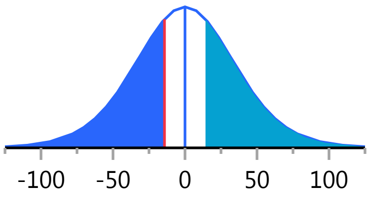


###### Analysis of Variance

| Source | DF | Sum of Squares | Mean Square | F Ratio | Prob > F |
|---|---|---|---|---|---|
| Weight | 1 | 652.192 | 652.19 | 0.1704 | 0.6870 |
| Error | 12 | 45921.022 | 3826.75 | | |
| C. Total | 13 | 46573.214 | | | |

###### Means for Oneway Anova

| Level | Number | Mean | Std Error | Lower 95% | Upper 95% |
|---|---|---|---|---|---|
| 0 | 5 | 137.800 | 27.665 | 77.523 | 198.08 |
| 1 | 9 | 123.556 | 20.620 | 78.628 | 168.48 |

Std Error uses a pooled estimate of error variance

##### Means and Std Deviations

| Level | Number | Mean | Std Dev | Std Err Mean | Lower 95% | Upper 95% |
|---|---|---|---|---|---|---|
| 0 | 5 | 137.800 | 37.9895 | 16.989 | 90.630 | 184.97 |
| 1 | 9 | 123.556 | 70.8416 | 23.614 | 69.102 | 178.01 |

##### t Test

1-0

Assuming unequal variances

| | | | |
|---|---|---|---|
| Difference | -14.244 | t Ratio | -0.48966 |
| Std Err Dif | 29.090 | DF | 11.99676 |
| Upper CL Dif | 49.140 | Prob > \|t\| | 0.6332 |
| Lower CL Dif | -77.629 | Prob > t | 0.6834 |
| Confidence | 0.95 | Prob < t | 0.3166 |

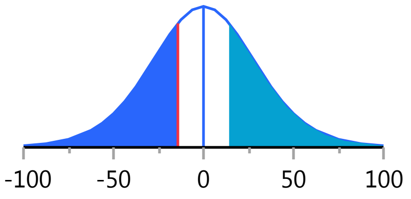


#### Oneway Analysis of IgG By Weight IBS-subtype=1

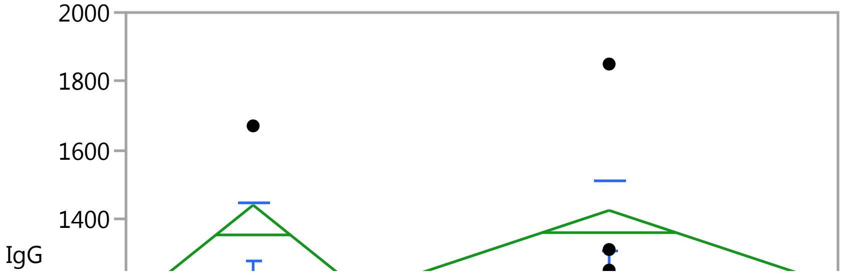

#### Fit Group

##### Oneway Analysis of IgG By Weight IBS-subtype=1

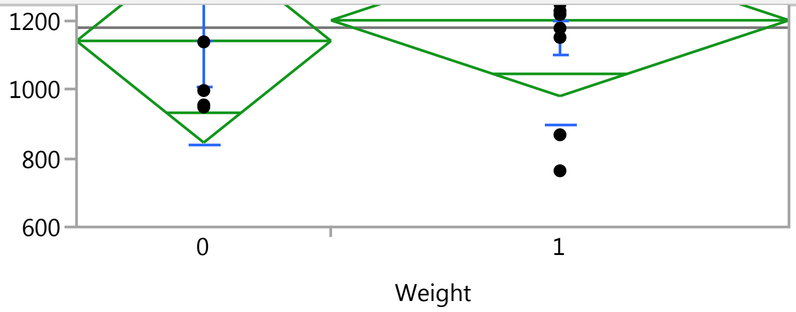


#### Oneway Anova

##### Summary of Fit

| | |
|---|---|
| Rsquare | 0.010333 |
| Adj Rsquare | -0.07214 |
| Root Mean Square Error | 304.3534 |
| Mean of Response | 1181.429 |
| Observations (or Sum Wgts) | 14 |

##### t Test

1-0

Assuming equal variances

| | | | |
|---|---|---|---|
| Difference | 60.09 | t Ratio | 0.353963 |
| Std Err Dif | 169.76 | DF | 12 |
| Upper CL Dif | 429.96 | Prob > \|t\| | 0.7295 |
| Lower CL Dif | -309.79 | Prob > t | 0.3648 |
| Confidence | 0.95 | Prob < t | 0.6352 |

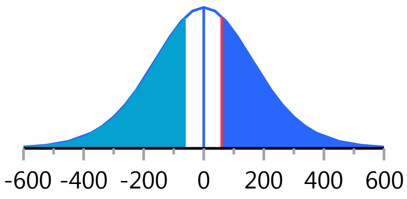


##### Analysis of Variance

| Source | DF | Sum of Squares | Mean Square | F Ratio | Prob > F |
|---|---|---|---|---|---|
| Weight | 1 | 11605.7 | 11605.7 | 0.1253 | 0.7295 |
| Error | 12 | 1111571.7 | 92631.0 | | |
| C. Total | 13 | 1123177.4 | | | |

##### Means for Oneway Anova

| Level | Number | Mean | Std Error | Lower 95% | Upper 95% |
|---|---|---|---|---|---|
| 0 | 5 | 1142.80 | 136.11 | 846.24 | 1439.4 |
| 1 | 9 | 1202.89 | 101.45 | 981.85 | 1423.9 |

Std Error uses a pooled estimate of error variance

#### Means and Std Deviations

| Level | Number | Mean | Std Dev | Std Err Mean | Lower 95% | Upper 95% |
|---|---|---|---|---|---|---|
| 0 | 5 | 1142.80 | 304.541 | 136.19 | 764.66 | 1520.9 |
| 1 | 9 | 1202.89 | 304.260 | 101.42 | 969.01 | 1436.8 |

#### t Test

1-0

### Fit Group

#### Oneway Analysis of IgG By Weight IBS-subtype=1

##### t Test

Assuming unequal variances

| | | | |
|---|---|---|---|
| Difference | 60.09 | t Ratio | 0.353862 |
| Std Err Dif | 169.81 | DF | 8.3781 |
| Upper CL Dif | 448.61 | Prob > \|t\| | 0.7322 |
| Lower CL Dif | -328.44 | Prob > t | 0.3661 |
| Confidence | 0.95 | Prob < t | 0.6339 |

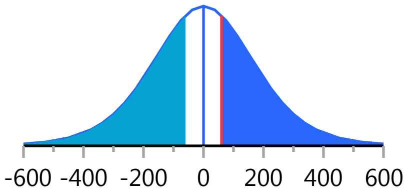


#### Oneway Analysis of HCT By Weight IBS-subtype=1

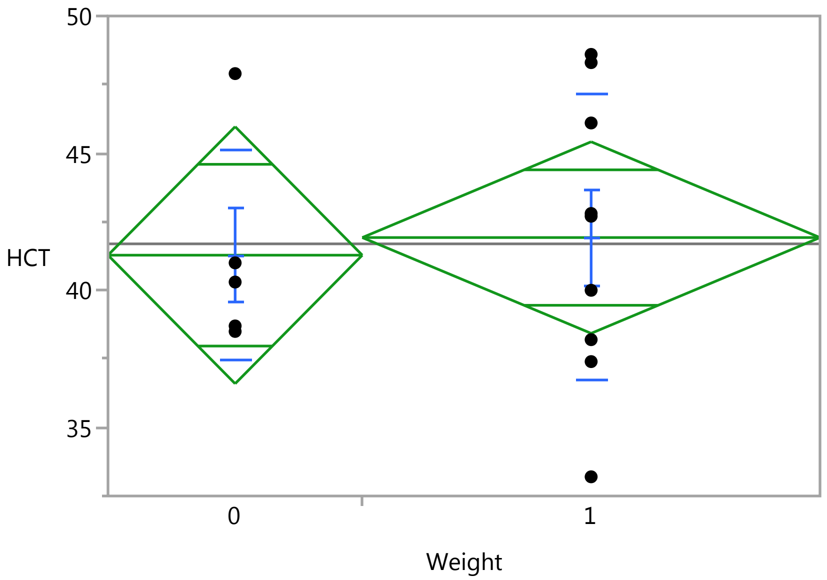


##### Oneway Anova

###### Summary of Fit

| | |
|---|---|
| Rsquare | 0.004756 |
| Adj Rsquare | -0.07818 |
| Root Mean Square Error | 4.808184 |
| Mean of Response | 41.69286 |
| Observations (or Sum Wgts) | 14 |

###### t Test

1-0

Assuming equal variances

| | | | |
|---|---|---|---|
| Difference | 0.6422 | t Ratio | 0.239467 |
| Std Err Dif | 2.6819 | DF | 12 |
| Upper CL Dif | 6.4855 | Prob > \|t\| | 0.8148 |
| Lower CL Dif | -5.2011 | Prob > t | 0.4074 |
| Confidence | 0.95 | Prob < t | 0.5926 |

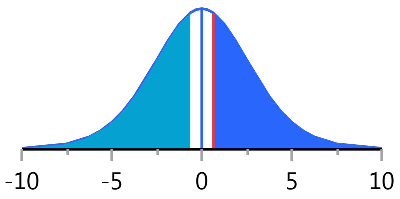

#### Fit Group

##### Oneway Analysis of HCT By Weight IBS-subtype=1

###### Oneway Anova

###### Analysis of Variance

| Source | DF | Sum of Squares | Mean Square | F Ratio | Prob > F |
|---|---|---|---|---|---|
| Weight | 1 | 1.32573 | 1.3257 | 0.0573 | 0.8148 |
| Error | 12 | 277.42356 | 23.1186 | | |
| C. Total | 13 | 278.74929 | | | |

###### Means for Oneway Anova

| Level | Number | Mean | Std Error | Lower 95% | Upper 95% |
|---|---|---|---|---|---|
| 0 | 5 | 41.2800 | 2.1503 | 36.595 | 45.965 |
| 1 | 9 | 41.9222 | 1.6027 | 38.430 | 45.414 |

Std Error uses a pooled estimate of error variance

###### Means and Std Deviations

| Level | Number | Mean | Std Dev | Std Err Mean | Lower 95% | Upper 95% |
|---|---|---|---|---|---|---|
| 0 | 5 | 41.2800 | 3.84864 | 1.7212 | 36.501 | 46.059 |
| 1 | 9 | 41.9222 | 5.22225 | 1.7408 | 37.908 | 45.936 |

###### t Test

1-0

Assuming unequal variances

| | | | |
|---|---|---|---|
| Difference | 0.6422 | t Ratio | 0.262348 |
| Std Err Dif | 2.4480 | DF | 10.74637 |
| Upper CL Dif | 6.0458 | Prob > \|t\| | 0.7980 |
| Lower CL Dif | -4.7613 | Prob > t | 0.3990 |
| Confidence | 0.95 | Prob < t | 0.6010 |

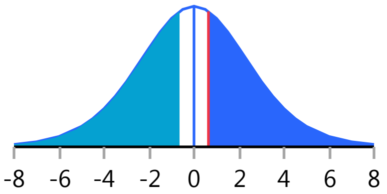


##### Oneway Analysis of MCH By Weight IBS-subtype=1

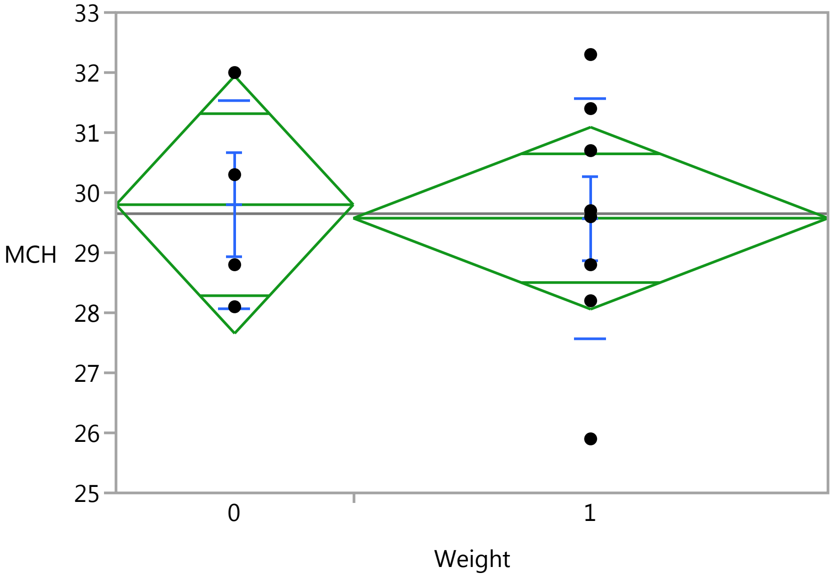

### Fit Group

#### Oneway Analysis of MCH By Weight IBS-subtype=1

Weight

##### Oneway Anova

###### Summary of Fit

| | |
|---|---|
| Rsquare | 0.003634 |
| Adj Rsquare | -0.096 |
| Root Mean Square Error | 1.923928 |
| Mean of Response | 29.65 |
| Observations (or Sum Wgts) | 12 |

###### t Test

1-0

Assuming equal variances

| | | | |
|---|---|---|---|
| Difference | -0.2250 | t Ratio | -0.19098 |
| Std Err Dif | 1.1782 | DF | 10 |
| Upper CL Dif | 2.4001 | Prob > \|t\| | 0.8524 |
| Lower CL Dif | -2.8501 | Prob > t | 0.5738 |
| Confidence | 0.95 | Prob < t | 0.4262 |

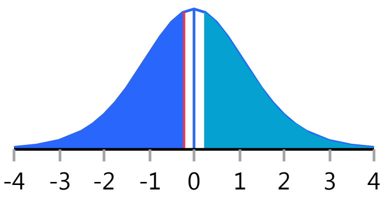


###### Analysis of Variance

| Source | DF | Sum of Squares | Mean Square | F Ratio | Prob > F |
|---|---|---|---|---|---|
| Weight | 1 | 0.135000 | 0.13500 | 0.0365 | 0.8524 |
| Error | 10 | 37.015000 | 3.70150 | | |
| C. Total | 11 | 37.150000 | | | |

###### Means for Oneway Anova

| Level | Number | Mean | Std Error | Lower 95% | Upper 95% |
|---|---|---|---|---|---|
| 0 | 4 | 29.8000 | 0.96196 | 27.657 | 31.943 |
| 1 | 8 | 29.5750 | 0.68021 | 28.059 | 31.091 |

Std Error uses a pooled estimate of error variance

##### Means and Std Deviations

| Level | Number | Mean | Std Dev | Std Err Mean | Lower 95% | Upper 95% |
|---|---|---|---|---|---|---|
| 0 | 4 | 29.8000 | 1.73013 | 0.86506 | 27.047 | 32.553 |
| 1 | 8 | 29.5750 | 2.00125 | 0.70755 | 27.902 | 31.248 |

##### t Test

1-0

Assuming unequal variances

| | | | |
|---|---|---|---|
| Difference | -0.2250 | t Ratio | -0.20133 |
| Std Err Dif | 1.1176 | DF | 7.01168 |
| Upper CL Dif | 2.4167 | Prob > \|t\| | 0.8462 |
| Lower CL Dif | -2.8667 | Prob > t | 0.5769 |
| Confidence | 0.95 | Prob < t | 0.4231 |

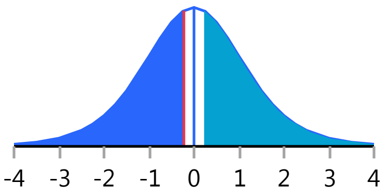


Missing Rows 2

### Fit Group

#### Oneway Analysis of LBP By Weight IBS-subtype=1

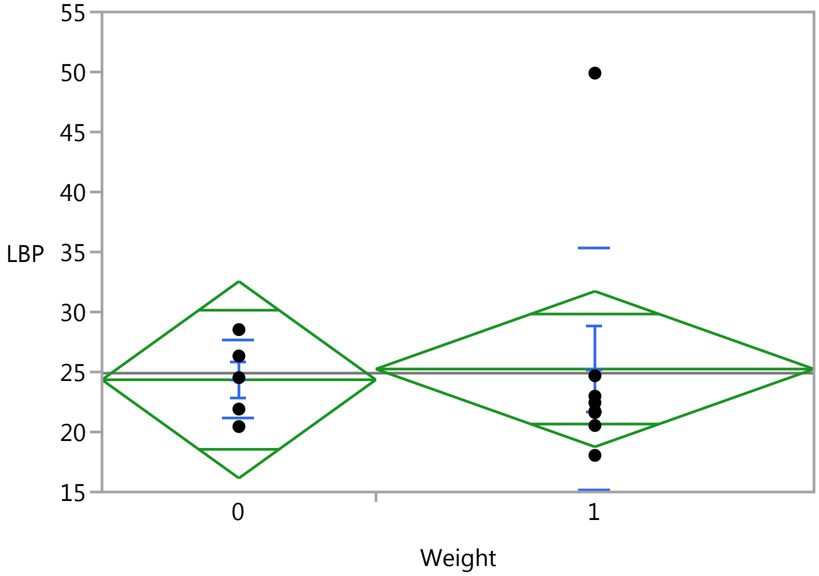


#### Oneway Anova

##### Summary of Fit

| | |
|---|---|
| Rsquare | 0.003215 |
| Adj Rsquare | -0.0874 |
| Root Mean Square Error | 8.330332 |
| Mean of Response | 24.90269 |
| Observations (or Sum Wgts) | 13 |

##### t Test

1-0

Assuming equal variances

| | | | |
|---|---|---|---|
| Difference | 0.895 | t Ratio | 0.188365 |
| Std Err Dif | 4.749 | DF | 11 |
| Upper CL Dif | 11.347 | Prob > \|t\| | 0.8540 |
| Lower CL Dif | -9.558 | Prob > t | 0.4270 |
| Confidence | 0.95 | Prob < t | 0.5730 |

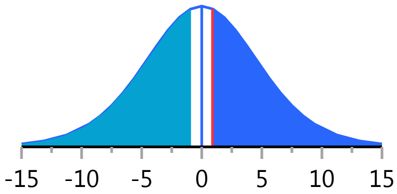


##### Analysis of Variance

| Source | DF | Sum of Squares | Mean Square | F Ratio | Prob > F |
|---|---|---|---|---|---|
| Weight | 1 | 2.46221 | 2.4622 | 0.0355 | 0.8540 |
| Error | 11 | 763.33874 | 69.3944 | | |
| C. Total | 12 | 765.80096 | | | |

##### Means for Oneway Anova

| Level | Number | Mean | Std Error | Lower 95% | Upper 95% |
|---|---|---|---|---|---|
| 0 | 5 | 24.3522 | 3.7254 | 16.153 | 32.552 |
| 1 | 8 | 25.2468 | 2.9452 | 18.764 | 31.729 |

Std Error uses a pooled estimate of error variance

#### Fit Group

##### Oneway Analysis of LBP By Weight IBS-subtype=1

###### Means and Std Deviations

| Level | Number | Mean | Std Dev | Std Err Mean | Lower 95% | Upper 95% |
|---|---|---|---|---|---|---|
| 0 | 5 | 24.3522 | 3.2632 | 1.4594 | 20.300 | 28.404 |
| 1 | 8 | 25.2468 | 10.1471 | 3.5875 | 16.764 | 33.730 |

###### t Test

1-0

Assuming unequal variances

| | | | |
|---|---|---|---|
| Difference | 0.8945 | t Ratio | 0.230971 |
| Std Err Dif | 3.8730 | DF | 9.07351 |
| Upper CL Dif | 9.6451 | Prob > \|t\| | 0.8225 |
| Lower CL Dif | -7.8560 | Prob > t | 0.4112 |
| Confidence | 0.95 | Prob < t | 0.5888 |

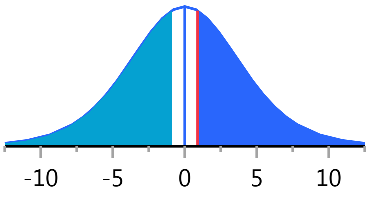


Missing Rows 1

##### Oneway Analysis of Lymphocytes_PCT By Weight IBS-subtype=1

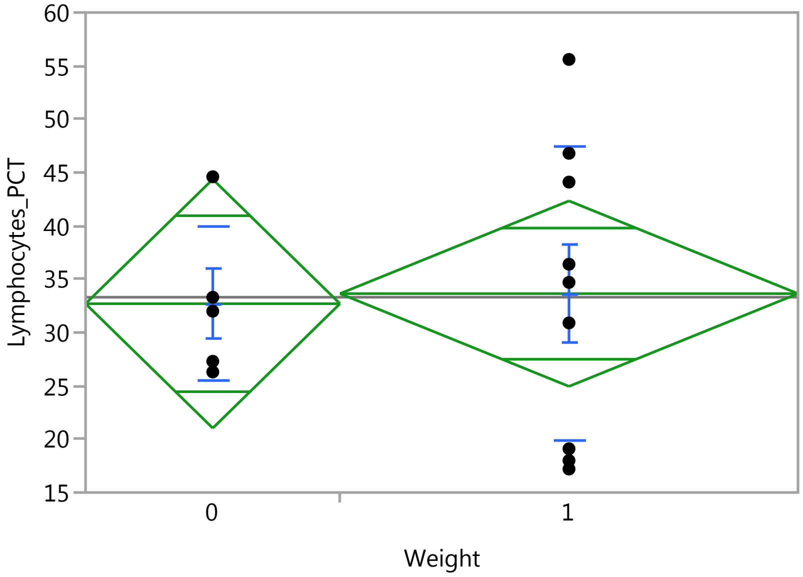


###### Oneway Anova

###### Summary of Fit

| | |
|---|---|
| Rsquare | 0.001664 |
| Adj Rsquare | -0.08153 |
| Root Mean Square Error | 11.97206 |
| Mean of Response | 33.30714 |
| Observations (or Sum Wgts) | 14 |

###### t Test

1-0

### Fit Group

#### Oneway Analysis of Lymphocytes_PCT By Weight IBS-subtype=1

##### Oneway Anova

###### t Test

Assuming equal variances

| | | | |
|---|---|---|---|
| Difference | 0.944 | t Ratio | 0.141433 |
| Std Err Dif | 6.678 | DF | 12 |
| Upper CL Dif | 15.494 | Prob > \|t\| | 0.8899 |
| Lower CL Dif | -13.605 | Prob > t | 0.4449 |
| Confidence | 0.95 | Prob < t | 0.5551 |

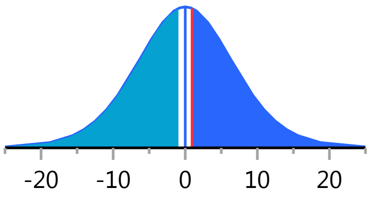


###### Analysis of Variance

| Source | DF | Sum of Squares | Mean Square | F Ratio | Prob > F |
|---|---|---|---|---|---|
| Weight | 1 | 2.8671 | 2.867 | 0.0200 | 0.8899 |
| Error | 12 | 1719.9622 | 143.330 | | |
| C. Total | 13 | 1722.8293 | | | |

###### Means for Oneway Anova

| Level | Number | Mean | Std Error | Lower 95% | Upper 95% |
|---|---|---|---|---|---|
| 0 | 5 | 32.7000 | 5.3541 | 21.034 | 44.366 |
| 1 | 9 | 33.6444 | 3.9907 | 24.949 | 42.339 |

Std Error uses a pooled estimate of error variance

##### Means and Std Deviations

| Level | Number | Mean | Std Dev | Std Err Mean | Lower 95% | Upper 95% |
|---|---|---|---|---|---|---|
| 0 | 5 | 32.7000 | 7.2901 | 3.2602 | 23.648 | 41.752 |
| 1 | 9 | 33.6444 | 13.7267 | 4.5756 | 23.093 | 44.196 |

##### t Test

1-0

Assuming unequal variances

| | | | |
|---|---|---|---|
| Difference | 0.944 | t Ratio | 0.168103 |
| Std Err Dif | 5.618 | DF | 11.99938 |
| Upper CL Dif | 13.186 | Prob > \|t\| | 0.8693 |
| Lower CL Dif | -11.297 | Prob > t | 0.4347 |
| Confidence | 0.95 | Prob < t | 0.5653 |

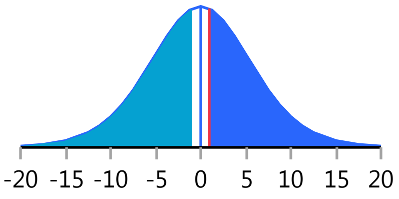


#### Oneway Analysis of Basophils By Weight IBS-subtype=1

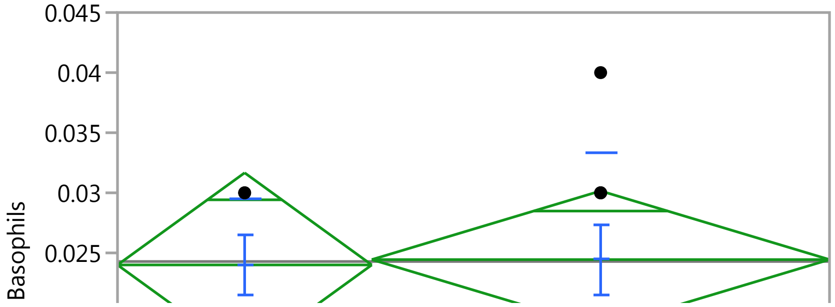

#### Fit Group

##### Oneway Analysis of Basophils By Weight IBS-subtype=1

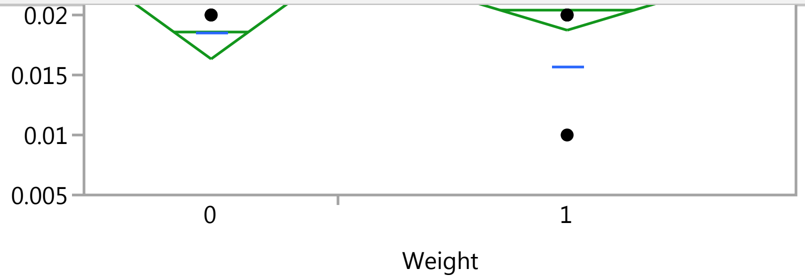


##### Oneway Anova

###### Summary of Fit

| | |
|---|---|
| Rsquare | 0.000855 |
| Adj Rsquare | -0.08241 |
| Root Mean Square Error | 0.007865 |
| Mean of Response | 0.024286 |
| Observations (or Sum Wgts) | 14 |

###### t Test

1-0

Assuming equal variances

| | | | |
|---|---|---|---|
| Difference | 0.00044 | t Ratio | 0.101317 |
| Std Err Dif | 0.00439 | DF | 12 |
| Upper CL Dif | 0.01000 | Prob > \|t\| | 0.9210 |
| Lower CL Dif | -0.00911 | Prob > t | 0.4605 |
| Confidence | 0.95 | Prob < t | 0.5395 |

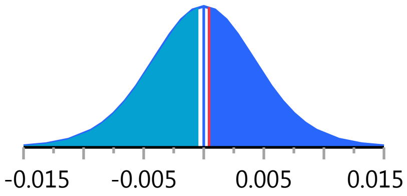


###### Analysis of Variance

| Source | DF | Sum of Squares | Mean Square | F Ratio | Prob > F |
|---|---|---|---|---|---|
| Weight | 1 | 0.00000063 | 6.349e-7 | 0.0103 | 0.9210 |
| Error | 12 | 0.00074222 | 0.000062 | | |
| C. Total | 13 | 0.00074286 | | | |

###### Means for Oneway Anova

| Level | Number | Mean | Std Error | Lower 95% | Upper 95% |
|---|---|---|---|---|---|
| 0 | 5 | 0.024000 | 0.00352 | 0.01634 | 0.03166 |
| 1 | 9 | 0.024444 | 0.00262 | 0.01873 | 0.03016 |

Std Error uses a pooled estimate of error variance

##### Means and Std Deviations

| Level | Number | Mean | Std Dev | Std Err Mean | Lower 95% | Upper 95% |
|---|---|---|---|---|---|---|
| 0 | 5 | 0.024000 | 0.005477 | 0.00245 | 0.01720 | 0.03080 |
| 1 | 9 | 0.024444 | 0.008819 | 0.00294 | 0.01767 | 0.03122 |

##### t Test

1-0

#### Fit Group

##### Oneway Analysis of Basophils By Weight IBS-subtype=1

###### t Test

Assuming unequal variances

| | | | |
|---|---|---|---|
| Difference | 0.00044 | t Ratio | 0.11615 |
| Std Err Dif | 0.00383 | DF | 11.6925 |
| Upper CL Dif | 0.00881 | Prob > \|t\| | 0.9095 |
| Lower CL Dif | -0.00792 | Prob > t | 0.4548 |
| Confidence | 0.95 | Prob < t | 0.5452 |

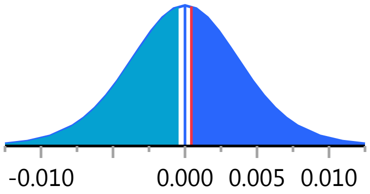


##### Oneway Analysis of Neutrophil_PCT By Weight IBS-subtype=1

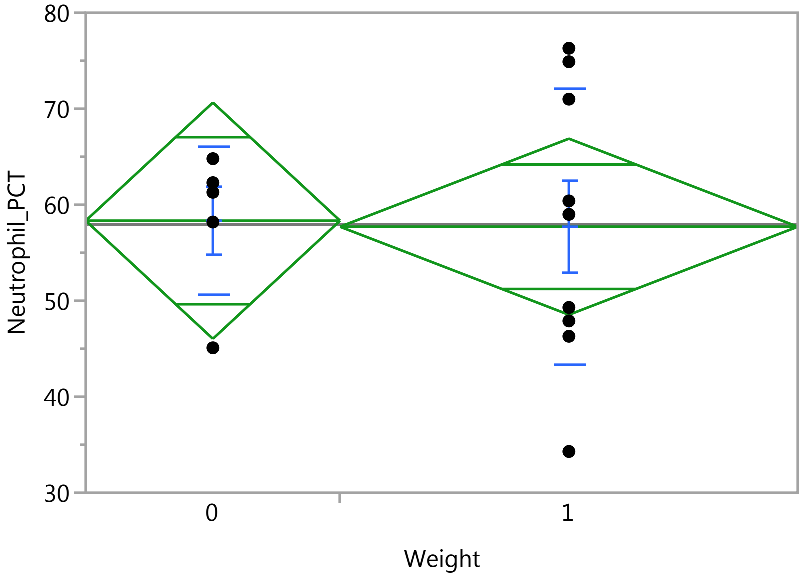


###### Oneway Anova

###### Summary of Fit

| | |
|---|---|
| Rsquare | 0.000664 |
| Adj Rsquare | -0.08261 |
| Root Mean Square Error | 12.62696 |
| Mean of Response | 57.93571 |
| Observations (or Sum Wgts) | 14 |

###### t Test

1-0

Assuming equal variances

| | | | |
|---|---|---|---|
| Difference | -0.629 | t Ratio | -0.08929 |
| Std Err Dif | 7.043 | DF | 12 |
| Upper CL Dif | 14.716 | Prob > \|t\| | 0.9303 |
| Lower CL Dif | -15.974 | Prob > t | 0.5348 |
| Confidence | 0.95 | Prob < t | 0.4652 |

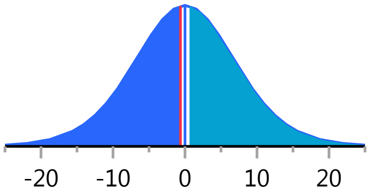

### Fit Group

#### Oneway Analysis of Neutrophil_PCT By Weight IBS-subtype=1

##### Oneway Anova

###### Analysis of Variance

| Source | DF | Sum of Squares | Mean Square | F Ratio | Prob > F |
|---|---|---|---|---|---|
| Weight | 1 | 1.2713 | 1.271 | 0.0080 | 0.9303 |
| Error | 12 | 1913.2809 | 159.440 | | |
| C. Total | 13 | 1914.5521 | | | |

###### Means for Oneway Anova

| Level | Number | Mean | Std Error | Lower 95% | Upper 95% |
|---|---|---|---|---|---|
| 0 | 5 | 58.3400 | 5.6469 | 46.036 | 70.644 |
| 1 | 9 | 57.7111 | 4.2090 | 48.541 | 66.882 |

Std Error uses a pooled estimate of error variance

##### Means and Std Deviations

| Level | Number | Mean | Std Dev | Std Err Mean | Lower 95% | Upper 95% |
|---|---|---|---|---|---|---|
| 0 | 5 | 58.3400 | 7.7700 | 3.4749 | 48.692 | 67.988 |
| 1 | 9 | 57.7111 | 14.4559 | 4.8186 | 46.599 | 68.823 |

##### t Test

1-0

Assuming unequal variances

| | | | |
|---|---|---|---|
| Difference | -0.629 | t Ratio | -0.10586 |
| Std Err Dif | 5.941 | DF | 11.99584 |
| Upper CL Dif | 12.316 | Prob > \|t\| | 0.9174 |
| Lower CL Dif | -13.573 | Prob > t | 0.5413 |
| Confidence | 0.95 | Prob < t | 0.4587 |

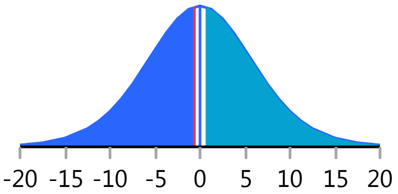


#### Oneway Analysis of RBC By Weight IBS-subtype=1

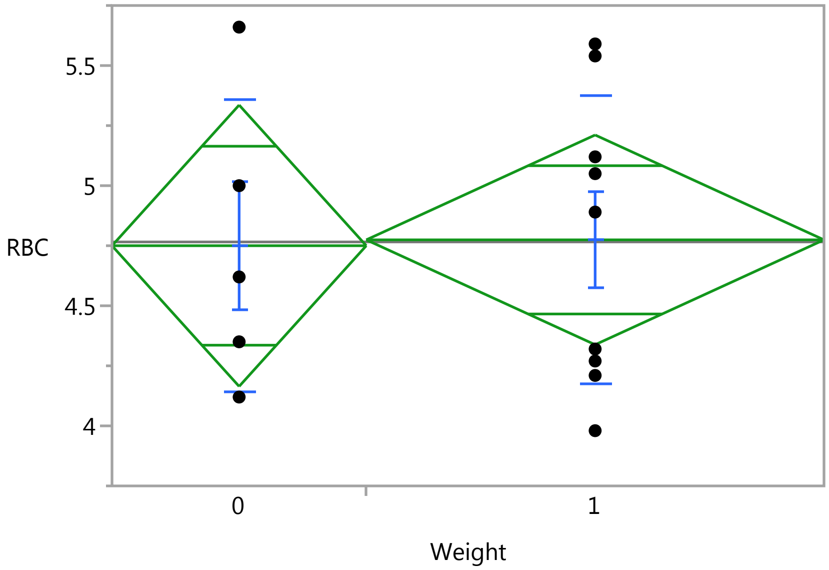

### Fit Group

#### Oneway Analysis of RBC By Weight IBS-subtype=1

Weight

##### Oneway Anova

###### Summary of Fit

| | |
|---|---|
| Rsquare | 0.000443 |
| Adj Rsquare | -0.08285 |
| Root Mean Square Error | 0.601042 |
| Mean of Response | 4.765714 |
| Observations (or Sum Wgts) | 14 |

###### t Test

1-0

Assuming equal variances

| | | | |
|---|---|---|---|
| Difference | 0.02444 | t Ratio | 0.072915 |
| Std Err Dif | 0.33525 | DF | 12 |
| Upper CL Dif | 0.75488 | Prob > \|t\| | 0.9431 |
| Lower CL Dif | -0.70599 | Prob > t | 0.4715 |
| Confidence | 0.95 | Prob < t | 0.5285 |

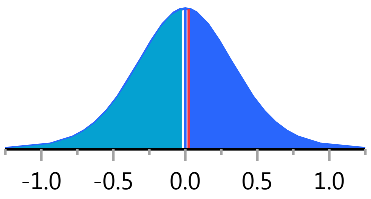


###### Analysis of Variance

| Source | DF | Sum of Squares | Mean Square | F Ratio | Prob > F |
|---|---|---|---|---|---|
| Weight | 1 | 0.0019206 | 0.001921 | 0.0053 | 0.9431 |
| Error | 12 | 4.3350222 | 0.361252 | | |
| C. Total | 13 | 4.3369429 | | | |

###### Means for Oneway Anova

| Level | Number | Mean | Std Error | Lower 95% | Upper 95% |
|---|---|---|---|---|---|
| 0 | 5 | 4.75000 | 0.26879 | 4.1643 | 5.3357 |
| 1 | 9 | 4.77444 | 0.20035 | 4.3379 | 5.2110 |

Std Error uses a pooled estimate of error variance

##### Means and Std Deviations

| Level | Number | Mean | Std Dev | Std Err Mean | Lower 95% | Upper 95% |
|---|---|---|---|---|---|---|
| 0 | 5 | 4.75000 | 0.605062 | 0.27059 | 3.9987 | 5.5013 |
| 1 | 9 | 4.77444 | 0.599022 | 0.19967 | 4.3140 | 5.2349 |

##### t Test

1-0

Assuming unequal variances

| | | | |
|---|---|---|---|
| Difference | 0.02444 | t Ratio | 0.072689 |
| Std Err Dif | 0.33629 | DF | 8.310176 |
| Upper CL Dif | 0.79492 | Prob > \|t\| | 0.9438 |
| Lower CL Dif | -0.74603 | Prob > t | 0.4719 |
| Confidence | 0.95 | Prob < t | 0.5281 |

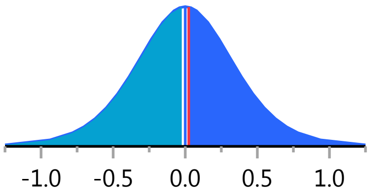

#### Fit Group

##### Oneway Analysis of SerumCortisol By Weight IBS-subtype=2

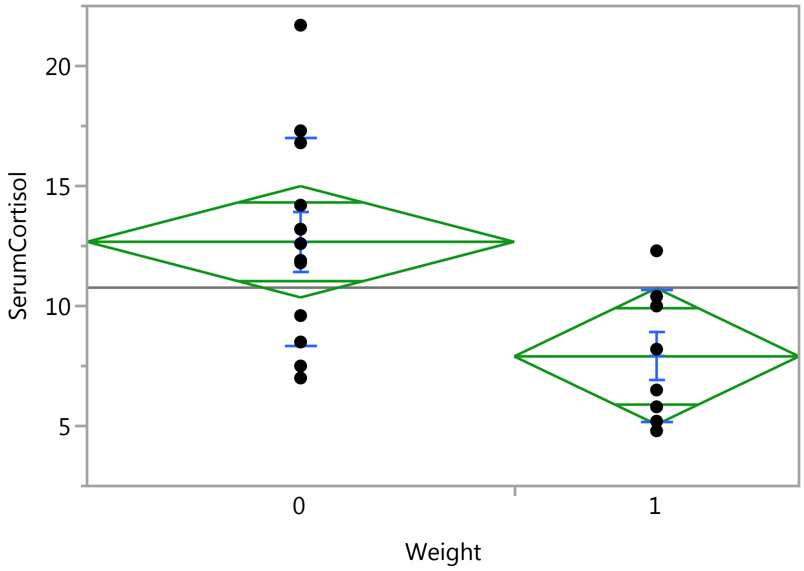


##### Oneway Anova

###### Summary of Fit

| | |
|---|---|
| Rsquare | 0.293787 |
| Adj Rsquare | 0.254553 |
| Root Mean Square Error | 3.823048 |
| Mean of Response | 10.765 |
| Observations (or Sum Wgts) | 20 |

###### t Test

1-0

Assuming equal variances

| | | | |
|---|---|---|---|
| Difference | -4.7750 | t Ratio | -2.73643 |
| Std Err Dif | 1.7450 | DF | 18 |
| Upper CL Dif | -1.1089 | Prob > \|t\| | 0.0136* |
| Lower CL Dif | -8.4411 | Prob > t | 0.9932 |
| Confidence | 0.95 | Prob < t | 0.0068* |

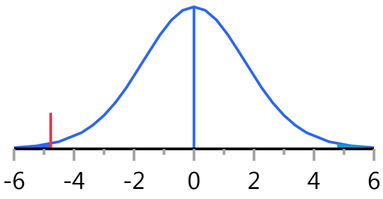


###### Analysis of Variance

| Source | DF | Sum of Squares | Mean Square | F Ratio | Prob > F |
|---|---|---|---|---|---|
| Weight | 1 | 109.44300 | 109.443 | 7.4880 | 0.0136* |
| Error | 18 | 263.08250 | 14.616 | | |
| C. Total | 19 | 372.52550 | | | |

###### Means for Oneway Anova

| Level | Number | Mean | Std Error | Lower 95% | Upper 95% |
|---|---|---|---|---|---|
| 0 | 12 | 12.6750 | 1.1036 | 10.356 | 14.994 |
| 1 | 8 | 7.9000 | 1.3517 | 5.060 | 10.740 |

Std Error uses a pooled estimate of error variance

### Fit Group

#### Oneway Analysis of SerumCortisol By Weight IBS-subtype=2

##### Means and Std Deviations

| Level | Number | Mean | Std Dev | Std Err Mean | Lower 95% | Upper 95% |
|---|---|---|---|---|---|---|
| 0 | 12 | 12.6750 | 4.36622 | 1.2604 | 9.9008 | 15.449 |
| 1 | 8 | 7.9000 | 2.76147 | 0.9763 | 5.5914 | 10.209 |

##### t Test

1-0

Assuming unequal variances

| | | | |
|---|---|---|---|
| Difference | -4.7750 | t Ratio | -2.995 |
| Std Err Dif | 1.5943 | DF | 17.98541 |
| Upper CL Dif | -1.4253 | Prob > \|t\| | 0.0078* |
| Lower CL Dif | -8.1247 | Prob > t | 0.9961 |
| Confidence | 0.95 | Prob < t | 0.0039* |

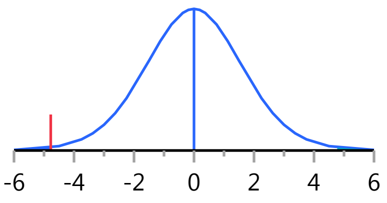


#### Oneway Analysis of ACTH By Weight IBS-subtype=2

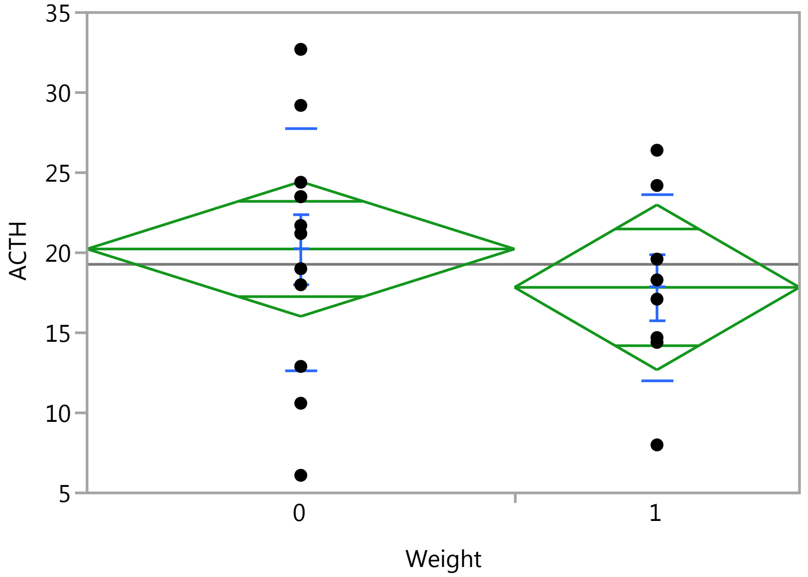


##### Oneway Anova

###### Summary of Fit

| | |
|---|---|
| Rsquare | 0.030816 |
| Adj Rsquare | -0.02303 |
| Root Mean Square Error | 6.938401 |
| Mean of Response | 19.275 |
| Observations (or Sum Wgts) | 20 |

###### t Test

1-0

### Fit Group

#### Oneway Analysis of ACTH By Weight IBS-subtype=2

##### Oneway Anova

###### t Test

Assuming equal variances

| | | | |
|---|---|---|---|
| Difference | -2.3958 | t Ratio | -0.75652 |
| Std Err Dif | 3.1669 | DF | 18 |
| Upper CL Dif | 4.2576 | Prob > \|t\| | 0.4591 |
| Lower CL Dif | -9.0493 | Prob > t | 0.7704 |
| Confidence | 0.95 | Prob < t | 0.2296 |

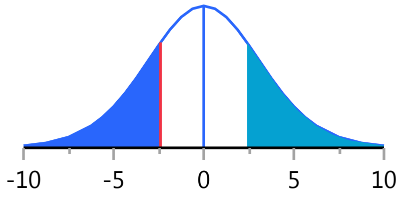


###### Analysis of Variance

| Source | DF | Sum of Squares | Mean Square | F Ratio | Prob > F |
|---|---|---|---|---|---|
| Weight | 1 | 27.55208 | 27.5521 | 0.5723 | 0.4591 |
| Error | 18 | 866.54542 | 48.1414 | | |
| C. Total | 19 | 894.09750 | | | |

###### Means for Oneway Anova

| Level | Number | Mean | Std Error | Lower 95% | Upper 95% |
|---|---|---|---|---|---|
| 0 | 12 | 20.2333 | 2.0029 | 16.025 | 24.441 |
| 1 | 8 | 17.8375 | 2.4531 | 12.684 | 22.991 |

Std Error uses a pooled estimate of error variance

##### Means and Std Deviations

| Level | Number | Mean | Std Dev | Std Err Mean | Lower 95% | Upper 95% |
|---|---|---|---|---|---|---|
| 0 | 12 | 20.2333 | 7.57056 | 2.1854 | 15.423 | 25.043 |
| 1 | 8 | 17.8375 | 5.80762 | 2.0533 | 12.982 | 22.693 |

##### t Test

1-0

Assuming unequal variances

| | | | |
|---|---|---|---|
| Difference | -2.3958 | t Ratio | -0.79896 |
| Std Err Dif | 2.9987 | DF | 17.52833 |
| Upper CL Dif | 3.9164 | Prob > \|t\| | 0.4350 |
| Lower CL Dif | -8.7080 | Prob > t | 0.7825 |
| Confidence | 0.95 | Prob < t | 0.2175 |

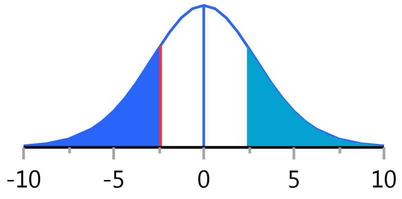


#### Oneway Analysis of CRP By Weight IBS-subtype=2

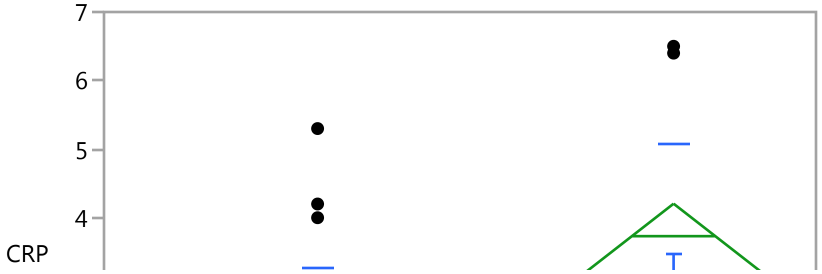

#### Fit Group

##### Oneway Analysis of CRP By Weight IBS-subtype=2

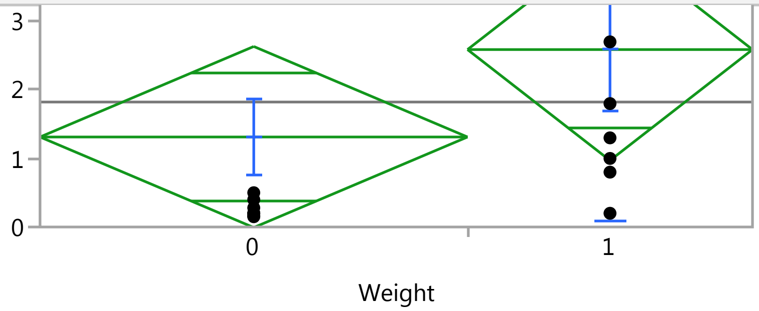


##### Oneway Anova

###### Summary of Fit

| | |
|---|---|
| Rsquare | 0.08382 |
| Adj Rsquare | 0.032921 |
| Root Mean Square Error | 2.176758 |
| Mean of Response | 1.8225 |
| Observations (or Sum Wgts) | 20 |

###### t Test

1-0

Assuming equal variances

| | | | |
|---|---|---|---|
| Difference | 1.2750 | t Ratio | 1.283278 |
| Std Err Dif | 0.9935 | DF | 18 |
| Upper CL Dif | 3.3624 | Prob > \|t\| | 0.2157 |
| Lower CL Dif | -0.8124 | Prob > t | 0.1078 |
| Confidence | 0.95 | Prob < t | 0.8922 |

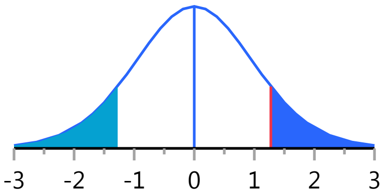


###### Analysis of Variance

| Source | DF | Sum of Squares | Mean Square | F Ratio | Prob > F |
|---|---|---|---|---|---|
| Weight | 1 | 7.803000 | 7.80300 | 1.6468 | 0.2157 |
| Error | 18 | 85.288975 | 4.73828 | | |
| C. Total | 19 | 93.091975 | | | |

###### Means for Oneway Anova

| Level | Number | Mean | Std Error | Lower 95% | Upper 95% |
|---|---|---|---|---|---|
| 0 | 12 | 1.31250 | 0.62838 | -0.0077 | 2.6327 |
| 1 | 8 | 2.58750 | 0.76960 | 0.9706 | 4.2044 |

Std Error uses a pooled estimate of error variance

##### Means and Std Deviations

| Level | Number | Mean | Std Dev | Std Err Mean | Lower 95% | Upper 95% |
|---|---|---|---|---|---|---|
| 0 | 12 | 1.31250 | 1.94796 | 0.56233 | 0.07482 | 2.5502 |
| 1 | 8 | 2.58750 | 2.49424 | 0.88185 | 0.50226 | 4.6727 |

##### t Test

1-0

### Fit Group

#### Oneway Analysis of CRP By Weight IBS-subtype=2

##### t Test

Assuming unequal variances

| | | | |
|---|---|---|---|
| Difference | 1.2750 | t Ratio | 1.219066 |
| Std Err Dif | 1.0459 | DF | 12.53159 |
| Upper CL Dif | 3.5431 | Prob > \|t\| | 0.2453 |
| Lower CL Dif | -0.9931 | Prob > t | 0.1226 |
| Confidence | 0.95 | Prob < t | 0.8774 |

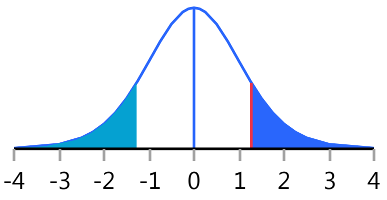


#### Oneway Analysis of ESR By Weight IBS-subtype=2

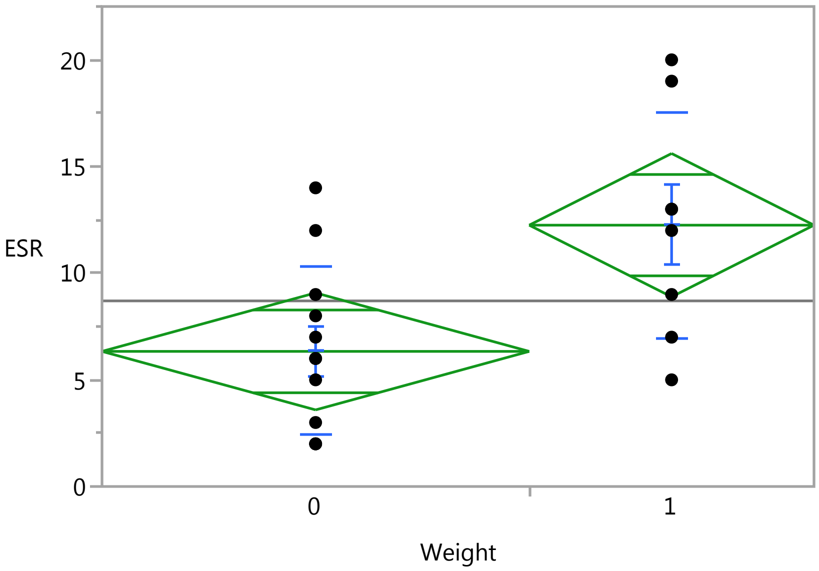


##### Oneway Anova

###### Summary of Fit

| | |
|---|---|
| Rsquare | 0.313378 |
| Adj Rsquare | 0.275232 |
| Root Mean Square Error | 4.522577 |
| Mean of Response | 8.7 |
| Observations (or Sum Wgts) | 20 |

###### t Test

1-0

Assuming equal variances

| | | | |
|---|---|---|---|
| Difference | 5.9167 | t Ratio | 2.866235 |
| Std Err Dif | 2.0643 | DF | 18 |
| Upper CL Dif | 10.2535 | Prob > \|t\| | 0.0103* |
| Lower CL Dif | 1.5798 | Prob > t | 0.0051* |
| Confidence | 0.95 | Prob < t | 0.9949 |

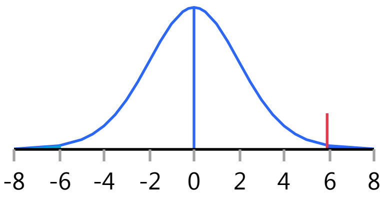

### Fit Group

#### Oneway Analysis of ESR By Weight IBS-subtype=2

##### Oneway Anova

###### Analysis of Variance

| Source | DF | Sum of Squares | Mean Square | F Ratio | Prob > F |
|---|---|---|---|---|---|
| Weight | 1 | 168.03333 | 168.033 | 8.2153 | 0.0103* |
| Error | 18 | 368.16667 | 20.454 | | |
| C. Total | 19 | 536.20000 | | | |

###### Means for Oneway Anova

| Level | Number | Mean | Std Error | Lower 95% | Upper 95% |
|---|---|---|---|---|---|
| 0 | 12 | 6.3333 | 1.3056 | 3.5905 | 9.076 |
| 1 | 8 | 12.2500 | 1.5990 | 8.8907 | 15.609 |

Std Error uses a pooled estimate of error variance

##### Means and Std Deviations

| Level | Number | Mean | Std Dev | Std Err Mean | Lower 95% | Upper 95% |
|---|---|---|---|---|---|---|
| 0 | 12 | 6.3333 | 3.93893 | 1.1371 | 3.8307 | 8.836 |
| 1 | 8 | 12.2500 | 5.31171 | 1.8780 | 7.8093 | 16.691 |

##### t Test

1-0

Assuming unequal variances

| | | | |
|---|---|---|---|
| Difference | 5.9167 | t Ratio | 2.695047 |
| Std Err Dif | 2.1954 | DF | 12.04322 |
| Upper CL Dif | 10.6981 | Prob > \|t\| | 0.0194* |
| Lower CL Dif | 1.1352 | Prob > t | 0.0097* |
| Confidence | 0.95 | Prob < t | 0.9903 |

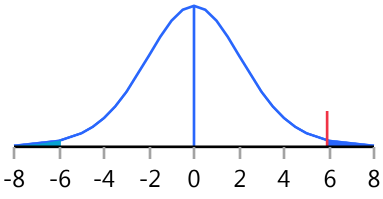


#### Oneway Analysis of LBP By Weight IBS-subtype=2

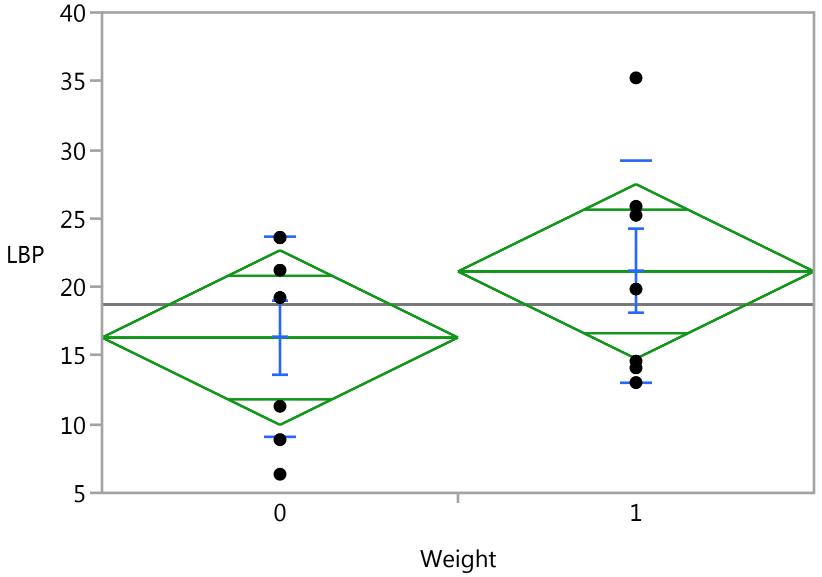

### Fit Group

#### Oneway Analysis of LBP By Weight IBS-subtype=2

Weight

##### Oneway Anova

###### Summary of Fit

| | |
|---|---|
| Rsquare | 0.101958 |
| Adj Rsquare | 0.027122 |
| Root Mean Square Error | 7.7223 |
| Mean of Response | 18.72686 |
| Observations (or Sum Wgts) | 14 |

###### t Test

1-0

Assuming equal variances

| | | | |
|---|---|---|---|
| Difference | 4.818 | t Ratio | 1.167224 |
| Std Err Dif | 4.128 | DF | 12 |
| Upper CL Dif | 13.812 | Prob > \|t\| | 0.2658 |
| Lower CL Dif | -4.176 | Prob > t | 0.1329 |
| Confidence | 0.95 | Prob < t | 0.8671 |

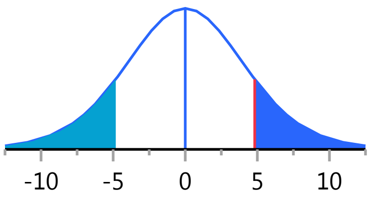


###### Analysis of Variance

| Source | DF | Sum of Squares | Mean Square | F Ratio | Prob > F |
|---|---|---|---|---|---|
| Weight | 1 | 81.24593 | 81.2459 | 1.3624 | 0.2658 |
| Error | 12 | 715.60709 | 59.6339 | | |
| C. Total | 13 | 796.85303 | | | |

###### Means for Oneway Anova

| Level | Number | Mean | Std Error | Lower 95% | Upper 95% |
|---|---|---|---|---|---|
| 0 | 7 | 16.3179 | 2.9188 | 9.958 | 22.677 |
| 1 | 7 | 21.1359 | 2.9188 | 14.776 | 27.495 |

Std Error uses a pooled estimate of error variance

##### Means and Std Deviations

| Level | Number | Mean | Std Dev | Std Err Mean | Lower 95% | Upper 95% |
|---|---|---|---|---|---|---|
| 0 | 7 | 16.3179 | 7.28301 | 2.7527 | 9.582 | 23.054 |
| 1 | 7 | 21.1359 | 8.13791 | 3.0758 | 13.610 | 28.662 |

##### t Test

1-0

Assuming unequal variances

| | | | |
|---|---|---|---|
| Difference | 4.818 | t Ratio | 1.167224 |
| Std Err Dif | 4.128 | DF | 11.85515 |
| Upper CL Dif | 13.824 | Prob > \|t\| | 0.2661 |
| Lower CL Dif | -4.188 | Prob > t | 0.1330 |
| Confidence | 0.95 | Prob < t | 0.8670 |

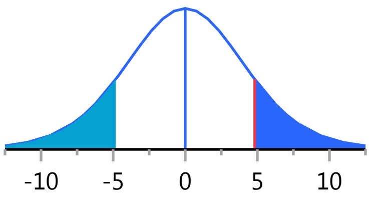


Missing Rows 6

#### Fit Group

##### Oneway Analysis of sCD14 By Weight IBS-subtype=2

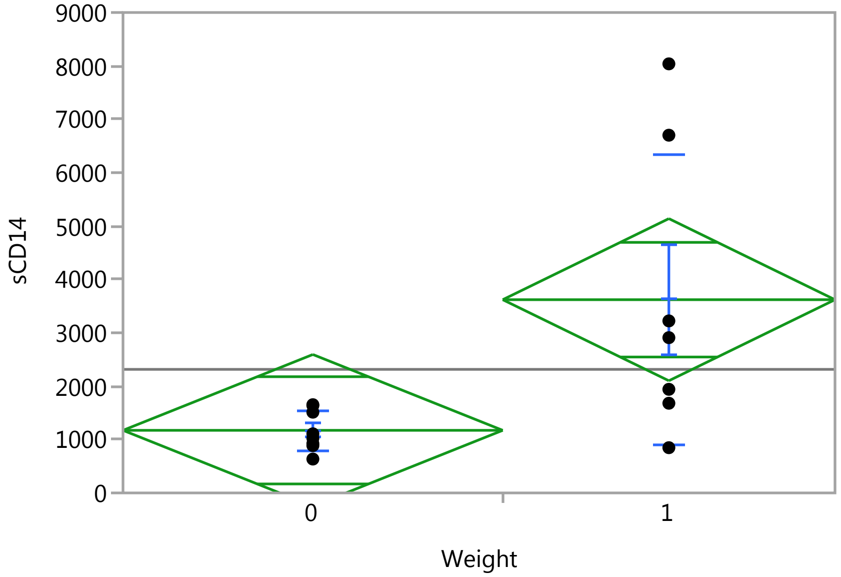


##### Oneway Anova

###### Summary of Fit

| | |
|---|---|
| Rsquare | 0.332104 |
| Adj Rsquare | 0.280727 |
| Root Mean Square Error | 1859.899 |
| Mean of Response | 2314.807 |
| Observations (or Sum Wgts) | 15 |

###### t Test

1-0

Assuming equal variances

| | | | |
|---|---|---|---|
| Difference | 2447.34 | t Ratio | 2.54246 |
| Std Err Dif | 962.59 | DF | 13 |
| Upper CL Dif | 4526.89 | Prob > \|t\| | 0.0245* |
| Lower CL Dif | 367.80 | Prob > t | 0.0123* |
| Confidence | 0.95 | Prob < t | 0.9877 |

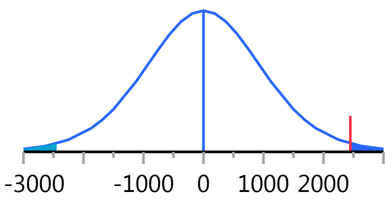


###### Analysis of Variance

| Source | DF | Sum of Squares | Mean Square | F Ratio | Prob > F |
|---|---|---|---|---|---|
| Weight | 1 | 22360784 | 22360784 | 6.4641 | 0.0245* |
| Error | 13 | 44969910 | 3459223.8 | | |
| C. Total | 14 | 67330694 | | | |

###### Means for Oneway Anova

| Level | Number | Mean | Std Error | Lower 95% | Upper 95% |
|---|---|---|---|---|---|
| 0 | 8 | 1172.71 | 657.57 | -248 | 2593.3 |
| 1 | 7 | 3620.06 | 702.98 | 2101 | 5138.7 |

Std Error uses a pooled estimate of error variance

### Fit Group

#### Oneway Analysis of sCD14 By Weight IBS-subtype=2

##### Means and Std Deviations

| Level | Number | Mean | Std Dev | Std Err Mean | Lower 95% | Upper 95% |
|---|---|---|---|---|---|---|
| 0 | 8 | 1172.71 | 382.01 | 135.1 | 853.3 | 1492.1 |
| 1 | 7 | 3620.06 | 2706.42 | 1022.9 | 1117.0 | 6123.1 |

##### t Test

1-0

Assuming unequal variances

| | | | |
|---|---|---|---|
| Difference | 2447.3 | t Ratio | 2.371895 |
| Std Err Dif | 1031.8 | DF | 6.209402 |
| Upper CL Dif | 4951.6 | Prob > \|t\| | 0.0540 |
| Lower CL Dif | -56.9 | Prob > t | 0.0270* |
| Confidence | 0.95 | Prob < t | 0.9730 |

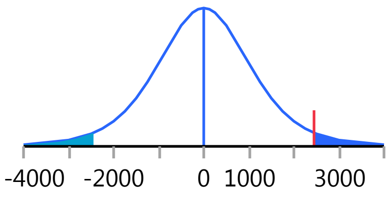


Missing Rows 5

#### Oneway Analysis of HCT By Weight IBS-subtype=2

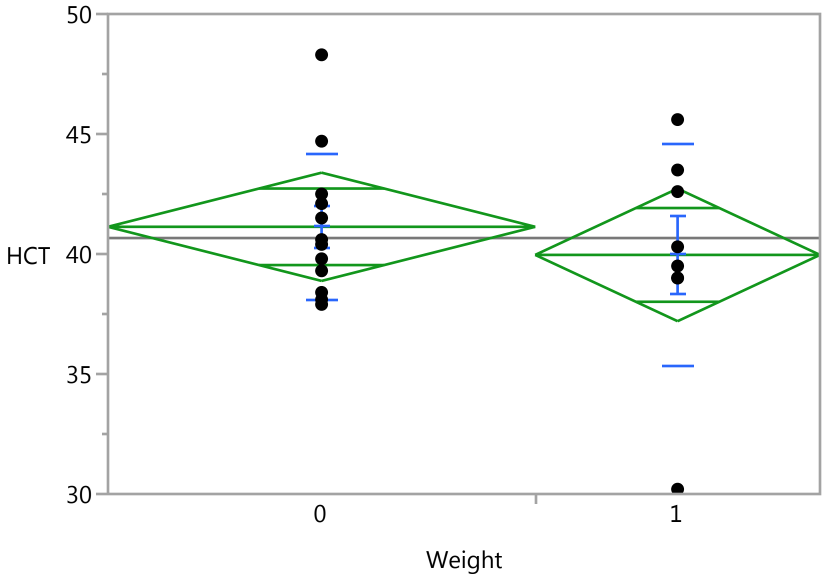


##### Oneway Anova

###### Summary of Fit

| | |
|---|---|
| Rsquare | 0.025761 |
| Adj Rsquare | -0.02836 |
| Root Mean Square Error | 3.718164 |
| Mean of Response | 40.665 |
| Observations (or Sum Wgts) | 20 |

###### t Test

1-0

### Fit Group

#### Oneway Analysis of HCT By Weight IBS-subtype=2

##### Oneway Anova

###### t Test

Assuming equal variances

| | | | |
|---|---|---|---|
| Difference | -1.1708 | t Ratio | -0.6899 |
| Std Err Dif | 1.6971 | DF | 18 |
| Upper CL Dif | 2.3946 | Prob > \|t\| | 0.4991 |
| Lower CL Dif | -4.7363 | Prob > t | 0.7505 |
| Confidence | 0.95 | Prob < t | 0.2495 |

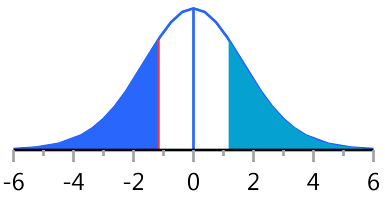


###### Analysis of Variance

| Source | DF | Sum of Squares | Mean Square | F Ratio | Prob > F |
|---|---|---|---|---|---|
| Weight | 1 | 6.58008 | 6.5801 | 0.4760 | 0.4991 |
| Error | 18 | 248.84542 | 13.8247 | | |
| C. Total | 19 | 255.42550 | | | |

###### Means for Oneway Anova

| Level | Number | Mean | Std Error | Lower 95% | Upper 95% |
|---|---|---|---|---|---|
| 0 | 12 | 41.1333 | 1.0733 | 38.878 | 43.388 |
| 1 | 8 | 39.9625 | 1.3146 | 37.201 | 42.724 |

Std Error uses a pooled estimate of error variance

##### Means and Std Deviations

| Level | Number | Mean | Std Dev | Std Err Mean | Lower 95% | Upper 95% |
|---|---|---|---|---|---|---|
| 0 | 12 | 41.1333 | 3.01672 | 0.8709 | 39.217 | 43.050 |
| 1 | 8 | 39.9625 | 4.60960 | 1.6297 | 36.109 | 43.816 |

##### t Test

1-0

Assuming unequal variances

| | | | |
|---|---|---|---|
| Difference | -1.1708 | t Ratio | -0.63363 |
| Std Err Dif | 1.8478 | DF | 10.99756 |
| Upper CL Dif | 2.8963 | Prob > \|t\| | 0.5393 |
| Lower CL Dif | -5.2380 | Prob > t | 0.7304 |
| Confidence | 0.95 | Prob < t | 0.2696 |

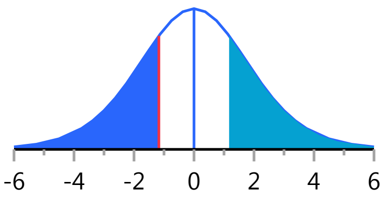


#### Oneway Analysis of MCH By Weight IBS-subtype=2

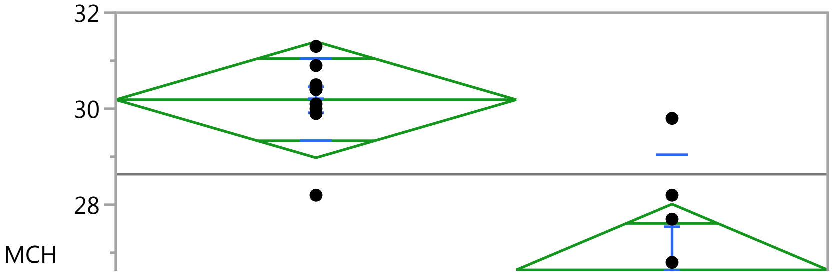

#### Fit Group

##### Oneway Analysis of MCH By Weight IBS-subtype=2

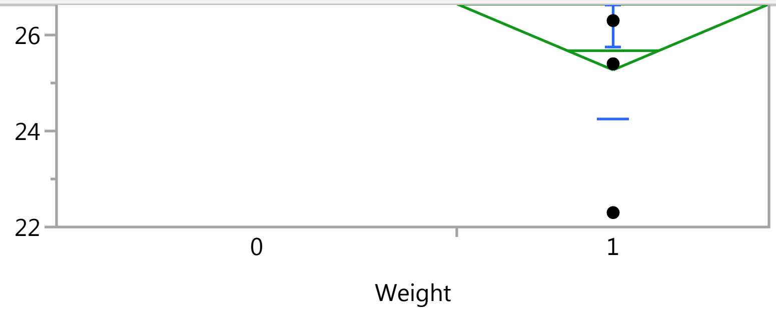


#### Oneway Anova

##### Summary of Fit

| | |
|---|---|
| Rsquare | 0.552722 |
| Adj Rsquare | 0.520774 |
| Root Mean Square Error | 1.691703 |
| Mean of Response | 28.6375 |
| Observations (or Sum Wgts) | 16 |

##### t Test

1-0

Assuming equal variances

| | | | |
|---|---|---|---|
| Difference | -3.5460 | t Ratio | -4.15938 |
| Std Err Dif | 0.8525 | DF | 14 |
| Upper CL Dif | -1.7175 | Prob > \|t\| | 0.0010* |
| Lower CL Dif | -5.3745 | Prob > t | 0.9995 |
| Confidence | 0.95 | Prob < t | 0.0005* |

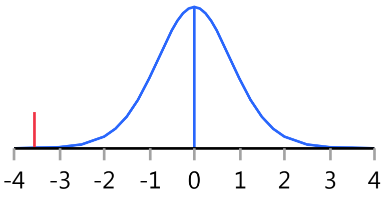


##### Analysis of Variance

| Source | DF | Sum of Squares | Mean Square | F Ratio | Prob > F |
|---|---|---|---|---|---|
| Weight | 1 | 49.511468 | 49.5115 | 17.3005 | 0.0010* |
| Error | 14 | 40.066032 | 2.8619 | | |
| C. Total | 15 | 89.577500 | | | |

##### Means for Oneway Anova

| Level | Number | Mean | Std Error | Lower 95% | Upper 95% |
|---|---|---|---|---|---|
| 0 | 9 | 30.1889 | 0.56390 | 28.979 | 31.398 |
| 1 | 7 | 26.6429 | 0.63940 | 25.271 | 28.014 |

Std Error uses a pooled estimate of error variance

#### Means and Std Deviations

| Level | Number | Mean | Std Dev | Std Err Mean | Lower 95% | Upper 95% |
|---|---|---|---|---|---|---|
| 0 | 9 | 30.1889 | 0.86667 | 0.28889 | 29.523 | 30.855 |
| 1 | 7 | 26.6429 | 2.38248 | 0.90049 | 24.439 | 28.846 |

#### t Test

1-0

#### Fit Group

##### Oneway Analysis of MCH By Weight IBS-subtype=2

###### t Test

Assuming unequal variances

| | | | |
|---|---|---|---|
| Difference | -3.5460 | t Ratio | -3.74965 |
| Std Err Dif | 0.9457 | DF | 7.241078 |
| Upper CL Dif | -1.3248 | Prob > \|t\| | 0.0067* |
| Lower CL Dif | -5.7672 | Prob > t | 0.9966 |
| Confidence | 0.95 | Prob < t | 0.0034* |

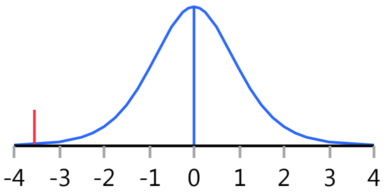


Missing Rows 4

##### Oneway Analysis of MPV By Weight IBS-subtype=2

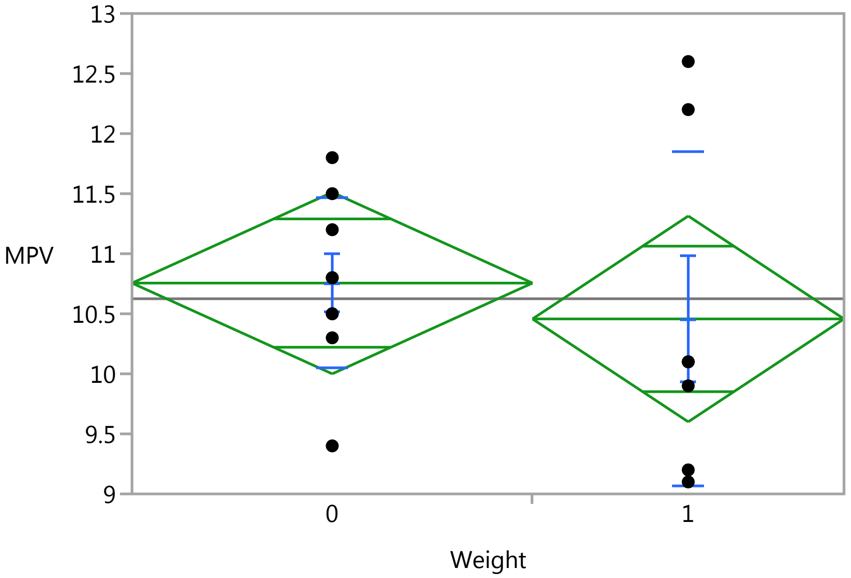


###### Oneway Anova

###### Summary of Fit

| | |
|---|---|
| Rsquare | 0.021928 |
| Adj Rsquare | -0.04793 |
| Root Mean Square Error | 1.056928 |
| Mean of Response | 10.625 |
| Observations (or Sum Wgts) | 16 |

###### t Test

1-0

Assuming equal variances

| | | | |
|---|---|---|---|
| Difference | -0.2984 | t Ratio | -0.56025 |
| Std Err Dif | 0.5326 | DF | 14 |
| Upper CL Dif | 0.8440 | Prob > \|t\| | 0.5842 |
| Lower CL Dif | -1.4408 | Prob > t | 0.7079 |
| Confidence | 0.95 | Prob < t | 0.2921 |

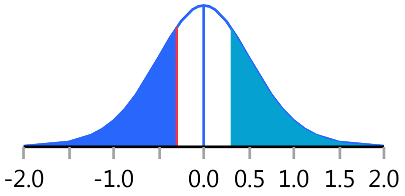

### Fit Group

#### Oneway Analysis of MPV By Weight IBS-subtype=2

##### Oneway Anova

###### Analysis of Variance

| Source | DF | Sum of Squares | Mean Square | F Ratio | Prob > F |
|---|---|---|---|---|---|
| Weight | 1 | 0.350635 | 0.35063 | 0.3139 | 0.5842 |
| Error | 14 | 15.639365 | 1.11710 | | |
| C. Total | 15 | 15.990000 | | | |

###### Means for Oneway Anova

| Level | Number | Mean | Std Error | Lower 95% | Upper 95% |
|---|---|---|---|---|---|
| 0 | 9 | 10.7556 | 0.35231 | 9.9999 | 11.511 |
| 1 | 7 | 10.4571 | 0.39948 | 9.6003 | 11.314 |

Std Error uses a pooled estimate of error variance

##### Means and Std Deviations

| Level | Number | Mean | Std Dev | Std Err Mean | Lower 95% | Upper 95% |
|---|---|---|---|---|---|---|
| 0 | 9 | 10.7556 | 0.70907 | 0.23636 | 10.211 | 11.301 |
| 1 | 7 | 10.4571 | 1.39147 | 0.52593 | 9.170 | 11.744 |

##### t Test

1-0

Assuming unequal variances

| | | | |
|---|---|---|---|
| Difference | -0.2984 | t Ratio | -0.51754 |
| Std Err Dif | 0.5766 | DF | 8.411046 |
| Upper CL Dif | 1.0200 | Prob > \|t\| | 0.6181 |
| Lower CL Dif | -1.6168 | Prob > t | 0.6909 |
| Confidence | 0.95 | Prob < t | 0.3091 |

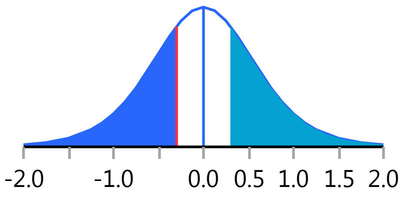


Missing Rows 4

#### Oneway Analysis of LDH By Weight IBS-subtype=2

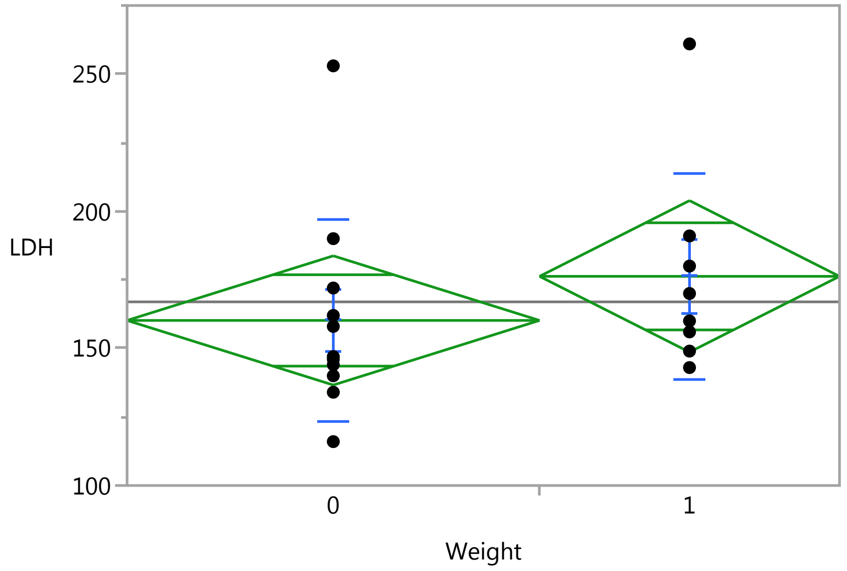

### Fit Group

#### Oneway Analysis of LDH By Weight IBS-subtype=2

Weight

##### Oneway Anova

###### Summary of Fit

| | |
|---|---|
| Rsquare | 0.048799 |
| Adj Rsquare | -0.00715 |
| Root Mean Square Error | 37.02871 |
| Mean of Response | 166.9474 |
| Observations (or Sum Wgts) | 19 |

###### t Test

1-0

Assuming equal variances

| | | | |
|---|---|---|---|
| Difference | 16.068 | t Ratio | 0.933883 |
| Std Err Dif | 17.206 | DF | 17 |
| Upper CL Dif | 52.369 | Prob > \|t\| | 0.3634 |
| Lower CL Dif | -20.233 | Prob > t | 0.1817 |
| Confidence | 0.95 | Prob < t | 0.8183 |

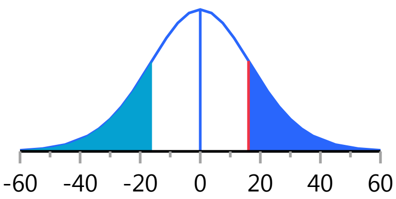


###### Analysis of Variance

| Source | DF | Sum of Squares | Mean Square | F Ratio | Prob > F |
|---|---|---|---|---|---|
| Weight | 1 | 1195.811 | 1195.81 | 0.8721 | 0.3634 |
| Error | 17 | 23309.136 | 1371.13 | | |
| C. Total | 18 | 24504.947 | | | |

###### Means for Oneway Anova

| Level | Number | Mean | Std Error | Lower 95% | Upper 95% |
|---|---|---|---|---|---|
| 0 | 11 | 160.182 | 11.165 | 136.63 | 183.74 |
| 1 | 8 | 176.250 | 13.092 | 148.63 | 203.87 |

Std Error uses a pooled estimate of error variance

##### Means and Std Deviations

| Level | Number | Mean | Std Dev | Std Err Mean | Lower 95% | Upper 95% |
|---|---|---|---|---|---|---|
| 0 | 11 | 160.182 | 36.5153 | 11.010 | 135.65 | 184.71 |
| 1 | 8 | 176.250 | 37.7501 | 13.347 | 144.69 | 207.81 |

##### t Test

1-0

Assuming unequal variances

| | | | |
|---|---|---|---|
| Difference | 16.068 | t Ratio | 0.928705 |
| Std Err Dif | 17.302 | DF | 14.92897 |
| Upper CL Dif | 52.961 | Prob > \|t\| | 0.3678 |
| Lower CL Dif | -20.825 | Prob > t | 0.1839 |
| Confidence | 0.95 | Prob < t | 0.8161 |

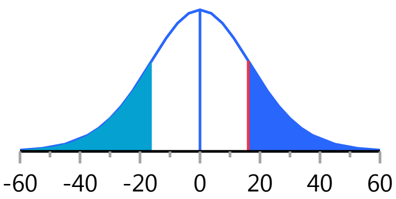


Missing Rows 1

#### Fit Group

##### Oneway Analysis of Monocytes By Weight IBS-subtype=2

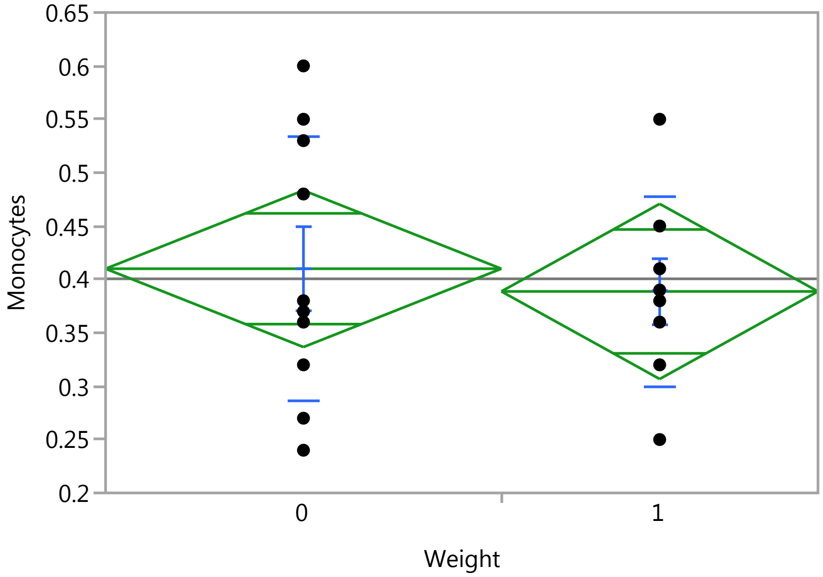


##### Oneway Anova

###### Summary of Fit

| | |
|---|---|
| Rsquare | 0.010361 |
| Adj Rsquare | -0.05149 |
| Root Mean Square Error | 0.109455 |
| Mean of Response | 0.400556 |
| Observations (or Sum Wgts) | 18 |

###### t Test

1-0

Assuming equal variances

| | | | |
|---|---|---|---|
| Difference | -0.02125 | t Ratio | -0.40929 |
| Std Err Dif | 0.05192 | DF | 16 |
| Upper CL Dif | 0.08881 | Prob > \|t\| | 0.6878 |
| Lower CL Dif | -0.13131 | Prob > t | 0.6561 |
| Confidence | 0.95 | Prob < t | 0.3439 |

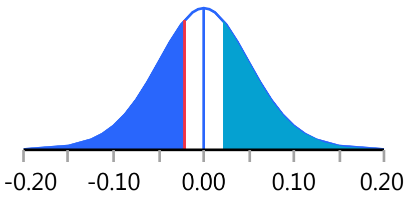


###### Analysis of Variance

| Source | DF | Sum of Squares | Mean Square | F Ratio | Prob > F |
|---|---|---|---|---|---|
| Weight | 1 | 0.00200694 | 0.002007 | 0.1675 | 0.6878 |
| Error | 16 | 0.19168750 | 0.011980 | | |
| C. Total | 17 | 0.19369444 | | | |

###### Means for Oneway Anova

| Level | Number | Mean | Std Error | Lower 95% | Upper 95% |
|---|---|---|---|---|---|
| 0 | 10 | 0.410000 | 0.03461 | 0.33662 | 0.48338 |
| 1 | 8 | 0.388750 | 0.03870 | 0.30671 | 0.47079 |

Std Error uses a pooled estimate of error variance

### Fit Group

#### Oneway Analysis of Monocytes By Weight IBS-subtype=2

##### Means and Std Deviations

| Level | Number | Mean | Std Dev | Std Err Mean | Lower 95% | Upper 95% |
|---|---|---|---|---|---|---|
| 0 | 10 | 0.410000 | 0.123198 | 0.03896 | 0.32187 | 0.49813 |
| 1 | 8 | 0.388750 | 0.088711 | 0.03136 | 0.31459 | 0.46291 |

##### t Test

1-0

Assuming unequal variances

| | | | |
|---|---|---|---|
| Difference | -0.02125 | t Ratio | -0.42487 |
| Std Err Dif | 0.05001 | DF | 15.87369 |
| Upper CL Dif | 0.08485 | Prob > \|t\| | 0.6766 |
| Lower CL Dif | -0.12735 | Prob > t | 0.6617 |
| Confidence | 0.95 | Prob < t | 0.3383 |

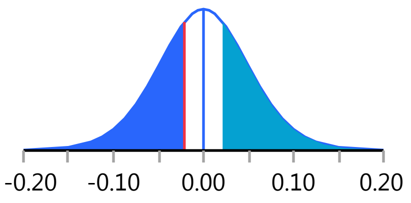


Missing Rows 2

#### Oneway Analysis of Monocytes_PCT By Weight IBS-subtype=2

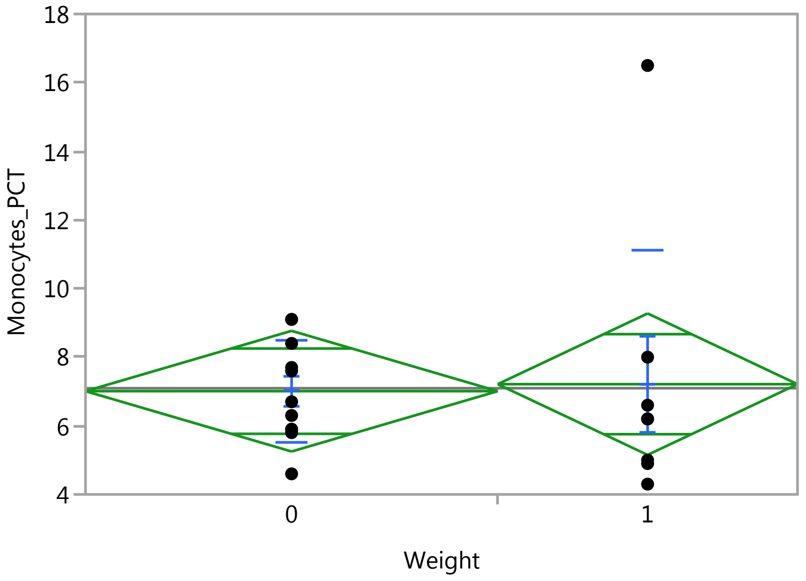


##### Oneway Anova

###### Summary of Fit

| | |
|---|---|
| Rsquare | 0.001474 |
| Adj Rsquare | -0.05726 |
| Root Mean Square Error | 2.763819 |
| Mean of Response | 7.094737 |
| Observations (or Sum Wgts) | 19 |

###### t Test

1-0

#### Fit Group

##### Oneway Analysis of Monocytes_PCT By Weight IBS-subtype=2

###### Oneway Anova

###### t Test

Assuming equal variances

| | | | |
|---|---|---|---|
| Difference | 0.2034 | t Ratio | 0.158389 |
| Std Err Dif | 1.2842 | DF | 17 |
| Upper CL Dif | 2.9129 | Prob > \|t\| | 0.8760 |
| Lower CL Dif | -2.5061 | Prob > t | 0.4380 |
| Confidence | 0.95 | Prob < t | 0.5620 |

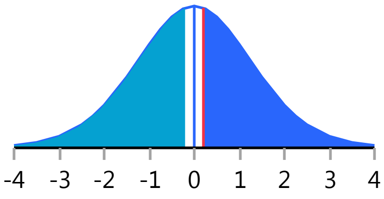


###### Analysis of Variance

| Source | DF | Sum of Squares | Mean Square | F Ratio | Prob > F |
|---|---|---|---|---|---|
| Weight | 1 | 0.19163 | 0.19163 | 0.0251 | 0.8760 |
| Error | 17 | 129.85784 | 7.63870 | | |
| C. Total | 18 | 130.04947 | | | |

###### Means for Oneway Anova

| Level | Number | Mean | Std Error | Lower 95% | Upper 95% |
|---|---|---|---|---|---|
| 0 | 11 | 7.00909 | 0.83332 | 5.2509 | 8.7672 |
| 1 | 8 | 7.21250 | 0.97716 | 5.1509 | 9.2741 |

Std Error uses a pooled estimate of error variance

###### Means and Std Deviations

| Level | Number | Mean | Std Dev | Std Err Mean | Lower 95% | Upper 95% |
|---|---|---|---|---|---|---|
| 0 | 11 | 7.00909 | 1.47747 | 0.4455 | 6.0165 | 8.002 |
| 1 | 8 | 7.21250 | 3.92844 | 1.3889 | 3.9282 | 10.497 |

###### t Test

1-0

Assuming unequal variances

| | | | |
|---|---|---|---|
| Difference | 0.2034 | t Ratio | 0.139454 |
| Std Err Dif | 1.4586 | DF | 8.451659 |
| Upper CL Dif | 3.5359 | Prob > \|t\| | 0.8924 |
| Lower CL Dif | -3.1291 | Prob > t | 0.4462 |
| Confidence | 0.95 | Prob < t | 0.5538 |

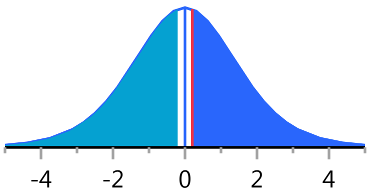


Missing Rows 1

##### Oneway Analysis of Basophil_PCT By Weight IBS-subtype=2

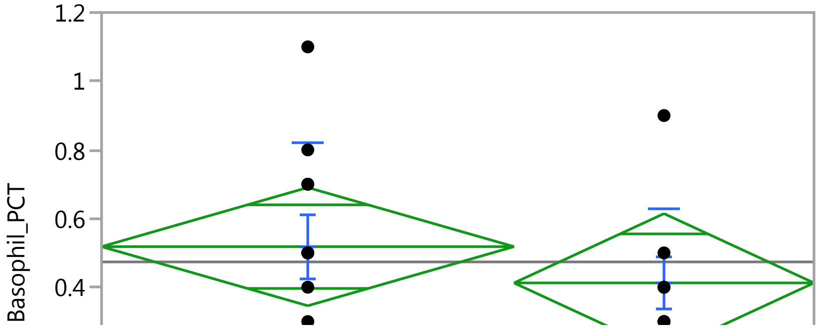

#### Fit Group

##### Oneway Analysis of Basophil_PCT By Weight IBS-subtype=2

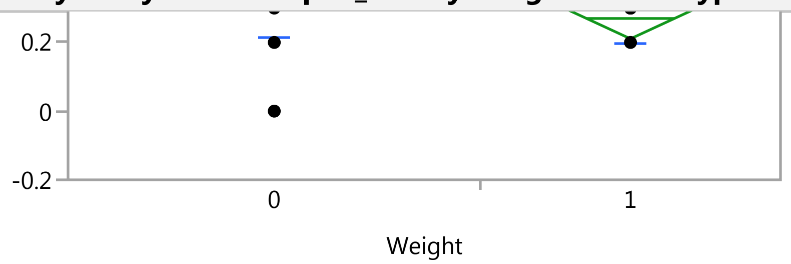


##### Oneway Anova

###### Summary of Fit

| | |
|---|---|
| Rsquare | 0.039888 |
| Adj Rsquare | -0.01659 |
| Root Mean Square Error | 0.270633 |
| Mean of Response | 0.473684 |
| Observations (or Sum Wgts) | 19 |

###### t Test

1-0

Assuming equal variances

| | | | |
|---|---|---|---|
| Difference | -0.10568 | t Ratio | -0.8404 |
| Std Err Dif | 0.12575 | DF | 17 |
| Upper CL Dif | 0.15963 | Prob > \|t\| | 0.4123 |
| Lower CL Dif | -0.37100 | Prob > t | 0.7938 |
| Confidence | 0.95 | Prob < t | 0.2062 |

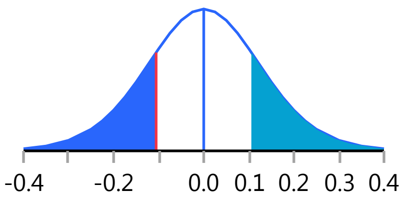


###### Analysis of Variance

| Source | DF | Sum of Squares | Mean Square | F Ratio | Prob > F |
|---|---|---|---|---|---|
| Weight | 1 | 0.0517285 | 0.051728 | 0.7063 | 0.4123 |
| Error | 17 | 1.2451136 | 0.073242 | | |
| C. Total | 18 | 1.2968421 | | | |

###### Means for Oneway Anova

| Level | Number | Mean | Std Error | Lower 95% | Upper 95% |
|---|---|---|---|---|---|
| 0 | 11 | 0.518182 | 0.08160 | 0.34602 | 0.69034 |
| 1 | 8 | 0.412500 | 0.09568 | 0.21063 | 0.61437 |

Std Error uses a pooled estimate of error variance

##### Means and Std Deviations

| Level | Number | Mean | Std Dev | Std Err Mean | Lower 95% | Upper 95% |
|---|---|---|---|---|---|---|
| 0 | 11 | 0.518182 | 0.302715 | 0.09127 | 0.31482 | 0.72155 |
| 1 | 8 | 0.412500 | 0.216712 | 0.07662 | 0.23132 | 0.59368 |

##### t Test

1-0

#### Fit Group

##### Oneway Analysis of Basophil_PCT By Weight IBS-subtype=2

###### t Test

Assuming unequal variances

| | | | |
|---|---|---|---|
| Difference | -0.10568 | t Ratio | -0.88683 |
| Std Err Dif | 0.11917 | DF | 16.99982 |
| Upper CL Dif | 0.14574 | Prob > \|t\| | 0.3875 |
| Lower CL Dif | -0.35711 | Prob > t | 0.8062 |
| Confidence | 0.95 | Prob < t | 0.1938 |

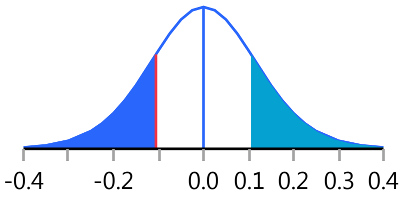


Missing Rows 1

##### Oneway Analysis of Basophils By Weight IBS-subtype=2

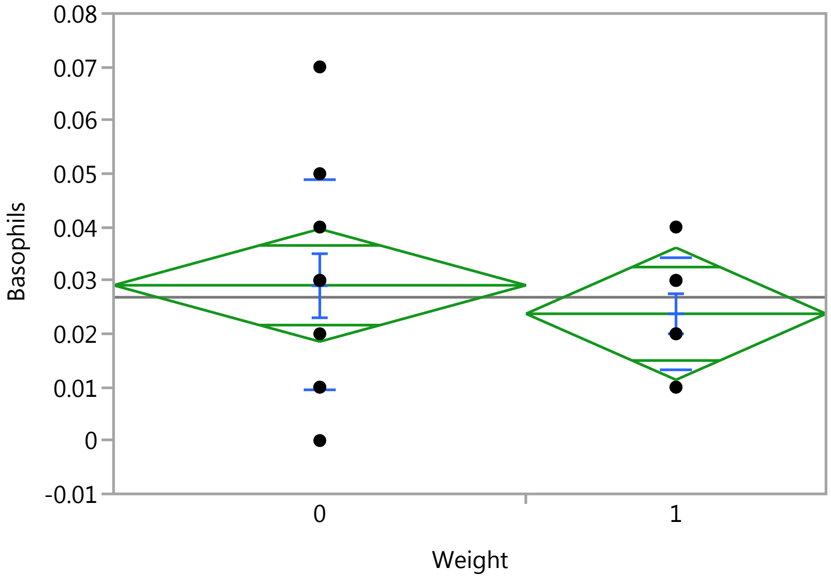


###### Oneway Anova

###### Summary of Fit

| | |
|---|---|
| Rsquare | 0.027464 |
| Adj Rsquare | -0.02974 |
| Root Mean Square Error | 0.016589 |
| Mean of Response | 0.026842 |
| Observations (or Sum Wgts) | 19 |

###### t Test

1-0

Assuming equal variances

| | | | |
|---|---|---|---|
| Difference | -0.00534 | t Ratio | -0.69288 |
| Std Err Dif | 0.00771 | DF | 17 |
| Upper CL Dif | 0.01092 | Prob > \|t\| | 0.4977 |
| Lower CL Dif | -0.02160 | Prob > t | 0.7511 |
| Confidence | 0.95 | Prob < t | 0.2489 |

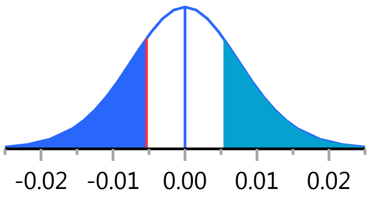

### Fit Group

#### Oneway Analysis of Basophils By Weight IBS-subtype=2

##### Oneway Anova

###### Analysis of Variance

| Source | DF | Sum of Squares | Mean Square | F Ratio | Prob > F |
|---|---|---|---|---|---|
| Weight | 1 | 0.00013212 | 0.000132 | 0.4801 | 0.4977 |
| Error | 17 | 0.00467841 | 0.000275 | | |
| C. Total | 18 | 0.00481053 | | | |

###### Means for Oneway Anova

| Level | Number | Mean | Std Error | Lower 95% | Upper 95% |
|---|---|---|---|---|---|
| 0 | 11 | 0.029091 | 0.00500 | 0.01854 | 0.03964 |
| 1 | 8 | 0.023750 | 0.00587 | 0.01138 | 0.03612 |

Std Error uses a pooled estimate of error variance

##### Means and Std Deviations

| Level | Number | Mean | Std Dev | Std Err Mean | Lower 95% | Upper 95% |
|---|---|---|---|---|---|---|
| 0 | 11 | 0.029091 | 0.019725 | 0.00595 | 0.01584 | 0.04234 |
| 1 | 8 | 0.023750 | 0.010607 | 0.00375 | 0.01488 | 0.03262 |

##### t Test

1-0

Assuming unequal variances

| | | | |
|---|---|---|---|
| Difference | -0.00534 | t Ratio | -0.75963 |
| Std Err Dif | 0.00703 | DF | 15.93399 |
| Upper CL Dif | 0.00957 | Prob > \|t\| | 0.4586 |
| Lower CL Dif | -0.02025 | Prob > t | 0.7707 |
| Confidence | 0.95 | Prob < t | 0.2293 |

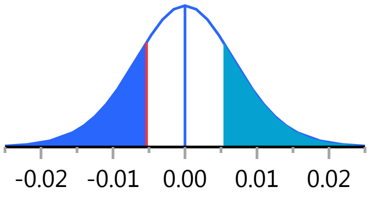


Missing Rows 1

#### Oneway Analysis of Eosinophil_PCT By Weight IBS-subtype=2

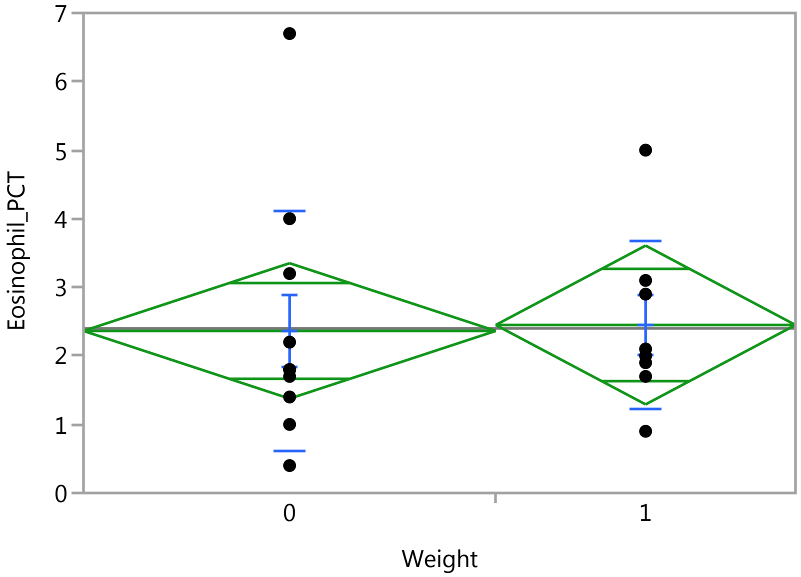

### Fit Group

#### Oneway Analysis of Eosinophil_PCT By Weight IBS-subtype=2

Weight

##### Oneway Anova

###### Summary of Fit

| | |
|---|---|
| Rsquare | 0.000843 |
| Adj Rsquare | -0.05793 |
| Root Mean Square Error | 1.552331 |
| Mean of Response | 2.4 |
| Observations (or Sum Wgts) | 19 |

###### t Test

1-0

Assuming equal variances

| | | | |
|---|---|---|---|
| Difference | 0.0864 | t Ratio | 0.119732 |
| Std Err Dif | 0.7213 | DF | 17 |
| Upper CL Dif | 1.6082 | Prob > \|t\| | 0.9061 |
| Lower CL Dif | -1.4355 | Prob > t | 0.4530 |
| Confidence | 0.95 | Prob < t | 0.5470 |

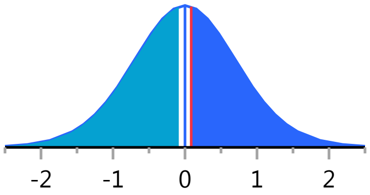


###### Analysis of Variance

| Source | DF | Sum of Squares | Mean Square | F Ratio | Prob > F |
|---|---|---|---|---|---|
| Weight | 1 | 0.034545 | 0.03455 | 0.0143 | 0.9061 |
| Error | 17 | 40.965455 | 2.40973 | | |
| C. Total | 18 | 41.000000 | | | |

###### Means for Oneway Anova

| Level | Number | Mean | Std Error | Lower 95% | Upper 95% |
|---|---|---|---|---|---|
| 0 | 11 | 2.36364 | 0.46805 | 1.3761 | 3.3511 |
| 1 | 8 | 2.45000 | 0.54883 | 1.2921 | 3.6079 |

Std Error uses a pooled estimate of error variance

##### Means and Std Deviations

| Level | Number | Mean | Std Dev | Std Err Mean | Lower 95% | Upper 95% |
|---|---|---|---|---|---|---|
| 0 | 11 | 2.36364 | 1.73912 | 0.52437 | 1.1953 | 3.5320 |
| 1 | 8 | 2.45000 | 1.23751 | 0.43753 | 1.4154 | 3.4846 |

##### t Test

1-0

Assuming unequal variances

| | | | |
|---|---|---|---|
| Difference | 0.0864 | t Ratio | 0.126461 |
| Std Err Dif | 0.6829 | DF | 16.99988 |
| Upper CL Dif | 1.5272 | Prob > \|t\| | 0.9009 |
| Lower CL Dif | -1.3545 | Prob > t | 0.4504 |
| Confidence | 0.95 | Prob < t | 0.5496 |

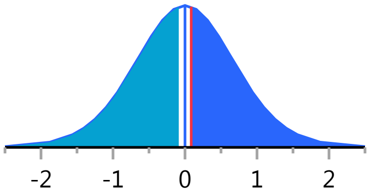


Missing Rows 1

#### Fit Group

##### Oneway Analysis of Eosinophils By Weight IBS-subtype=2

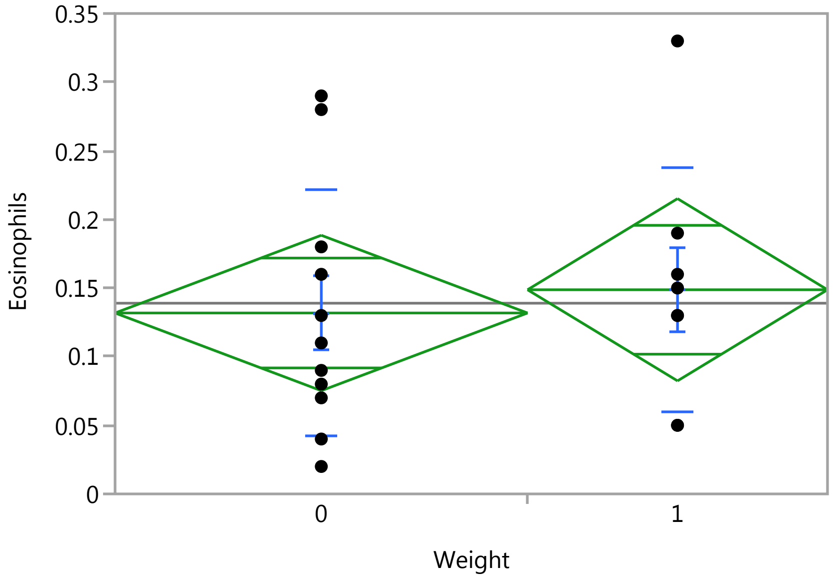


##### Oneway Anova

###### Summary of Fit

| | |
|---|---|
| Rsquare | 0.009765 |
| Adj Rsquare | -0.04848 |
| Root Mean Square Error | 0.088998 |
| Mean of Response | 0.138947 |
| Observations (or Sum Wgts) | 19 |

###### t Test

1-0

Assuming equal variances

| | | | |
|---|---|---|---|
| Difference | 0.01693 | t Ratio | 0.409437 |
| Std Err Dif | 0.04135 | DF | 17 |
| Upper CL Dif | 0.10418 | Prob > \|t\| | 0.6873 |
| Lower CL Dif | -0.07032 | Prob > t | 0.3437 |
| Confidence | 0.95 | Prob < t | 0.6563 |

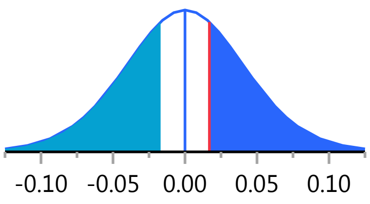


###### Analysis of Variance

| Source | DF | Sum of Squares | Mean Square | F Ratio | Prob > F |
|---|---|---|---|---|---|
| Weight | 1 | 0.00132781 | 0.001328 | 0.1676 | 0.6873 |
| Error | 17 | 0.13465114 | 0.007921 | | |
| C. Total | 18 | 0.13597895 | | | |

###### Means for Oneway Anova

| Level | Number | Mean | Std Error | Lower 95% | Upper 95% |
|---|---|---|---|---|---|
| 0 | 11 | 0.131818 | 0.02683 | 0.07520 | 0.18843 |
| 1 | 8 | 0.148750 | 0.03147 | 0.08236 | 0.21514 |

Std Error uses a pooled estimate of error variance

### Fit Group

#### Oneway Analysis of Eosinophils By Weight IBS-subtype=2

##### Means and Std Deviations

| Level | Number | Mean | Std Dev | Std Err Mean | Lower 95% | Upper 95% |
|---|---|---|---|---|---|---|
| 0 | 11 | 0.131818 | 0.089310 | 0.02693 | 0.07182 | 0.19182 |
| 1 | 8 | 0.148750 | 0.088550 | 0.03131 | 0.07472 | 0.22278 |

##### t Test

1-0

Assuming unequal variances

| | | | |
|---|---|---|---|
| Difference | 0.01693 | t Ratio | 0.410023 |
| Std Err Dif | 0.04129 | DF | 15.31944 |
| Upper CL Dif | 0.10479 | Prob > \|t\| | 0.6875 |
| Lower CL Dif | -0.07093 | Prob > t | 0.3437 |
| Confidence | 0.95 | Prob < t | 0.6563 |

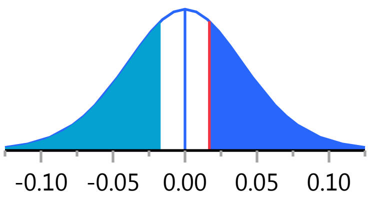


Missing Rows 1

#### Oneway Analysis of Lymphocytes_PCT By Weight IBS-subtype=2

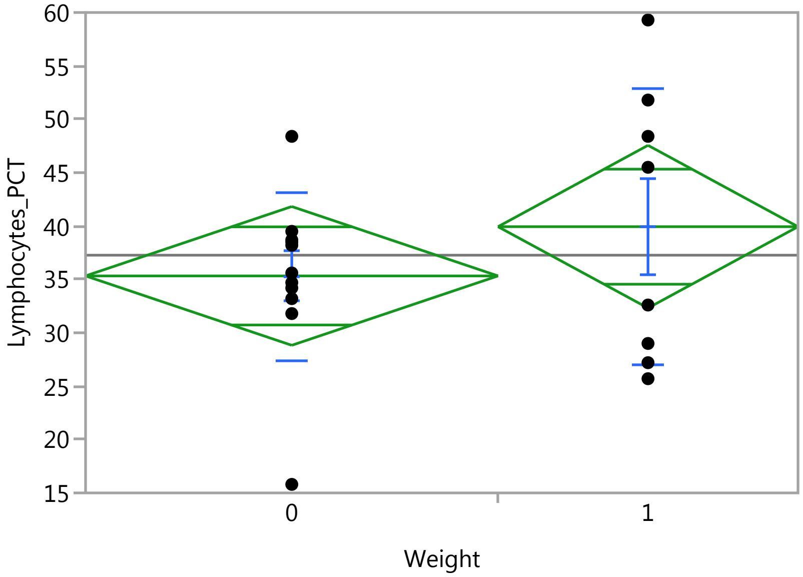


##### Oneway Anova

###### Summary of Fit

| | |
|---|---|
| Rsquare | 0.052556 |
| Adj Rsquare | -0.00318 |
| Root Mean Square Error | 10.21712 |
| Mean of Response | 37.26842 |
| Observations (or Sum Wgts) | 19 |

###### t Test

1-0

### Fit Group

#### Oneway Analysis of Lymphocytes_PCT By Weight IBS-subtype=2

##### Oneway Anova

###### t Test

Assuming equal variances

| | | | |
|---|---|---|---|
| Difference | 4.610 | t Ratio | 0.971088 |
| Std Err Dif | 4.747 | DF | 17 |
| Upper CL Dif | 14.627 | Prob > \|t\| | 0.3451 |
| Lower CL Dif | -5.406 | Prob > t | 0.1726 |
| Confidence | 0.95 | Prob < t | 0.8274 |

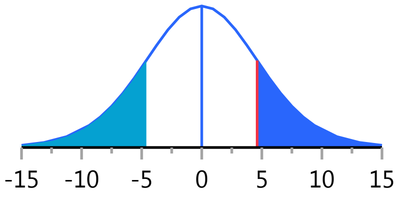


###### Analysis of Variance

| Source | DF | Sum of Squares | Mean Square | F Ratio | Prob > F |
|---|---|---|---|---|---|
| Weight | 1 | 98.4405 | 98.440 | 0.9430 | 0.3451 |
| Error | 17 | 1774.6206 | 104.389 | | |
| C. Total | 18 | 1873.0611 | | | |

###### Means for Oneway Anova

| Level | Number | Mean | Std Error | Lower 95% | Upper 95% |
|---|---|---|---|---|---|
| 0 | 11 | 35.3273 | 3.0806 | 28.828 | 41.827 |
| 1 | 8 | 39.9375 | 3.6123 | 32.316 | 47.559 |

Std Error uses a pooled estimate of error variance

##### Means and Std Deviations

| Level | Number | Mean | Std Dev | Std Err Mean | Lower 95% | Upper 95% |
|---|---|---|---|---|---|---|
| 0 | 11 | 35.3273 | 7.8614 | 2.3703 | 30.046 | 40.609 |
| 1 | 8 | 39.9375 | 12.8541 | 4.5446 | 29.191 | 50.684 |

##### t Test

1-0

Assuming unequal variances

| | | | |
|---|---|---|---|
| Difference | 4.610 | t Ratio | 0.899449 |
| Std Err Dif | 5.126 | DF | 10.7686 |
| Upper CL Dif | 15.921 | Prob > \|t\| | 0.3881 |
| Lower CL Dif | -6.701 | Prob > t | 0.1940 |
| Confidence | 0.95 | Prob < t | 0.8060 |

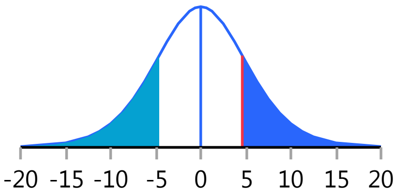


Missing Rows 1

#### Oneway Analysis of Lymphocytes By Weight IBS-subtype=2

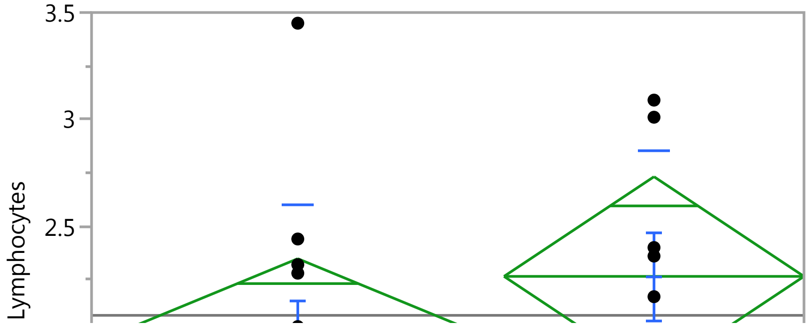

#### Fit Group

##### Oneway Analysis of Lymphocytes By Weight IBS-subtype=2

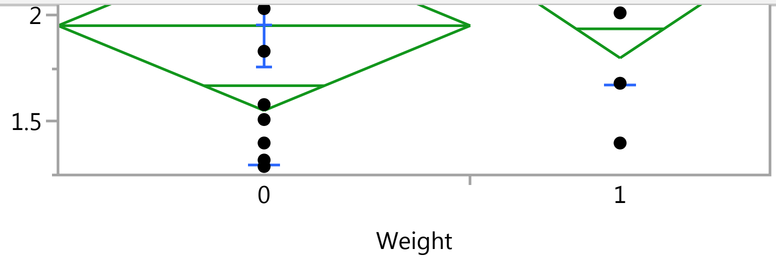


##### Oneway Anova

###### Summary of Fit

| | |
|---|---|
| Rsquare | 0.064644 |
| Adj Rsquare | 0.009624 |
| Root Mean Square Error | 0.625422 |
| Mean of Response | 2.082632 |
| Observations (or Sum Wgts) | 19 |

###### t Test

1-0

Assuming equal variances

| | | | |
|---|---|---|---|
| Difference | 0.31500 | t Ratio | 1.083931 |
| Std Err Dif | 0.29061 | DF | 17 |
| Upper CL Dif | 0.92813 | Prob > \|t\| | 0.2935 |
| Lower CL Dif | -0.29813 | Prob > t | 0.1468 |
| Confidence | 0.95 | Prob < t | 0.8532 |

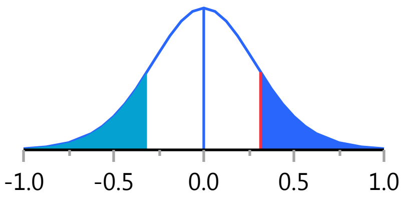


###### Analysis of Variance

| Source | DF | Sum of Squares | Mean Square | F Ratio | Prob > F |
|---|---|---|---|---|---|
| Weight | 1 | 0.4595684 | 0.459568 | 1.1749 | 0.2935 |
| Error | 17 | 6.6496000 | 0.391153 | | |
| C. Total | 18 | 7.1091684 | | | |

###### Means for Oneway Anova

| Level | Number | Mean | Std Error | Lower 95% | Upper 95% |
|---|---|---|---|---|---|
| 0 | 11 | 1.95000 | 0.18857 | 1.5521 | 2.3479 |
| 1 | 8 | 2.26500 | 0.22112 | 1.7985 | 2.7315 |

Std Error uses a pooled estimate of error variance

##### Means and Std Deviations

| Level | Number | Mean | Std Dev | Std Err Mean | Lower 95% | Upper 95% |
|---|---|---|---|---|---|---|
| 0 | 11 | 1.95000 | 0.649785 | 0.19592 | 1.5135 | 2.3865 |
| 1 | 8 | 2.26500 | 0.588873 | 0.20820 | 1.7727 | 2.7573 |

##### t Test

1-0

#### Fit Group

##### Oneway Analysis of Lymphocytes By Weight IBS-subtype=2

###### t Test

Assuming unequal variances

| | | | |
|---|---|---|---|
| Difference | 0.31500 | t Ratio | 1.101843 |
| Std Err Dif | 0.28588 | DF | 16.06701 |
| Upper CL Dif | 0.92084 | Prob > \|t\| | 0.2868 |
| Lower CL Dif | -0.29084 | Prob > t | 0.1434 |
| Confidence | 0.95 | Prob < t | 0.8566 |

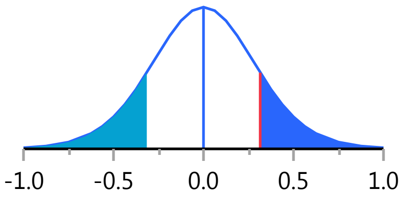


Missing Rows 1

##### Oneway Analysis of Neutrophil_PCT By Weight IBS-subtype=2

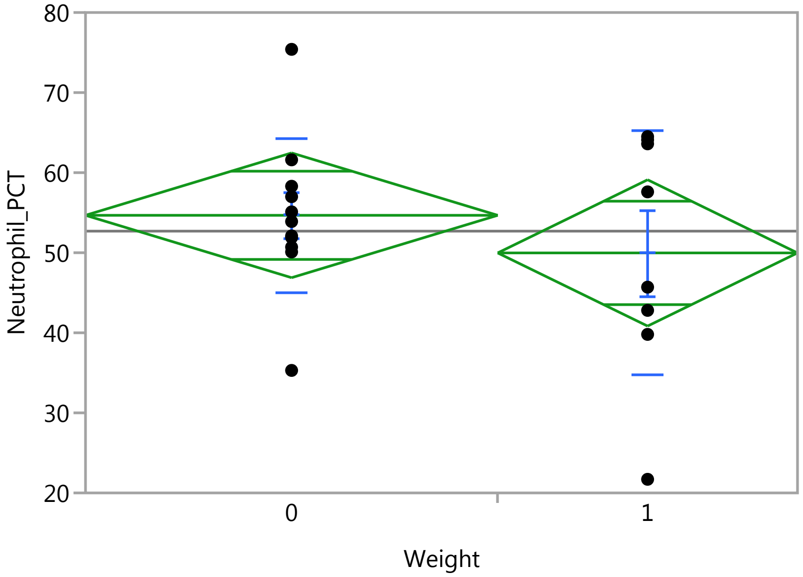


###### Oneway Anova

###### Summary of Fit

| | |
|---|---|
| Rsquare | 0.038534 |
| Adj Rsquare | -0.01802 |
| Root Mean Square Error | 12.24816 |
| Mean of Response | 52.69474 |
| Observations (or Sum Wgts) | 19 |

###### t Test

1-0

Assuming equal variances

| | | | |
|---|---|---|---|
| Difference | -4.698 | t Ratio | -0.82543 |
| Std Err Dif | 5.691 | DF | 17 |
| Upper CL Dif | 7.310 | Prob > \|t\| | 0.4206 |
| Lower CL Dif | -16.705 | Prob > t | 0.7897 |
| Confidence | 0.95 | Prob < t | 0.2103 |

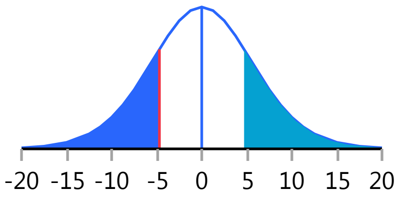

### Fit Group

#### Oneway Analysis of Neutrophil_PCT By Weight IBS-subtype=2

##### Oneway Anova

###### Analysis of Variance

| Source | DF | Sum of Squares | Mean Square | F Ratio | Prob > F |
|---|---|---|---|---|---|
| Weight | 1 | 102.2127 | 102.213 | 0.6813 | 0.4206 |
| Error | 17 | 2550.2968 | 150.017 | | |
| C. Total | 18 | 2652.5095 | | | |

###### Means for Oneway Anova

| Level | Number | Mean | Std Error | Lower 95% | Upper 95% |
|---|---|---|---|---|---|
| 0 | 11 | 54.6727 | 3.6930 | 46.881 | 62.464 |
| 1 | 8 | 49.9750 | 4.3304 | 40.839 | 59.111 |

Std Error uses a pooled estimate of error variance

##### Means and Std Deviations

| Level | Number | Mean | Std Dev | Std Err Mean | Lower 95% | Upper 95% |
|---|---|---|---|---|---|---|
| 0 | 11 | 54.6727 | 9.6087 | 2.8971 | 48.218 | 61.128 |
| 1 | 8 | 49.9750 | 15.2458 | 5.3902 | 37.229 | 62.721 |

##### t Test

1-0

Assuming unequal variances

| | | | |
|---|---|---|---|
| Difference | -4.698 | t Ratio | -0.76767 |
| Std Err Dif | 6.119 | DF | 10.98673 |
| Upper CL Dif | 8.773 | Prob > \|t\| | 0.4589 |
| Lower CL Dif | -18.168 | Prob > t | 0.7706 |
| Confidence | 0.95 | Prob < t | 0.2294 |

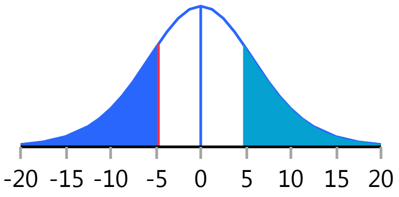


Missing Rows 1

#### Oneway Analysis of Neutrophils By Weight IBS-subtype=2

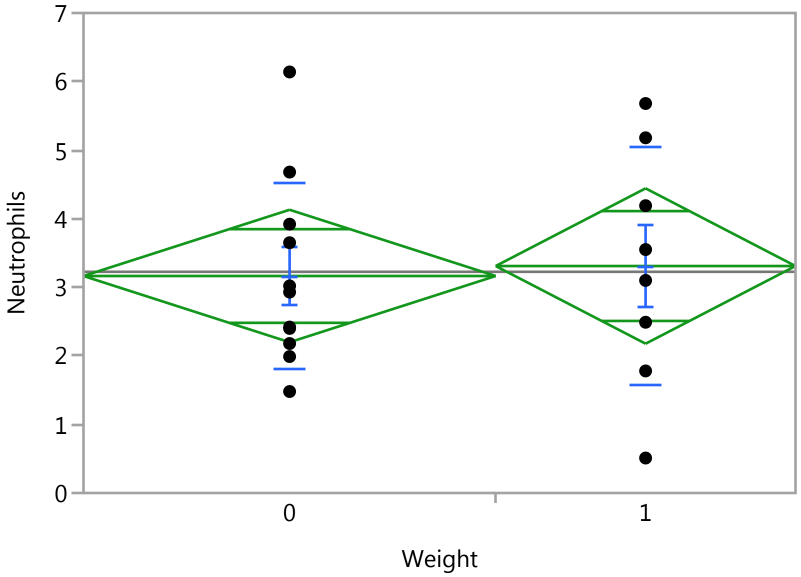

### Fit Group

#### Oneway Analysis of Neutrophils By Weight IBS-subtype=2

Weight

##### Oneway Anova

###### Summary of Fit

| | |
|---|---|
| Rsquare | 0.002493 |
| Adj Rsquare | -0.05618 |
| Root Mean Square Error | 1.51853 |
| Mean of Response | 3.225789 |
| Observations (or Sum Wgts) | 19 |

###### t Test

1-0

Assuming equal variances

| | | | |
|---|---|---|---|
| Difference | 0.1455 | t Ratio | 0.206143 |
| Std Err Dif | 0.7056 | DF | 17 |
| Upper CL Dif | 1.6341 | Prob > \|t\| | 0.8391 |
| Lower CL Dif | -1.3432 | Prob > t | 0.4196 |
| Confidence | 0.95 | Prob < t | 0.5804 |

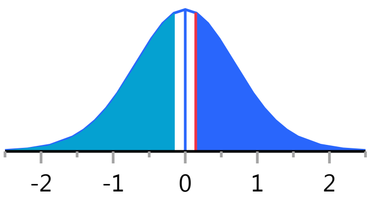


###### Analysis of Variance

| Source | DF | Sum of Squares | Mean Square | F Ratio | Prob > F |
|---|---|---|---|---|---|
| Weight | 1 | 0.097990 | 0.09799 | 0.0425 | 0.8391 |
| Error | 17 | 39.200873 | 2.30593 | | |
| C. Total | 18 | 39.298863 | | | |

###### Means for Oneway Anova

| Level | Number | Mean | Std Error | Lower 95% | Upper 95% |
|---|---|---|---|---|---|
| 0 | 11 | 3.16455 | 0.45785 | 2.1986 | 4.1305 |
| 1 | 8 | 3.31000 | 0.53688 | 2.1773 | 4.4427 |

Std Error uses a pooled estimate of error variance

##### Means and Std Deviations

| Level | Number | Mean | Std Dev | Std Err Mean | Lower 95% | Upper 95% |
|---|---|---|---|---|---|---|
| 0 | 11 | 3.16455 | 1.35490 | 0.40852 | 2.2543 | 4.0748 |
| 1 | 8 | 3.31000 | 1.72557 | 0.61008 | 1.8674 | 4.7526 |

##### t Test

1-0

Assuming unequal variances

| | | | |
|---|---|---|---|
| Difference | 0.1455 | t Ratio | 0.198106 |
| Std Err Dif | 0.7342 | DF | 12.87303 |
| Upper CL Dif | 1.7332 | Prob > \|t\| | 0.8461 |
| Lower CL Dif | -1.4423 | Prob > t | 0.4230 |
| Confidence | 0.95 | Prob < t | 0.5770 |

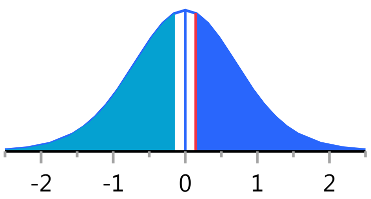


Missing Rows 1

#### Fit Group

##### Oneway Analysis of PlateletCount By Weight IBS-subtype=2

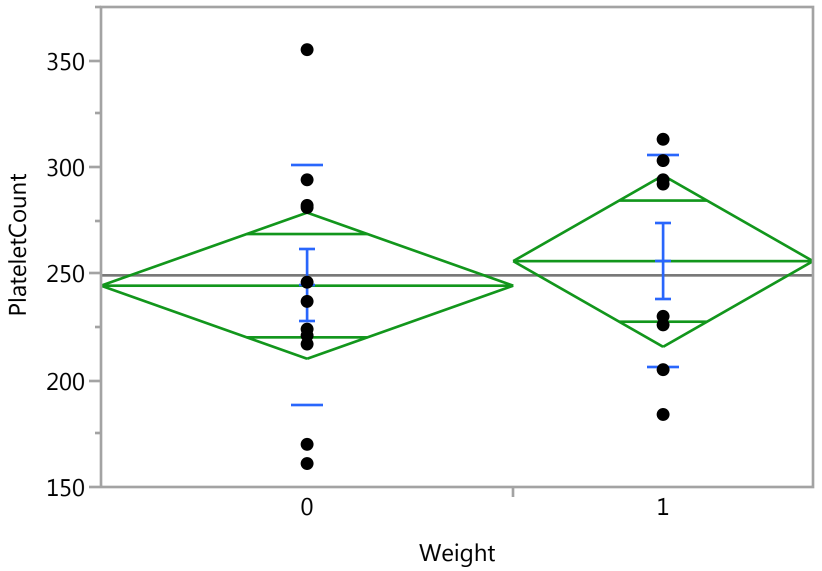


##### Oneway Anova

###### Summary of Fit

| | |
|---|---|
| Rsquare | 0.012304 |
| Adj Rsquare | -0.0458 |
| Root Mean Square Error | 53.83385 |
| Mean of Response | 249.2105 |
| Observations (or Sum Wgts) | 19 |

###### t Test

1-0

Assuming equal variances

| | | | |
|---|---|---|---|
| Difference | 11.511 | t Ratio | 0.460189 |
| Std Err Dif | 25.014 | DF | 17 |
| Upper CL Dif | 64.287 | Prob > \|t\| | 0.6512 |
| Lower CL Dif | -41.265 | Prob > t | 0.3256 |
| Confidence | 0.95 | Prob < t | 0.6744 |

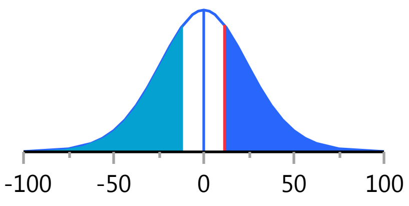


###### Analysis of Variance

| Source | DF | Sum of Squares | Mean Square | F Ratio | Prob > F |
|---|---|---|---|---|---|
| Weight | 1 | 613.737 | 613.74 | 0.2118 | 0.6512 |
| Error | 17 | 49267.420 | 2898.08 | | |
| C. Total | 18 | 49881.158 | | | |

###### Means for Oneway Anova

| Level | Number | Mean | Std Error | Lower 95% | Upper 95% |
|---|---|---|---|---|---|
| 0 | 11 | 244.364 | 16.232 | 210.12 | 278.61 |
| 1 | 8 | 255.875 | 19.033 | 215.72 | 296.03 |

Std Error uses a pooled estimate of error variance

#### Fit Group

##### Oneway Analysis of PlateletCount By Weight IBS-subtype=2

###### Means and Std Deviations

| Level | Number | Mean | Std Dev | Std Err Mean | Lower 95% | Upper 95% |
|---|---|---|---|---|---|---|
| 0 | 11 | 244.364 | 56.3103 | 16.978 | 206.53 | 282.19 |
| 1 | 8 | 255.875 | 50.0840 | 17.707 | 214.00 | 297.75 |

###### t Test

1-0

Assuming unequal variances

| | | | |
|---|---|---|---|
| Difference | 11.511 | t Ratio | 0.469242 |
| Std Err Dif | 24.532 | DF | 16.20166 |
| Upper CL Dif | 63.464 | Prob > \|t\| | 0.6451 |
| Lower CL Dif | -40.441 | Prob > t | 0.3226 |
| Confidence | 0.95 | Prob < t | 0.6774 |

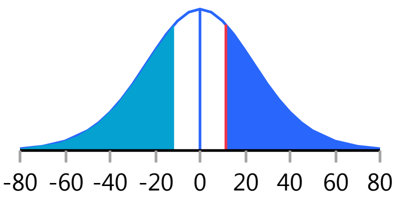


Missing Rows 1

##### Oneway Analysis of RBC By Weight IBS-subtype=2

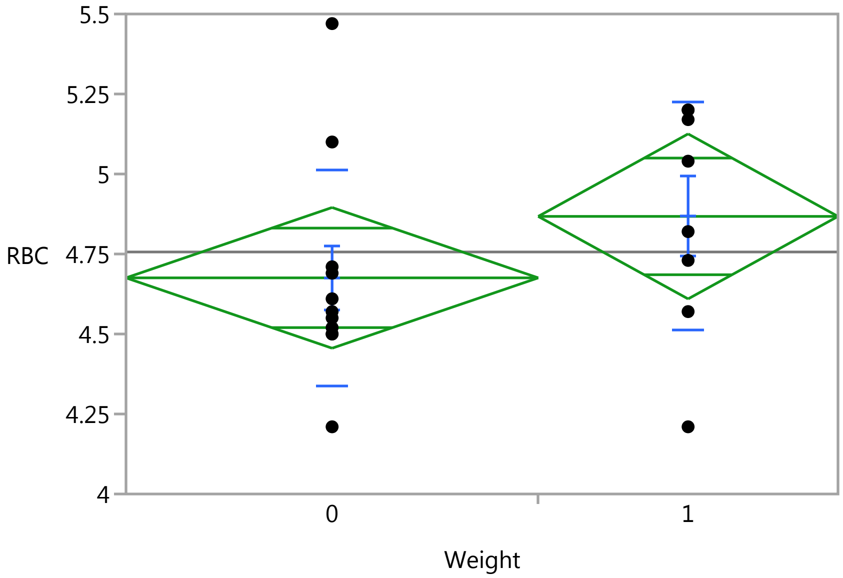


###### Oneway Anova

###### Summary of Fit

| | |
|---|---|
| Rsquare | 0.077587 |
| Adj Rsquare | 0.023328 |
| Root Mean Square Error | 0.34563 |
| Mean of Response | 4.756316 |
| Observations (or Sum Wgts) | 19 |

###### t Test

1-0

### Fit Group

#### Oneway Analysis of RBC By Weight IBS-subtype=2

##### Oneway Anova

###### t Test

Assuming equal variances

| | | | |
|---|---|---|---|
| Difference | 0.19205 | t Ratio | 1.195796 |
| Std Err Dif | 0.16060 | DF | 17 |
| Upper CL Dif | 0.53088 | Prob > \|t\| | 0.2482 |
| Lower CL Dif | -0.14679 | Prob > t | 0.1241 |
| Confidence | 0.95 | Prob < t | 0.8759 |

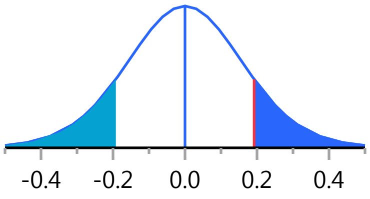


###### Analysis of Variance

| Source | DF | Sum of Squares | Mean Square | F Ratio | Prob > F |
|---|---|---|---|---|---|
| Weight | 1 | 0.1708194 | 0.170819 | 1.4299 | 0.2482 |
| Error | 17 | 2.0308227 | 0.119460 | | |
| C. Total | 18 | 2.2016421 | | | |

###### Means for Oneway Anova

| Level | Number | Mean | Std Error | Lower 95% | Upper 95% |
|---|---|---|---|---|---|
| 0 | 11 | 4.67545 | 0.10421 | 4.4556 | 4.8953 |
| 1 | 8 | 4.86750 | 0.12220 | 4.6097 | 5.1253 |

Std Error uses a pooled estimate of error variance

##### Means and Std Deviations

| Level | Number | Mean | Std Dev | Std Err Mean | Lower 95% | Upper 95% |
|---|---|---|---|---|---|---|
| 0 | 11 | 4.67545 | 0.338596 | 0.10209 | 4.4480 | 4.9029 |
| 1 | 8 | 4.86750 | 0.355437 | 0.12567 | 4.5703 | 5.1647 |

##### t Test

1-0

Assuming unequal variances

| | | | |
|---|---|---|---|
| Difference | 0.19205 | t Ratio | 1.186134 |
| Std Err Dif | 0.16191 | DF | 14.7818 |
| Upper CL Dif | 0.53759 | Prob > \|t\| | 0.2543 |
| Lower CL Dif | -0.15350 | Prob > t | 0.1271 |
| Confidence | 0.95 | Prob < t | 0.8729 |

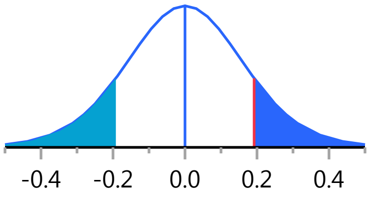


Missing Rows 1

#### Oneway Analysis of WBC By Weight IBS-subtype=2

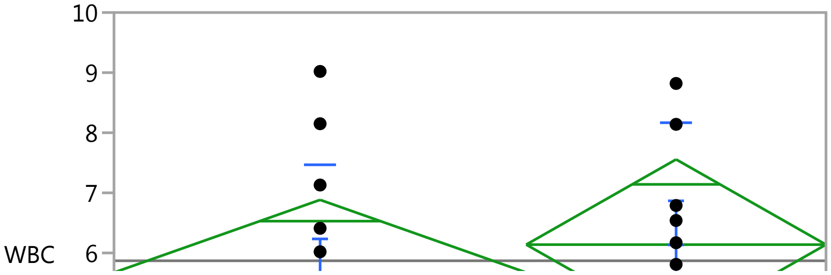

#### Fit Group

##### Oneway Analysis of WBC By Weight IBS-subtype=2

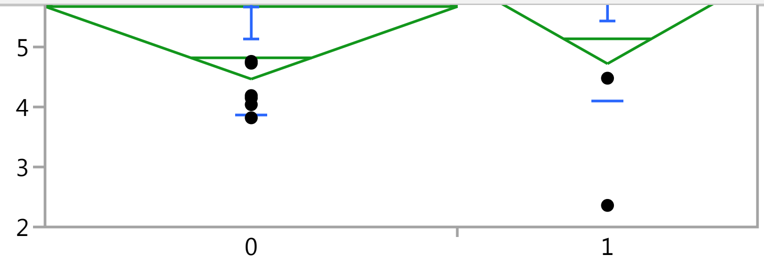


##### Oneway Anova

###### Summary of Fit

| | |
|---|---|
| Rsquare | 0.015999 |
| Adj Rsquare | -0.04188 |
| Root Mean Square Error | 1.900232 |
| Mean of Response | 5.87 |
| Observations (or Sum Wgts) | 19 |

###### t Test

1-0

Assuming equal variances

| | | | |
|---|---|---|---|
| Difference | 0.4642 | t Ratio | 0.525736 |
| Std Err Dif | 0.8830 | DF | 17 |
| Upper CL Dif | 2.3271 | Prob > \|t\| | 0.6059 |
| Lower CL Dif | -1.3987 | Prob > t | 0.3029 |
| Confidence | 0.95 | Prob < t | 0.6971 |

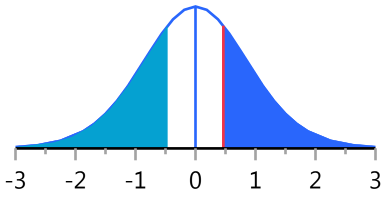


###### Analysis of Variance

| Source | DF | Sum of Squares | Mean Square | F Ratio | Prob > F |
|---|---|---|---|---|---|
| Weight | 1 | 0.998040 | 0.99804 | 0.2764 | 0.6059 |
| Error | 17 | 61.384960 | 3.61088 | | |
| C. Total | 18 | 62.383000 | | | |

###### Means for Oneway Anova

| Level | Number | Mean | Std Error | Lower 95% | Upper 95% |
|---|---|---|---|---|---|
| 0 | 11 | 5.67455 | 0.57294 | 4.4657 | 6.8833 |
| 1 | 8 | 6.13875 | 0.67183 | 4.7213 | 7.5562 |

Std Error uses a pooled estimate of error variance

##### Means and Std Deviations

| Level | Number | Mean | Std Dev | Std Err Mean | Lower 95% | Upper 95% |
|---|---|---|---|---|---|---|
| 0 | 11 | 5.67455 | 1.80184 | 0.54328 | 4.4641 | 6.8850 |
| 1 | 8 | 6.13875 | 2.03255 | 0.71861 | 4.4395 | 7.8380 |

##### t Test

1-0

### Fit Group

#### Oneway Analysis of WBC By Weight IBS-subtype=2

##### t Test

Assuming unequal variances

| | | | |
|---|---|---|---|
| Difference | 0.4642 | t Ratio | 0.515289 |
| Std Err Dif | 0.9009 | DF | 14.07077 |
| Upper CL Dif | 2.3955 | Prob > \|t\| | 0.6144 |
| Lower CL Dif | -1.4670 | Prob > t | 0.3072 |
| Confidence | 0.95 | Prob < t | 0.6928 |

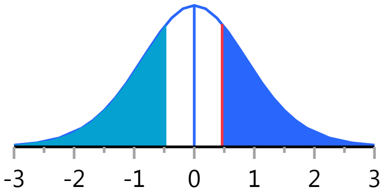


Missing Rows 1

#### Oneway Analysis of IgA By Weight IBS-subtype=2

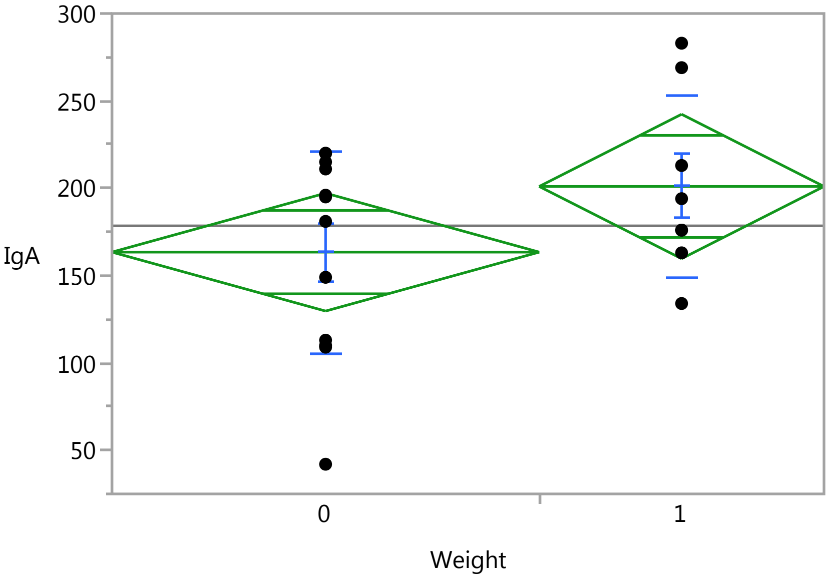


##### Oneway Anova

###### Summary of Fit

| | |
|---|---|
| Rsquare | 0.108468 |
| Adj Rsquare | 0.058939 |
| Root Mean Square Error | 55.64117 |
| Mean of Response | 178.45 |
| Observations (or Sum Wgts) | 20 |

###### t Test

1-0

Assuming equal variances

| | | | |
|---|---|---|---|
| Difference | 37.583 | t Ratio | 1.479857 |
| Std Err Dif | 25.397 | DF | 18 |
| Upper CL Dif | 90.940 | Prob > \|t\| | 0.1562 |
| Lower CL Dif | -15.773 | Prob > t | 0.0781 |
| Confidence | 0.95 | Prob < t | 0.9219 |

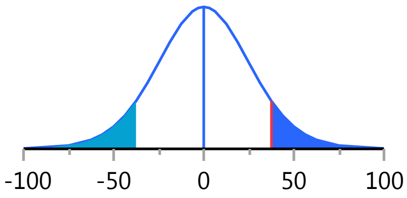

### Fit Group

#### Oneway Analysis of IgA By Weight IBS-subtype=2

##### Oneway Anova

###### Analysis of Variance

| Source | DF | Sum of Squares | Mean Square | F Ratio | Prob > F |
|---|---|---|---|---|---|
| Weight | 1 | 6780.033 | 6780.03 | 2.1900 | 0.1562 |
| Error | 18 | 55726.917 | 3095.94 | | |
| C. Total | 19 | 62506.950 | | | |

###### Means for Oneway Anova

| Level | Number | Mean | Std Error | Lower 95% | Upper 95% |
|---|---|---|---|---|---|
| 0 | 12 | 163.417 | 16.062 | 129.67 | 197.16 |
| 1 | 8 | 201.000 | 19.672 | 159.67 | 242.33 |

Std Error uses a pooled estimate of error variance

##### Means and Std Deviations

| Level | Number | Mean | Std Dev | Std Err Mean | Lower 95% | Upper 95% |
|---|---|---|---|---|---|---|
| 0 | 12 | 163.417 | 57.9992 | 16.743 | 126.57 | 200.27 |
| 1 | 8 | 201.000 | 51.7190 | 18.285 | 157.76 | 244.24 |

##### t Test

1-0

Assuming unequal variances

| | | | |
|---|---|---|---|
| Difference | 37.583 | t Ratio | 1.515898 |
| Std Err Dif | 24.793 | DF | 16.3462 |
| Upper CL Dif | 90.051 | Prob > \|t\| | 0.1486 |
| Lower CL Dif | -14.885 | Prob > t | 0.0743 |
| Confidence | 0.95 | Prob < t | 0.9257 |

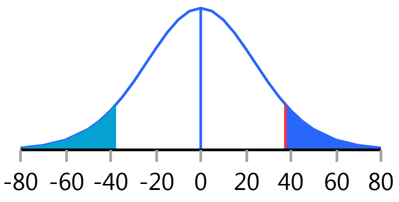


#### Oneway Analysis of IgE By Weight IBS-subtype=2

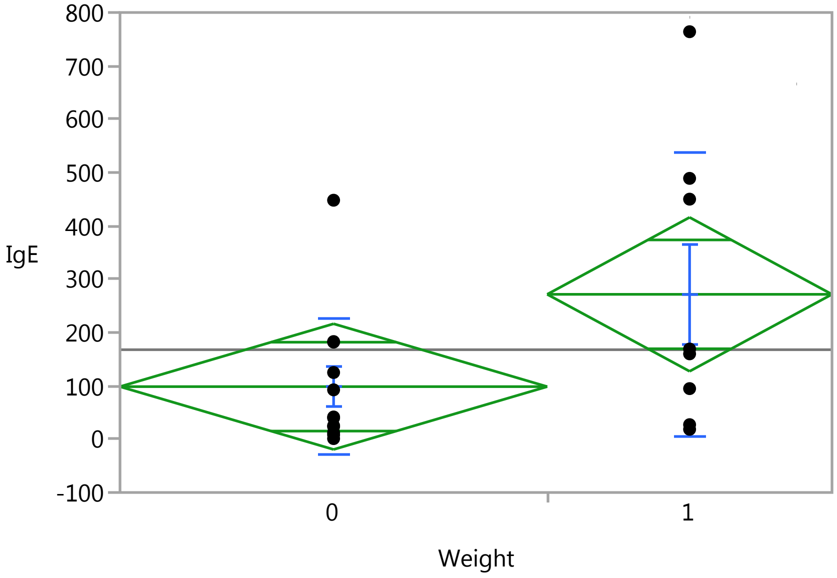

### Fit Group

#### Oneway Analysis of IgE By Weight IBS-subtype=2

Weight

##### Oneway Anova

###### Summary of Fit

| | |
|---|---|
| Rsquare | 0.174653 |
| Adj Rsquare | 0.1288 |
| Root Mean Square Error | 194.2753 |
| Mean of Response | 167.725 |
| Observations (or Sum Wgts) | 20 |

###### t Test

1-0

Assuming equal variances

| | | | |
|---|---|---|---|
| Difference | 173.06 | t Ratio | 1.951668 |
| Std Err Dif | 88.67 | DF | 18 |
| Upper CL Dif | 359.36 | Prob > \|t\| | 0.0667 |
| Lower CL Dif | -13.23 | Prob > t | 0.0334* |
| Confidence | 0.95 | Prob < t | 0.9666 |

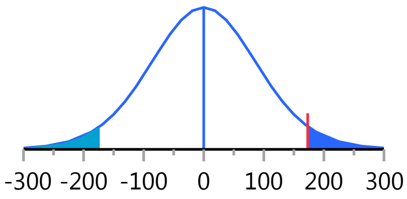


###### Analysis of Variance

| Source | DF | Sum of Squares | Mean Square | F Ratio | Prob > F |
|---|---|---|---|---|---|
| Weight | 1 | 143763.02 | 143763 | 3.8090 | 0.0667 |
| Error | 18 | 679372.12 | 37743 | | |
| C. Total | 19 | 823135.14 | | | |

###### Means for Oneway Anova

| Level | Number | Mean | Std Error | Lower 95% | Upper 95% |
|---|---|---|---|---|---|
| 0 | 12 | 98.500 | 56.082 | -19.3 | 216.32 |
| 1 | 8 | 271.563 | 68.687 | 127.3 | 415.87 |

Std Error uses a pooled estimate of error variance

##### Means and Std Deviations

| Level | Number | Mean | Std Dev | Std Err Mean | Lower 95% | Upper 95% |
|---|---|---|---|---|---|---|
| 0 | 12 | 98.500 | 127.847 | 36.906 | 17.270 | 179.73 |
| 1 | 8 | 271.563 | 267.148 | 94.451 | 48.221 | 494.90 |

##### t Test

1-0

Assuming unequal variances

| | | | |
|---|---|---|---|
| Difference | 173.06 | t Ratio | 1.706635 |
| Std Err Dif | 101.41 | DF | 9.164772 |
| Upper CL Dif | 401.83 | Prob > \|t\| | 0.1215 |
| Lower CL Dif | -55.71 | Prob > t | 0.0607 |
| Confidence | 0.95 | Prob < t | 0.9393 |

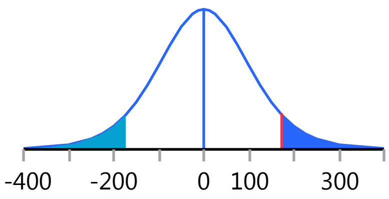

#### Fit Group

##### Oneway Analysis of IgG By Weight IBS-subtype=2

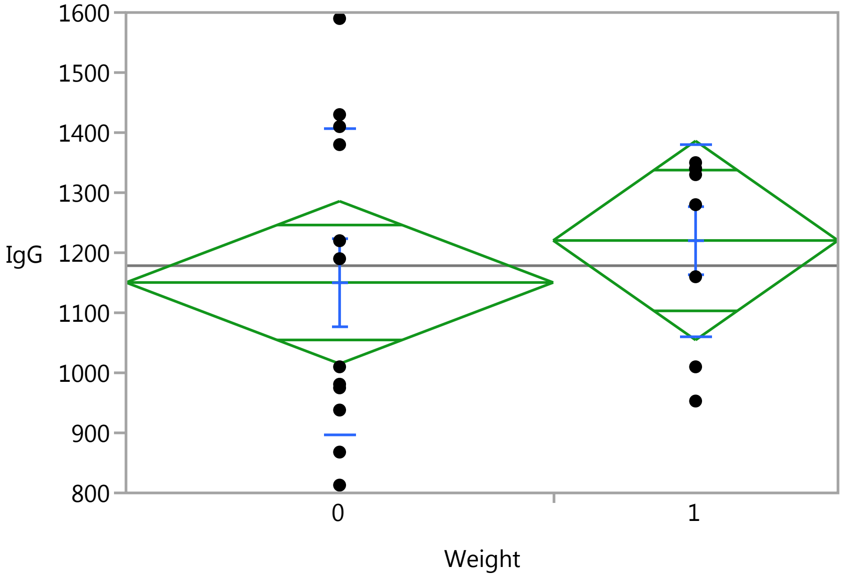


##### Oneway Anova

###### Summary of Fit

| | |
|---|---|
| Rsquare | 0.025543 |
| Adj Rsquare | -0.02859 |
| Root Mean Square Error | 223.1353 |
| Mean of Response | 1178.4 |
| Observations (or Sum Wgts) | 20 |

###### t Test

1-0

Assuming equal variances

| | | | |
|---|---|---|---|
| Difference | 69.96 | t Ratio | 0.686897 |
| Std Err Dif | 101.85 | DF | 18 |
| Upper CL Dif | 283.93 | Prob > \|t\| | 0.5009 |
| Lower CL Dif | -144.01 | Prob > t | 0.2505 |
| Confidence | 0.95 | Prob < t | 0.7495 |

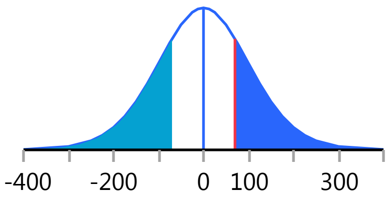


###### Analysis of Variance

| Source | DF | Sum of Squares | Mean Square | F Ratio | Prob > F |
|---|---|---|---|---|---|
| Weight | 1 | 23492.01 | 23492.0 | 0.4718 | 0.5009 |
| Error | 18 | 896208.79 | 49789.4 | | |
| C. Total | 19 | 919700.80 | | | |

###### Means for Oneway Anova

| Level | Number | Mean | Std Error | Lower 95% | Upper 95% |
|---|---|---|---|---|---|
| 0 | 12 | 1150.42 | 64.414 | 1015.1 | 1285.7 |
| 1 | 8 | 1220.38 | 78.890 | 1054.6 | 1386.1 |

Std Error uses a pooled estimate of error variance

### Fit Group

#### Oneway Analysis of IgG By Weight IBS-subtype=2

##### Means and Std Deviations

| Level | Number | Mean | Std Dev | Std Err Mean | Lower 95% | Upper 95% |
|---|---|---|---|---|---|---|
| 0 | 12 | 1150.42 | 255.097 | 73.640 | 988.3 | 1312.5 |
| 1 | 8 | 1220.38 | 160.529 | 56.756 | 1086.2 | 1354.6 |

##### t Test

1-0

Assuming unequal variances

| | | | |
|---|---|---|---|
| Difference | 69.96 | t Ratio | 0.752453 |
| Std Err Dif | 92.97 | DF | 17.98005 |
| Upper CL Dif | 265.30 | Prob > \|t\| | 0.4615 |
| Lower CL Dif | -125.39 | Prob > t | 0.2308 |
| Confidence | 0.95 | Prob < t | 0.7692 |

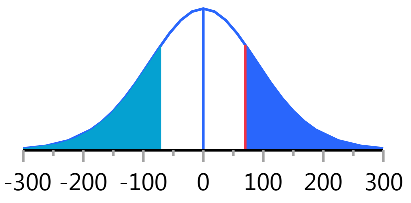


#### Oneway Analysis of IgM By Weight IBS-subtype=2

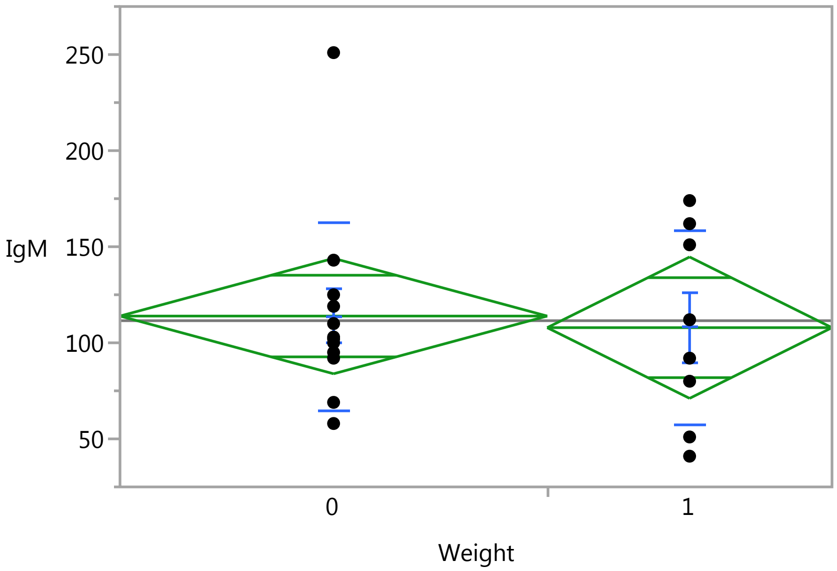


##### Oneway Anova

###### Summary of Fit

| | |
|---|---|
| Rsquare | 0.003952 |
| Adj Rsquare | -0.05138 |
| Root Mean Square Error | 49.52765 |
| Mean of Response | 111.5 |
| Observations (or Sum Wgts) | 20 |

###### t Test

1-0

### Fit Group

#### Oneway Analysis of IgM By Weight IBS-subtype=2

##### Oneway Anova

###### t Test

Assuming equal variances

| | | | |
|---|---|---|---|
| Difference | -6.042 | t Ratio | -0.26726 |
| Std Err Dif | 22.606 | DF | 18 |
| Upper CL Dif | 41.452 | Prob > \|t\| | 0.7923 |
| Lower CL Dif | -53.535 | Prob > t | 0.6038 |
| Confidence | 0.95 | Prob < t | 0.3962 |

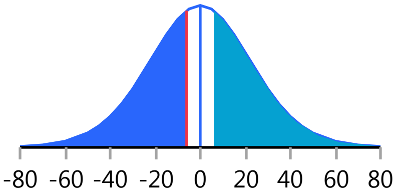


###### Analysis of Variance

| Source | DF | Sum of Squares | Mean Square | F Ratio | Prob > F |
|---|---|---|---|---|---|
| Weight | 1 | 175.208 | 175.21 | 0.0714 | 0.7923 |
| Error | 18 | 44153.792 | 2452.99 | | |
| C. Total | 19 | 44329.000 | | | |

###### Means for Oneway Anova

| Level | Number | Mean | Std Error | Lower 95% | Upper 95% |
|---|---|---|---|---|---|
| 0 | 12 | 113.917 | 14.297 | 83.879 | 143.95 |
| 1 | 8 | 107.875 | 17.511 | 71.086 | 144.66 |

Std Error uses a pooled estimate of error variance

##### Means and Std Deviations

| Level | Number | Mean | Std Dev | Std Err Mean | Lower 95% | Upper 95% |
|---|---|---|---|---|---|---|
| 0 | 12 | 113.917 | 48.8401 | 14.099 | 82.885 | 144.95 |
| 1 | 8 | 107.875 | 50.5892 | 17.886 | 65.581 | 150.17 |

##### t Test

1-0

Assuming unequal variances

| | | | |
|---|---|---|---|
| Difference | -6.042 | t Ratio | -0.26528 |
| Std Err Dif | 22.775 | DF | 14.77227 |
| Upper CL Dif | 42.567 | Prob > \|t\| | 0.7945 |
| Lower CL Dif | -54.650 | Prob > t | 0.6028 |
| Confidence | 0.95 | Prob < t | 0.3972 |

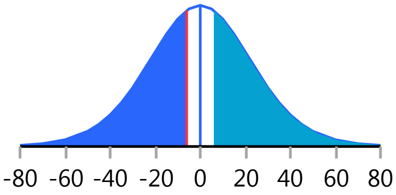
